# Spatial Proximity Moderates Genotype Uncertainty in Genetic Tagging Studies

**DOI:** 10.1101/2020.01.01.892463

**Authors:** Ben C. Augustine, J. Andrew Royle, Daniel W. Linden, Angela K. Fuller

**Affiliations:** Atkinson Center for a Sustainable Future and Department of Natural Resources, Cornell University, Ithaca, NY 14843; U.S. Geological Survey, Patuxent Wildlife Research Center, Laurel, MD, 20708; NOAA National Marine Fisheries Service, Gloucester, MA, 01930; U.S. Geological Survey, New York Cooperative Fish and Wildlife Research Unit, Department of Natural Resources, Cornell University, Ithaca, NY 14843

**Keywords:** Spatial capture-recapture, partial identity, spatial partial identity, genetic capture-recapture, microsatellite, capture-recapture

## Abstract

Accelerating declines of an increasing number of animal populations worldwide necessitate methods to reliably and efficiently estimate demographic parameters such as population density and trajectory. Standard methods for estimating demographic parameters from noninvasive genetic samples are inefficient because lower quality samples cannot be used, and they do not allow for errors in individual identification. We introduce the Genotype Spatial Partial Identity Model (SPIM), which integrates a genetic classification model with a spatial population model to combine both spatial and genetic information, thus reducing genotype uncertainty and increasing the precision of demographic parameter estimates. We apply this model to data from a study of fishers (*Pekania pennanti*) in which 37% of samples were originally discarded because of uncertainty in individual identity. The Genotype SPIM density estimate using all collected samples was 25% more precise than the original density estimate, and the model identified and corrected 3 errors in the original individual identity assignments. A simulation study demonstrated that our model increased the accuracy and precision of density estimates 63% and 42%, respectively, using 3 PCRs per genetic sample. Further, the simulations showed that the Genotype SPIM model parameters are identifiable with only one PCR per sample, and that accuracy and precision are relatively insensitive to the number of PCRs for high quality samples. Current genotyping protocols devote the majority of resources to replicating and confirming high quality samples, but when using the Genotype SPIM, genotyping protocols could be more efficient by devoting more resources to low quality samples.

**Significance:** We present a new statistical framework for the estimation of animal demographic parameters, such as abundance, density, and growth rate, from noninvasive genetic samples (e.g., hair, scat). By integrating a genetic classification model with a spatial population model, we show that accounting for spatial proximity of samples reduces genotype uncertainty and improves parameter estimation. Our method produces a fundamentally different approach to genetic capture-recapture by sharing information between the normally disjunct steps of assigning individual identities to genetic samples and modeling population processes. Further, it leads to more efficient protocols for processing genetic samples, which can lower project costs and expand opportunities for applying noninvasive genetics to conservation and management problems.

## Introduction

Species extinction risk is tied to the loss of individual populations, with recent studies demonstrating range contractions of 94-99% in some of the world’s large carnivores (Wolf and Ripple 2017). Accelerating declines of an increasing number of animal populations worldwide necessitate methods to reliably and efficiently estimate demographic parameters such as population size, and vital rates such as survival probability, recruitment rate, and the population trajectory through time. Unfortunately, many species of conservation concern are managed without having the necessary information on population status or trends, which is largely a consequence of the cost and difficulty of studying species in decline and the difficulty of applying statistical models to sparse data, which can produce imprecise and biased estimates of demographic parameters.

Noninvasive genetic monitoring has become an invaluable tool for estimating population parameters and quantifying population status because genetic samples are efficient to collect for a large number of species (Lamb et al. 2019). The DNA contained in noninvasive samples, such as tissue, hair, or scat, can be used to extract microsatellite markers (Taberlet et al. 1996), or more recently, single nucleotide polymorphisms (SNPs; Natesh et al. 2019), which serve as the basis for estimating population genetics or population dynamics parameters. The role of genetic markers in genetic capture-recapture studies is to provide individual identities for the collected samples, which are then used to construct capture histories required by capture-recapture models, or more recently, spatial capture-recapture models (SCR; Borchers and Efford 2008; Royle et al. 2013). Individual identities are constructed by observing the combination of allele values at enough genetic loci that it is improbable that multiple individuals in the collected sample share the same multilocus genotype (the “shadow effect”; Taberlet et al. 1996).

Two key challenges in the application of noninvasive genetic sampling are that genetic markers from noninvasive samples are observed with error and the markers may provide insufficient power to discriminate between individuals, particularly when not all loci amplify successfully. These challenges are especially problematic when using observed multilocus genotypes to establish individual identities for the use in capture-recapture models because errors in assigning individual identities can severely bias parameter estimates(Mills et al. 2000; Creel et al. 2003; Lukacs and Burnham 2005). The most widely applied solution to these problems involves a process of data curation and filtering aimed at identifying only the highest certainty samples and discarding the remainder (Lampa et al. 2013; Sethi et al. 2014).

The genotyping process involves (1) DNA extraction and amplification via polymerase chain reaction (PCR) (Waits and Paetkau 2005), and (2) decision making and analysis by an expert to interpret genetic samples and assign them multilocus genotypes and individual identities. In practice, these are regarded to be error-free, reorganized into capture histories, and then used in capture-recapture models to estimate animal demographic parameters. In practice, DNA extraction and amplification are prone to errors, especially for noninvasive samples that typically contain a low quantity and quality of DNA (Taberlet et al. 1996; Lampa et al. 2013). Both shadow effects and genotyping errors lead to incorrect assignment of individual identities to samples, and error rates as low as 1-5% introduce strong bias into population parameter estimates using typical capture-recapture models (Mills et al. 2000; Creel et al. 2003; Lukacs and Burnham 2005) that strictly assume that all samples are identified to individual correctly (Otis et al. 1978). In addition, because only a single capture history is produced, based on the “consensus genotype” of each sample, uncertainty inherent to the decision making process of assigning individual identities to samples is not propagated through to the inferences.

The extreme sensitivity of capture-recapture models to even small error rates in individual identity (Mills et al. 2000; Lukacs and Burnham 2005) has largely determined the structure of genotyping protocols used for genetic capture-recapture to date. Genetic markers were established as a reliable tool for capture-recapture analyses by the development of standardized lab protocols that minimized genotyping errors (Taberlet et al. 1996; Paetkau 2003; Waits and Paetkau 2005) and statistical tools that aid in determining the number of required markers and identifying the reliable samples (e.g., Waits et al. 2001; Miller et al. 2002). These tools were originally developed for microsatellite markers, which we focus on here; however, much of our discussion also applies to SNPs or other genetic markers. Genotyping protocols vary across studies, but common features used to reduce errors in assigning individual identities are 1) the use of enough highly variable loci to minimize shadow events, 2) some form of replicate genotyping to identify and limit genotyping errors, and 3) the removal of samples judged to be unreliable. We will briefly describe this “traditional approach” to assigning individual identities to samples–see Lampa et al. (2013) and Sethi et al. (2014) for comprehensive reviews.

First, shadow events are minimized by using a marker set with sufficient discriminatory power. The main statistic used to measure the discriminatory power of a marker set is *P*_*ID*_, which quantifies the probability that two randomly selected individuals in the population will have the same genotype by chance, given the number of loci and estimated allele frequencies (Paetkau et al. 1998). In practice, the more conservative *P*_*IDsib*_, the probability two randomly selected *siblings* in the population will have the same genotype by chance (Waits et al. 2001), is typically used. For simplicity, we will refer to both of these statistics as *P*_*ID*_. Generally, a low *P*_*ID*_ threshold is used to determine how many markers to use, and this threshold varies widely across studies–Lampa et al. (2013) documented studies with *P*_*ID*_ thresholds spanning 7 orders of magnitude (8.2 x 10^−4^ to 2.7^−11^). One factor partially accounting for this variability is that, in order to limit the absolute number of shadow events, the *P*_*ID*_ threshold must scale with the number of individuals captured (Paetkau 2003), a product of the population size and individual capture probability. Unfortunately, the population size, capture probability, and number of individuals captured are quantities to be estimated and are by definition unknown, or imprecisely known, in advance. Given this and other sources of uncertainty, *P*_*ID*_ thresholds are typically chosen to be overly conservative, which can lead to the culling of large numbers of samples that do not amplify at all loci or otherwise cannot be scored at all loci due to potential genotyping errors. We refer to these samples as *partial genotypes*.

Genotyping errors are then minimized by using some form of replication of the genotyping process from which *consensus genotypes* are constructed. Generally, the most rigorous (and expensive) method for generating consensus genotypes from low quality DNA samples is the “multitubes” approach (Taberlet et al. 1996; Miller et al. 2002) where the DNA product from each sample is split across multiple subsamples and then amplified and scored independently. The consensus genotype is then determined by comparing the scores across replications and only scoring a locus if the same single-locus genotype is seen a minimum number of times across replicates (Taberlet et al. 1996). This minimum for reaching consensus varies somewhat subjectively across studies and by zygosity (homozygotes vs. heterozygotes; Lampa et al. 2013). Less comprehensive and more efficient protocols for generating consensus genotypes are also in use, where only samples suspected of containing errors are replicated (Paetkau 2003; Schwartz et al. 2006). After consensus genotypes are generated, samples are matched to individual using one of a number of algorithms (e.g., Creel et al. 2003; Macbeth et al. 2011), while discarding samples whose consensus genotypes are missing too many locus scores due to failed amplification or insufficient genotype confirmation.

This typical approach to producing individual identities from replicated genotype scores (the ‘*individual identity observation process*’) can be conceptualized as a random thinning process (depicted in Figure I) where the true capture history, ***Y***^*true*^ is split into a capture history of known identity samples, ***Y***^*ID*^, and a vector of trap-level or a matrix of trap by occasion-level counts, ***Y***^*unk*^, which is discarded. A possible thinning process for an individual by trap capture history is 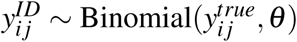, where the *θ* parameter determines the probability that a sample can be identified to individual. *θ* is then a function of the overall quantity and quality of DNA in the samples, but also, the level of conservativeness used for accepting samples as reliable. For the same set of samples, a more conservative genotyping protocol will raise *θ*, leading to fewer individual identity errors in ***Y***^*ID*^ at the cost of discarding more samples. This trade-off cannot be avoided if no errors are allowed in individual identity. One further thing to note is that the individual identity observation process is only partially connected to the ecological and capture processes–the information in the data associated with density, the detection parameters, and the spatial location of samples is not used to assign individual identities to the samples, despite the fact that these information sources contain abundant information about the true genotypes and their individual identities (Augustine et al. 2019). In fact, the spatial location where a sample was collected constitutes a continuous partial identity, observed with error, where the true state for each individual is its center of activity during a survey and the dispersion of samples around an individual’s activity center is a function of its home range size and thus, spatial scale of detection (Augustine et al. 2019). Therefore, the spatial locations where samples are collected are analogous to an additional genetic marker, but to date, this information has not been used as part of the genotyping process. All of these limitations can be resolved with an appropriately structured capture-recapture model that allows for errors and uncertainty in individual identity. Recently, capture-recapture models that allow for various types of errors and uncertainty in individual identity have been developed (Lukacs and Burnham 2005; Link et al. 2010; Wright et al. 2009; Augustine et al. 2019; Knapp et al. 2009), though to date, no comprehensive model exists that includes all the features relevant to the genotyping process.

Here, we present the Genotype Spatial Partial Identity Model (SPIM)–a single probabilistic framework which removes the process of subjective processing and interpretation of genetic data from the “traditional genotyping process”. Our approach combines an explicit model of genetic and individual classification with a model of spatially-explicit individual encounter histories for making inference about animal population parameters. This model, with clearly articulated probability assumptions about each component of the system, allows for uncertainty to be propagated among the ecological, capture, and genotyping processes, uses all available sources of information, and removes the need for data culling. The net result is increased efficiency in noninvasive genetic capture-recapture studies by making use of all available data and the concomitant improvement in statistical accuracy and precision of estimates of population parameters. Our model accounts for the shadow effect and genotyping errors–both allelic dropout and false alleles. Further, we use the spatial locations where samples were collected to reduce uncertainty in probabilistically assigning their individual identities. We apply this model to a previously analyzed data set of fishers (*Pekania pennanti*) in New York, USA (Linden et al. 2017), making use of the 37% of samples that were originally discarded due to uncertain individual identity (Figure 1). We then investigate the performance of the model via simulation.

**Figure 1:**
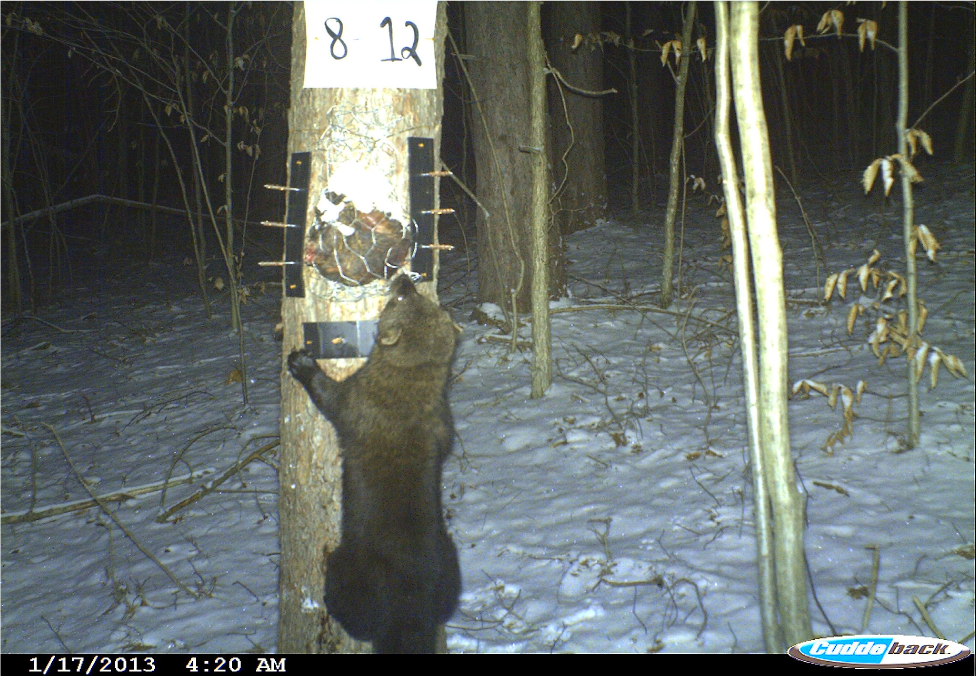
A fisher at a baited hair snare.

## Methods – Model Description

The Genotype SPIM is a 3-level ecological hierarchical model (Royle and Dorazio 2008, see Figure I), with the first two levels being models for the ecological and capture processes of the same general structure as the catSPIM. We use a joint ecological process model, the first component of which describes the number and distribution of individuals across a two-dimensional state space, 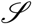, by use of a spatial point process in which realized point locations ***S***_*N*×2_ represent each individual’s mean location (activity center) during the survey. We assume activity centers are distributed uniformly – ***s***_*i*_ ∼ Uniform(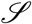), *i* = 1, *…, N*, though other models could be used. The second process model component describes each individual’s multilocus genotype, which we define to be an individual’s true values at *n*^*cat*^ genetic loci. Each loci *l* has 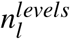 possible single locus genotypes, *l* = 1, *…, n*^*cat*^, which are enumerated 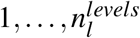 for each *l*. Associated with each loci *l* are single locus genotype frequencies ***γ***_*l*_–the probabilities with which each single locus genotype occurs in the population. The multilocus genotypes of all individuals are organized into the *N* × *n*^*cat*^ matrix ***G***^*true*^, with rows of ***G***^*true*^ corresponding to the same individuals as the rows of ***S***. We assume each element of ***G***^*true*^ is independent of the others: 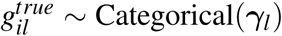. Together, these models for the activity centers and genotypes provides a spatially-explicit description for the distribution of genotypes across space. Note that this genotype distribution model allows multiple individuals in the population to have the same multilocus genotype (shadow effect), but they will not share a spatial location.

The capture process model also has two components. The first component describes how animals are detected during *K* capture occasions, conditional on their activity center locations. We assume individuals are detected at specific point locations in the state space (i.e. “traps”), ***X***_*J*×2_ = {***x*** _*j*_; *j* = 1, 2, *…, J*}. The capture data are organized in ***Y***^*true*^, recording the number of counts for each individual at each trap summed across capture occasions. This data structure is given a superscript “true” to distinguish it from the observed capture data, described below–in the presence of genotyping error and/or shadow events, the true encounter histories are latent variables. We assume the individual by trap detection data arise following a detection function, describing the trap by occasion-level detection probability or rate as a function of distance from an individual activity center. We further assume the individual by trap by occasion detection process is Poisson, where the baseline detection rate of individual *i* at trap *j* is 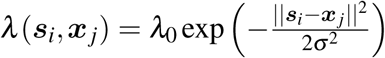, and 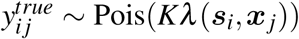. The observed capture data are defined to be ***Y*** ^*obs*^, an *n*^*obs*^ × *J* matrix, where *n*^*obs*^ is the sum of all observed counts. Each row of ***Y*** ^*obs*^ corresponds to one count member (e.g., a count of 3 has 3 count members that cannot be deterministically linked when individual identity is unknown), with all zero entries, except for a single 1 indicating the trap of capture. The second component of the capture process model describes how the genetic loci are observed or “captured” conditional on the true multilocus genotypes. In a model without observation error (e.g., catSPIM), the observed loci reflect their true values or are recorded as missing data.

We now consider that the loci-level genotypes may not always be observed correctly using an explicit genotype observation process (see Figure I). Let *n*^*rep*^ be the maximum number of times each of the *n*^*obs*^ samples are observed (e.g., number of replicate PCRs), with the observed scores recorded in the *n*^*obs*^ × *n*^*cat*^ × *n*^*rep*^ array *G*^*obs*^ for the loci that amplify and recorded as missing data for loci that do not amplify or for replication numbers of samples with fewer than the maximum number of replicates. We allow for 3 observation events–correct observation, allelic dropout (heterozygote observed as homozygote), and false allele (any other error). We use a simple model for these genotype observation events that assumes that each possible allelic dropout event is equally likely (previously assumed by Wright et al. 2009; Sethi et al. 2016), each possible false allele event is equally likely (previously assumed by Sethi et al. 2016), and the allelic dropout and false allele probabilities do not vary across sample (relaxed below), locus, individual, or replicate number. We define ***p***^*hom*^ and ***p***^*het*^ to be vectors of the observation probabilities for homozygous and heterozygous loci-level genotypes, respectively. Then, 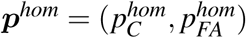 for homozygous correct and false allele observation and 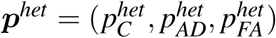 for heterozygous correct, allelic dropout, and false allele observation. These observation probabilities then make up the elements of ***pi***_*l*_,the observation probability matrix for locus *l*, conditional on the true locus-level genotypes. Details of how these matrices are constructed along with other technical details of the model can be found in Appendix A.

## Methods – Fisher Application

We applied the Genotype SPIM to a hair snare data set from fisher surveyed in New York, USA, collected in 2014. The full details of this study can be found in Linden et al. (2017). Four hundred and twenty hair samples were collected at 608 traps, each operated for 3 1-week occasions. Each sample was amplified 1 to 7 times across 9 microsatellite loci. In the original study, 263 samples were assigned individual identities using the methods of Creel et al. (2003) with a *P*_*ID*−*sib*_ criterion of 0.005, though some additional matches were made using the methods of Macbeth et al. (2011). The 263 samples were assigned to 189 distinct individuals. One hundred fifty-seven samples (37%) were discarded because they produced partial genotypes not informative enough to be confidently assigned individual identities (105 samples) or they did not amplify at all (52 samples). Among the individually identified samples in the original study, there were 74 recaptures from 50 individuals and 9 spatial recaptures across 8 individuals. The low rate of spatial recaptures was largely due to a survey that was primarily designed for an occupancy analysis aiming for independent trap sites.

We applied the Genotype SPIM to all 420 collected samples and compared it to the estimates from the regular SCR analysis using the 263 samples originally assigned certain individual identities using traditional methods. For both the Genotype SPIM and SCR analyses, we used the individual heterogeneity model for the detection function parameters described in Appendix A, because the distribution of spatial recapture numbers and distances suggested strong individual heterogeneity in space use. We allowed genotyping error rates to vary by two sample quality categories as described in Appendix A. We defined “high quality” samples to be those that amplified at an average of ≥8 of 9 loci across the first 3 replication attempts and “low quality” samples as those amplifying at an average of *<*8 loci across the same 3 replication attempts. This criterion is somewhat arbitrary, but it is more realistic than assuming all samples have the same genotyping error probabilities and is consistent with the fact that samples with less DNA product have both higher genotyping error rates and lower loci-level amplification rates (Taberlet et al. 1996; McKelvey and Schwartz 2004). To compare the precision of selected parameter estimates between the Genotype SPIM and SCR analyses, we used the coefficient of variation (CV)–the posterior standard deviation divided by the posterior mode. See Appendix B for the MCMC specifications for this analysis.

### Methods – Simulation Study

We conducted a small simulation study motivated by the fisher analysis with a large proportion of originally discarded samples to demonstrate the performance of the Genotype SPIM in general (e.g., bias and coverage) and to compare 1) the regular SCR estimate using only high quality samples to the Genotype SPIM estimates using all samples, 2) the Genotype SPIM estimates with 1-3 PCRs per sample and 3) the Genotype SPIM estimates using or discarding the low quality samples. The simulation study was designed to replicate the design that produced the fisher data set with a few caveats. Specifically, we did not consider the individual heterogeneity model for the detection function parameters to reduce computation time, we used a smaller trapping array with traps spaced optimally for SCR to better reflect the typical resources available for survey effort, and we considered a higher population density so that the scenario is more challenging for uncertain identity methods (more home range overlap; Augustine et al. 2019). See Appendix C for the full simulation study specifications.

## Results

### Results – Fisher Application

The Genotype SPIM analysis produced abundance and density estimates that were 25% more precise than the SCR estimate as judged by the coefficient of variation (Table 1). The increased precision was largely driven by an increase in individual detectability and more precise spatial scale parameter estimates when including the 157 samples originally discarded. The overall detection parameter (*a*_0_) point estimate increased 44% from 3.28 to 4.73 and the number of individuals detected was estimated at 272, a 45% increase over the 187 detected individuals in the original, curated data set. The Genotype SPIM abundance point estimate was 15% higher than the SCR estimates; however, given the level of uncertainty in both estimates, there is no indication that these two estimators would differ on average in their point estimates. The certain identity assignments made by the Genotype SPIM (posterior match probability of 1) corresponded to those made in the original study except for 5 cases described in Supplement 1. Two of these 5 cases were examples where the Genotype SPIM assignment implied that the geneticist was too confident in probably correct assignments (assigned a match when the Genotype SPIM estimated match probability was ¡1, but ¿ 0.75), but in 2 cases, the Genotype SPIM assignments implied the geneticist assigned different individual identities to samples that were actually from the same individual with probability 1 and 0.92, and in one case, the Genotype SPIM assignment implied the geneticist incorrectly assigned 2 samples to the same individual with probability 1. Spatially-explicit depictions of the posterior identity matches can be visualized for every sample using code provided in the Data Supplement, with a subset of these matches illustrated in Supplement 1.

**Table 1:**
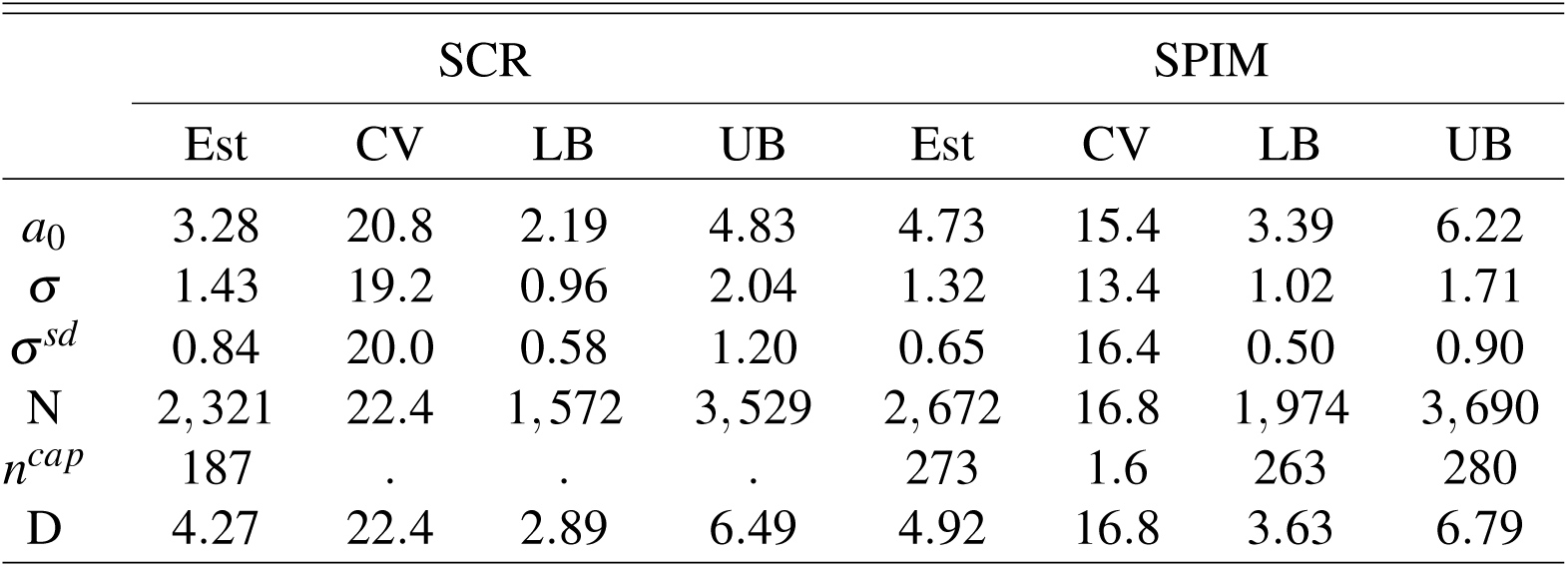
SCR process and observation model parameter estimates from the SCR and SPIM analyses of the fisher data set. *a*_0_ is the overall detection parameter, *σ* is the population-level detection function spatial scale parameter in km, *σ*^*sd*^ is the standard deviation of the individual-level variance in the spatial scale parameter, *N* is the population abundance, *n*^*cap*^ is the number of individuals captured, and *D* is the population density (individuals/100km^2^. Posterior modes are presented as point estimates, posterior standard deviations/posterior modes are presented as the coefficient of variation, and 95% HPD interval upper and lower bounds (LB and UB) are presented as interval estimates.

|  | SCR |  |  |  | SPIM |  |  |  |
| --- | --- | --- | --- | --- | --- | --- | --- | --- |
|  | Est | CV | LB | UB | Est | CV | LB | UB |
| $a_0$ | 3.28 | 20.8 | 2.19 | 4.83 | 4.73 | 15.4 | 3.39 | 6.22 |
| $\sigma$ | 1.43 | 19.2 | 0.96 | 2.04 | 1.32 | 13.4 | 1.02 | 1.71 |
| $\sigma^{sd}$ | 0.84 | 20.0 | 0.58 | 1.20 | 0.65 | 16.4 | 0.50 | 0.90 |
| $N$ | 2,321 | 22.4 | 1,572 | 3,529 | 2,672 | 16.8 | 1,974 | 3,690 |
| $n^{cap}$ | 187 | . | . | . | 273 | 1.6 | 263 | 280 |
| $D$ | 4.27 | 22.4 | 2.89 | 6.49 | 4.92 | 16.8 | 3.63 | 6.79 |

The detection function spatial scale point estimates, *σ* and *σ*^*sd*^, were roughly similar between the Genotype SPIM and SCR models (Table 3, Figure 2), but these parameter estimates were more precise for the Genotype SPIM. This gain in precision is likely due to the greater number of spatial recaptures (individuals captured in more than 1 locations), especially high probability spatial recaptures, contained in the partial genotype samples. The 105 partial genotype samples effectively doubled the number of certain spatial recaptures as seen through the posterior for the number of spatial recaptures (Figure 2), which takes a minimum value of 14, compared to the 9 used in the original data set and SCR analysis. The posterior mode for spatial recaptures (Figure 2) was 24, with a 95% HPD interval of (18 - 31), indicating a high probability that there were more than 2 times as many spatial recaptures than included in the original data set. Among the 157 samples originally discarded, the Genotype SPIM matched 6 to another sample with probability greater than 0.99, 17 with probability greater than 0.9, and 30 with probability greater than 0.75. For these same discarded samples, the Genotype SPIM assigned 11 samples to unique individuals (individuals with 1 capture event) with a probability greater than 0.99, 31 with a probability greater than 0.9, and 49 with a probability greater than 0.75.

**Table 2:**
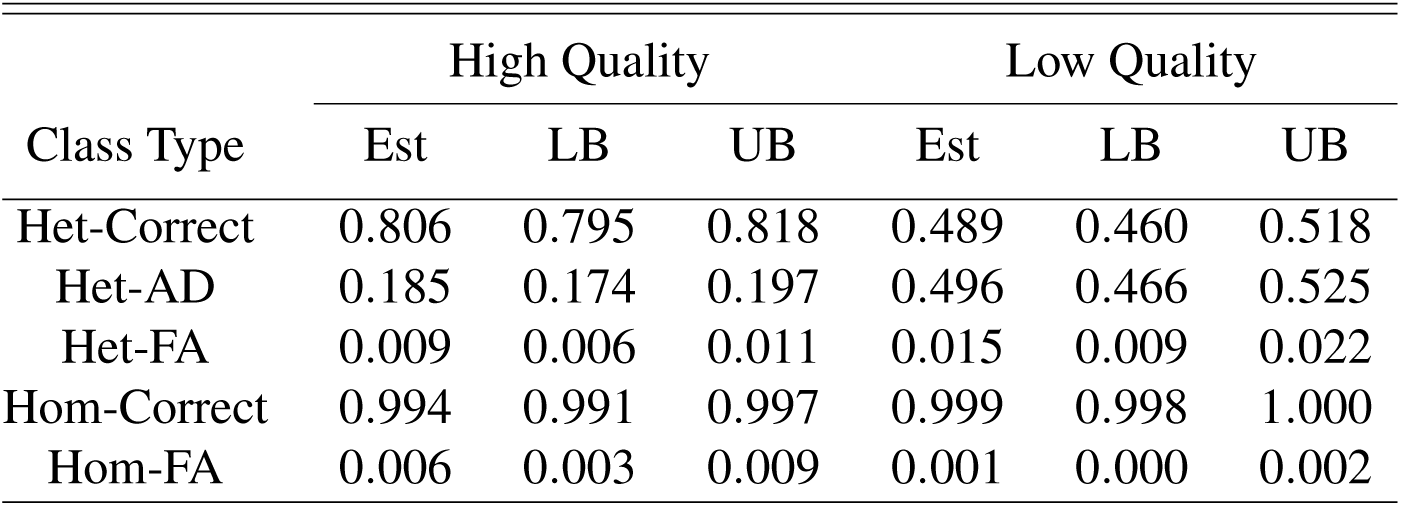
Single locus genotype observation probability parameter estimates for high and low quality samples. “Het” indicates a heterozygous single locus genotype and “hom” indicates a homozygous single locus genotype. “Correct”, “AD”, and “FA” indicate a correct, allelic dropout, and false allele observation, respectively. Posterior means are presented as point estimates and 95% HPD interval upper and lower bounds are presented as interval estimates.

**Table 3:** Genotype SPIM simulation results for the detection model and abundance. Scenarios indicate the model used (SPIM or SCR), the number of replicated assignments (1-3) and whether the low quality samples were included (A) or not (B). The low quality samples were not included in the SCR analysis by default. *λ*_0_ is the baseline detection rate, *σ* is the detection function spatial scale parameter, N is abundance, and *n*^*cap*^ is the number of individuals captured (fewer when excluding the low quality samples). The values listed here are the mean point estimates across 120 simulated data sets. “Cov” indicates the coverage of the 95% credible intervals, “Wid” indicates the mean width of the 95% credible intervals, and “MSE” indicates the mean squared error of the point estimates.

| Scenario | $\lambda_0$ | $\sigma$ | N | $n^{cap}$ | N Cov | n cov | N Wid | n Wid | N MSE |
| --- | --- | --- | --- | --- | --- | --- | --- | --- | --- |
| True | 0.570 | 1.320 | 166.0 | 115.8 | . | . | . | . | . |
| SPIM3A | 0.562 | 1.321 | 165.2 | 115.6 | 0.933 | 0.983 | 36.9 | 2.3 | 105.0 |
| SPIM2A | 0.566 | 1.321 | 164.0 | 115.0 | 0.958 | 0.950 | 37.6 | 4.8 | 119.1 |
| SPIM1A | 0.574 | 1.329 | 159.8 | 112.8 | 0.908 | 0.908 | 41.5 | 10.8 | 171.2 |
| True | 0.297 | 1.320 | 166.0 | 88.4 | . | . | . | . | . |
| SPIM3B | 0.296 | 1.311 | 163.8 | 88.3 | 0.950 | 0.992 | 64.0 | 0.01 | 279.7 |
| SPIM2B | 0.296 | 1.310 | 163.9 | 88.3 | 0.958 | 0.975 | 64.3 | 0.07 | 277.6 |
| SPIM1B | 0.297 | 1.313 | 162.5 | 88.2 | 0.967 | 0.950 | 64.0 | 1.62 | 281.1 |
| SCR | 0.295 | 1.311 | 163.9 | . | 0.983 | . | 64.1 | . | 282.7 |

**Figure 2:**
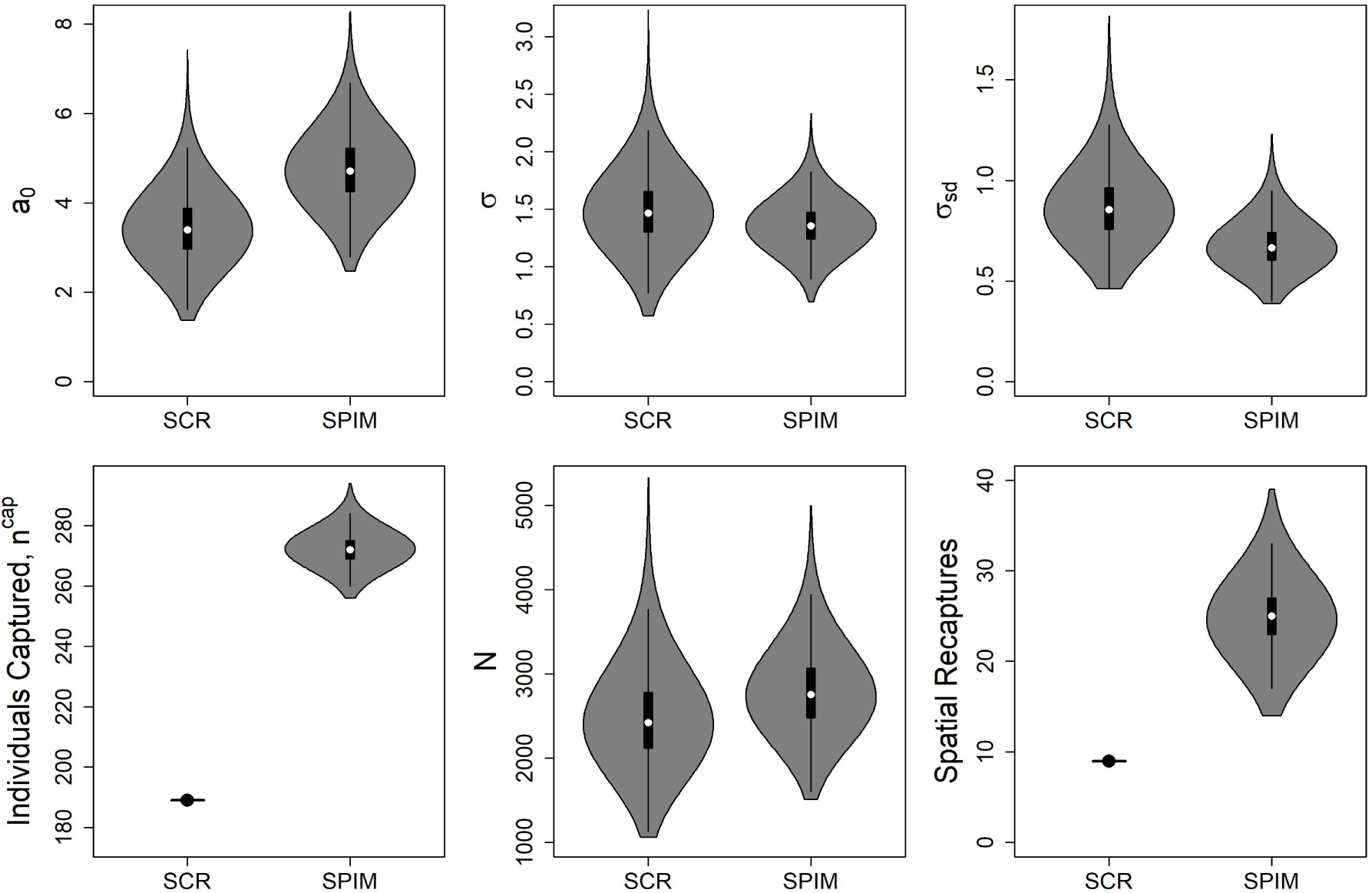
SCR process and observation model posterior distributions from the SCR and SPIM analyses of the fisher data set. *a*_0_ is the overall detection parameter, *σ* is the population-level detection function spatial scale parameter in km, *σ*^*sd*^ is the standard deviation of the individual-level variance in the spatial scale parameter, *n*^*cap*^ is the number of individuals captured, and *N* is the population abundance. The number of captured individuals and spatial recaptures in the SCR analysis are known statistics.

The genotype observation probabilities differed between sample types (high vs. low quality; Table 2). Because allelic dropout can only occur for heterozygous genotypes and false allele rates were very low, homozygous single-locus genotypes were estimated to be almost always scored correctly for both sample types (*>*0.994). The major difference in reliability between sample types was the probability of an allelic dropout observation, which was roughly 1.7 times more likely for low quality samples (0.185 vs 0.496). Despite the general unreliability of the poor quality samples, the overall improvement in the precision of the abundance estimate stemming from their use demonstrates that they still contain substantial information about the population parameters of interest. The single-locus genotype frequency estimates can be found in Supplement 1.

### Results – Simulation Study

The Genotype SPIM abundance estimates were approximately unbiased (≈-1%) with near nominal coverage (Table 3) except when using only 1 genotype assignment (e.g., PCR). In this case, bias was −3.6% when including the low quality samples and −2.1% when only using the high quality samples. The 95% coverage for abundance was only less than nominal in the scenario including the low quality samples with only 1 genotype assignment, where it was 0.91. The Genotype SPIM including the low quality samples (37% of total samples) with 3 replicated assignments was 42% more precise, as judged by the mean 95% CI width, and 63% more accurate, as judged by the mean squared error, than the SCR estimator that did not use the low quality samples. With only 2 replicated assignments, the Genotype SPIM was 41% more precise and 59% more accurate than the SCR estimator. With just 1 assignment, the Genotype SPIM was 35% more precise and 39% more accurate than the SCR estimator. By including the low quality samples, 27.4 more individuals were captured, on average, representing an increase of 31% over the number captured in the high quality samples alone. The uncertainty in the number of individuals captured, *n*^*cap*^, came almost entirely from the low quality samples except when using only 1 genotype assignment. The mean 95% CI width for *n*^*cap*^ when not using low quality samples was effectively 0 when using 2 and 3 replicated assignments (Scenarios SPIM2B and SPIM3B) and the posterior modes of *n*^*cap*^ matched the true value exactly 98% and 99% of the time, respectively. Thus, the individual identities of all high quality samples were assigned correctly with probability 1 nearly 100% of the time, except when there was only 1 genotype assignment, where nearly all individuals were assigned correctly with probability 1, on average.

The genotype observation probability estimates (correct assignment, allelic dropout, false allele) for heterozygous and homozygous true genotypes were approximately unbiased when using more than 1 genotype assignment, except when low quality samples were included, where they were approximately unbiased when using 3 replicated assignments (Table 4). In scenarios with bias, the allelic dropout probability 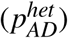 and false allele probabilities (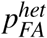 and 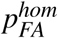) estimates were positively biased, with a corresponding negative bias in the correct observation probability estimates (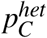 and 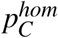). There was more bias in the genotyping error probabilities for the low quality samples; however, there was less overall bias in the genotyping error probabilities for the high quality samples when including the low quality samples and using only 1 genotype assignment (Scenarios SPIM1A vs. SPIM 1B), likely due to the overall greater precision in the detection and abundance parameters when including these samples. The estimates of ***p***^*het*^ and ***p***^*hom*^ for high quality samples with 1 genotype assignment were 1 - 6% more precise (depending on the parameter), as judged by the posterior standard deviation when including the low quality samples compared to when they were excluded. There was also some precision gain in the estimates of ***p***^*het*^ and ***p***^*hom*^ by including the low quality samples with 2 replicated assignments, but they were less pronounced (1-2% precision gain). Precision gains in the estimates of ***p***^*het*^ and ***p***^*hom*^ including the low quality samples with 1 genotype assignment were negligible (*<*1%).

**Table 4:** Genotype SPIM simulation results for the genotype observation model. Scenarios indicate the model used (SPIM), the number of replicated assignments (1-3) and whether the low quality samples were included (A) or not (B). “C”, “AD”, and “FA” indicate correct, allelic dropout, and false allele probabilities, respectively. “Het” indicates heterozygous genotypes and “hom” indicates homozygous genotypes. The high quality sample parameters are indicated with a “1” and low quality indicated with a “2”.

| | $p_C^{het1}$ | $p_{AD}^{het1}$ | $p_{FA}^{het1}$ | $p_C^{het2}$ | $p_{AD}^{het2}$ | $p_{FA}^{het2}$ | $p_C^{hom1}$ | $p_{FA}^{hom1}$ | $p_C^{hom2}$ | $p_{FA}^{hom2}$ |
| --- | --- | --- | --- | --- | --- | --- | --- | --- | --- | --- |
| True | 0.806 | 0.185 | 0.009 | 0.489 | 0.496 | 0.015 | 0.994 | 0.006 | 0.999 | 0.001 |
| SPIM3A | 0.806 | 0.185 | 0.009 | 0.486 | 0.498 | 0.016 | 0.993 | 0.007 | 0.997 | 0.003 |
| SPIM2A | 0.805 | 0.185 | 0.009 | 0.484 | 0.500 | 0.016 | 0.993 | 0.007 | 0.996 | 0.004 |
| SPIM1A | 0.797 | 0.194 | 0.010 | 0.479 | 0.503 | 0.018 | 0.993 | 0.007 | 0.989 | 0.011 |
| SPIM3B | 0.806 | 0.185 | 0.009 | . | . | . | 0.994 | 0.006 | . | . |
| SPIM2B | 0.805 | 0.186 | 0.009 | . | . | . | 0.994 | 0.006 | . | . |
| SPIM1B | 0.792 | 0.198 | 0.010 | . | . | . | 0.993 | 0.007 | . | . |

The simulation results for *n*^*cap*^, the number of individuals captured, indicate some suboptimal performance of the Genotype SPIM when including the low quality samples (Table 3). In these scenarios with low quality samples, the *n*^*cap*^ estimates are slightly negatively biased, with these effects increasing as the number of replicated assignments decreases. With only 1 genotype assignment, 95% coverage of *n*^*cap*^ was less than nominal (0.91). Inspection of these simulation results revealed that low quality samples for individuals only captured once were rarely assigned incorrectly to neighboring individuals that had similar genotypes at the loci observed in the low quality samples. In cases where this happened, the low quality sample was incorrectly scored at one or more loci in all replicated assignments where a score was made (e.g., 1 allelic dropout score and 2 failed amplifications or 2 allelic dropout scores and 1 failed amplification). The observed bias in *n*^*cap*^ increased with a decreasing number of replicated assignments because the expected number of loci with no score and the expected number of loci where the correct genotype was not included in any replicated assignment increases with fewer replicated assignments. This source of bias in *n*^*cap*^ also likely explains the bias seen in the genotype observation probability estimates for low quality samples with 1 PCR.

## Discussion

We developed the Genotype SPIM, a unified probabilistic framework for the processes of determining the true genotypes of samples, matching samples to individuals, and estimating population parameters using spatial capture-recapture. The Genotype SPIM recognizes that uncertainty in the genetic classification and population estimation processes is sequentially connected and propagates this uncertainty from one process to another, using a hierarchical model. The Genotype SPIM allows for a fundamental shift in the use of genetic data in capture-recapture, eliminating the need for decision rules that determine the (minimized) expected level of error in the individual identity assignments and that do not propagate identity error probabilities to the population parameter estimates. Perhaps most importantly, the Genotype SPIM eliminates the need for data culling, which can be extreme in many non-invasive data sets where DNA quality is typically poor, and leverages the additional information to increase the precision and accuracy of population parameter estimates. This is especially important in conservation applications of many species that are of concern largely because of their extremely low population sizes.

Unlike all existing attempts to address genotype uncertainty in capture-recapture, the Genotype SPIM recognizes that ecological systems are spatially explicit and it uses the spatial location where genetic samples were collected to reduce genotype uncertainty. Two key features of our model that allow it to exploit the spatial information of genetic samples are a spatially explicit model for the number and distribution of individuals and genotypes across the study area and a spatially-constrained model for individual detection (see Figure I). This genotype-augmented ecological and capture model provides the scaffolding that allows for the shadow effect to be efficiently resolved (disallowed in the most similar nonspatial model of Wright et al. 2009) and which formally links the ecological concepts of population density and home range size to the uncertainty in assigning samples to individuals (Augustine et al. 2019). The Genotype SPIM recognizes that samples collected closer together in space are more likely to come from the same individual, so each sample carries information about the true genotype of their neighboring samples and the genotyping errors that likely did or did not occur in these neighboring samples. This contrasts with previous approaches for matching samples to individuals in the presence of genotyping errors that assume the samples are independent of one another (e.g., Kalinowski et al. 2006; Macbeth et al. 2011; Sethi et al. 2016). The spatial locations where samples were collected reduce the uncertainty in each captured individual’s estimated genotype, improve the estimation of genotype frequencies and genotyping error rates, and improve the probabilistic assignment of samples to individuals. The net result is improved estimates of population parameters. Posterior states for the probabilistic identity assignments can be visualized for every sample, providing an understanding of how the ecological, capture, and genotype observation models combine to produce the probabilistic assignments (see examples in Supplement 1)

We believe it is this spatial structure of the Genotype SPIM that allows for the high probability identity assignments for many of the “low quality” samples in the Fisher application (see Supplement 1), for the ability of the low quality samples to substantially improve inference in the simulation study, and for the identifiability of the model parameters with only 1 genotype assignment in the simulation study and supplementary application (see Appendix E). Unlike the most similar nonspatial model of Wright et al. (2009), the Genotype SPIM is identifiable with no genotype information at all, at least when the level of uncertainty in individual identity is not too high (Augustine et al. 2019), as it reduces to the model of Chandler and Royle (2013) for unmarked SCR. Augustine et al. (2019) showed that introducing categorical identity covariates (e.g., genotypes) known with certainty can greatly improve estimation over unmarked SCR and not much categorical identity information is required to achieve certainty in individual identification when individual space use is restricted relative to the complete extent of the population under study. The Genotype SPIM builds on the model of Augustine et al. (2019) to allow for the genotypes to be observed with error. Assuming that false allele rates will always be very low, the homozygous single-locus genotypes will be recorded with near certainty, leaving just the heterozygous single-locus genotypes for which there is nonnegligible uncertainty. Thus, the Genotype SPIM will match many samples with high certainty that would not pass typical *P*_*ID*_ thresholds and it should match samples with higher certainty than the nonspatial model of Wright et al. (2009) in many scenarios. No simulation study of the Wright et al. (2009) model has been published, but we expect low quality samples to be less useful in that model without their associated spatial information and we expect that model to rely more heavily on the assumptions of the genotype distribution and error models than the Genotype SPIM, which does not rely on these submodels for parameter identifiability.

An alternative SPIM that allows all samples to be used without modeling the genotyping error process is hinted at in Figure I in the individual identity observation process under the “typical approach” to applying SCR to genetic samples. We conceptualized the process of assigning individual identities to samples as a “random thinning” process (following others, see below), where samples lose their individual identities at random with probability 1-*θ*. This produces two data sets–one with individual identities, and one without. The data set with no individual identities is typically discarded, which can be extremely detrimental in conservation decision making for species that are most information poor and of high conservation concern. Alternatively, the unknown identity samples could be used to probabilistically reconstruct the full data set without consideration for their replicated genotype scores. This “random thinning” model has previously been developed (Richard Chandler, pers. comm.), but not published, and is identical to the random thinning process specified for marked individuals in spatial mark-resight by Jimenez et al. (2019). While not requiring the replicated assignment data nor assumptions about the genotype distributions or the genotyping observation process, this approach has two disadvantages. First, it does not incorporate any uncertainty in the individual identity assignments for the samples assigned individual identities–this data set may still contain errors as we found in the fisher application. Second, the information contained in the replicated assignments increases the probability with which samples can be assigned to individuals. Discarding this information can significantly reduce the information about individual identity–the only information remaining is the spatial location. Thus, the value of using the unidentified individual samples will decrease much more rapidly as population density increases, compared to the full information samples.

While the genotyping error rates are effectively nuisance parameters whose estimation is required for correct inference about the population parameters of interest, the Genotype SPIM may provide a more statistically powerful framework for exploring the mechanisms of genotyping error in low quality DNA samples and quantifying their relative influence absent known identity, reference genotypes. As expected, we estimated that false allele events were very rare, with a per locus per replicate error probability of approximately 0.01 (Table 2). Allelic dropout events were estimated to be much more common, with probabilities of 0.19 and 0.50, for high and low quality samples, respectively. Interestingly, we estimated that the false allele probability varied by zygosity, with false alleles being more common for true heterozygous single-locus genotypes than true homozygotes (Table 2). This difference was more pronounced for low quality samples (0.015 vs. 0.001) than for high quality samples (0.009 vs. 0.006). A possible explanation for this result is that a major source of false alleles is allele-calling errors which disproportionately occur for true heterozygotes (Johnson and Haydon 2007)–a false allele event is more likely when reading 2 allele peaks for a heterozygote than when reading the single peak for a homozygote. One apparently anomalous result is that the false allele probability for homozygous loci from low quality samples is lower than that for high quality samples. A possible explanation for this result is that the majority of homozygous loci for low quality samples that would have produced a false allele if they amplified failed to amplify. PCR artifacts are a source of false alleles for homozygotes (Johnson and Haydon 2007), and the mechanisms leading to PCR artifacts may lead instead to failed amplification more often in the lower quality samples. The genotyping error rates estimated by our model are for the loci that amplify only. If there is a correlation between any of the genotyping error rates and the amplification rates, the most error prone samples/loci will drop out disproportionately and the genotyping error rates estimated for different sample qualities must be interpreted accordingly. This estimation of error rates from censored data is not a problem for estimating the population parameters in the presence of genotyping error, as only the error rates of the amplified samples/loci are relevant.

We believe the Genotype SPIM will be relatively robust to misspecification of the genotyping error model, especially in lower density populations, because there may only be a few possible individuals with activity centers near any particular focal sample, and even fewer with similar genotypes. This property should be investigated via simulation and through application to existing data sets. Genotyping error rates may vary as a function of error type (allelic dropout vs. false allele), zygosity (heterozygote vs. homozygote), sample quality, loci, replicate number, or individual. Of these factors, we expect the largest differences to be between error types and sample quality, both of which we accommodated in the Genotype SPIM. Error rates could be generalized to vary by loci and replicate number, though we don’t see a plausible reason why they would vary by individual. Paetkau (2003) identified that faulty lab procedures can lead to a lack of independence between samples, and that sample quality can lead to a lack of independence across markers for the same sample. The former source of dependence could be accommodated with replicate covariates and the latter with sample type covariates. Instead of dividing samples into “low” and “high” quality categories, one could use sample covariates that correlate with the amount and/or quality of the DNA in each sample, if they are available. In the absence of these covariates, crude sample type categories based on the amplification rates across replicated assignments should improve inference.

Thousands of genetic capture-recapture data sets exist to which the Genotype SPIM could be applied and the probabilistic individual identity assignments it makes can be compared with those made using typical methods to identify possible inadequacies with the Genotype SPIM model structure for any particular application. For the fisher application, the Genotype SPIM made all the same identity assignments as the geneticist did for the samples originally assigned certain individual identities, except for 3 cases where it appears the Genotype SPIM identified erroneous assignments made by the geneticist (see Supplement 1). This near perfect agreement in individual identity assignments for the subset of samples assigned identities by the geneticist supports the adequacy of the genotyping error model we used, though we cannot independently assess the performance of the model for the lowest quality samples that were not originally assigned individual identities.

Violations of the genotype distribution model are also possible (see Augustine et al. 2019). We will discuss one way in which the possibility of genotyping error may exacerbate an assumption violation for the genotype distribution over the model without genotyping error. We assume, as is typical in methods for assigning individual identities to samples (e.g., Wright et al. 2009; Kalinowski et al. 2006; Macbeth et al. 2011), that the distribution of genotypes are independent across individuals; though spatial genetic structure due to relatedness may be present due to philopatry in some species. In this case, nearby genotypes may be more similar than expected under independence. When genotyping error is allowed, it is possible that samples from two individuals with *very similar* genotypes will erroneously be combined because they only differ at one or a few loci where the differences could plausibly be due to genotyping errors. Therefore, spatially correlated genotypes due to philopatry could introduce negative bias into the Genotype SPIM abundance and density estimates, especially when the overall detection rate is lower and the proportion of samples with sparse genotype information is higher.

Perhaps the most critical assumption of the Genotype SPIM is the proper specification of the detection model. Specifically, individual heterogeneity in detection function parameters has been identified as a challenge for SPIMs in general (Augustine et al. 2018*b*, 2019). We were able to accommodate rather extreme individual heterogeneity in detection function parameters in the fisher application using a modified version of the model of Efford and Mowat (2014), which specifies a deterministic, negative relationship between 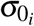 and 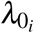 with a single individual random effect on 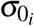. However, this detection model was not sufficient for all long distance spatial recaptures to be linked up during convergence using our default algorithm to initialize individual identities to samples (see Appendix B) and required an informative prior for the *σ* standard deviation. Alternatively, individual heterogeneity in *λ*_0_ and *σ* with independent random effects on each parameter could be considered, but this model is likely challenging to fit with typically sparse SCR data sets and it would be more difficult to specify an informative prior on *λ*_0_ than it is for *σ*, which is related to home range size. A second alternative model that may be useful for accommodating individual heterogeneity in space use is that of Royle et al. (2016) where a subset of or all individuals are allowed to have transient activity centers across sampling occasions. This model will not be helpful if the very large spatial recaptures occur on the same capture occasion, which did occur for the 1 individual in the fisher data set that prevented convergence. Given the importance of individual heterogeneity in detection function parameters to the performance of SPIMs, we recommend further research and model development, though given the typical sparsity of individual recaptures and spatial recaptures in SCR data sets, feasible models will need to be very simple and/or rely on parameters for which ecologically-informed priors can be set (e.g. *σ*^*sd*^). An alternative solution is to disallow the shadow effect for particular samples by deterministically linking certain long distance spatial recaptures as can be done in the categorical spatial mark-resight model (Augustine et al. 2018*a*).

While the Genotype SPIM can improve inference for data sets produced using the current genotyping protocols that seek to maximize certainty in individual identity for just a subset of samples, it is possible that by relaxing the requirement that individual identities are assigned with certainty, current genotyping protocols are no longer optimal. The simulation study demonstrates that the amount of information obtained about individual identity from each replicate assignment declines exponentially, while the costs increase linearly, so less replication in general may provide a better compromise between project cost and the precision of population parameter estimates. Further, it is possible that more replication of the low quality samples that are typically discarded as unreliable leads to larger improvements in population parameter estimates than replication of the high quality samples which have been the focus to date. In the simulation study, adding replicate assignments over the first assignment improved the abundance estimate precision and accuracy very minimally when using only the high quality samples (Figure 3, Table 3). Including the low quality samples improved both the precision and accuracy of the abundance estimate and adding a second replicated assignment in this scenario reduced bias, substantially improved the accuracy, and modestly improved precision. Thus, it is possible that current applications of the multi-tubes approach may be allocating too much effort to replicating high quality samples, and protocols that preclude the replication of poorly performing samples may be misallocating resources when the Genotype SPIM can be used. Because the Genotype SPIM provides a single probabilistic framework for genetic capture-recapture, different genotyping protocols can be evaluated via simulation using clearly defined assumptions about the ecological, capture, and genotyping processes. The minimum level of replication required in practice will depend on the specifics of the data set, but see Appendix D for an application demonstrating that density can be estimated with only 1 PCR per sample using a real data set.

**Figure 3:**
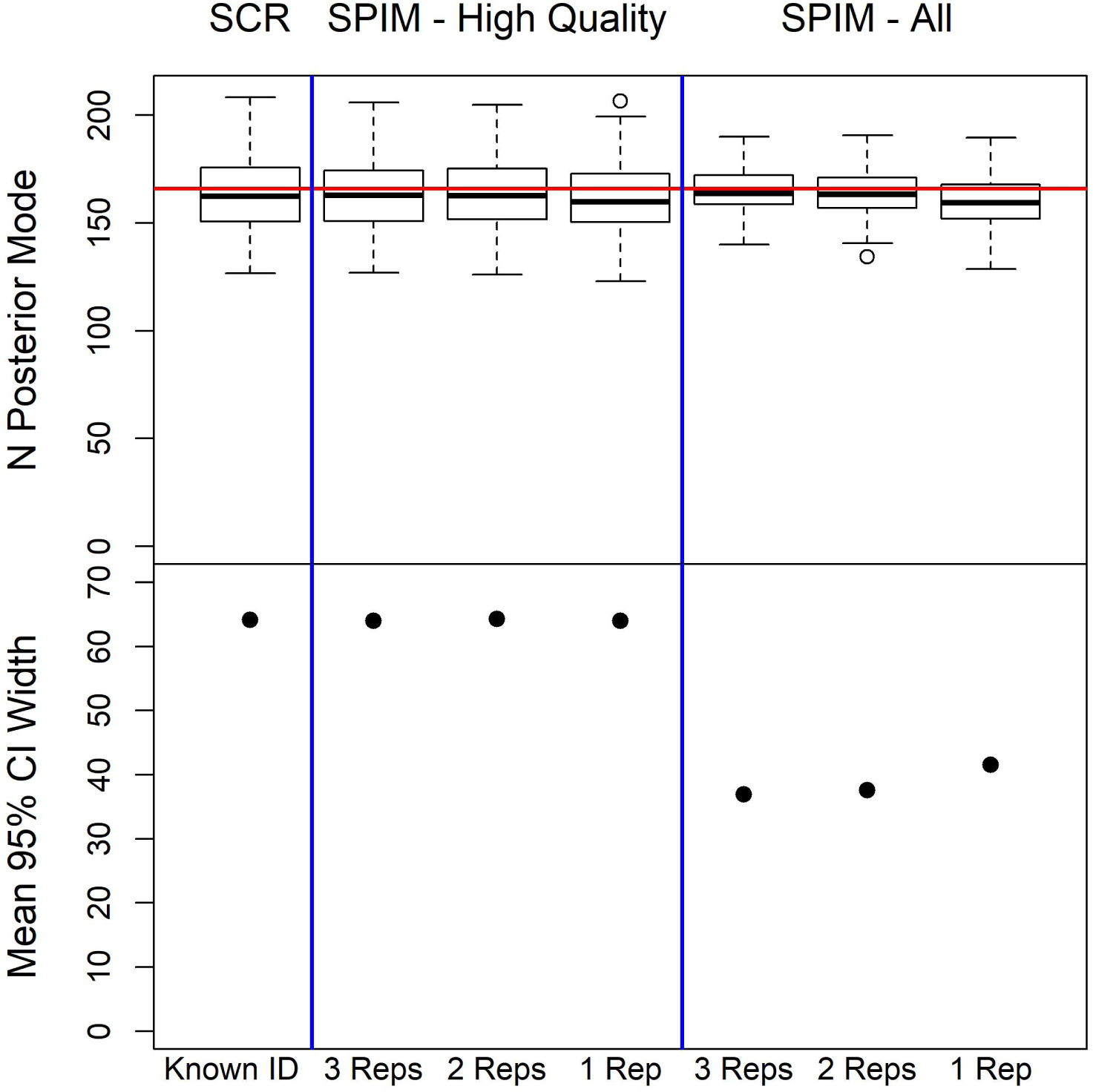
Plots of simulation study results for the estimation of abundance. The top row displays boxplots of the posterior modes for abundance (Simulated value is 166), showing estimator accuracy, and the bottom row displays the mean 95% CI width, a measure of estimator precision. The estimators from left to right are the SCR estimator using only certain identity samples, followed by the Genotype SPIM estimators using only the high quality samples with 3, 2, or 1 replicated assignments, followed by the Genotype SPIM estimators using both high and low quality samples with 3, 2, and 1 replicated assignment.

The general structure of the Genotype SPIM should also be useful for other noninvasive observation methods such as remote cameras, to which machine learning algorithms for classification are increasingly being applied (e.g., Arzoumanian et al. 2005; Norouzzadeh et al. 2018). Currently, these machine learning algorithms assign individual identities using the photographs alone, with no linkage to the ecological or capture processes, and typically require training data of known identity individuals, which is often not available. Individual classification should be improved by linking the process of identifying individuals to the ecological and capture processes, especially in situations of lower signal to noise ratios, and this linkage facilitates the propagation of uncertainty to the population parameters of interest. To date, the use of machine learning to produce individual identities from photographs has been mostly applied to problems with a high signal to noise ratio, for example, spot patterns on animal flanks (e.g., Arzoumanian et al. 2005; Crall et al. 2013), and the uncertainty in assigning the individual identities has not been propagated to the population parameters of interest (but see Ellis 2018). Similarly, linking the *species* identification process to the ecological and capture processes when using machine learning to produce species records for occupancy studies should also improve inference. We provide a small simulation study and an application of the general catSPIM with observation error model to an Andean bear data set in Appendix E, though we argue the model did not work well in this application due to a signal to noise ratio that was too low. However, this example shows how an application might work in a situation where more information about individual identity can be reliably extracted from photographs.

Noninvasive genetics has revolutionized the study of animal populations by capture-recapture, allowing for the study of many species that could not have been studied effectively using conventional methods based on physical capture or species that cannot be individually-identified from camera traps. However, noninvasive studies often result in sparse data sets, that can lead to imprecise population parameter estimates which are of limited use in conservation decisions because they lead to an inability to discriminate between a population in need of conservation action and one that is not. Biased estimates of population parameters can be even more problematic, potentially falsely indicating that an imperiled population does not warrant conservation action, risking extinction, or that a healthy population does warrant conservation action, leading to an ineffective allocation of resources. Therefore, statistical methods that produce unbiased and precise population parameter estimates are vital for making reliable, evidence based,conservation decisions. Our genotype SPIM model links genetic classification with the ecological and capture process, and accordingly results in increased accuracy and precision of density estimates with less bias and no data loss, which can lead to more informed conservation decision making.

## Acknowledgments

We would like thank Dana Morin and Chris Sutherland for contributing to model development and Richard Chandler for many ideas relevant to updating latent individual identities in SCR MCMC algorithms. Research funding was provided by Cornell University’s Atkinson Center for a Sustainable Future (BA). Any use of trade, firm, or product names is for descriptive purposes only and does not imply endorsement by the U.S. Government.

## Box 1

The Genotype SPIM is a 3-level hierarchical model, with submodels for the ecological, capture, and genotype observation processes. The ecological process determines the abundance (N), density, and spatial locations (***S***) of the individuals in the study area and associates a genotype (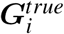 with each individual, which are governed by the loci-level genotype frequencies, ***γ***. These genotypes may not be unique (shadow effect)–we depict unique genotypes with unique colors, with 2 individuals sharing a “red” genotype. The capture process then determines where and how many times each individual will be captured, 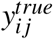, governed by a detection function between the location of individual *i* and trap location, ***x*** _*j*_. When captured, individuals leave a record of their genotype, not their unique individual identity, due to the possibility of the shadow effect. The genotype observation process then determines which genotype we observe, 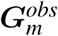, for sample *m* on replicate *l*, conditional on the true genotype of the individual that was captured and the genotype observation probabilities in *π*. We gray out the observed genotypes because the true genotypes are no longer observed perfectly, and indicate possible genotyping errors with red exes. The data for the Genotype SPIM are the spatially referenced observed genotypes, which are used to probabilistically reconstruct ***Y*** ^*true*^ and incorporate the uncertainty in individual identity into the population parameter estimates. To contrast the Genotype SPIM with the “typical approach”, we can conceptualize the individual identity observation process as a random thinning process where the true capture history, ***Y*** ^*true*^ is split into a capture history of known identity samples, ***Y*** ^*ID*^, and a vector of trap-level or a matrix of trap by occasion-level counts, ***Y*** ^*unk*^, which is discarded. A possible thinning process for an individual by trap capture history is 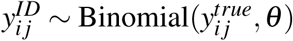, where the *θ* parameter determines the probability that a sample can be identified to individual. *θ* is then a function of the overall quantity and quality of DNA in the samples, but also, the level of conservativeness used for accepting samples as reliable. For the same set of samples, a more conservative genotyping protocol will raise *θ*, leading to fewer individual identity errors in ***Y*** ^*ID*^ at the cost of discarding more samples. This trade-off cannot be avoided if no errors are allowed in individual identity and we cannot guarantee that ***Y*** ^*ID*^ has no errors.

**Figure I:**
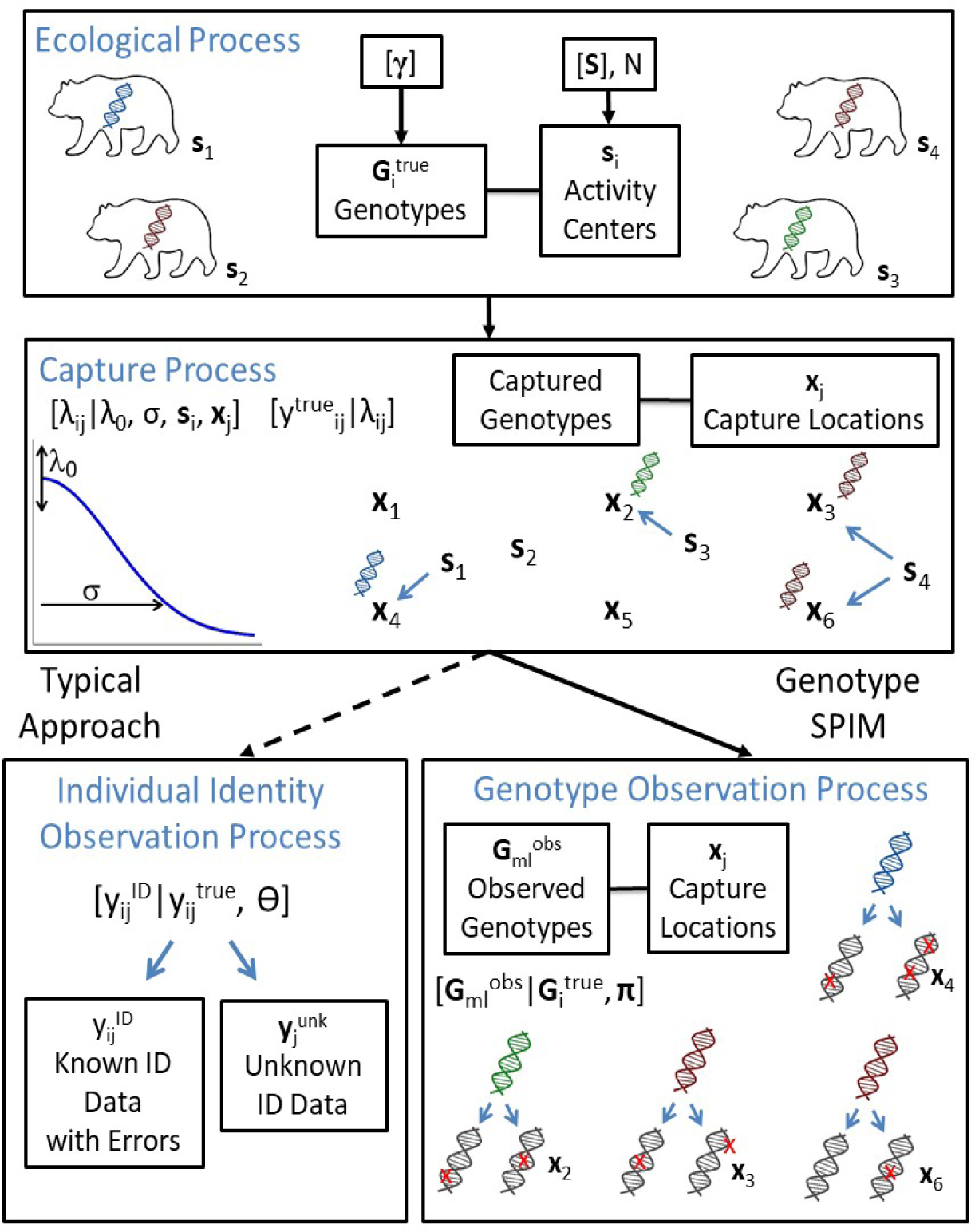
A graphical depiction of the Genotype SPIM contrasted with the typical approach to genetic capture-recapture

## Appendix A: Technical Details and MCMC Algorithm

Here, we describe the technical details of the model structure omitted in the main text and the novel features of the MCMC algorithm. We refer readers to Table A1 below, where we reproduce a table with definitions for model parameters and data structures (observed and latent) for the categorical Spatial Partial Identity Model (catSPIM) from Augustine et al. (2019), followed by the new parameters and data structures required to allow for observation error in the category levels. We retain the same notation from Augustine et al. (2019) for consistency.

### General Model for Observation Error

In order to relax the requirement of the catSPIM that category levels must be recorded correctly, we introduce the possibility that the individual identity category levels of each sample are observed multiple times, subject to error. For brevity, we will refer to these replicated category level observations or assignments as “replicated assignments”, which could be replicated category level observations from multiple observers looking at features in photographs (See Appendix E on Andean Bears) or replicated DNA scores across multiple amplifications and PCRs to determine the values of microsatellites or SNPs across loci. Let ***G***^*obs.true*^ be an *n*^*obs*^ × *n*^*cat*^ matrix, where each row, *l*, contains the full categorical identity of individual *i*, to which sample *l* belongs, with the trap of capture recorded in the *l*^*th*^ row of ***Y*** ^*obs*^. More specifically, 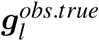 takes the same values as 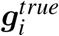 when sample *l* comes from individual *i*. ***G***^*obs.true*^ corresponds to ***G***^*obs*^ in the catSPIM model, except there are no missing covariate values because ***G***^*obs.true*^ is a partially latent, rather than fully observed, data structure. In this model with observation error, we define ***G***^*obs*^, containing the replicated assignments, to be an array of size *n*^*obs*^ × *n*^*cat*^ × *n*^*rep*^, where *n*^*rep*^ is the number of replicated assignments, indexed by *m*. Then, 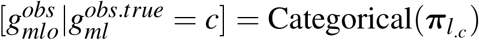, where ***π***_*l*_ is an 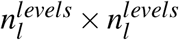 matrix of category level observation probabilities, and *c* is the matrix column matching the value of 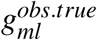. These observation probabilities may or may not differ across identity covariates, *l*. More specifically, 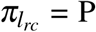 (observer classifies as category level *r |* true category level *c*) for covariate *l*. As with the catSPIM, missing category level values are allowed in ***G***^*obs*^, coded as a “0”. We assume these are missing at random with respect to individual and identity covariate values (i.e., missingness does not contain information about individual identity or the value of the identity covariate).

This general model structure for category level observation error can accommodate a wide variety of observation systems and error processes; however, it will need to be modified if there are more than just a few category levels in order to estimate error probabilities for an increasing number of possible observation events with typically sparse capture-recapture data sets. Multiple observation events may be combined into a reduced number of event types if a priori knowledge about the error processes is available. We will take this approach in the next section to modify the observation process above for genetic capture-recapture where we consider three possible outcomes–a genotype locus can be classified correctly, there can be allelic dropout, or there can be a false allele. A similar approach can be taken for other types of error processes for example, reading tags or bands, (Cowen and Schwarz 2006; Bonner et al. 2016). We will refer to the most general model structure presented above as “catSPIM-OE” to distinguish it from the genotyping model to follow, though the catSPIM-OE error structure may be appropriate for SNPs which can only take 2 values per locus, where there are only 2 possible error types. See Appendix E for a simulation study of catSPIM-OE and an application of catSPIM-OE to camera trapping data of Andean bears.

### Model for Genotyping Error

Before proceeding, a cursory description of microsatellites and genotyping errors is required. Microsatellite markers consist of genetic sequence repeats where the number of repeats determines the value of an allele, of which there are two per locus. There is no ordering of the alleles, and the allele with the fewest number of repeats is customarily listed first. For example, a microsatellite locus for one individual may have a value of 150.152, indicating one allele has 150 repeats and the other has 152. In this case, we say that 150.152 is a single locus genotype, while the combined values across multiple loci constitute a multilocus genotype. A single locus genotype is said to be heterozygous if the two alleles do not share the same value, say 150.152, and homozygous if they do, say 150.150. This is an important distinction for how genotyping errors arise. The most common genotyping error is allelic dropout (Roon et al. 2005) where a heterozygous single locus genotype is erroneously scored as a homozygote, taking the value of only one of the two alleles present. For example, the heterozygous single locus genotype 150.152 may be scored as either 150.150 or 152.152 as the result of an allelic dropout event. While typically more rare, other errors may occur, including false alleles, laboratory errors, and transcription errors (McKelvey and Schwartz 2004). For simplicity, we will classify genotyping errors into two categories based on their outcome, rather than their cause–we will use “allelic dropout” to describe any event causing a true heterozygote to be scored as a homozygote and “false allele” to describe any event leading to an error other than allelic dropout.

Now, we will describe how microsatellite loci can be used as categorical identity covariates. Similar to Wright et al. (2009), we enumerate the loci-level genotypes from 1 to 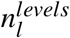 at each loci, *l*, in an arbitrary order. There are several possible ways to determine which genotypes to enumerate at each loci. One option is to include all genotypes that were observed in the current sample, either in all the replicated assignments or in the consensus genotypes. This option will identify all the most common genotypes, but may miss rare genotypes. A more comprehensive option, which will be assumed for the remainder of the model description, is to create a set of genotypes from all possible combinations of the observed set of alleles. This option may still miss some genotypes that occur in the population, but the total number of missed genotypes will necessarily be lower than enumerating only the observed genotypes unless all genotypes were observed. Enumerating all possible genotypes implied by the observed alleles may enumerate genotypes that do not exist in the population; however, their frequencies will be estimated near zero if not observed. Either of these two approaches may enumerate genotypes that do not exist in the population if false alleles are erroneously included in the list of possible alleles, but the frequencies of genotypes containing erroneous alleles should also be estimated near zero if they are rarely observed and/or if genotyping error is modeled as we will address below. Finally, ***γ*** now carries the interpretation of the loci-level *genotype* frequencies. This is distinguished from Wright et al. (2009), where ***γ*** is the *allele* frequencies, which are converted to genotype frequencies by assuming Hardy-Weinberg equilibrium. We do not make this assumption at the cost of estimating more parameters.

With the loci-level genotypes enumerated, the catSPIM-OE model described above can be directly applied to genetic capture-recapture studies using microsatellite markers. One caveat; however, is that we will simplify the elements of ***π***_*l*_, now with the interpretation of the genotype observation probabilities conditional on the true genotype at locus *l*, using the specific genotyping error mechanisms of allelic dropout and false allele events described above. We will refer to this model as the “Genotype SPIM” to distinguish it from the more general model above. We use a simple model for these events that assumes that each possible allelic dropout event is equally likely (previously assumed by Wright et al. 2009; Sethi et al. 2016), each possible false allele event is equally likely (previously assumed by Sethi et al. 2016), and the allelic dropout and false allele probabilities do not vary across sample (relaxed below), locus, individual, or replicate number. We define ***p***^*hom*^ and ***p***^*het*^ to be vectors of the observation probabilities for homozygous and heterozygous loci-level genotypes, respectively. Then, 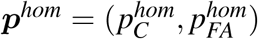 for homozygous correct and false allele observation and 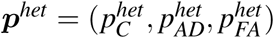 for heterozygous correct, allelic dropout, and false allele observation. Splitting the observation probabilities by zygosity is required due to the differing number of possible observation events for homozygotes and heterozygotes, but it also allows the false allele probabilities to vary by zygosity (see Johnson and Haydon 2007).

The probability of classifying true genotype *c* as observed genotype *r* depends on the values of both *c* and *r* as follows. For correct observation events, 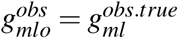, the probability of correctly classifying sample *m* at locus *l* on replicate *o*, 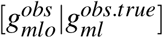 is 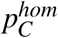 and 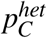 for homozygous and heterozygous genotypes, respectively. For allelic drop out events, i.e., 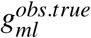 is heterozygous and 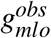 is homozygous matching one of the two alleles in 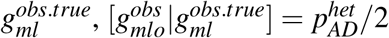. We divide the allelic dropout probability by two because there are two ways to observe allelic dropout. Finally, for false allele events, i.e., 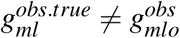 and 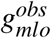 is not homozygous taking a value matching one of the alleles in 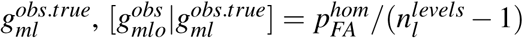 for homozygous genotypes since there are no possible allelic dropout events and one correct observation event and 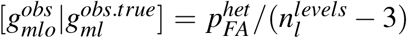 for heterozygous genotypes since there are 2 possible allelic dropout events and one possible correct observation event. These probabilities are plugged into the appropriate elements of ***π***_*l*_ for each loci *l*. The genotype observation probabilities will need to be modified from these listed if not all possible loci-level genotypes implied by the observed loci-level alleles are enumerated and included in the model.

To illustrate how the genotyping errors are modeled, consider a genetic capture-recapture study with three possible alleles at the first locus–150, 152, and 154. The set of all possible genotypes at this locus is (150.150, 150.152, 150.154, 152.152, 152.154, 154.154). Then, the genotype observation probabilities are:

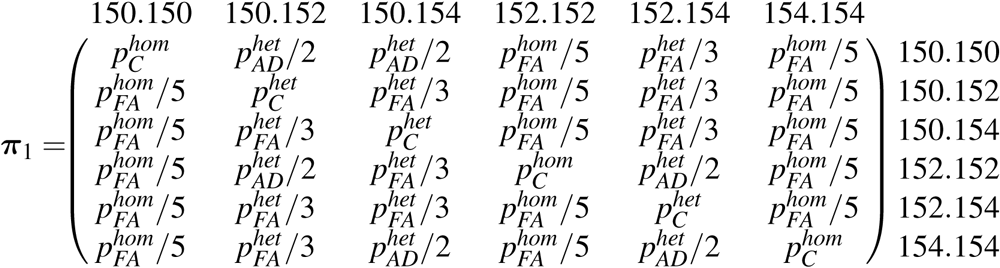

where the column labels indicate the true genotype and the row labels indicate the observed genotypes. Note, columns must sum to 1 and the elements of ***π*** will vary across loci, depending on the number and values of possible genotypes at each locus.

### Alternate Observation Models

Previous studies (Augustine et al. 2018*a*,*b*, 2019) have suggested individual heterogeneity in detection function parameters, especially *σ*, can erode the performance of SPIMs for real world data sets. Here, we consider a model for individual heterogeneity in detection that assumes there is an inverse relationship between *λ*_0_ and *σ*, specifically, *λ*_0_ = *a*_0_*/*(2*σ* ^2^) (Efford and Mowat 2014). This model is motivated by the idea that an individual’s detection rate in space should be proportional to its utilization distribution or rate of space use. Efford and Mowat (2014) interpret *a*_0_ as the “single detector effective sampling area” by showing that under idealized conditions, *a*_0_ is approximately equal to the total individual-level effective sampling area. However, we will interpret *a*_0_ more generally as the overall detection parameter that scales a bivariate normal space use model following *λ* (***x***) = *a*_0_ *f* (***x***) where *λ*(·) is the detection function for the expected counts described above, *f* (***x***) is a bivariate normal PDF, and ***x*** is a spatial location. We modeled an individual random effect on *σ* where 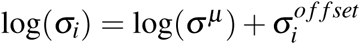 and 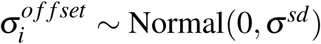. The induced random effect on baseline detection is then 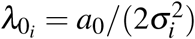. The estimated parameters of this model are *a*_0_, *σ*^*μ*^, *σ*^*sd*^, and the *σ* offsets vector ***σ***^*o f f set*^.

For the Genotype SPIM described above, we make the assumption that the the genotype observation probabilities do not vary across sample, locus, individual, or replication number. Here, we will relax the assumption that the genotype observation probabilities do not vary across samples because samples can vary considerably in the amount and quality of DNA they contain and thus the probabilities that genotyping errors will occur (Paetkau 2003). While this variability could be modeled as a function of continuous and/or categorical sample-level covariates using a mutlinomial logistic link, for conceptual and computational simplicity, we just consider that samples can be split into high and low quality categories with category definitions specific to the application. We estimate separate genotype observation probabilities for each category, ***p***^*hom*-*high*^, ***p***^*hom*-*low*^, ***p***^*het*-*high*^, ***p***^*het*-*low*^, which are organized in genotype observation matrices, ***π***^*high*^ and ***π***^*low*^. Further covariate effects for samples, locus, or replication number could be accommodated using the multinomial logistic link and fit using an auxiliary variable model (Holmes et al. 2006).

### Data Augmentation and Inference

We use a process similar to data augmentation (Royle et al. 2007) to probabilistically resolve ***Y*** ^*true*^ and ***G***^*true*^ and estimate *N*. In typical applications of data augmentation in capture-recapture MCMC algorithms, the capture histories of the *n* captured individuals are observed, which are then augmented with all zero capture histories up to *M* possible individuals where *M* >> *N*. In SPIMs, *n* is unknown, but we can specify an *M* × *J* matrix, ***Y*** ^*true*^, and initialize it with a possibly true capture history using the spatial proximity of observed samples. Similarly, we specify an *M* × *J* matrix ***G***^*true*^ and initialize it with possibly true full categorical identities consistent with the matched samples in ***Y*** ^*true*^ and their associated observed category levels. Then, we proceed with data augmentation as normal, specifying a vector ***z***, of length *M*, to indicate whether individual *i* is in the population (*z*_*i*_ = 1) or not (*z*_*i*_ = 0), assuming *z*_*i*_ ∼ Bernoulli(*ψ*). This individual-level Bernoulli assumption induces the relationship *N* ∼ Binomial(*M, ψ*), where *ψ* is a nuisance parameter (Royle et al. 2007). Population abundance is a derived parameter, 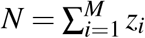, and population density, *D*, is 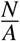. While the number of individuals captured is not known, it can be estimated as a derived parameter following 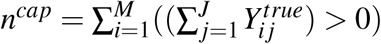 (the number of individuals assigned at least one sample; Augustine et al. 2019). The number of spatial recaptures can be calculated in a similar manner. A final derived parameter of interest is the posterior probabilities of pairwise sample matches, all sets of P(sample A and sample B came from the same individual), which are just the proportion of the posterior samples for which samples A and B are assigned to the same individual.

### 1 MCMC Details

The joint posterior of the categorical SPIM with observation error is very similar to that of the categorical SPIM. The joint posterior is:

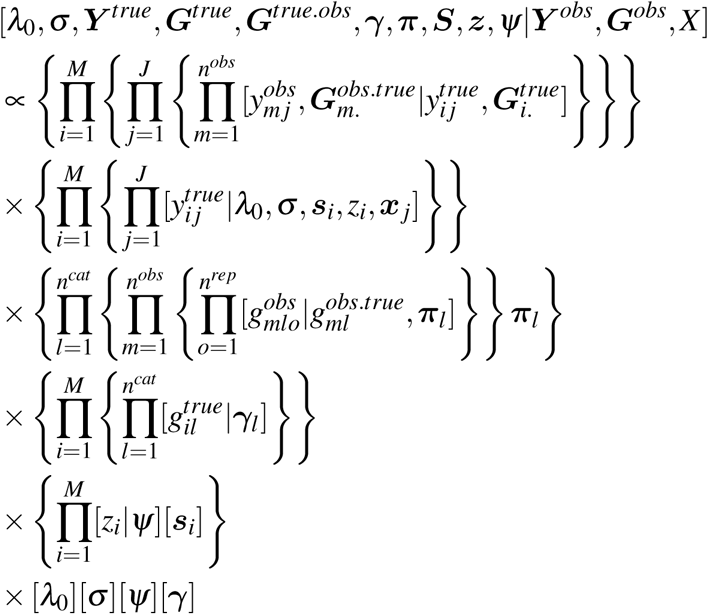

The prior distributions are

1. *p*(*λ*_0_) ∼ Uniform(0, ∞)
2. *p*(*σ*) ∼ Uniform(0, ∞)
3. *p*(*ψ*) ∼ Uniform(0, 1)
4. *p*(***s***_*i*_) ∼ Uniform(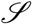)
5. *p*(***γ***_*l*_) ∼ Dirichlet(***α***_*l*_), where ***α***_*l*_ is the vector of Dirichlet parameters of length 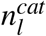, indexed by *g* below. All ***α***_*l*_ were set to vectors of 1.
6. 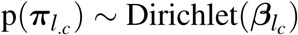, where 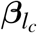 is the vector of Dirichlet parameters of length 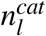, indexed by *r* below. All 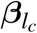 were set to vectors of 1.

Here, we list the full conditional distributions or the distributions that the full conditionals are proportional to, used to update the parameters and latent variables, that are unique to the categorical SPIM with observation error.

1. 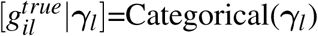
2. 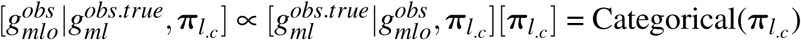 where 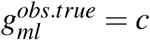.
3. 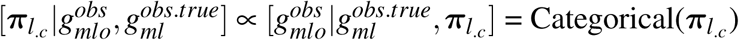 where 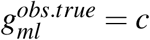.

Here we outline the new steps of the MCMC algorithm required to introduce observation error in the category levels. See Augustine et al. (2019) for the remaining parameter updates.

1. Update ***Y*** ^*true*^. The categorical SPIM uses a proposal distribution based on the SCR observation model to update ***Y*** ^*true*^ in a Metropolis-Hastings update where the only likelihood to be evaluated is the SCR observation model.

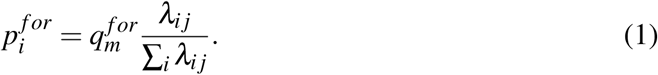

where 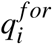 is the probability the randomly selected focal sample belongs to individual *i* at trap *j*. In fact, when the SCR observation model is Poisson, this proposal distribution is the full conditional distribution. With the introduction of category level observation error, the proposal distribution also needs to consider the proposal probabilities of different category level observation types and we need to evaluate the category level observation likelihood in the MH ratio. In order to update ***Y*** ^*true*^, on each iteration, we randomly select a user-specified number of indices *m* of ***Y*** ^*obs*^ to propose new individual identities for (because updating all individual identities on each iteration may not be most efficient). The current individual identity of 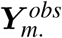 is stored in *ID*_*m*_, indicating which individual *i* to which sample *m* is assigned. So for each selected index *m*, we select a new individual *i* that we will propose for sample *m*. Once we select 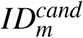, we construct 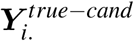 for 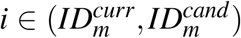 by moving sample *m* from individual 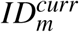 to individual 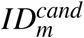 and set 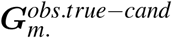 equal to 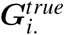 for 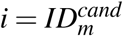. The forward proposal distribution for selecting 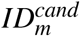 is:

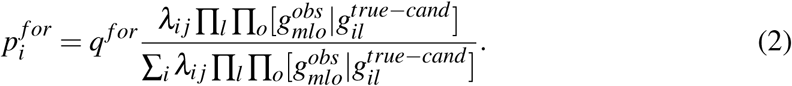

where *j* is the trap that focal sample *m* was recorded at and 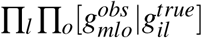 is the category level likelihood of the focal sample if it belonged to individual *i*. The backwards proposal probability is calculated as above, with the current and proposed arguments exchanged. We then accept the proposal with probability:

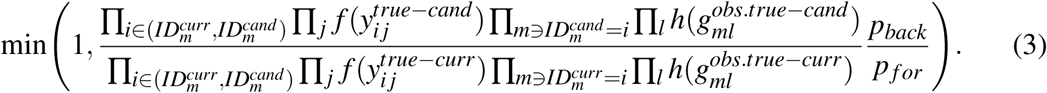

where *f* (·) is the SCR observation model likelihood and h(·) is the category level observation likelihood, and *p* _*f or*_ and *p*_*back*_ are obtained from the proposal distributions evaluated at 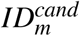 and 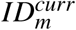, respectively. As in the categorical SPIM, this modified proposal distribution is the full conditional when the SCR observation model is Poisson.
2. Update ***G***^*true*^ and ***G***^*obs.true*^ jointly. In the categorical SPIM, latent values of 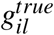 are updated using the full conditional Categorical(***γ***_*l*_) if 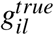 is latent and cannot be updated otherwise. 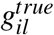 entries are not latent if at least one sample *m* is currently assigned to individual *i* whose *l*^*th*^ index of 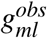 is not missing (*G*^*obs*^ is equivalent to *G*^*obs.true*^ in the current model). This can be described as requiring at least one *m* such that *ID*_*m*_ = *i*. In this model with observation error, we update 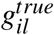 in the same manner when no samples *m* are assigned to individual *I* whose *l*^*th*^ index of 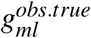 is not missing. However, once we introduce observation error, updating 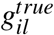 is possible when observed samples *are* assigned to it. More specifically, 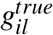 values are deterministically linked to 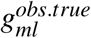 values for samples *m* currently assigned to individual *i* (for *m* ∋ *ID*_*m*_ = *i*). Because 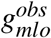 are observed with error, we can update both 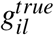 and its associated 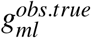 values, changing the category level observation likelihood in the process. We will use a Metropolis-Hastings update for these linked indices of 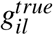 and 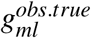 that are not currently unobserved. First, we propose 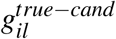 from Categorical(***γ***_*l*_), which also updates 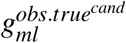 for *m* indicies currently assigned to individual *i*. The forward proposal probability, *p*^*f or*^, is then Categorical(***γ***_*l*_) evaluated at 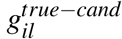 and the backwards proposal probability, *p*^*back*^ is Categorical(***γ***_*l*_) evaluated at 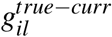. We accept the proposal with probability:

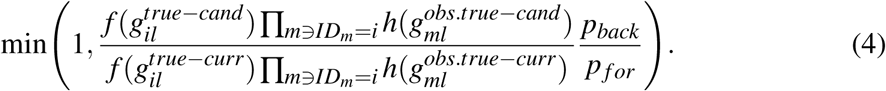

where *f* (·) is the category level likelihood and h(·) is the category level observation likelihood (distributions 1 and 2 listed above).
3. Update 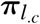. This update is for the general observation error model–see the next step for the Genotype SPIM update. Consider the following ***π*** structure for *n*^*cat*^ = 2 and *n*^*levels*^ = {2, 3}.

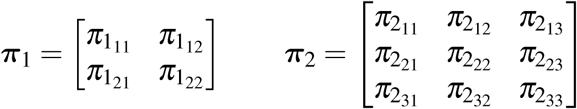 Column 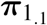 corresponds to the category level observation probabilities for identity covariate 1, which has 2 levels, conditional on the true value being 1, and column 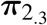 corresponds to the category level observation probabilities for identity covariate 2, which has 3 levels, conditional on the true value being 3. The diagonals of these matrices are the probabilities of correct observation. We use a standard Dirichlet-multinomial update (e.g., Wright et al. 2009). 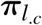 is updated with a Gibbs step. By adopting a Dirichlet prior for 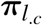, we get a Dirichlet full conditional with parameter vector 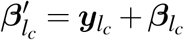 where 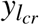 is the number of 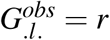 values for which 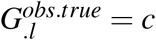 and 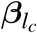 is the prior parameter vector. To draw values from the full conditional, we simulate a vector of Gamma random variables 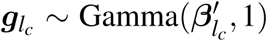, where 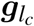 is of length 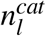. Then, after renormalizing these gamma random variables by 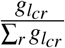, we have the Dirichlet full conditional.
4. Update ***p***^*hom*^ and ***p***^*het*^. For the Genotype SPIM, the elements of each ***π***_*l*_ are constrained by the correct classification, allellic dropout, and false allele probabilities. As in the more general model, we use a standard Dirichlet-multinomial to update ***p***^*hom*^ and ***p***^*het*^ independently, adopting independent Dirichlet priors for each. For ***p***^*hom*^, the Dirichlet full conditional parameter vector 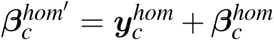 where 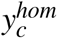 is the number of homozygous genotype correct classification events for *c* = 1 and the number of homozygous genotype false allele events for *c* = 2. For ***p***^*het*^, the Dirichlet full conditional parameter vector 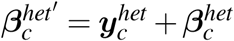 where 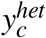 is the number of heterozygous genotype correct classification events for *c* = 1, the number of heterozygous genotype allelic dropout events for *c* = 2, and the number of heterozygous genotype false allele events for *c* = 3. 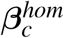 and 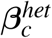 are the prior parameter vectors. After updating ***p***^*hom*^ and ***p***^*het*^, their values are plugged into the appropriate elements of ***π***. For the model with genotype observation probabilities that vary by sample type, we update the high and low sample quality parameters exactly as above using only the high and low quality samples, respectively.

---

| Term | Definition |
| --- | --- |
| $N$ | Population abundance |
| $J$ | The number of traps |
| $K$ | The number of capture occasions |
| $M$ | Level of data augmentation |
| $X$ | $J \times 2$ matrix of trap locations |
| $n^{obs}$ | The number of observed samples |
| $n^{cap}$ | The number of captured individuals (latent) |
| $n^{cat}$ | The number of categorical identity covariates |
| $n_l^{levels}$ | The number of category levels for categorical identity covariate $l$ |
| $\gamma_l$ | A vector of category level probabilities for categorical identity covariate $l$, corresponding to values $1, \dots, n_l^{levels}$ |
| $\psi$ | The data augmentation individual inclusion probability |
| $\lambda_0$ | The baseline detection rate (count data) |
| $p_0$ | The baseline detection probability (detection data) |
| $\sigma$ | The detection function spatial scale parameter |
| $\lambda_{ij}$ | The individual by trap detection rate |
| $p_{ij}$ | The individual by trap detection probability |
| $z$ | The data augmentation indicator vector of length $M$ |
| $S$ | $M \times 2$ matrix of individual activity centers |
| $G^{true}$ | An $M \times n^{cat}$ matrix of full categorical identities |
| $Y^{true}$ | The latent $M \times J$ capture history |
| $Y^{obs}$ | The observed $n^{obs} \times J$ capture history, with one capture per row indicating the trap of capture |
| $n^{reps}$ | The number of independent, replicate assignments of the categorical identity covariates |
| $G^{obs.true}$ | An $n^{obs} \times n^{cat}$ matrix of true, latent sample-level categorical identities |
| $G^{obs}$ | An $n^{obs} \times n^{cat} \times n^{obs}$ array of sample-level observed (possibly erroneously) categorical identities |
| $\pi_l$ | An $n_l^{levels} \times n_l^{levels}$ matrix of observation probabilities for covariate $l$ conditional on the true category level. The diagonal elements represent correct observation probabilities and off-diagonal elements represent erroneous observation probabilities |
| $ID$ | A derived vector of length $n^{obs}$ recording the individual index $i$ to which the $m^{th}$ row of $Y^{obs}$ and $G^{obs.true}$ are linked |
| $p^{hom}$ | A vector of genotype observation probabilities that determine the element values of $\pi$ for homozygous loci-level genotypes. $p^{hom} = (p_C^{hom}, p_{FA}^{hom})$, the probabilities of correct and false allele observation, respectively. |
| $p^{het}$ | A vector of genotype observation probabilities that determine the element values of $\pi$ for heterozygous loci-level genotypes. $p^{het} = (p_C^{het}, p_{AD}^{het}, p_{FA}^{het})$, the probabilities of correct, allelic dropout, and false allele observation, respectively. |

**Figure A1:**
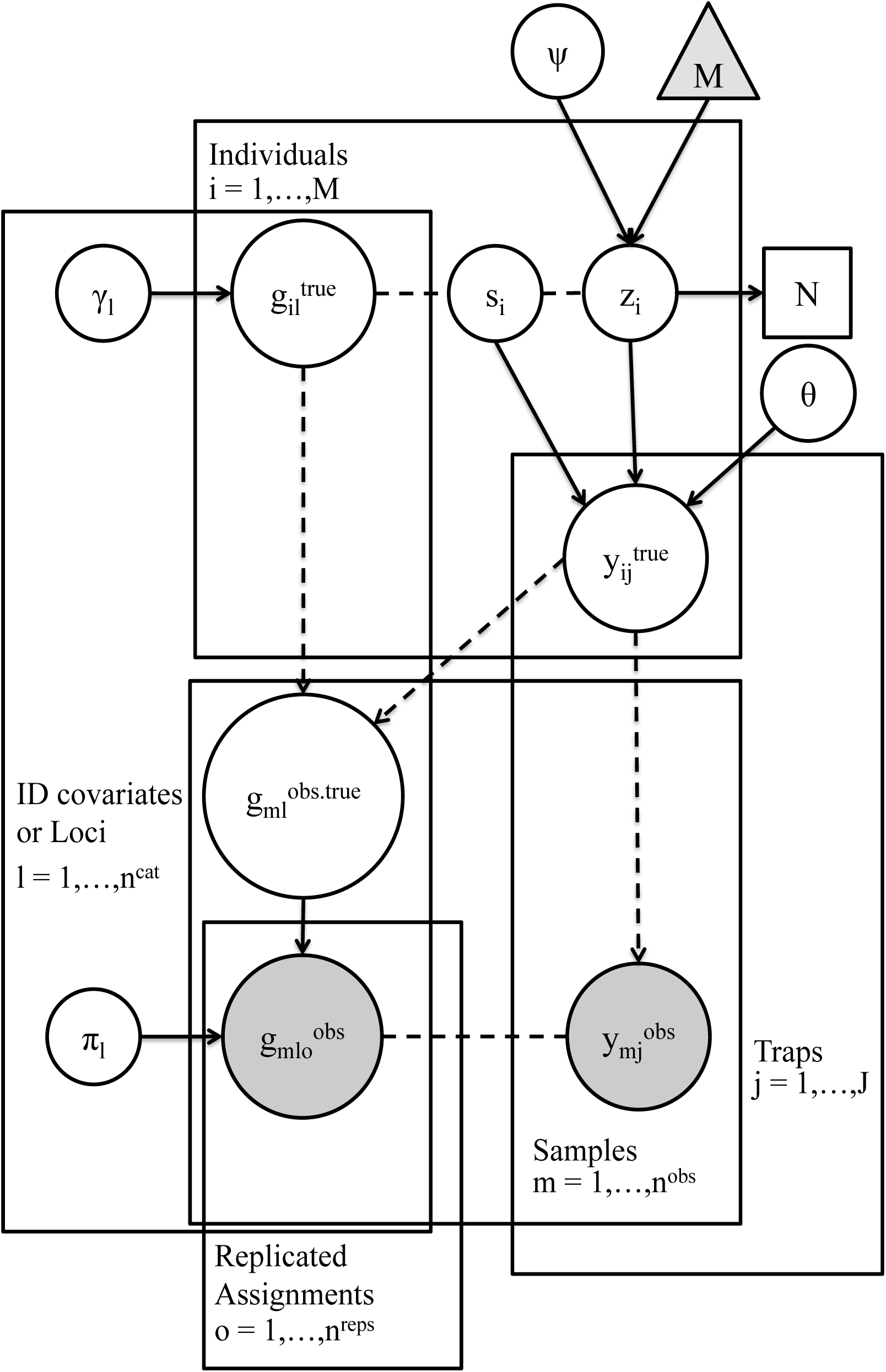
Directed acyclic graph of the catSPIM-OE model. Dashed lines represent objects linked along the first dimension and dashed arrows represent disaggregation of data from the individual level to the sample level. *θ* represents the detection parameters. All other terms are described in the table above.

## Appendix B: MCMC Details for Fisher Data Set

### MCMC Specifications

For the SCR analysis, we ran 5 MCMC chains for 200,000 iterations, thinning by 10, and discarding the first 25,000 of each chain as burn in, leaving 87,500 samples for posterior inference. Due to convergence problems with the Genotype SPIM for this data set, we initialized the samples for one particular individual with a long-distance spatial recapture to belong to the same individual (discussed further below). After this deviation from the typical algorithm for initializing all latent states, we ran 40,000 iterations as burn in across 5 chains, followed by 240,000 iterations for each chain, thinned by 10, leaving 120,000 iterations for posterior inference. For both analyses, we used a continuous, polygonal state space with a minimum distance between polygon vertices and traps of at least 3*σ*. We set the data augmentation level, *M*, to 5500 in the SCR analysis and 5000 in the SPIM analysis (lower due to greater precision for *N*).

Due to the low number of spatial recaptures, we put an informative prior of Gamma(3,7) on *σ*^*sd*^, the standard deviation of the individual-specific detection function spatial scale parameter, which ruled out most unreasonably large point estimates of *σ*_*i*_. We used the uninformative priors listed in Appendix A for the remaining parameters in both models, including a Uniform(0,∞) prior for *σ*.

### Computation Time

Both the SCR and SPIM analyses required long computation times, mostly due to the large data augmentation level, a large number of traps, and the slow mixing of the *σ*^*sd*^ parameter, which required more posterior samples than would be required for a model without individual heterogeneity in detection function parameters. The MCMC samplers for both models completed roughly 1000 iterations per hour on a machine with a 2.2GHz processor. The simpler SCR MCMC sampler was about as fast as the more complicated Genotype SPIM sampler because the lower precision for abundance required a larger level of data augmentation (M=5500 vs 5000). The long run times per chain were offset substantially by the use of multiple chains on multiple cores. Recording all posteriors for the Genotype SPIM model (especially *G*^*true*^) could potentially use a prohibitive amount of RAM, but this was ameliorated by thinning the posterior and running the sampler for a relatively short number of iterations (40,000), saving the posterior, and then restarting the MCMC chain from the previous final state.

### Convergence Issue

The model using the fisher data set and our typical algorithm for initializing the parameters and latent variables did not converge. What we mean when we say that the model did not converge is that all of the samples for one particular individual that were assigned the same individual identity in the original study were not linked by the Genotype SPIM and the reason for the failed linkage was not due to the observed genotypes of the samples–all 5 samples in question had matching consensus genotypes. We argue that the failure of the Genotype SPIM to link these samples to the same individual is a violation of the SCR observation model assumptions.

The algorithm we use to initialize the latent variables, specifically, how we initialize samples to individuals to create the starting latent capture history can be seen as starting from an “overdispersed” state in the sense that we start from a very unlikely assignment of samples to individuals with respect to the expected true *σ*. We initialize the latent capture history by only linking samples with similar observed genotype scores *at the same trap*, implying that *σ* is very small because there are no spatial recaptures in this state. Then for the model to converge, the spatial recaptures are gradually linked up, or at least linked with some probability, and the *σ* value converges upwards. This did not occur for the focal individual in question within the 800,000 iterations that we ran starting from this overdispersed state.

It is possible the Genotype SPIM was correct in not linking up these 5 samples and that 2 individuals in the population had the same genotype–a shadow event. We do not believe this is the case. First, while this is the longest distance spatial recapture if all 5 samples belong to the same individual, it is of a similar distance as other observed spatial recaptures (Figure B1). Therefore, if these samples do belong to two individuals, they must live very close to one another, which is less likely under our model assumptions than two individuals with the same genotype living anywhere in the state space.

Figure B2 provides some clues about why these samples are not being linked to the same individual. Depicted there are the activity center posteriors from a 40,000 iteration subset of the MCMC chain that we claim has not converged. It links the 4 samples in trap 165 to one proposed individual and the 1 sample in trap 179 to another proposed individual. Notice that the activity center posterior for the proposed individual linked to trap 165 is much more spatially constrained than the activity center posterior for the proposed individual linked to trap 179. Further, notice that these posteriors do not overlap in space. In order for these 5 samples to be combined into 1 individual, the activity centers of proposed individuals 1 and 2 must overlap in space. For example, in order to have a non-negligible probability of moving the 1 sample at trap 179 to the same individual with 4 samples at trap 165, the activity center for individual 1 in the current configuration must be a similar distance from trap 179 as the activity center for individual 2 in the current configuration. This is required because the expected number of detections is a function of the distance between the activity center and the trap.

The differential spatial extents covered by the activity center posterior distributions for these two proposed individuals is a consequence of the individual heterogeneity model for the detection function parameters where *σ*_*i*_ is inversely related to 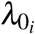 for individual *i*. Because proposed individual 1 was captured 4 times in trap 165, it must have a larger 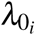 than proposed individual 2 captured 1 time in trap 179. This necessarily implies proposed individual 1 must have a smaller *σ*_*i*_. As a consequence of the smaller *σ*_*i*_, the activity center for proposed individual 1 is estimated much more precisely. The estimated detection functions for these two proposed individuals are displayed in Figure B3. There, we see that the activity center for proposed individual 2 must move within a distance of roughly 2 km to trap 179 before it has any chance that the sample at the trap will be combined with the other 4 samples. Where the two detection functions cross is the distance at which the 1 sample at trap 179 would be equally likely to belong to proposed individual 1 or 2 if both their activity centers were exactly this distance away from trap 179. This never occurs in Figure B2.

Now, consider what *σ*_*i*_ and 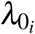 would be when these 5 samples are allocated to 1 individual if all model assumptions are met. The activity center would necessarily be somewhere between traps 165 and 179, though closer to trap 165 because there are more captures there. In order for these samples to be linked, *σ*_*i*_ must be relatively large. But the expected 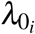 corresponding to a large *σ*_*i*_ is relatively small. The 4 detections at trap 165 are then very improbable. This apparently unlikely set of events could be due to this individual’s detection function deviating from the assumption that detection is exactly proportional to space use, or it could be because the Poisson model for the expected number of detections as a function of the distance between the activity center and the trap is not exactly correct. There could be overdispersion in the expected counts, for example, because factors other than the distance between the activity center and trap determine the observed count. In this survey, there were 3 week long capture occasions and the 4 samples at trap 165 were observed as a count of 2 on occasions 2 and 3. If the difference between a count of 1 and 2 is at least partially influenced by the behavior of an individual encountering a hair snare, rather than the distance between the activity center and the trap, overdispersion in the Poisson observation model should be expected. Perhaps a better SCR observation model is one where detection is a function of the distance between an activity center and a trap and the number of detections given at least 1 detection is not a function of the distance between an activity center and a trap (e.g., a hurdle model).

One final note is that if the violations of the SCR observation model do explain the lack of convergence for the Genotype SPIM for the fisher data set, they may not prevent convergence if the two traps in question were closer together in space. It may be the combination of a long distance spatial recapture with the violation of the SCR observation model that led to this convergence problem. Regardless, when we initialized all 5 of these samples as belonging to the same individual, they remained in this state for 240,000 iterations across each of 5 MCMC chains, so it appears the problem leading to the lack of convergence did not prevent us from sampling from the true posterior when we initialized the latent variables closer to the presumably true values. Once these samples are connected to one individual, it is extremely unlikely that a second individual with the exact same genotype will be proposed, much less one that is close enough to these samples to split them across two individuals. Further, no convergence problems were identified in the simulation study where we know all model assumptions were met.

### Figures

**Figure B1:**
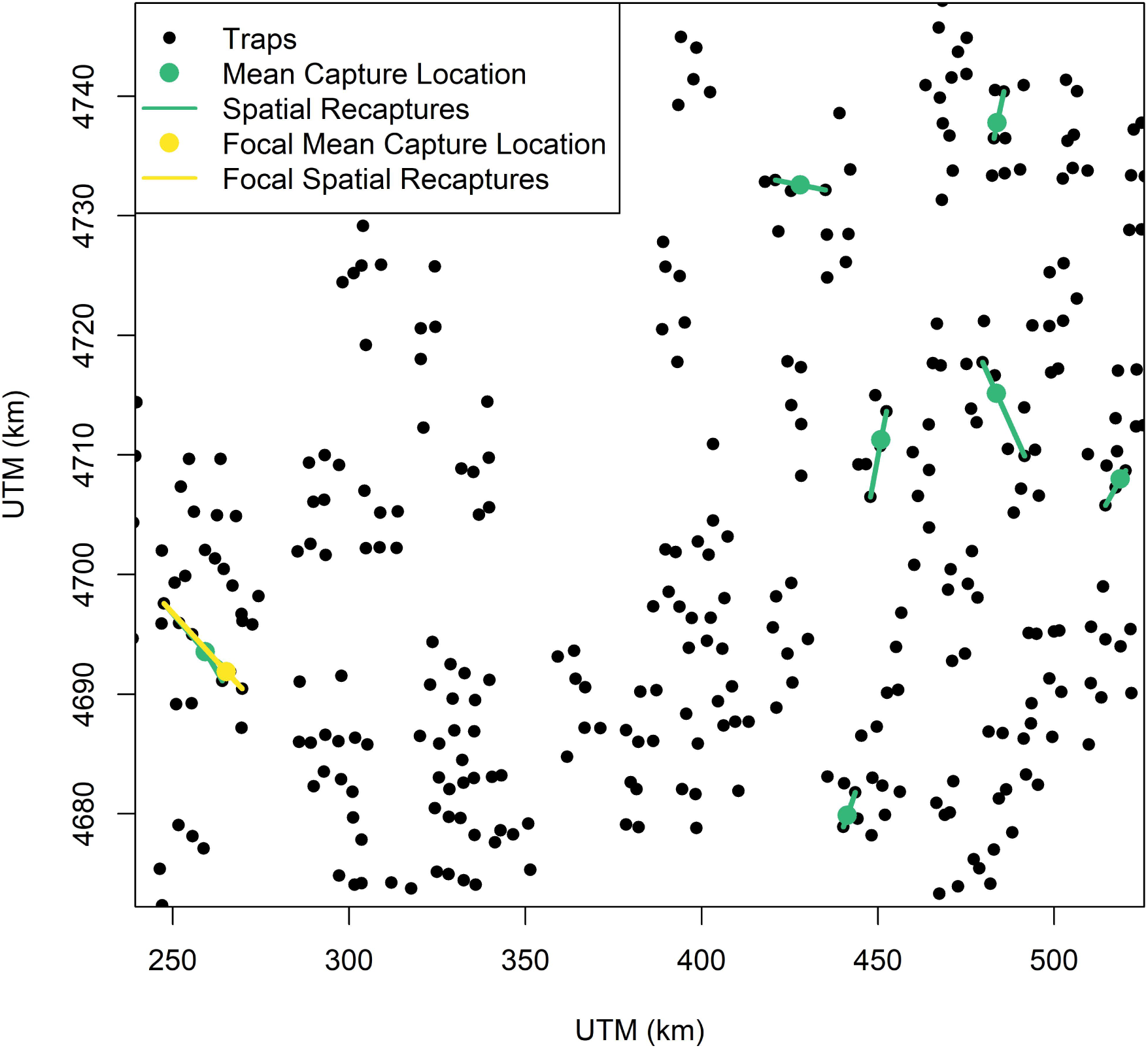
The mean capture locations and spatial recaptures for the 8 individuals with spatial recaptures in the original fisher data set, with the focal individual responsible for the lack of convergence distinguished from the other individuals in yellow.

**Figure B2:**
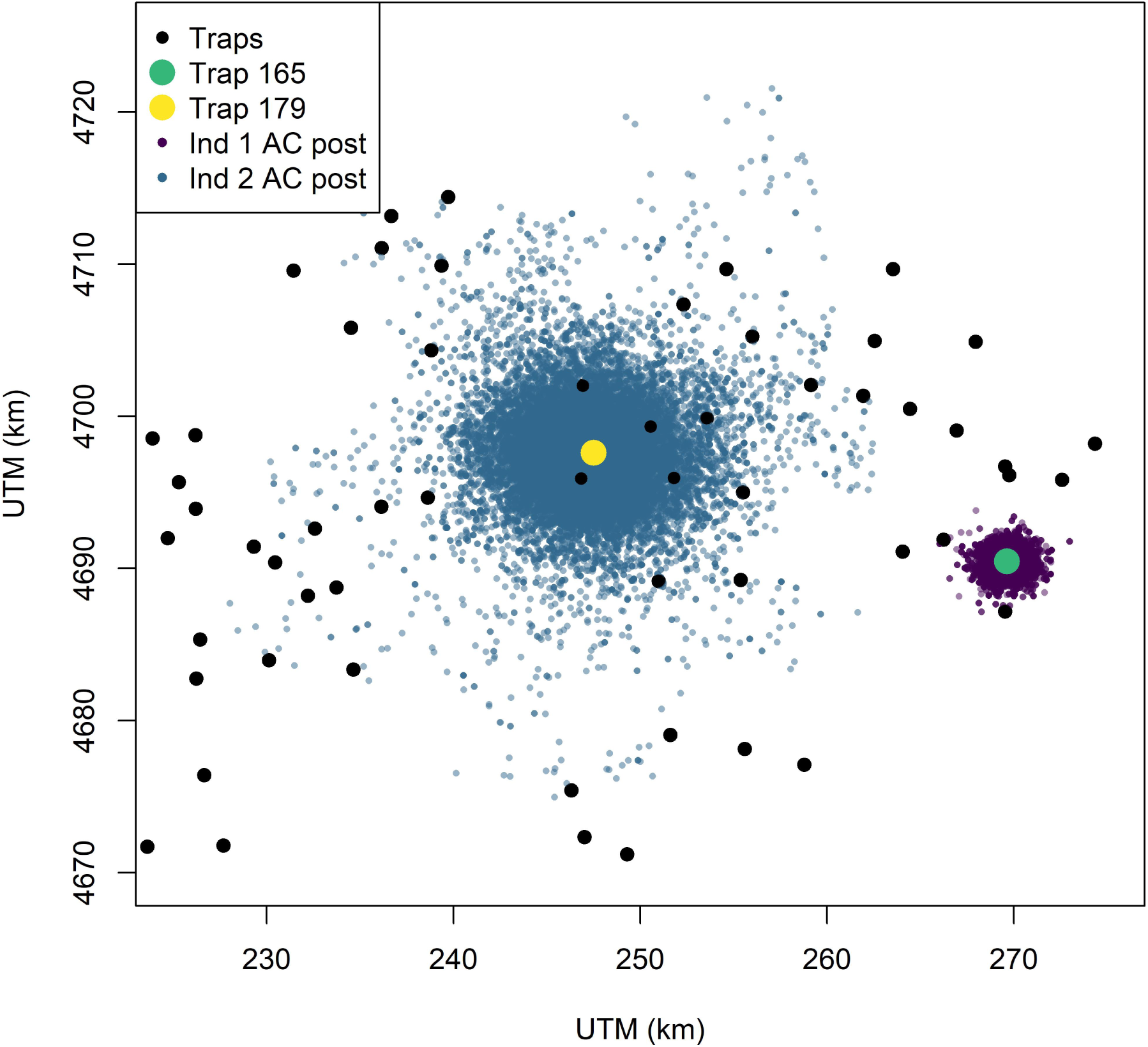
The activity center posteriors for the samples in question at traps 165 and 179 when they are split between two individuals. There were 4 samples observed at trap 165 and 1 at trap 179.

**Figure B3:**
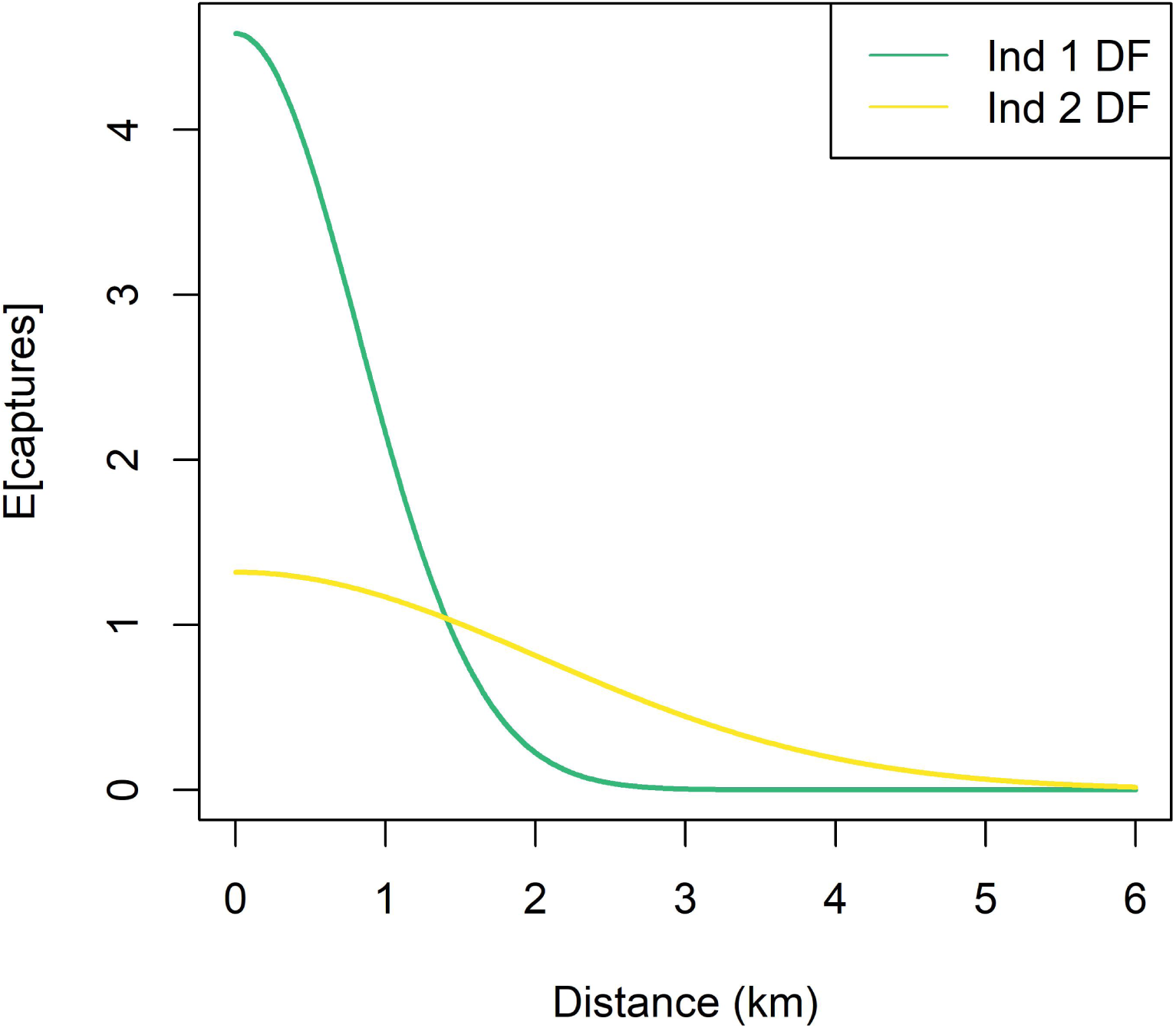
The estimated detection functions for the 2 proposed individuals in question.

## Appendix C: Simulation Study Details

We evaluated estimator performance using a simulation study in which the data-generating parameter values were set to those estimated in the fisher analysis, except for density which we set to a larger value to make resolving the individual identities of samples more challenging (Augustine et al. 2019). We set *σ* to 1.32 km and converted the fisher *a*_0_ estimate of 4.73 to a *λ*_0_ value of 0.570. We quadrupled the fisher density estimate of 4.27/100 km^2^ to 17.08/100 km^2^. We specified a 9 x 9 rectangular trapping array of 81 traps with a 2*σ* spacing of 2.64 km and a buffer of 3*σ* to define the rectangular state space. The abundance implied by the density and state space area stated above was 166 individuals.

We simulated from the same genotype frequency and genotype observation models as used for the fisher data set using the parameter estimates from that analysis. We simulated “high” and “low” quality samples in the proportion of 52:48, similar to that observed in the fisher data set. We set the locus by replication amplification probability (not an explicit parameter in our model) to 0.995 and 0.437 for high and low quality samples, respectively, matching that observed on average across each sample type for the first 3 replicated assignments in the fisher data set. We used the parameter estimates for ***γ, p***^*hom*-*high*^, ***p***^*hom*-*low*^, ***p***^*het*-*high*^, ***p***^*het*-*low*^ from the fisher data set to simulate from. We considered a maximum of *n*^*rep*^ = 3 replicated assignments, fewer than the maximum of 7 in the fisher data set, though the probability of loci amplification declined after 3 replicated assignments for the fisher data set.

We simulated 120 data sets from the Genotype SPIM and fit the Genotype SPIM model using 1, 2, and 3 of the replicated assignments, with and without the low quality samples. Then, we fit a regular SCR model using only the “high quality” samples, to roughly mimic what happens in practice where only high confidence samples are retained, though we assumed a best case scenario where there were no errors in individual identity assignment. For all models, we ran 2 chains of 100,000 iterations, discarding 5,000 iterations as burn in and thinning by 50 to reduce output file sizes. We used the posterior modes for point estimates except for parameters that sum to 1 (e.g., genotype observation probabilities) for which we used the posterior means. We used 95% percentile intervals for interval estimates, which had better coverage than HPD intervals for *n*^*cap*^ because it is a discrete parameter with very low uncertainty leading to very narrow intervals. Because all 4 of these estimators should be approximately unbiased, we used the posterior standard deviations and mean 95% CI widths to compare precision. We used the mean squared error to compare accuracy. The uninformative priors listed in Appendix A were used for all parameters.

The analysis time for the simulated data sets was much less than for the fisher application because we did not consider individual heterogeneity in detection function parameters; we also considered a much lower abundance and number of traps, though the expected number of detections per individual was higher. A run of 100,000 iterations took roughly 8 hours, while the same number of iterations for the fisher application took roughly 4 days.

**Table 1:** Simulation settings for baseline detection rate, *λ*_0_, detection function spatial scale, *σ*, abundance, *N*, number of replicated genotype assignments, *n*^*rep*^, and whether or not to use the low quality samples with lower amplification rates and higher error rates.

| Scenario | $\lambda_0$ | $\sigma$ | $N$ | $n^{rep}$ | Use Low Quality Samples |
| --- | --- | --- | --- | --- | --- |
| SPIM3A | 0.570 | 1.320 | 166 | 3 | Yes |
| SPIM2A | 0.570 | 1.320 | 166 | 2 | Yes |
| SPIM1A | 0.570 | 1.320 | 166 | 1 | Yes |
| SPIM3B | 0.297 | 1.320 | 166 | 3 | No |
| SPIM2B | 0.297 | 1.320 | 166 | 2 | No |
| SPIM1B | 0.297 | 1.320 | 166 | 1 | No |
| SCR | 0.297 | 1.320 | 166 | . | No |

## Appendix D: Black Bear Application

Here, we will provide an application of the Genotype SPIM to a data set with no replicated assignments (only 1 PCR), or at least any replicated assignments were not provided by the lab that did the genotyping (Wildlife Genetics International; WGI). WGI uses the methods described in Paetkau (2003) where 1/3 of the DNA product of each sample is amplified in a single PCR. Poorly performing samples are then culled and any remaining samples with 1–2 mismatched pairs are reanalyzed. Therefore, we do not know how many replicated assignments there were for each sample. We will ignore this problem in the analysis here, admitting that it amounts to a misspecification of the genotyping error model, though this may not be of great consequence. Since we will estimate that the certainty in individual identity is 100% for the high quality samples in this data set, we are really only concerned with the number of times the poor quality samples were genotyped. It is likely that the majority of these samples were genotyped only once and then deemed unreliable because they produced partial or no genotypes. This application is presented as a proof of concept that the Genotype SPIM can work with no replicated assignments for data sets collected in practice.

### Survey Description

This data set comes from Murphy et al. (2016). We will describe the most pertinent details of the survey here and direct readers to the original publication for the full methods. This data set comes from the second year at one site of a 3 year (2011–2013), 4 site hair snare survey for black bears along the Kentucky-Virginia, USA border. We chose to use the site that encompasses parts of Pine and Black Mountains because it had the larger spatial extent of traps, encompassed the primary core of the population, and thus produced more captures and spatial recaptures. We chose the second year (2012) because it had the most samples that were not assigned individual identities by WGI.

The survey effort consisted of 81 hair snares with an average spacing of 1.6 km, operated over 8 consecutive 1-week long capture occasions. A total of 340 samples were collected and sent to WGI for microsatellite genotyping. To reduce the costs of genotyping, only 1 sample per trap per occasion was selected for genotyping, following sample randomization within each trap x occasion combination. A total of 179 samples remained after subsampling, for which genotyping was attempted using microsatellite markers G10H, G10L, G10M, MU23, G10J, G10B, G10P, and a sex marker. One hundred fifty-four samples were assigned an individual identity (63 individuals total), leaving 25 samples that were unassigned to individual. Of these originally discarded samples, 18 provided at least 1 scored locus, leaving 7 with no partial identity information at all, other than the location where the sample was collected. There were 1, 1, 1, 2, 6, and 7 samples with 6, 5, 4, 3, 2, and 1 scored loci, respectively. In total, there were only 40 observed loci among the 18 partial genotype samples, or on average, 2.2 loci per sample. This amounted to a 16% increase in total samples and a 3% increase in samples with 3 or more loci.

### Statistical Methods

As with the Fisher application in the main text, we fit the Genotype SPIM using all 179 samples and fit a regular SCR model using the 154 samples assigned individual identities by WGI for comparison. We used a Bernoulli observation model instead of the Poisson because the subsampling process for the hair samples ensured that a maximum of 1 sample per individual could be collected per trap on each occasion. Treating these subsampled detection data as Bernoulli was previously demonstrated via simulation to be appropriate by Murphy et al. (2016), which differs from typical single-catch traps because the capture order of individuals does not influence the subsampling protocol. For both the Genotype SPIM and SCR models, we used the individual heterogeneity detection function model described in the main text and we used a Gamma(3,7) prior for *σ*^*sd*^, the parameter determining the level of individual-level variability in *σ*_*i*_ and *λ*_*i*_. For the Genotype SPIM, we split samples into “high” and “low” quality categories, determined by whether or not WGI assigned them an individual identity, and estimated separate genotype observation parameters for each quality category.

Unlike the fisher application in the main text, we included the sex marker as a partial identity covariate that could be observed with error. We used the same genotyping error model for this covariate as we did for the other microsatellite markers. This treatment introduces a misspecification of the genotyping error process for the sex marker, but it should have a negligible impact on this analysis for two reasons. First, the female sex marker has a value of 250.250, while the male marker has a value of 204.250. We assumed that the value 204.204 was also in the population, but because 204.204 was never observed, the frequency of this marker is estimated at <1%. Second, we assumed that the mechanisms that lead to allelic dropout and false alleles in microsatellites also apply to this marker, but this marker presumably has different error mechanisms and a lower rate of error. Thus, this misspecification likely decreases the estimated rate of allelic dropout and false allele events which apply to the other microsatellites. However, the high quality samples were estimated to be genotyped with near certainty and only one poor quality sample was sexed, limiting the effect of this misspecification for the low quality samples of interest.

For the regular SCR model, we ran 3 chains for 200,000 iterations, thinned by 50, and discarded the first 5000 samples of each chain as burn in. For the Genotype SPIM, we ran 1 chain for 800,000 iterations, thinned by 50, and discarded the first 325,000 as burn in. We present posterior modes as point estimates, except for parameters that sum to 1 (e.g., genotype observation probabilities), where we used the posterior mean. We present 95% highest posterior density (HPD) intervals as interval estimates. We use the coefficient of variation (CV) to compare the precision of each model, which we define as the posterior standard deviation divided by the posterior mode.

### Results

The Genotype SPIM and SCR models produced nearly equivalent estimates for abundance (Figure D1, Table D1). The Genotype SPIM abundance point estimate was 2.7% lower than the SCR estimate, with a CV that was 1.1% lower, demonstrating that the partial genotype samples very slightly improved precision. Including the low quality samples in the Genotype SPIM increased the overall detection parameter, *a*_0_ from 1.42 to 1.81 and the estimated number of individuals captured was raised to 69, compared to the 63 available in the high quality samples. The Genotype SPIM produced slightly more precise estimates of the both spatial scale parameters, *σ* and *σ*^*sd*^, likely due to the increased number of spatial recaptures. There were thirty-five spatial recaptures in the high quality samples used in the SCR analysis and an estimated 50 when including the low quality samples in the Genotype SPIM analysis. The minimum value of the Genotype SPIM posterior for the number of spatial recaptures was 43, indicating that the poor quality samples added 7 spatial recaptures with near certainty.

The high quality samples were estimated to have a nearly perfect genotype correct observation probability, and the low quality samples were estimated to be relatively reliable, but with a large amount of uncertainty (Figure D1, Table D2). We compare 12 of the 18 partial genotype samples with their highest posterior probability matches to complete genotype samples in Table D3. Five partial genotype samples were matched with individuals in the complete genotype samples with a probability of greater than 0.97 (Table D3). These samples were scored at 2-6 loci. Generally, samples scored at 2 or fewer loci did not provide high probability matches. Samples with 1 or 2 scored loci were more likely to match more than one individual with similar probabilities (e.g., samples 8, 18, and 19). Of the 12 samples displayed in the table, 11 did not contain any genotyping errors when associated with their highest probability matches. Partial genotype sample 147 was scored at 4 loci and did not match any complete genotype individual at all loci. It did match a complete genotype individual at 2 loci, which was captured at the same trap. This assignment was given a 0.20 probability and implied 2 false alleles, though the event of a correct assignment of no match was assigned a higher probability 0.80.

### Discussion

We demonstrated that the Genotype SPIM can work with no replicated assignments for a real world data set. In this case, there was only a negligible improvement in the inference about abundance; however, this data set contained very minimal information in the added low quality samples–only 40 scored loci in total and only a 3% increase in samples when considering the more informative samples scored at 3 or more loci. Further, the paucity of partial genotype scores led to very imprecise estimates of the poor quality sample genotype observation probabilities (Figure D2). It is likely for this reason that sample 147 had a non-negligible posterior probability (0.20) of matching a complete genotype sample with 2 false allele scores captured at the same trap. Interestingly, the presumably correct assignment of no match was more probable (0.80). If no genotype information was available at all as is the case with unmarked SCR (Chandler and Royle 2013), it is likely the incorrect assignment would be made with a higher probability as these two samples were the only ones recorded at this trap. We expect that if more partial genotype samples were available, the false allele probability would be estimated more precisely around a very low value and this presumably incorrect match would be less likely or ruled out. Still, we believe the Genotype SPIM assignments like this made with very imprecise error probability estimates will be an improvement over a spatial partial identity model that discards this information (e.g., the “random thinning” model discussed in the main text).

The high quality samples for this bear data set were estimated to be scored correctly with near certainty, compared to a 0.185 estimated allelic dropout rate per replicated assignment for the fisher data set. In fact, this bear data set was estimated to contain 0 errors with probability 1. While there may have been differences in the DNA quantity and quality between the fisher and bear hair samples, the lab methods used by WGI (Paetkau 2003) likely reduced the error rate by including more DNA product in the first and often only PCR, compared to the modified multi-tubes approach (Frantz et al. 2003) used for the fisher data set, requiring the DNA product to be split across up to 7 PCRs (Waits and Paetkau 2005). We expect the multi-tubes approach to yield more reliable estimates from the Genotype SPIM due to the explicit replicated assignments; however, it is more expensive (Waits and Paetkau 2005). The reliability of the Genotype SPIM estimates using data from WGI or other labs using the methods of Paetkau (2003) could be improved by using the replicated assignment data for the samples where multiple PCRs were done.

## Acknowledgements

We thank Sean Murphy and John Cox for making this black bear data set available for us to use.

## Figures

**Figure D1:**
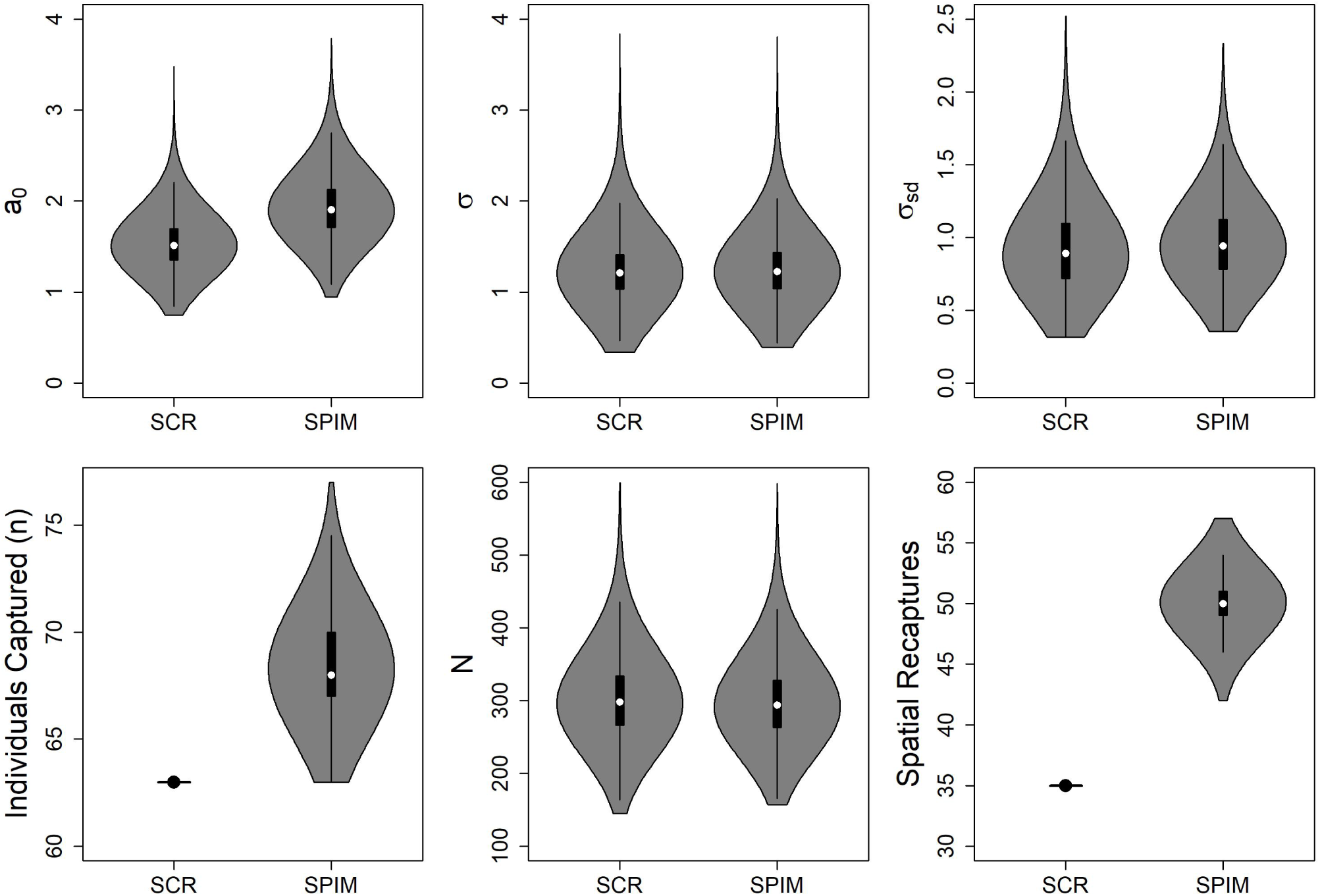
SCR process and observation model posterior distributions from the SCR and SPIM analyses. *a*_0_ is the overall detection parameter, *σ* is the population-level detection function spatial scale parameter in km, *σ*^*sd*^ is the standard deviation of the individual-level variance in the spatial scale parameter, *n* is the number of individuals captured, and *N* is the population abundance. The number of captured individuals and spatial recaptures in the SCR analysis are known statistics.

**Figure D2:**
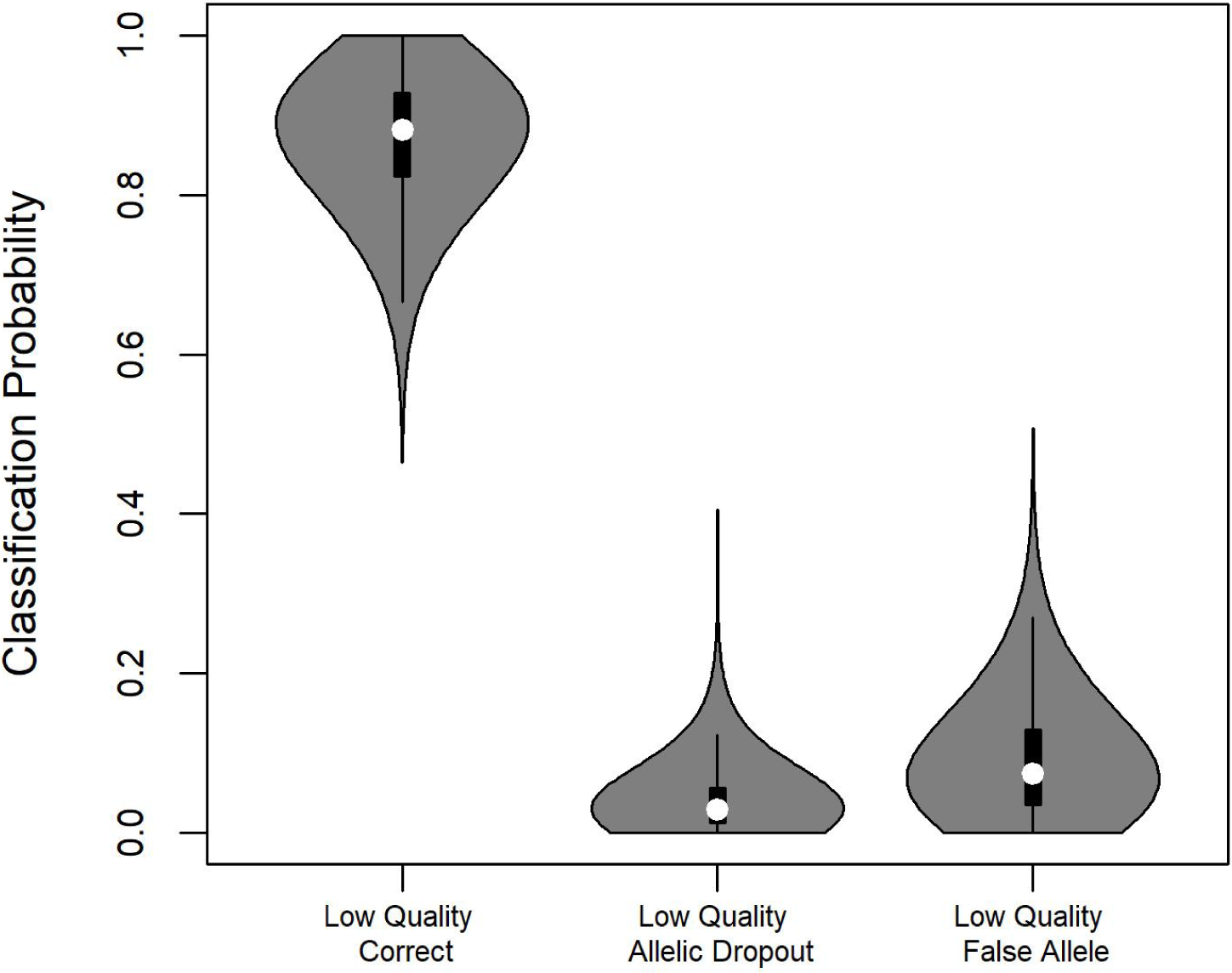
Heterozygous single locus genotype observation probability posterior distributions for low quality samples. See Table 2 for full observation probability results.

## Tables

**Table D1:** SCR process and observation model parameter estimates from the SCR and SPIM analyses. *a*_0_ is the overall detection parameter, *σ* is the population-level detection function spatial scale parameter in km, *σ*^*sd*^ is the standard deviation of the individual-level variance in the spatial scale parameter, *N* is the population abundance, *n* is the number of individuals captured, and *D* is the population density (individuals/100km^2^). Posterior modes are presented as point estimates, posterior standard deviations/posterior modes are presented as the coefficient of variation, and 95% HPD interval upper and lower bounds are presented as interval estimates.

|  | SCR |  |  |  | SPIM |  |  |  |
| --- | --- | --- | --- | --- | --- | --- | --- | --- |
|  | Est | CV | LB | UB | Est | CV | LB | UB |
| $a_0$ | 1.42 | 18.2 | 1.07 | 2.06 | 1.86 | 17.0 | 1.35 | 2.57 |
| $\sigma$ | 1.14 | 28.7 | 0.61 | 1.90 | 1.20 | 27.5 | 0.63 | 1.90 |
| $\sigma^{sd}$ | 0.75 | 38.1 | 0.44 | 1.49 | 0.86 | 30.3 | 0.49 | 1.46 |
| $N$ | 294 | 18.1 | 203 | 406 | 279 | 18.1 | 203 | 397 |
| $n$ | 63 | | | | 68 | 2.7 | 64 | 71 |
| $D$ | 17.99 | 18.1 | 12.43 | 24.86 | 17.08 | 18.1 | 12.49 | 24.37 |

**Table D2:** Single locus genotype observation probability parameter estimates for high and low quality samples. “Het” indicates a heterozygous single locus genotype and “hom” indicates a homozygous single locus genotype. “Correct”, “AD”, and “FA” indicate a correct, allelic dropout, and false allele observation, respectively. Posterior modes are presented as point estimates and 95% HPD interval upper and lower bounds are presented as interval estimates.

| Class Type | High Quality |  |  | Low Quality |  |  |
| --- | --- | --- | --- | --- | --- | --- |
|  | Est | LB | UB | Est | LB | UB |
| Het-Correct | 0.997 | 0.993 | 1.000 | 0.870 | 0.722 | 0.995 |
| Het-AD | 0.001 | 0.000 | 0.004 | 0.040 | 0.000 | 0.117 |
| Het-FA | 0.001 | 0.000 | 0.004 | 0.090 | 0.000 | 0.228 |
| Hom-Correct | 0.996 | 0.988 | 1.000 | 0.910 | 0.745 | 1.000 |
| Hom-FA | 0.004 | 0.000 | 0.012 | 0.090 | 0.000 | 0.255 |

**Table D3:**
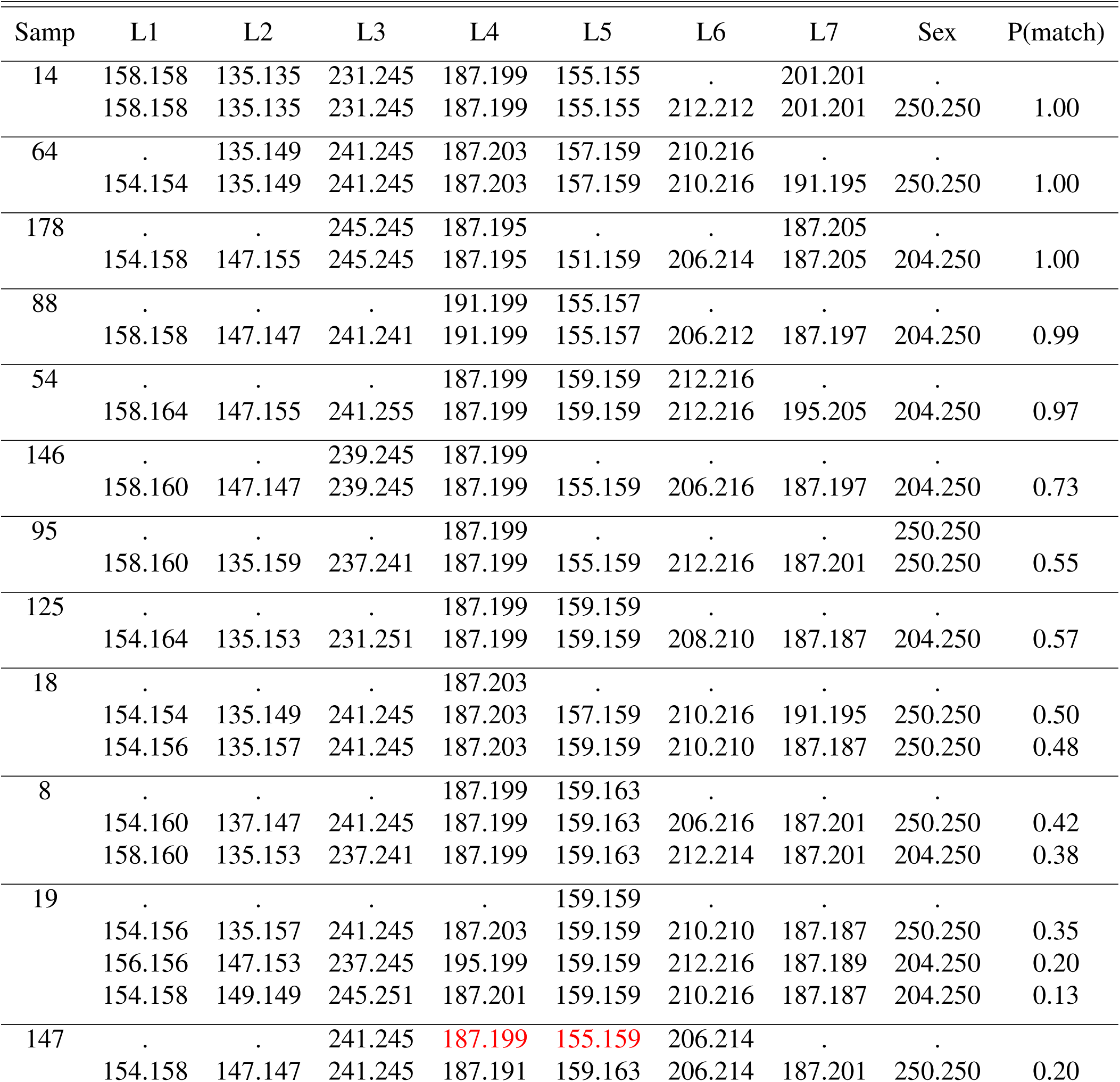
Twelve of the 18 partial genotype samples with compared to their highest posterior probabilities of matches among the complete genotype samples. The observed values at 7 microsatellite loci (L1 - L7) and the sex marker for the partial genotype sample are listed followed by the values for the complete genotype samples they matched with. P(match) is the posterior probability of the pairwise match between the partial and complete genotype sample. Mismatches are colored in red.

## Appendix E: General Observation Error Model Simulations and Andean Bear Application

In this Appendix, we provide a short simulation study of the catSPIM with general observation error (catSPIM-OE) described in Appendix A, followed by an application to an Andean bear camera trap data set where facial markings are used as categorical identity covariates.

### Simulation Study

#### Simulation Specifications

We conducted a simulation study to demonstrate the performance of catSPIM-OE, compare it to catSPIM without observation error, and illustrate the effects of varying the number of categorical identity covariates, *n*^*cat*^, and the number of replicated assignments, *n*^*reps*^. We replicated Scenario A2 from Augustine et al. (2019), which was the most optimal scenario considered in terms of expected catSPIM performance. In this scenario, the values of population density and *σ* led to the least amount of home range overlap across individuals considered and thus, the least amount of uncertainty in individual identity. We recreated this scenario using a 9 × 9 grid of traps with unit spacing and a 3*σ*. We set *N* = 39, *λ*_0_ = 0.25, *σ* = 0.5, and *K* = 10. We considered 2-5 2-level identity covariates, 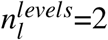, for *l* ∈ (2, 3, 4, 5) with correct observation probabilities for each category level of 1, 0.9, 0.8, and 0.7. For example, for correct observation probability of 0.9, 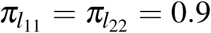 and 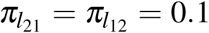. For the perfect classification rate, we fit the regular catSPIM model. For the imperfect observation probabilities, we considered *n*^*reps*^ ∈ (3, 5, 7). The two covariate value frequencies for all identity covariates were set to 0.5, i.e., *γ*_*l*_ = (0.5, 0.5). These specifications led to 4 perfect observation scenarios and 48 imperfect observation scenarios.

We simulated 96 data sets for each of the 52 scenarios. Then, we ran 3 MCMC chains for 100,000 - 150,000 iterations, depending on the MCMC mixing characteristics of each scenario, with chains thinned by 50 iterations, and discarded 12,500 iterations as burn in. We calculated the Gelman-Rubin statistic, *R*_*c*_ (Gelman et al. 1992), using *R*_*c*_ < 1.1 for *N* to indicate convergence. We discarded simulated data sets that did not meet this criterion. Within each set of scenarios with the same correct observation probability, if convergence was indicated for over 99% of the data sets for a given level of *n*^*rep*^, we reverted to running a single chain for the next level of *n*^*rep*^ (i.e., if *R*_*c*_ < 1.1 for *>*99% of simulations with *n*^*reps*^=3, we assumed all chains would converge for *n*^*reps*^ ∈ (5, 7) and used only 1 chain). We used posterior modes for point estimates and 95% highest posterior density (HPD) intervals for interval estimates. Finally, when estimating the elements of ***π*** and ***γ***, we pooled parameters across identity covariates *l*, so we discard the *l* indicator for ***π*** and ***γ*** in the results below.

#### Simulation Results

The abundance point estimates were approximately unbiased and the 95% credible intervals had approximately nominal coverage across all scenarios (Table E1). The precision of the abundance estimates was higher when the correct observation probability was higher, when there were more identity covariates, and when there were more replicated assignments (Table E1, Figure E1). The increased precision from including additional identity covariates and replicated assignments was more pronounced when the correct observation probability was lower (Table E1, Figure E1). Point estimates of the correct observation probabilities, *π*_11_ and *π*_22_, were approximately unbiased and interval estimates had approximately nominal coverage except in some scenarios with the lowest correct observation probability (0.7) and with fewer replicated assignments (Table E2). The point estimates for the category level probabilities were approximately unbiased in all scenarios and coverage was close to nominal in most scenarios except those with the lowest correct observation probabilities (0.7, Table E2).

**Figure E1:**
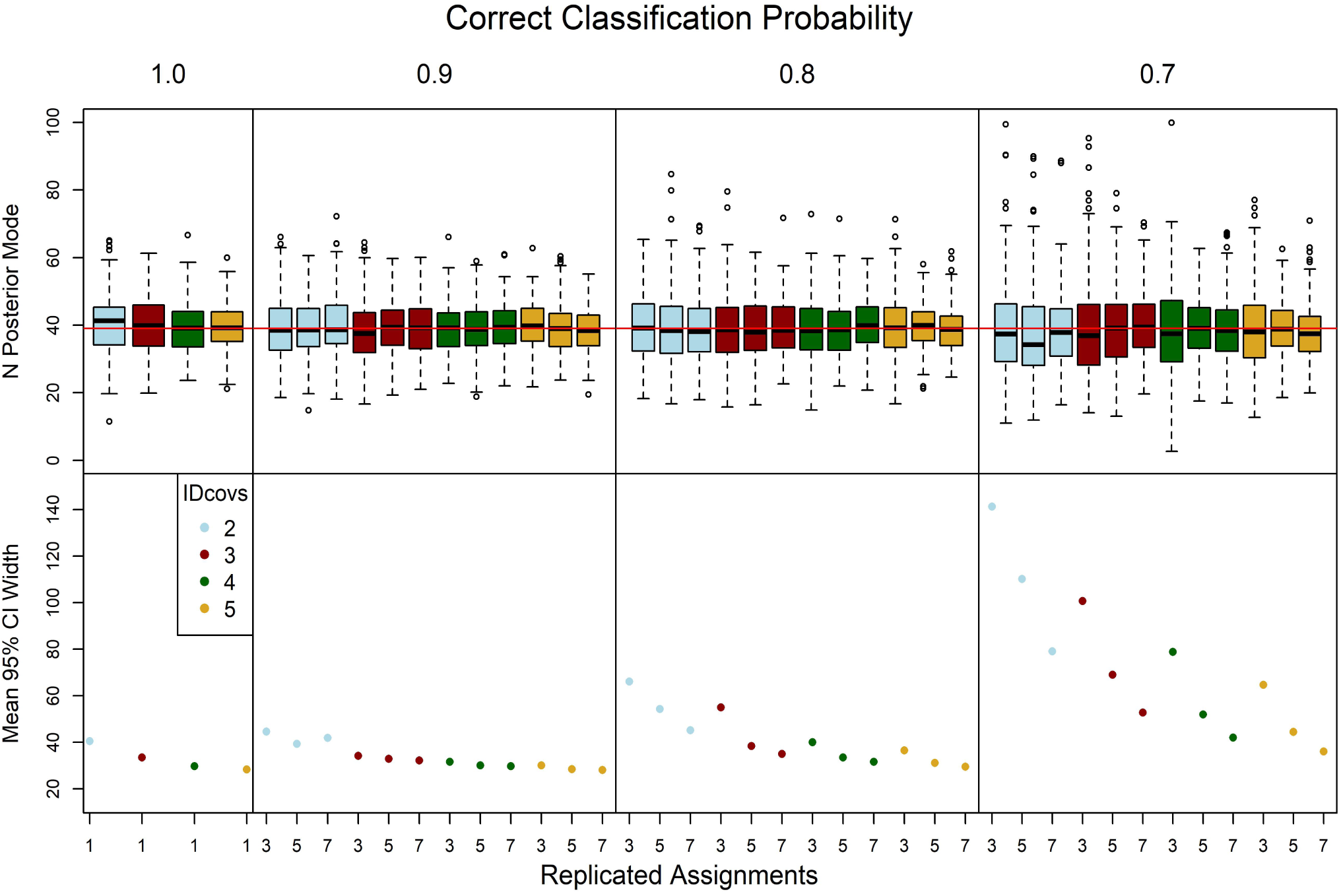
Simulation results of abundance estimate accuracy and precision as a function of the correct observation probability, number of identity covariates, and number of replicated assignments. The top row displays box plots of the abundance point estimates and the bottom row displays the mean 95% credible interval width. The red line in the point estimate box plots indicates the true value of abundance, 39.

**Table B1:** Simulation results for catSPIM-OE. *π* indicates the correct observation probability, *n*^*cat*^ indicates the number of identity covariates, and *n*^*reps*^ indicates the number of replicated assignments. 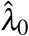 is the mean baseline detection point estimate, 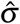 is the mean detection function spatial scale estimate, 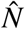 is the mean abundance estimate, 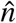 is the mean estimate of the number of individuals detected, N Cov is the coverage of the 95% credible interval, N wid is the mean width of the 95% credible interval, n is the mean number of individuals detected across simulations, and *R*_*c*_ is the proportion of simulated data sets for which the Gelman-Rubin convergence statistics was < 1.1.

| $\pi$ | $n^{cat}$ | $n^{reps}$ | $\hat{\lambda}_0$ | $\hat{\sigma}$ | $\hat{N}$ | $\hat{n}$ | N<br>Cov | N<br>Wid | n | $R_c$ |
| --- | --- | --- | --- | --- | --- | --- | --- | --- | --- | --- |
| 1.0 | 2 | 1 | 0.253 | 0.495 | 40.6 | 19.0 | 0.94 | 40.5 | 19.0 | 1.00 |
| 1.0 | 3 | 1 | 0.245 | 0.491 | 40.0 | 18.6 | 0.96 | 33.5 | 18.6 | 1.00 |
| 1.0 | 4 | 1 | 0.254 | 0.487 | 39.4 | 18.1 | 0.96 | 29.7 | 18.3 | 1.00 |
| 1.0 | 5 | 1 | 0.251 | 0.491 | 39.4 | 18.3 | 0.94 | 28.3 | 18.6 | 1.00 |
| 0.9 | 2 | 3 | 0.246 | 0.488 | 39.2 | 18.3 | 0.97 | 44.5 | 18.5 | 0.99 |
| 0.9 | 3 | 3 | 0.248 | 0.491 | 38.1 | 17.7 | 0.96 | 34.2 | 18.2 | 1.00 |
| 0.9 | 4 | 3 | 0.252 | 0.486 | 39.0 | 18.2 | 0.97 | 31.6 | 18.4 | 1.00 |
| 0.9 | 5 | 3 | 0.253 | 0.487 | 39.6 | 18.5 | 0.97 | 30.0 | 18.6 | 1.00 |
| 0.9 | 2 | 5 | 0.247 | 0.492 | 39.3 | 18.4 | 0.98 | 39.3 | 18.6 | . |
| 0.9 | 3 | 5 | 0.250 | 0.490 | 39.3 | 18.4 | 0.94 | 32.8 | 18.6 | . |
| 0.9 | 4 | 5 | 0.255 | 0.486 | 39.1 | 18.2 | 0.94 | 30.0 | 18.4 | . |
| 0.9 | 5 | 5 | 0.251 | 0.489 | 39.1 | 18.2 | 0.92 | 28.5 | 18.4 | . |
| 0.9 | 2 | 7 | 0.245 | 0.484 | 40.2 | 18.6 | 0.97 | 41.9 | 18.5 | . |
| 0.9 | 3 | 7 | 0.261 | 0.480 | 39.4 | 18.3 | 0.97 | 32.2 | 18.4 | . |
| 0.9 | 4 | 7 | 0.250 | 0.489 | 39.7 | 18.3 | 0.96 | 29.7 | 18.5 | . |
| 0.9 | 5 | 7 | 0.242 | 0.499 | 38.7 | 18.1 | 0.95 | 28.0 | 18.4 | . |
| 0.8 | 2 | 3 | 0.241 | 0.484 | 39.4 | 18.6 | 0.99 | 66.0 | 18.7 | 0.88 |
| 0.8 | 3 | 3 | 0.249 | 0.484 | 39.2 | 18.3 | 0.97 | 55.0 | 18.3 | 0.94 |
| 0.8 | 4 | 3 | 0.257 | 0.479 | 39.0 | 18.2 | 0.97 | 40.0 | 18.3 | 0.99 |
| 0.8 | 5 | 3 | 0.244 | 0.486 | 39.2 | 18.3 | 0.95 | 36.5 | 18.5 | 0.99 |
| 0.8 | 2 | 5 | 0.252 | 0.488 | 39.4 | 18.4 | 0.98 | 54.2 | 18.7 | 0.95 |
| 0.8 | 3 | 5 | 0.252 | 0.482 | 38.5 | 17.9 | 0.94 | 38.4 | 18.0 | 1.00 |
| 0.8 | 4 | 5 | 0.253 | 0.486 | 39.0 | 18.1 | 0.95 | 33.5 | 18.3 | 0.99 |
| 0.8 | 5 | 5 | 0.252 | 0.488 | 39.7 | 18.6 | 0.97 | 31.1 | 18.7 | 1.00 |
| 0.8 | 2 | 7 | 0.254 | 0.492 | 38.6 | 18.2 | 0.97 | 45.1 | 18.4 | . |
| 0.8 | 3 | 7 | 0.252 | 0.487 | 39.2 | 18.4 | 0.99 | 34.9 | 18.4 | . |
| 0.8 | 4 | 7 | 0.247 | 0.496 | 39.9 | 18.7 | 0.96 | 31.6 | 18.9 | . |
| 0.8 | 5 | 7 | 0.249 | 0.494 | 38.7 | 18.1 | 0.98 | 29.4 | 18.1 | . |
| 0.7 | 2 | 3 | 0.265 | 0.465 | 39.2 | 19.0 | 0.96 | 141.2 | 18.4 | 0.78 |
| 0.7 | 3 | 3 | 0.255 | 0.473 | 39.6 | 18.8 | 0.94 | 100.7 | 18.6 | 0.85 |
| 0.7 | 4 | 3 | 0.238 | 0.509 | 39.1 | 18.5 | 0.96 | 78.8 | 18.5 | 0.86 |
| 0.7 | 5 | 3 | 0.249 | 0.481 | 39.6 | 18.6 | 0.98 | 64.6 | 18.4 | 0.88 |
| 0.7 | 2 | 5 | 0.250 | 0.482 | 38.1 | 18.1 | 0.96 | 110.2 | 18.1 | 0.76 |
| 0.7 | 3 | 5 | 0.241 | 0.488 | 39.7 | 18.6 | 0.97 | 69.0 | 18.4 | 0.90 |
| 0.7 | 4 | 5 | 0.253 | 0.478 | 39.3 | 18.5 | 0.97 | 51.9 | 18.3 | 0.95 |
| 0.7 | 5 | 5 | 0.239 | 0.489 | 39.0 | 18.3 | 0.98 | 44.5 | 18.3 | 0.97 |
| 0.7 | 2 | 7 | 0.248 | 0.486 | 39.0 | 18.6 | 0.95 | 79.0 | 18.4 | 0.92 |
| 0.7 | 3 | 7 | 0.242 | 0.484 | 40.2 | 18.9 | 0.99 | 52.8 | 18.8 | 0.97 |
| 0.7 | 4 | 7 | 0.249 | 0.488 | 39.0 | 18.4 | 0.95 | 42.0 | 18.4 | 0.99 |
| 0.7 | 5 | 7 | 0.254 | 0.485 | 38.1 | 17.8 | 0.94 | 36.0 | 18.3 | 1.00 |

**Table B2:**
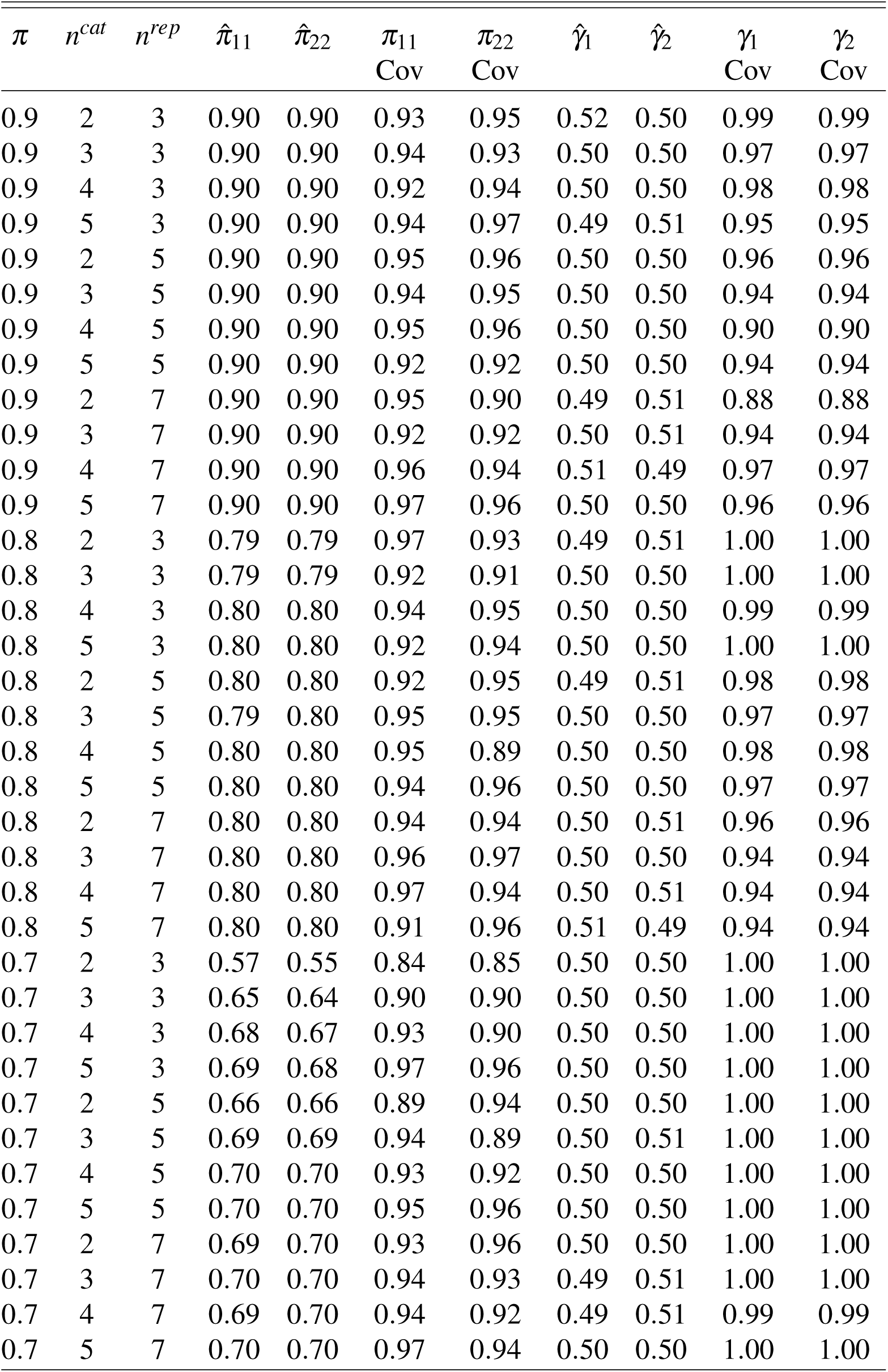
Continued simulation results for catSPIM-OE. This table contains point and interval estimate summaries for ***pi*** and *γ. π* indicates the correct observation probability, *n*^*cat*^ indicates the number of identity covariates, and *n*^*reps*^ indicates the number of replicated assignments. 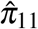 is the mean point estimate for the probability category level 1 is correctly classified, and 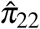 is the mean point estimate for the probability category level 2 is correctly classified. These are followed by the coverage of the 95% credible interval for each. 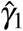 is the mean point estimate for the first category level probability and 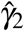 is the mean point estimate for the second category level probability. These are followed by the coverage of the 95% credible interval for each.

| $\pi$ | $n^{cat}$ | $n^{rep}$ | $\hat{\pi}_{11}$ | $\hat{\pi}_{22}$ | $\pi_{11}$<br>Cov | $\pi_{22}$<br>Cov | $\hat{\gamma}_1$ | $\hat{\gamma}_2$ | $\gamma_1$<br>Cov | $\gamma_2$<br>Cov |
| --- | --- | --- | --- | --- | --- | --- | --- | --- | --- | --- |
| 0.9 | 2 | 3 | 0.90 | 0.90 | 0.93 | 0.95 | 0.52 | 0.50 | 0.99 | 0.99 |
| 0.9 | 3 | 3 | 0.90 | 0.90 | 0.94 | 0.93 | 0.50 | 0.50 | 0.97 | 0.97 |
| 0.9 | 4 | 3 | 0.90 | 0.90 | 0.92 | 0.94 | 0.50 | 0.50 | 0.98 | 0.98 |
| 0.9 | 5 | 3 | 0.90 | 0.90 | 0.94 | 0.97 | 0.49 | 0.51 | 0.95 | 0.95 |
| 0.9 | 2 | 5 | 0.90 | 0.90 | 0.95 | 0.96 | 0.50 | 0.50 | 0.96 | 0.96 |
| 0.9 | 3 | 5 | 0.90 | 0.90 | 0.94 | 0.95 | 0.50 | 0.50 | 0.94 | 0.94 |
| 0.9 | 4 | 5 | 0.90 | 0.90 | 0.95 | 0.96 | 0.50 | 0.50 | 0.90 | 0.90 |
| 0.9 | 5 | 5 | 0.90 | 0.90 | 0.92 | 0.92 | 0.50 | 0.50 | 0.94 | 0.94 |
| 0.9 | 2 | 7 | 0.90 | 0.90 | 0.95 | 0.90 | 0.49 | 0.51 | 0.88 | 0.88 |
| 0.9 | 3 | 7 | 0.90 | 0.90 | 0.92 | 0.92 | 0.50 | 0.51 | 0.94 | 0.94 |
| 0.9 | 4 | 7 | 0.90 | 0.90 | 0.96 | 0.94 | 0.51 | 0.49 | 0.97 | 0.97 |
| 0.9 | 5 | 7 | 0.90 | 0.90 | 0.97 | 0.96 | 0.50 | 0.50 | 0.96 | 0.96 |
| 0.8 | 2 | 3 | 0.79 | 0.79 | 0.97 | 0.93 | 0.49 | 0.51 | 1.00 | 1.00 |
| 0.8 | 3 | 3 | 0.79 | 0.79 | 0.92 | 0.91 | 0.50 | 0.50 | 1.00 | 1.00 |
| 0.8 | 4 | 3 | 0.80 | 0.80 | 0.94 | 0.95 | 0.50 | 0.50 | 0.99 | 0.99 |
| 0.8 | 5 | 3 | 0.80 | 0.80 | 0.92 | 0.94 | 0.50 | 0.50 | 1.00 | 1.00 |
| 0.8 | 2 | 5 | 0.80 | 0.80 | 0.92 | 0.95 | 0.49 | 0.51 | 0.98 | 0.98 |
| 0.8 | 3 | 5 | 0.79 | 0.80 | 0.95 | 0.95 | 0.50 | 0.50 | 0.97 | 0.97 |
| 0.8 | 4 | 5 | 0.80 | 0.80 | 0.95 | 0.89 | 0.50 | 0.50 | 0.98 | 0.98 |
| 0.8 | 5 | 5 | 0.80 | 0.80 | 0.94 | 0.96 | 0.50 | 0.50 | 0.97 | 0.97 |
| 0.8 | 2 | 7 | 0.80 | 0.80 | 0.94 | 0.94 | 0.50 | 0.51 | 0.96 | 0.96 |
| 0.8 | 3 | 7 | 0.80 | 0.80 | 0.96 | 0.97 | 0.50 | 0.50 | 0.94 | 0.94 |
| 0.8 | 4 | 7 | 0.80 | 0.80 | 0.97 | 0.94 | 0.50 | 0.51 | 0.94 | 0.94 |
| 0.8 | 5 | 7 | 0.80 | 0.80 | 0.91 | 0.96 | 0.51 | 0.49 | 0.94 | 0.94 |
| 0.7 | 2 | 3 | 0.57 | 0.55 | 0.84 | 0.85 | 0.50 | 0.50 | 1.00 | 1.00 |
| 0.7 | 3 | 3 | 0.65 | 0.64 | 0.90 | 0.90 | 0.50 | 0.50 | 1.00 | 1.00 |
| 0.7 | 4 | 3 | 0.68 | 0.67 | 0.93 | 0.90 | 0.50 | 0.50 | 1.00 | 1.00 |
| 0.7 | 5 | 3 | 0.69 | 0.68 | 0.97 | 0.96 | 0.50 | 0.50 | 1.00 | 1.00 |
| 0.7 | 2 | 5 | 0.66 | 0.66 | 0.89 | 0.94 | 0.50 | 0.50 | 1.00 | 1.00 |
| 0.7 | 3 | 5 | 0.69 | 0.69 | 0.94 | 0.89 | 0.50 | 0.51 | 1.00 | 1.00 |
| 0.7 | 4 | 5 | 0.70 | 0.70 | 0.93 | 0.92 | 0.50 | 0.50 | 1.00 | 1.00 |
| 0.7 | 5 | 5 | 0.70 | 0.70 | 0.95 | 0.96 | 0.50 | 0.50 | 1.00 | 1.00 |
| 0.7 | 2 | 7 | 0.69 | 0.70 | 0.93 | 0.96 | 0.50 | 0.50 | 1.00 | 1.00 |
| 0.7 | 3 | 7 | 0.70 | 0.70 | 0.94 | 0.93 | 0.49 | 0.51 | 1.00 | 1.00 |
| 0.7 | 4 | 7 | 0.69 | 0.70 | 0.94 | 0.92 | 0.49 | 0.51 | 0.99 | 0.99 |
| 0.7 | 5 | 7 | 0.70 | 0.70 | 0.97 | 0.94 | 0.50 | 0.50 | 1.00 | 1.00 |

### Andean Bear Application

#### Data Description

Here, we describe an example of how the categorical Spatial Partial Identity Model with observation error (catSPIM-OE) can be applied to camera trap data using an Andean bear camera trap data set from Ecuador. This specific application did not work well, but we describe the general methodology and results to inform potential future applications to other camera trap data sets which may work better. The full details of this data set can be found in Springer (2018)–we will only describe the details relevant for this analysis here. The study was conducted in the Ecuadorian Andes across an area of 805 km^2^. The study area was divided into 1 km^2^ grid cells, with camera stations placed in 70 grid cells. Two cameras were deployed at camera stations in 31 grid cells, and one camera was deployed at camera stations in 39 grid cells. All camera stations were baited with a vanilla scent lure to increase the time individuals spent in front of cameras. These 70 camera stations were operated for 106 days. Photographs collected within a 5 minute period were assumed to be of the same individual and classified as a capture event. Capture events within 30 minutes of another capture event were discarded to reduce the probability that events from the same bear were counted multiple times, violating basic capture-recapture independence assumptions. These guidelines produced 139 capture events from an unknown number of Andean bears.

#### Image Processing

Categorical identity covariates were produced from each capture event using volunteers to digitize the visible facial and neck markings across all photographs within a capture event using Adobe Photoshop. Volunteers were provided with a bear face template (Figure E1b) and asked to fill in the areas of the face and neck which contained marks with white and areas not visible with gray (e.g., Figure E1c). Areas not colored by the volunteers were black, indicating no marks were present in these areas. A total of 9 volunteers digitized the bear markings, with 3 observers classifying each capture event. After all facial and neck markings were digitized by multiple observers in continuous space, categorical identity covariates were created by discretizing the face into grid cells. The brow region was identified as the area of the face that varied the most across individuals and where there was the most agreement across observers (Figure E2). The muzzle region was almost always classified as marked because almost all bears had markings in this area and agreement between observers was high because these markings were the easiest to digitize accurately. Conversely, almost all bears had markings in the neck region, but the observers varied considerably in whether or not they actually digitized these areas and agreement was lower because this region was difficult to digitize accurately. Thus, we focused on the brow region. We used 3, 5, 10, and 20 grid cells to represent the brow markings, but only present results using 10 grid cells. If a grid cell intersected an area colored white, it was classified as “2” (marked). If a grid cell intersected an area colored black, but no areas colored white, it was classified as “1” (unmarked). If a grid cell intersected an area colored gray and no areas colored white, it was classified as “0”, indicating a missing observation.

#### CatSPIM-OE Model

The image processing described above produced a data set of 139 capture locations stored in ***y***^*obs*^ and the observed categorical identity covariates stored in ***G***^*obs*^ of dimension 139 × 10 × 3 (10 categorical identity covariates and 3 observer classifications). We assumed that each categorical identity covariate had its own probability of containing a mark across individuals (*γ*_*l*_ estimated separately for all *l*); however, we fixed the category level observation probabilities, ***π*** across identity covariates due to data sparsity. Because there are only two possible identity covariate values (marked or not marked), there were 4 observation probabilities to be estimated,

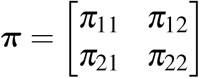

These probabilities correspond to correctly classifying an unmarked area as unmarked (*π*_11_), correctly classifying a marked area as marked (*π*_22_), incorrectly classifying an an unmarked area as marked (*π*_21_) and incorrectly classifying a marked area as unmarked (*π*_12_).

We fit the null model described in the main text that assumes the detection function parameters do not vary across individuals, but also tried models with individual heterogeneity in detection function parameters, and/or a behavioral response to capture. We also considered uninformative (Uniform(0,∞)) and various informative priors for *σ*. None of these models yielded plausible results, so we just describe the null model specifications with an uninformative prior here. We ran 1 MCMC chain for 100,000 iterations, discarding the first 50,000 as burn in. We used posterior modes for point estimates and 95% HPD intervals for interval estimates.

### Results and Discussion

We estimated that we captured 38 individuals in the 139 capture events and estimated a density of 10.69 individuals/100km^2^ (Table E1) Further, we estimated *σ* at 0.63 km. We can compare these estimates to previous estimates from a subset of this study area using SCR where the photographed individuals were presumably easier to identify confidently (Molina et al. 2017). Molina et al. (2017) estimated a lower density at 7.45 bears/100km^2^ and a substantially larger *σ* at 2.8 km. The *σ* estimate from Molina et al. (2017) is more in line with *σ* estimates from other bear populations. We believe we substantially underestimated *σ* and thus overestimated density. If so, this result would be consistent with previous simulations of unmarked SCR showing an underestimation of *σ* and overestimation of density when there is a high level of home range overlap among individuals (Augustine et al. 2019). A second line of evidence that we overestimated density is that we were able to identify 11 individuals from their facial markings with a relatively high level of confidence and we subjectively estimate there were probably 15 - 25 observed bears in the 139 capture events, not the 38 we estimated. Therefore, we do not view these parameter estimates as plausible.

The population level category level frequencies and category level observation probabilities; however, were plausible. The population frequency with which each of the 10 facial regions contained a mark were estimated from 0.174 to 0.716 and roughly corresponded with the pattern seen in the composite bear face drawing in Figure E2. Generally, areas closer to the center of the brow and lower on the brow had a higher probability of containing a mark than the outer and upper brow regions. We estimated that if a face region had no mark, it was classified as having no mark with high probability (0.97, Table E2). However, we estimated that observers less consistently recorded face regions with marks as having marks (0.74).

It is difficult to pinpoint one specific reason this application did not work as expected. We speculate that perhaps the largest two problems were that there was not enough variation in facial markings between bears and observers could not record them reliably enough. Multiple bears had no facial markings at all and others may have the same values in all facial regions when discretizing the face into just a few number of regions. Further, the photos were often blurry due to the low level of light under the forest canopy. Finally, there is a negative relationship between the number of facial regions used and the agreement between observers. As the face is broken into smaller regions, the observers are more likely to disagree about which facial regions contain marks. Therefore, this system may not have a high enough signal to noise ratio to reliably extract information about individual identity.

A second factor that might have prevented this application from working well is a violation of the SCR observation model. Individual heterogeneity in detection function parameters has been identified as a problem for spatial partial identity models (SPIM) such as this one (Augustine et al. 2018, 2019) and in our experience, including a behavioral response to capture can be required to correctly estimate *σ* and the number of individuals captured in SPIMs. We attempted to fit these models, but this data set was much to sparse for reliable estimation of these more complex models. We also suspect that there was a high level of variability in the detection rate across sites, with some sites recording many photographs of multiple bears, while other nearby sites recorded no photographs. In effect, this is caused by missing site covariates, perhaps related to fine scale habitat quality, or variability in bear use along different trails. We expect regular SCR is relatively robust to missing site covariates, but SPIMs will be less robust as it will tend to break the captures of single individuals into two or more individuals.

A third factor that might have prevented this application from working well is a violation of the face region identity covariates and/or their observation probabilities. In particular, the face region covariates are likely not independent as we assumed. For example, if a bear has markings at the edges of the brow, they almost certainly have markings in the center of the brow. Thus, we may need to consider spatial correlation in the mark patterns. Finally, there is likely heterogeneity in the category level observation probabilities across the face region covariates, which we could not accommodate with such a sparse data set and small number of observers. Observers may be more likely to agree about markings in the center of the brown than at the edges.

Despite the poor performance of this model for this application, we expect that there may be other camera-based applications that may work well, specifically ones where there is more variability in markings across individuals and a higher signal to noise ratio. Ideally, researchers will have independent density estimates for the same study area to which the SPIM estimates can be compared to increase confidence that the SPIM estimates are reliable.

### Figures

**Figure E1:**
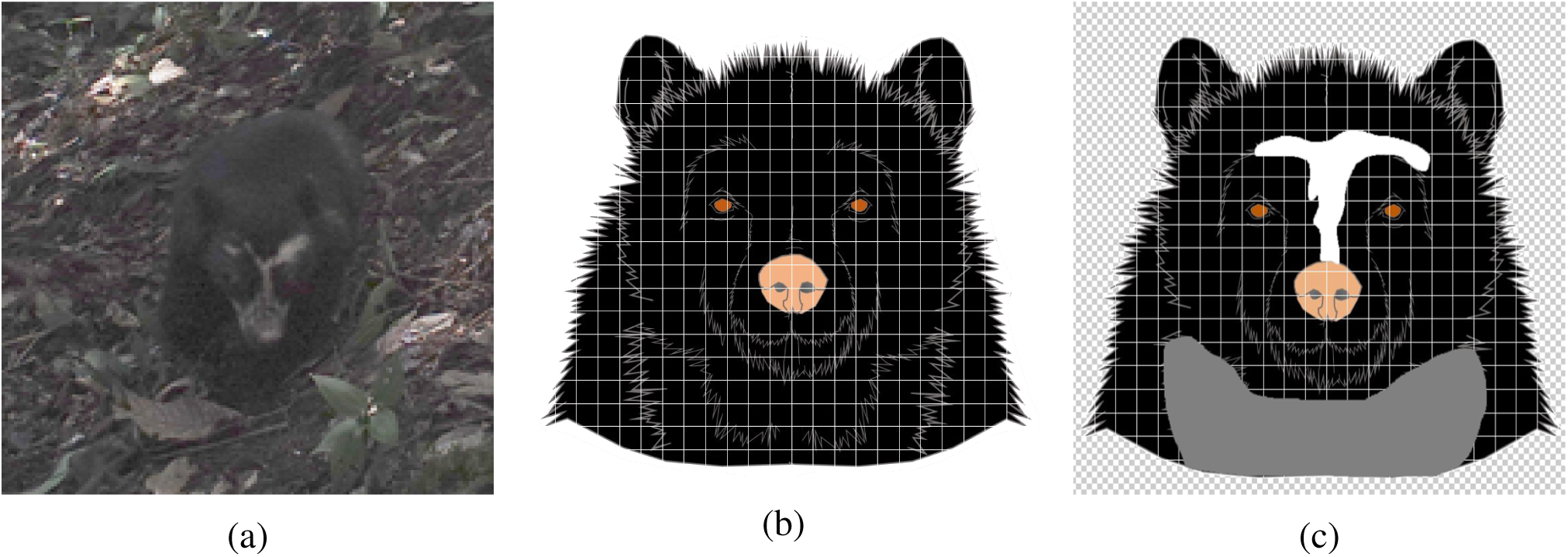
An example digitization of Andean bear face covariates. Figure B1a is a single photograph capture event, Figure B1b is the template used for digitizing marks, and Figure B1c is the digitized marks from one observer.

**Figure E2:**
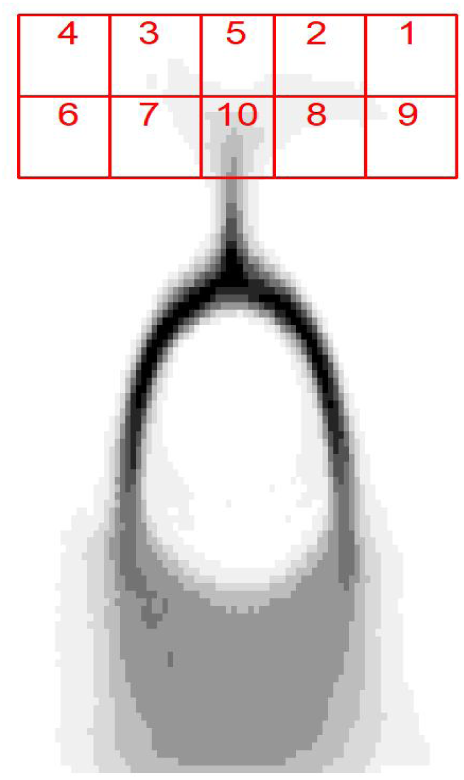
A composite of all bear face drawings across the 139 capture events with the 10 facial regions used as categorical identity covariates depicted in red. Darker areas indicate areas more consistently drawn as “marked” by observers across capture events. The numbers for the facial regions match those in Table E2 where the estimated probabilities of containing a mark are presented. All 10 facial regions were classified as “marked” in at least some drawings, despite the composite drawing not depicting marks in some regions due to the relative rarity of these events.

### Tables

**Table E3:** Detection function, abundance, and observation probability parameter estimates for the Andean bear analysis. *λ*_0_ is the baseline detection rate, *σ* is the detection function spatial scale parameter, *n* is the number of individuals captured, *N* is abundance, *D* is density, and *π*_*r,c*_ is the probability that identity covariates taking value *c* are recorded as *r*. Point estimates are posterior modes and “LB” and “UB” are the 95% HPD interval lower and upper bounds.

|  | Estimate | LB | UB |
| --- | --- | --- | --- |
| $\lambda_0$ | 0.07 | 0.05 | 0.10 |
| $\sigma$ | 0.63 | 0.53 | 0.77 |
| $N$ | 225 | 138 | 349 |
| $n$ | 38 | 33 | 41 |
| $D$ | 10.69 | 6.65 | 16.67 |
| $\pi_{11}$ | 0.97 | 0.96 | 0.98 |
| $\pi_{21}$ | 0.03 | 0.02 | 0.04 |
| $\pi_{12}$ | 0.26 | 0.23 | 0.29 |
| $\pi_{22}$ | 0.74 | 0.71 | 0.77 |

**Table E4:**
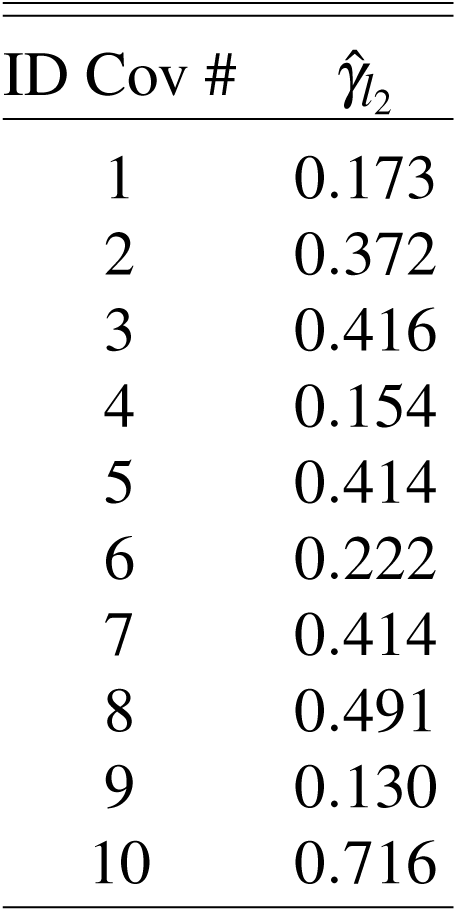
The estimated population frequencies with which each of the 10 facial regions contained a mark. ID Cov # is the facial region number depicted in Figure B2, and 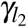 is the estimated frequency with which facial region *l* took value 2 (marked).

## Selected Detailed Sample Match Information

*Ben Augustine*

*December 21, 2019*

### Assignments that differed from the geneticist

Here, we will look at the five Genotype SPIM identity assignments that differed from those made by the geneticist. We will consider a posterior probability match of <0.99 as differing from the certain assignment made by the geneticist. Note, that the scoring rules used in the original study (Linden et al. 2017) were that a homozygote must be seen 3 times to be called and a heterozygote must be seen 2 times to be called.

#### Sample 3

This sample assignment only differed from that made by the geneticist because the geneticist used the sex information of the samples, which we did not use. However, this example is instructive of how the Genotype SPIM works.

Samples 2, 3, and 4 were assigned the same individual identity by the geneticist. The genotype SPIM assigns sample 3 to the same individual as the individual that produced samples 2 and 4 with probability 0.73. We will label this individual as individual A. The remainder of the match probability (0.27) is allocated to the event that sample 3 actually matches the same individual as the individual that produced sample 1, also collected at the same trap as samples 2-4. We will label this individual as individual B.

If sample 3 belongs to individual A, there was most likely 4 allelic dropouts for sample 3 at locus 2 (136.138 to 138.138). If sample 3 belongs to individual B, there was most likely 4 allelic dropouts for sample 3 at locus 5 (132.144 to 144.144). Given the most likely genotypes for individuals A and B, these 2 possible sets of allelic dropouts for sample 3 are equally likely under our genotyping error model. However, the assumed SCR observation model with compensation between *λ*_0_ and *σ* allocates different probabilities to these two identity assignments. Because there are two samples from individual A (2 and 4) and only one sample from individual B (1), and these samples were collected at a single trap, the 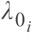 for individual B is lower (conversely, individual B has a larger *σ*_*i*_), and thus sample 3 matches individual A with higher probability than individual B under our model. This can be visualized in Plots 2 and 3 below showing the activity center posterior more strongly concentrated around the trap of capture for individual A than individual B due to the different capture numbers.

However, the geneticist knew that the focal sample and samples 2 and 4 came from females and sample 1 came from a male so the correct assignment was made. Interestingly, the Genotype SPIM gives a higher match probability for the correct assignment than the incorrect assignment. This is consistent with the sex of this individual being female. If detection is proportional to space use as our observation model assumes, the greater number of detections at a single trap for the correct match leads to this individual having a larger estimated 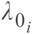 and lower estimated *σ*_*i*_ as we would expect for females which have smaller home range sizes than males.

The Genotype SPIM did not place any probability that all four of these samples came from the same individual. If so, there would have been 8 allelic dropouts for samples 1 and 3 at locus 2, and 12 allelic dropouts for samples 2, 3, and 4 at locus 5. This is a total of 20 allelic dropouts versus 4 if there were actually 2 individuals, making it much more likely there were indeed 2 individuals. The sex information of the samples that we did not use confirms this was the case, unless there were errors in sex determination.

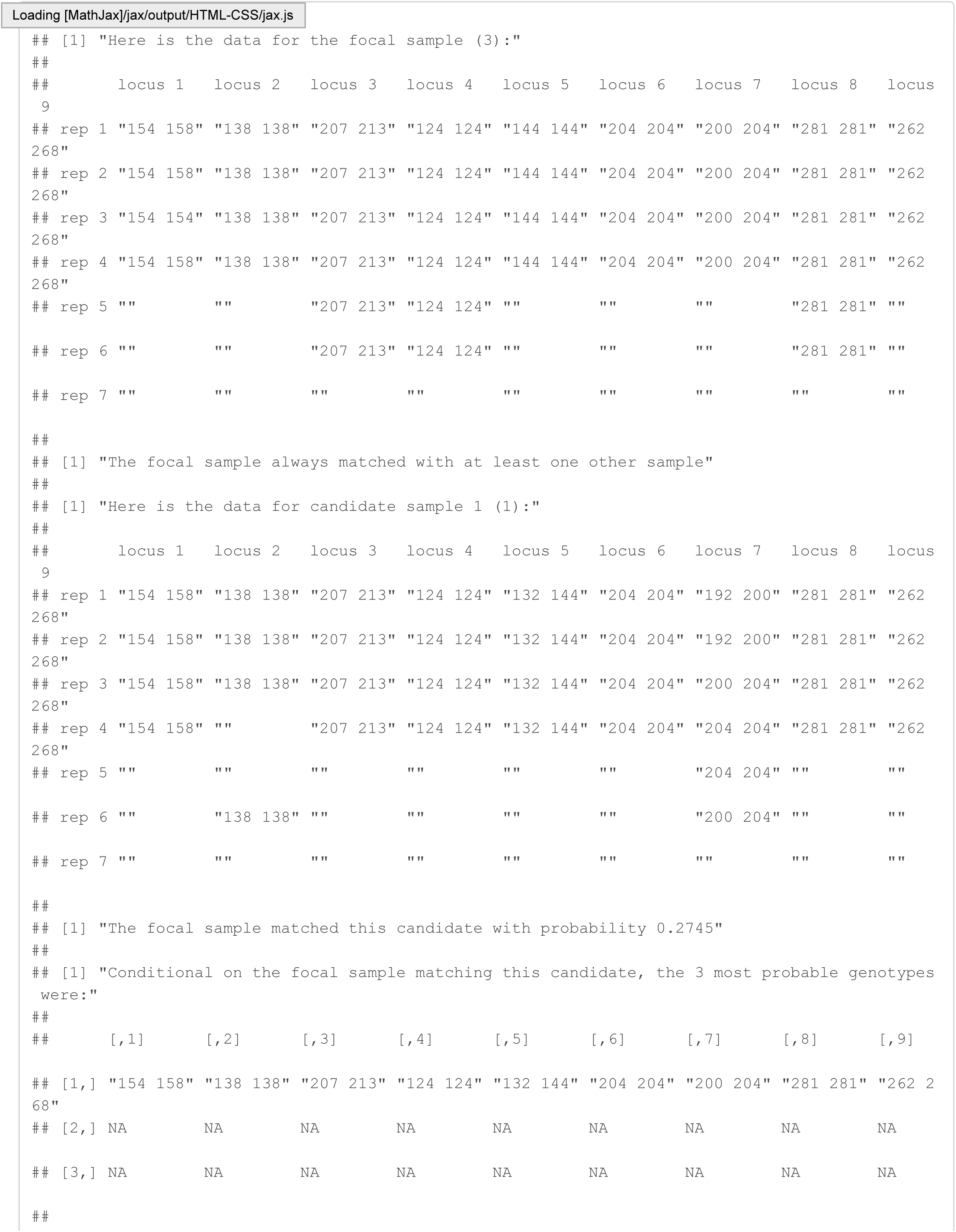

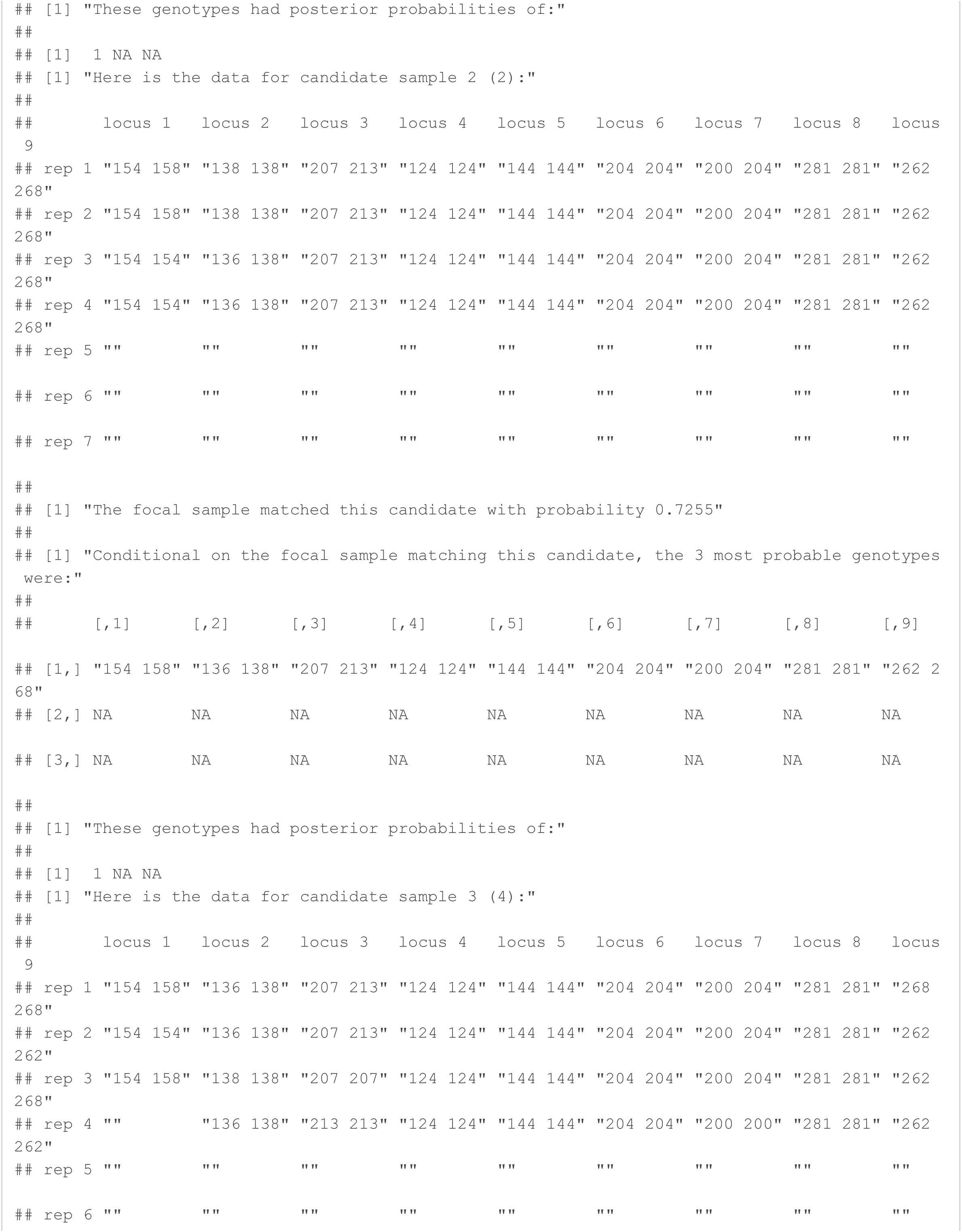

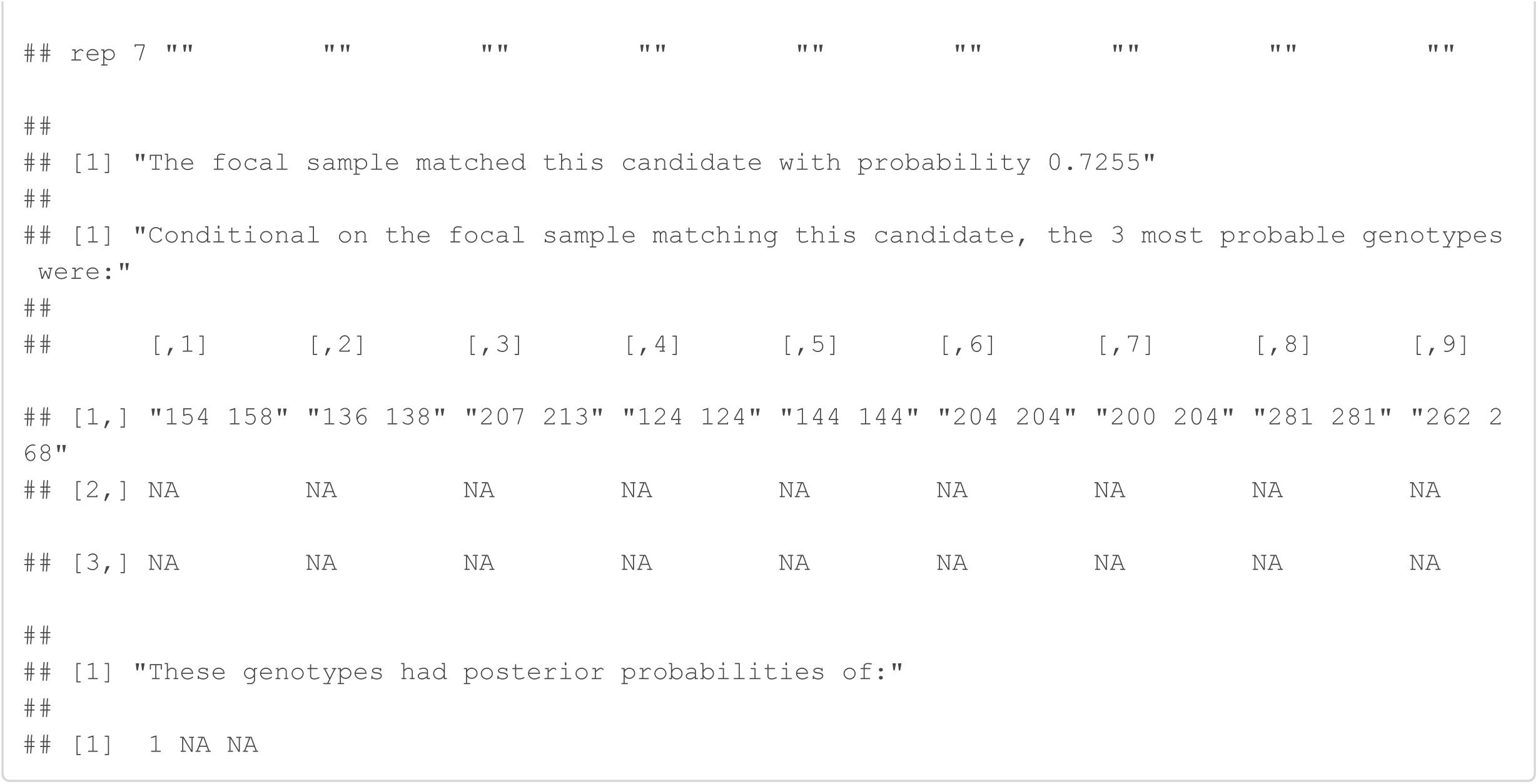

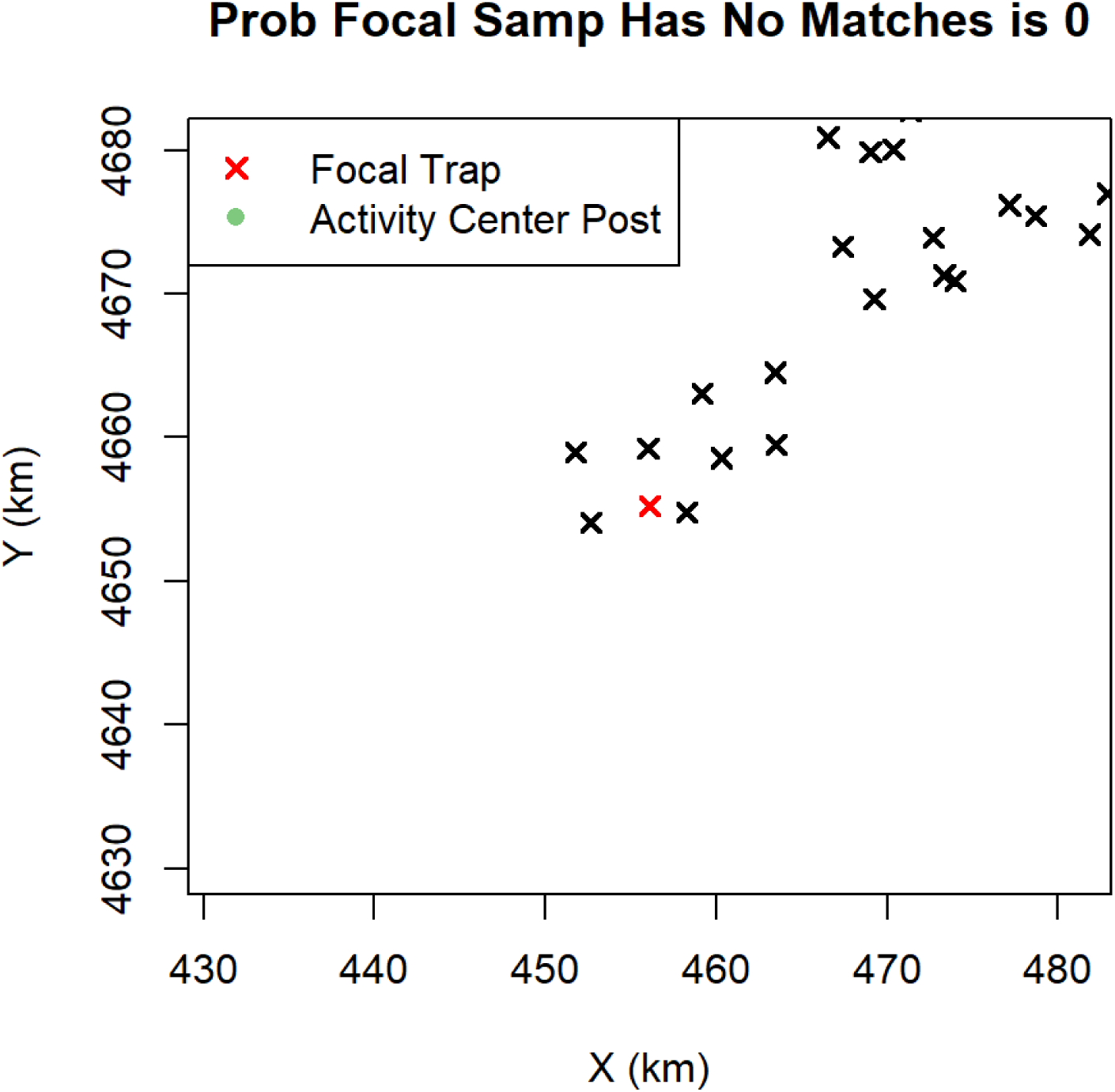

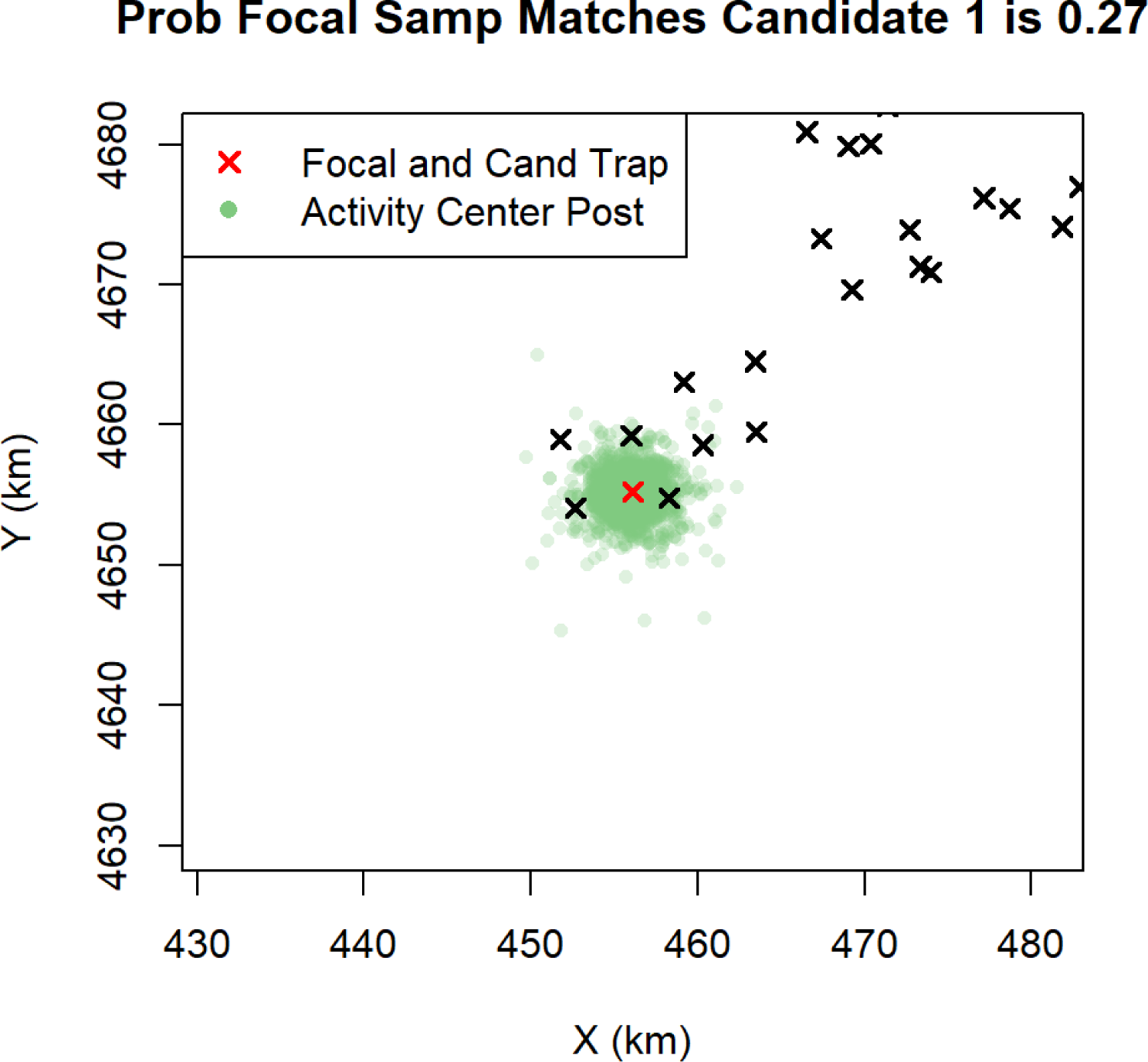

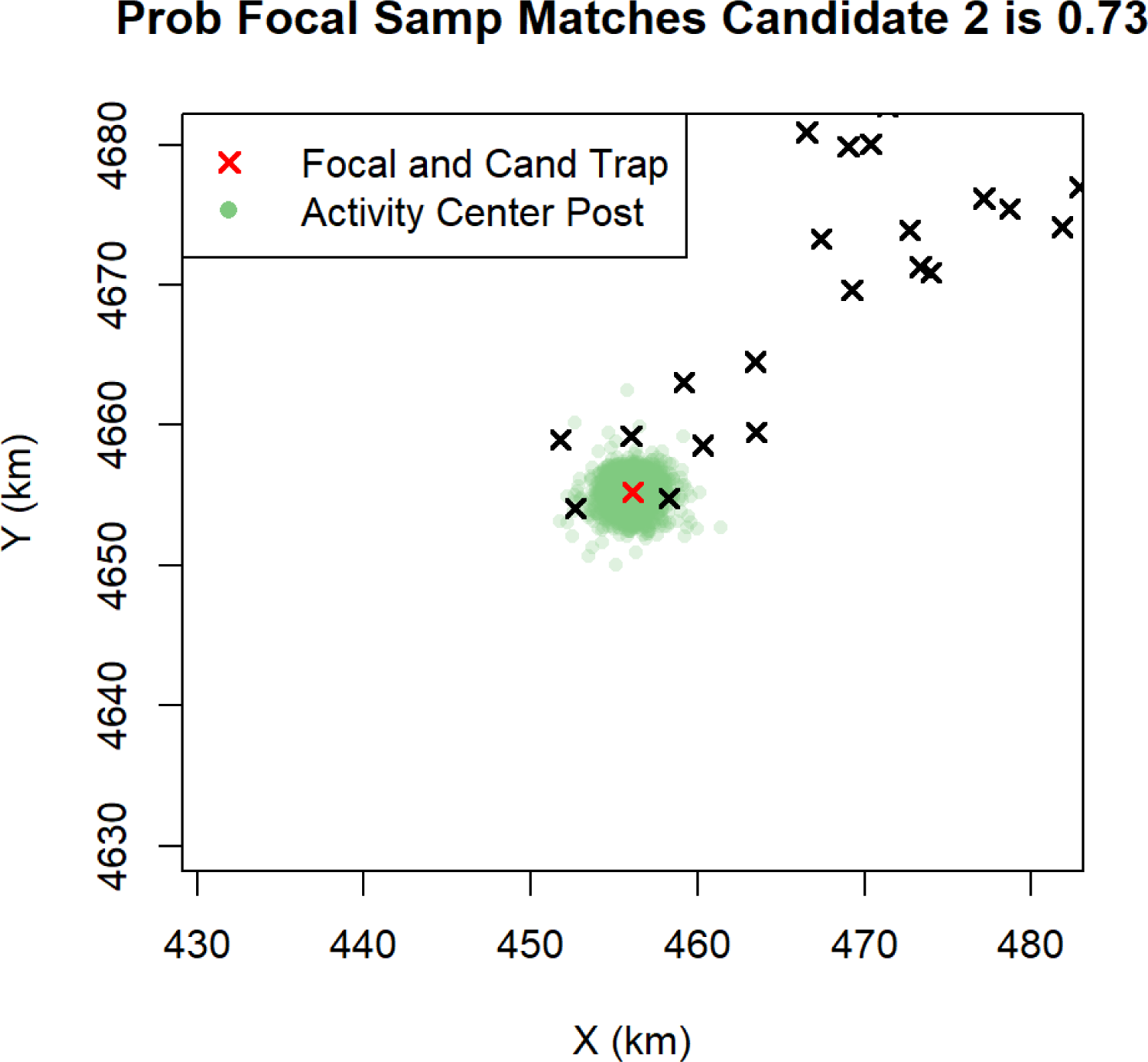

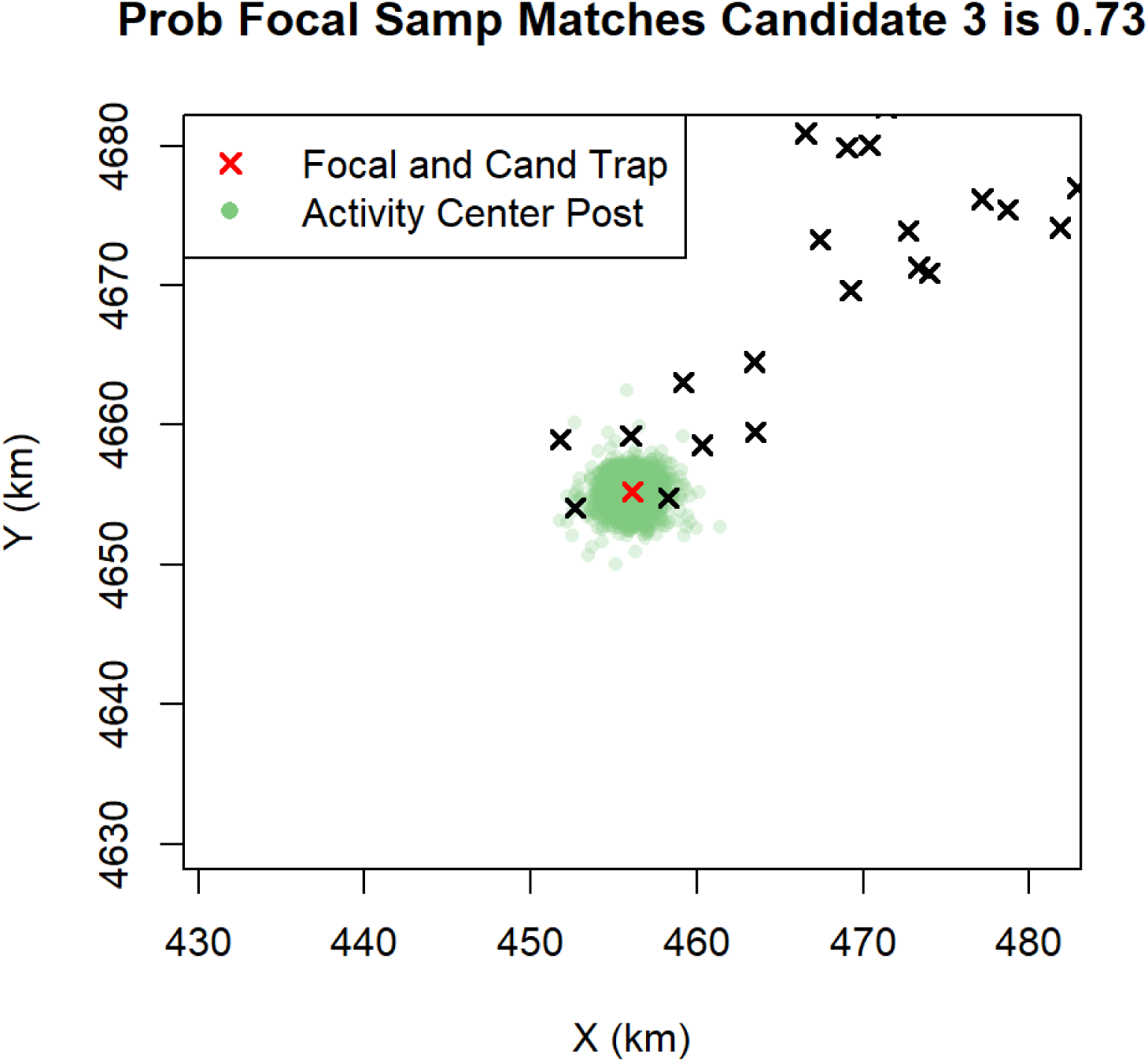

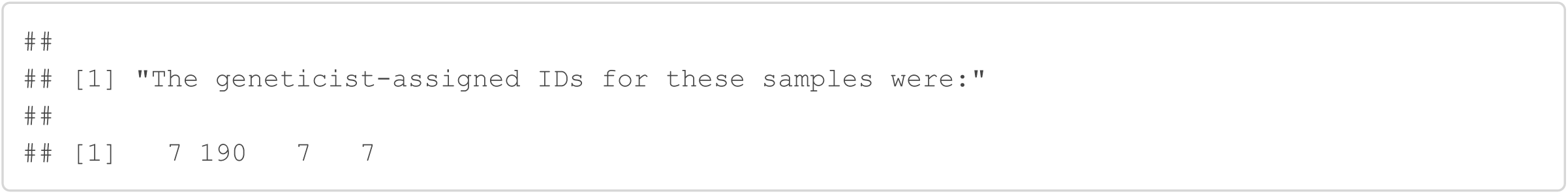

#### Sample 197

The geneticist grouped sample 197 with samples 195, 196, and 198 because the consensus genotypes only mismatched at 1 locus, locus 3. Sample 197 was scored as 213.215 at locus 3 across 4 replications, while the other 3 samples had a consensus score of 205.213 at locus 3. Thus, the geneticist’s grouping implies a false allele assignment occurred in sample 197 4 times; however, false alleles were estimated to be extremely rare in this data set. Therefore, the genotype SPIM assigned a probability of 1 that sample 197 was a unique individual in the data set. This example illustrates that the type of error (allelic dropout vs. false allele) that explains the mismatch between two samples and the number of times the error occurred, can be very informative about whether the two samples belong to the same individual or not. This information is typically not used.

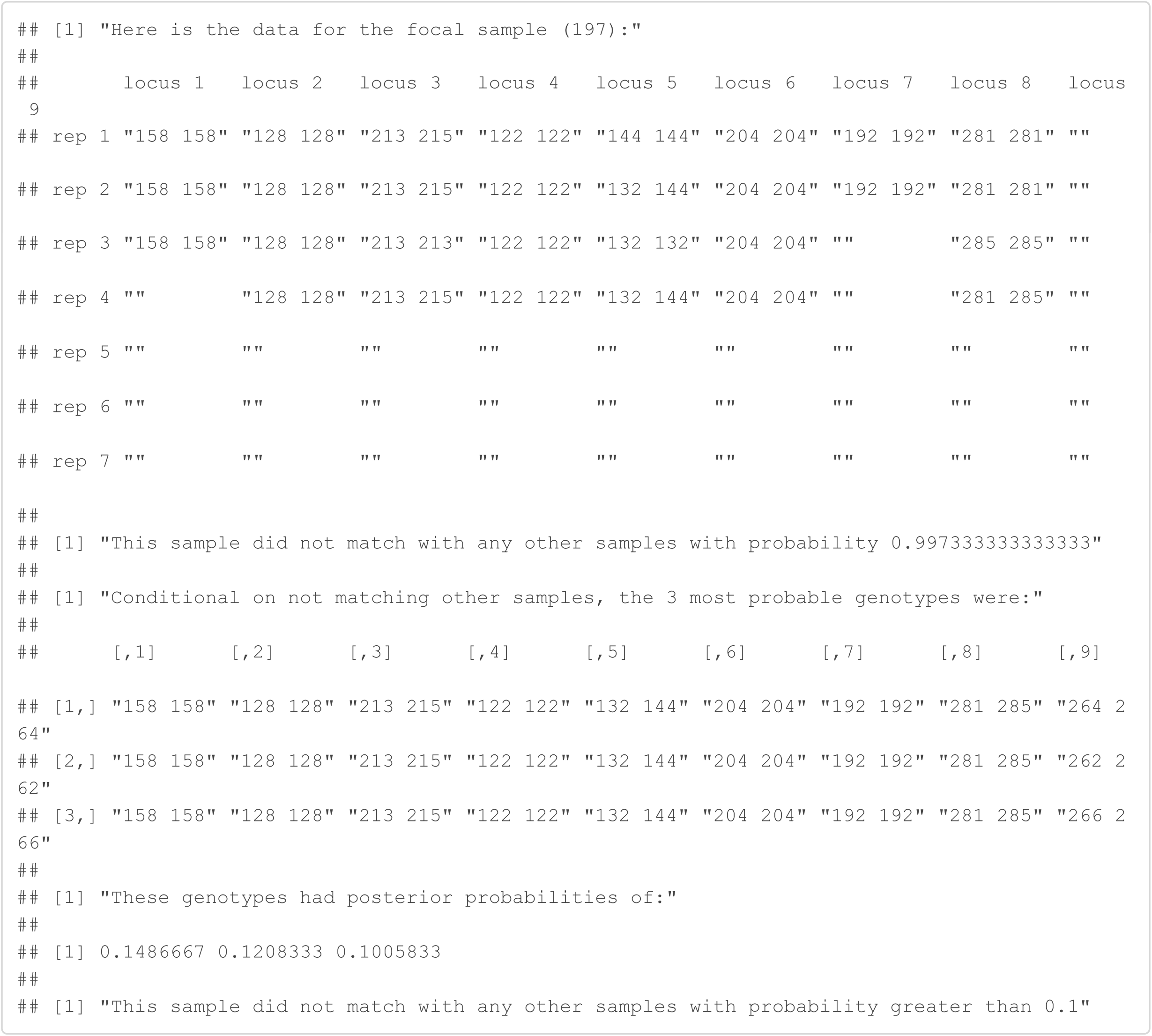

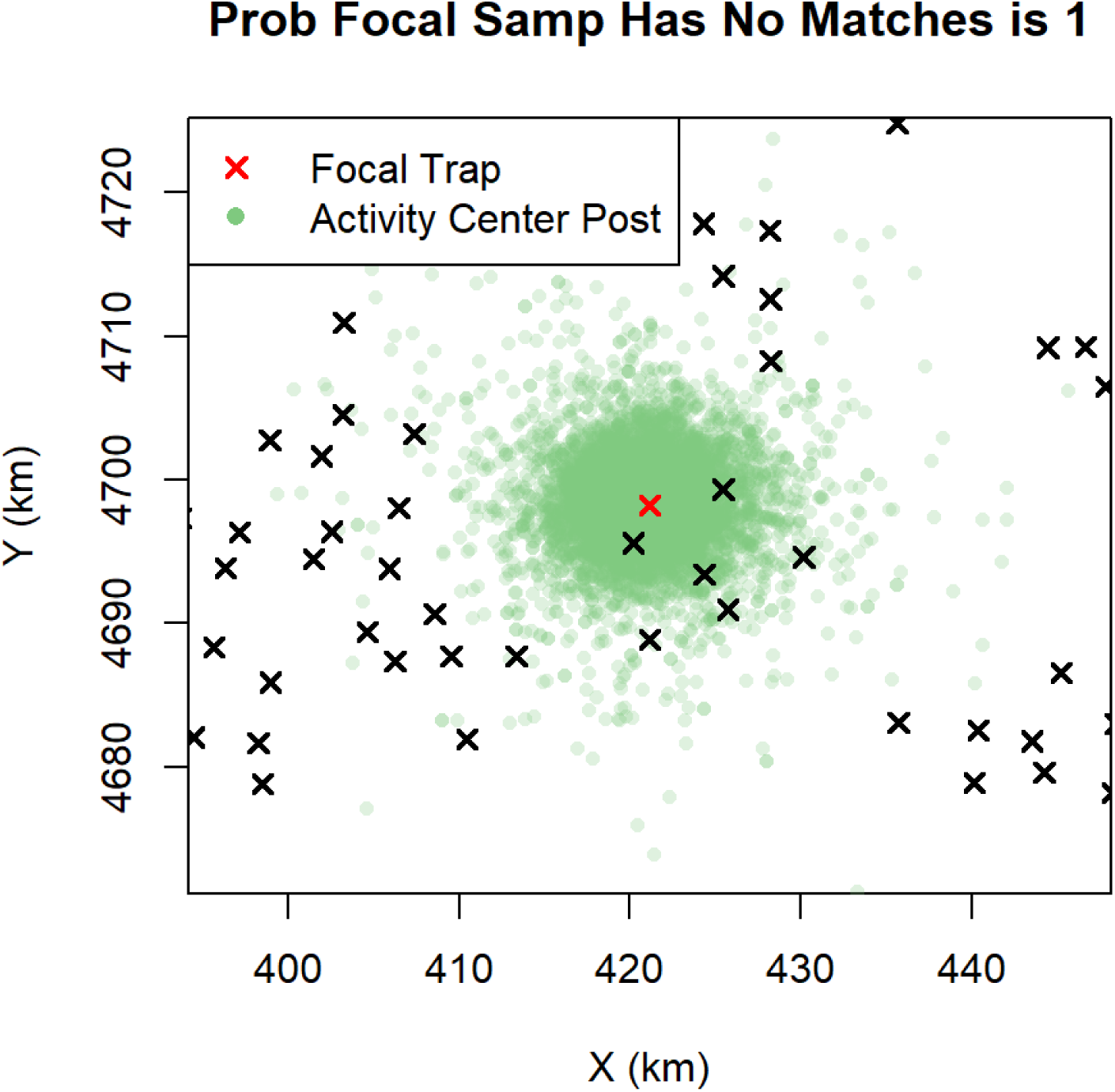

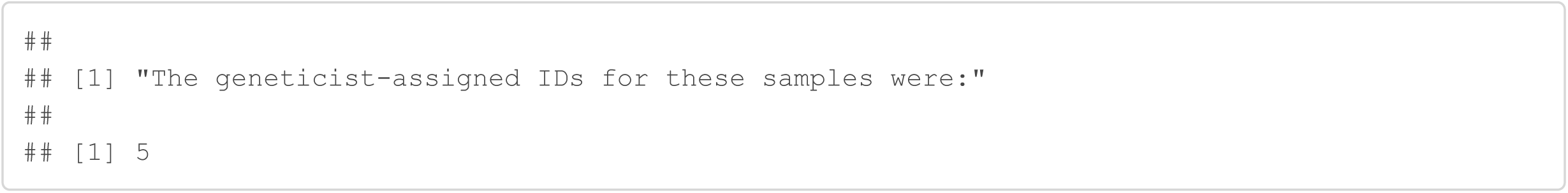

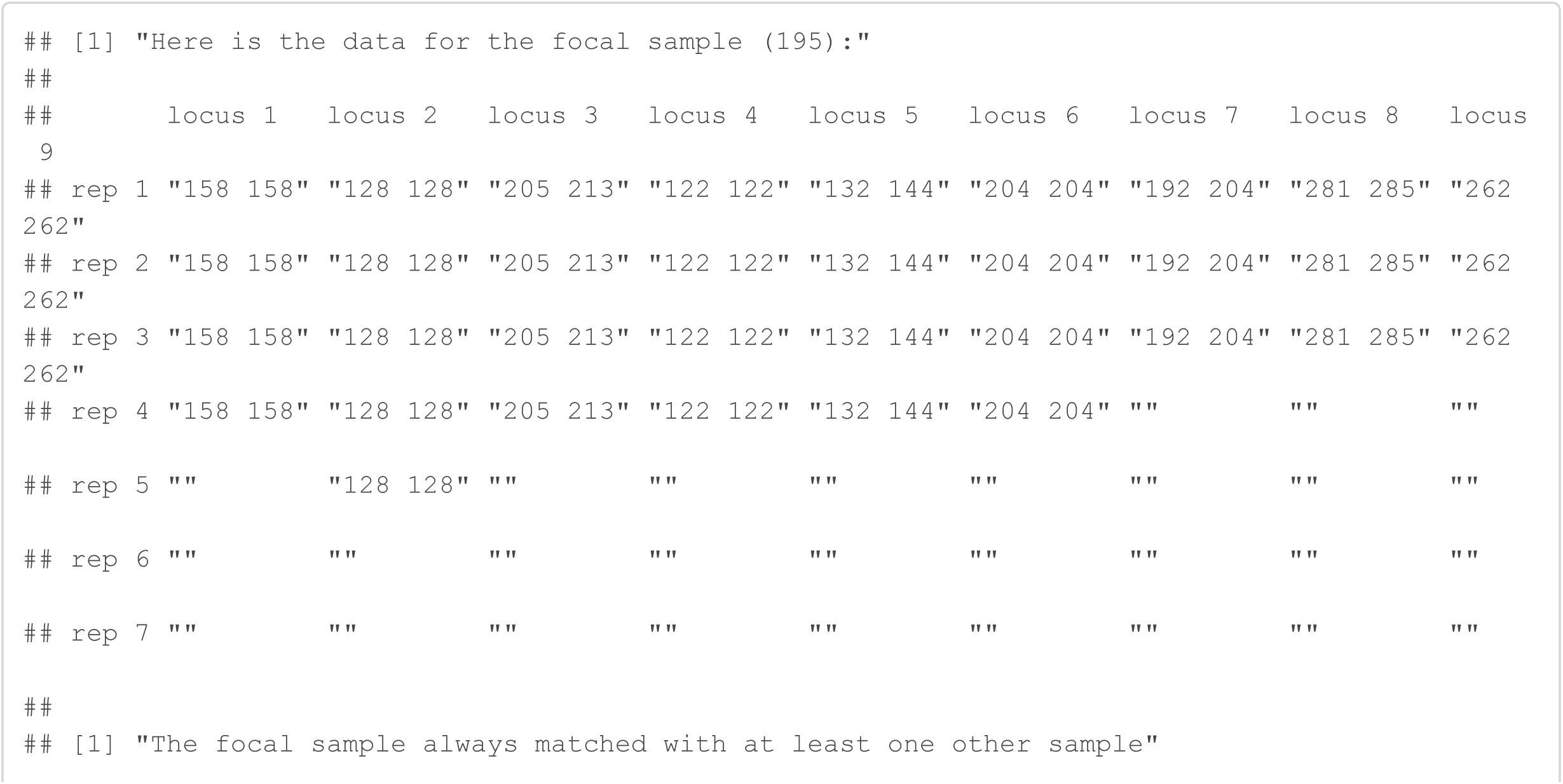

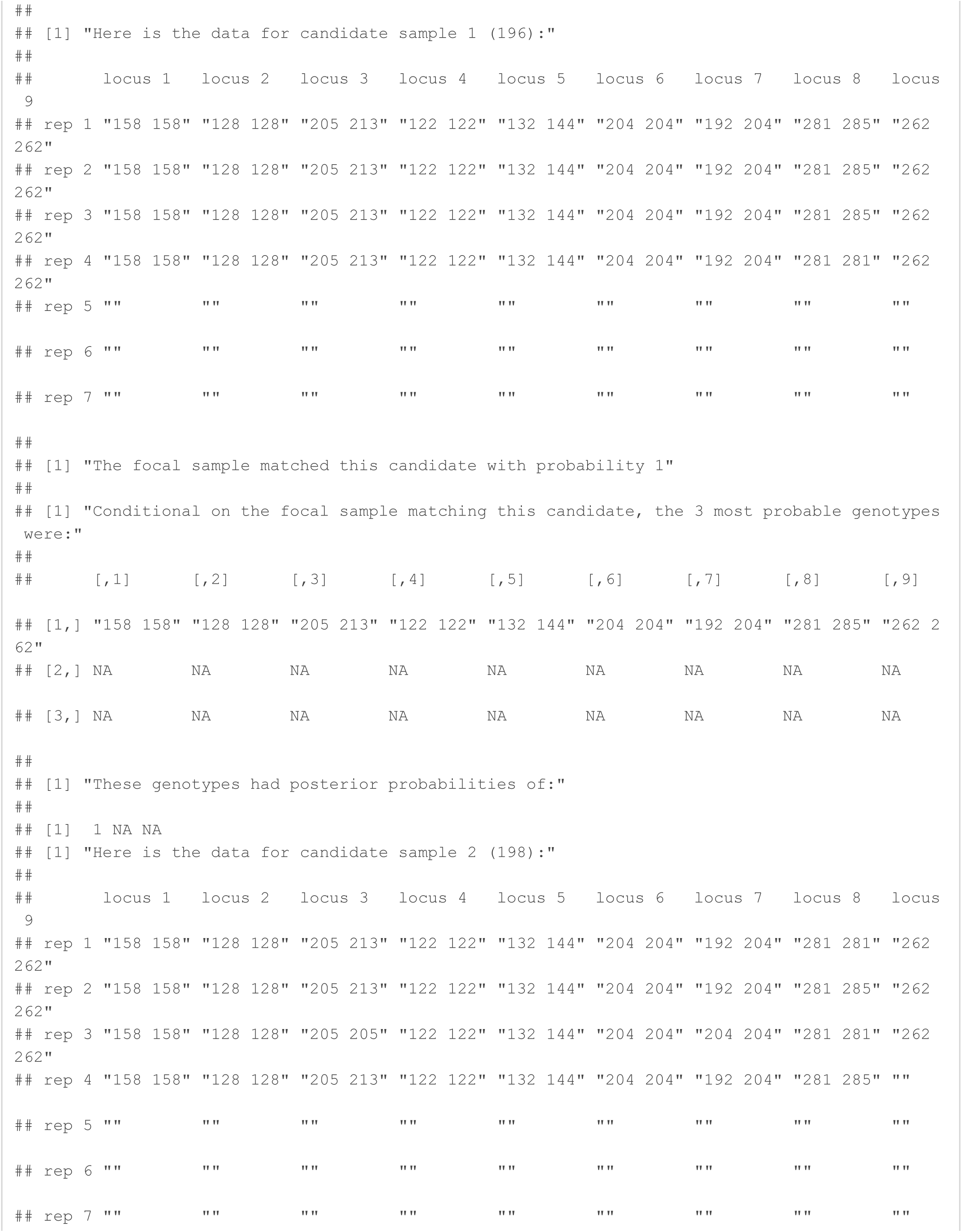

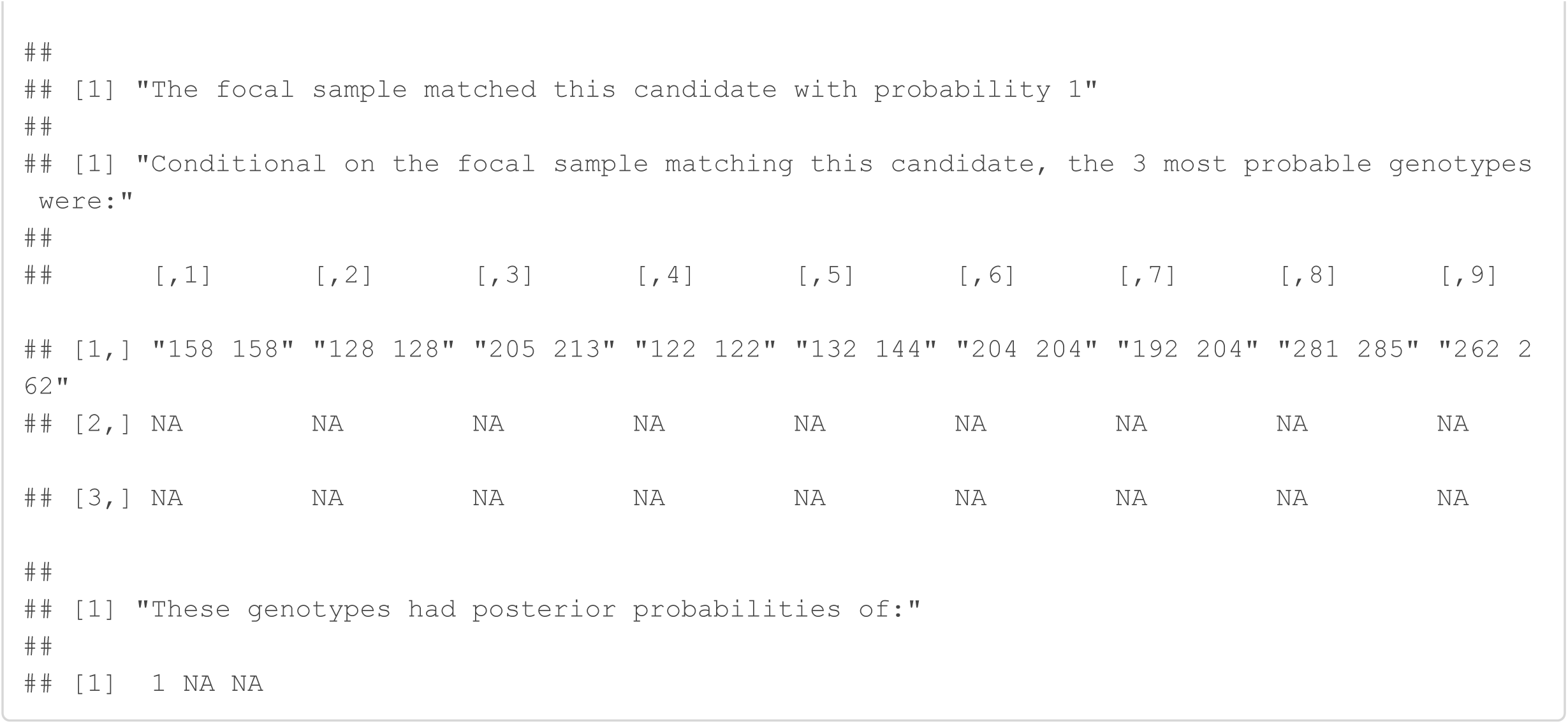

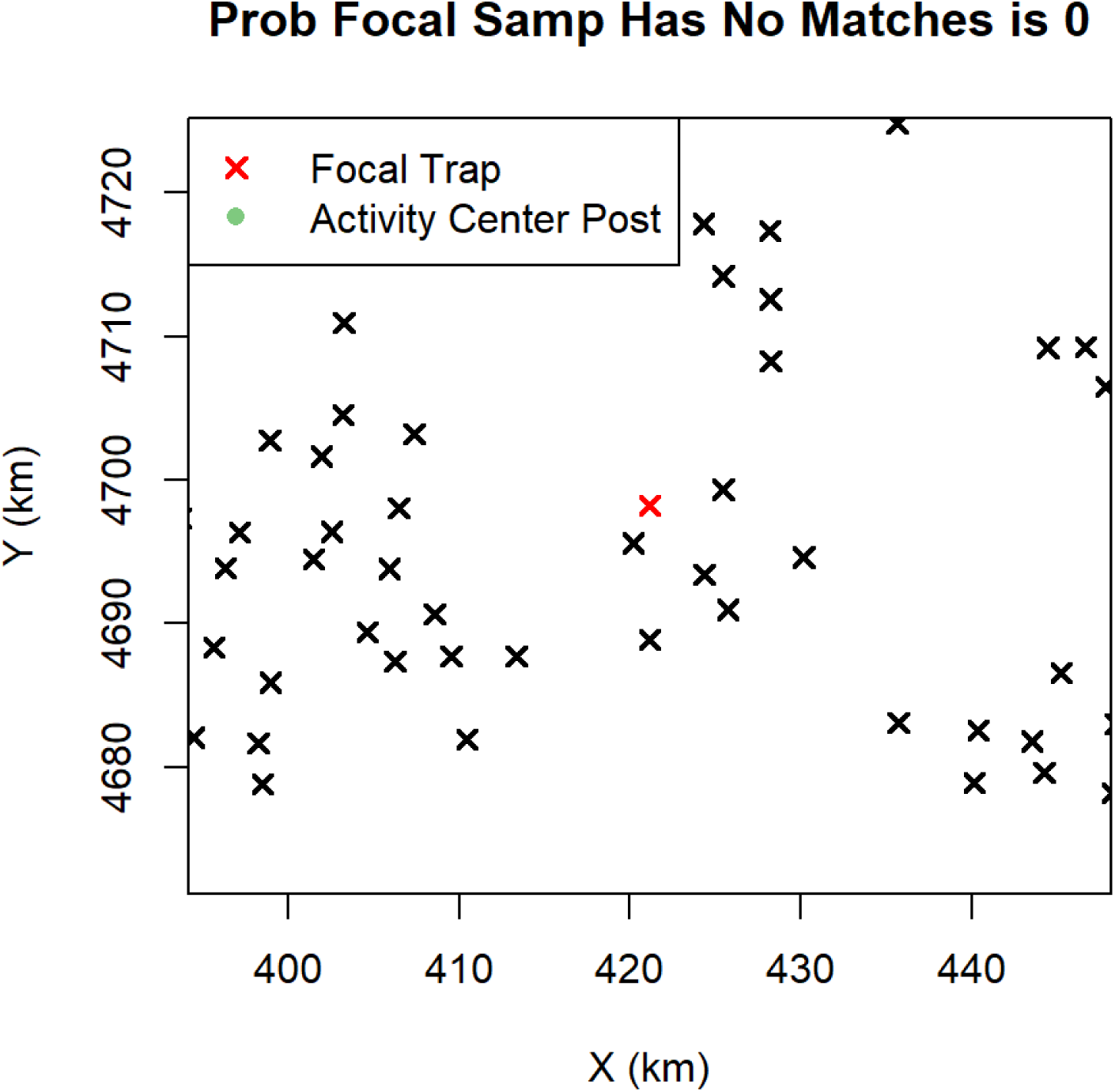

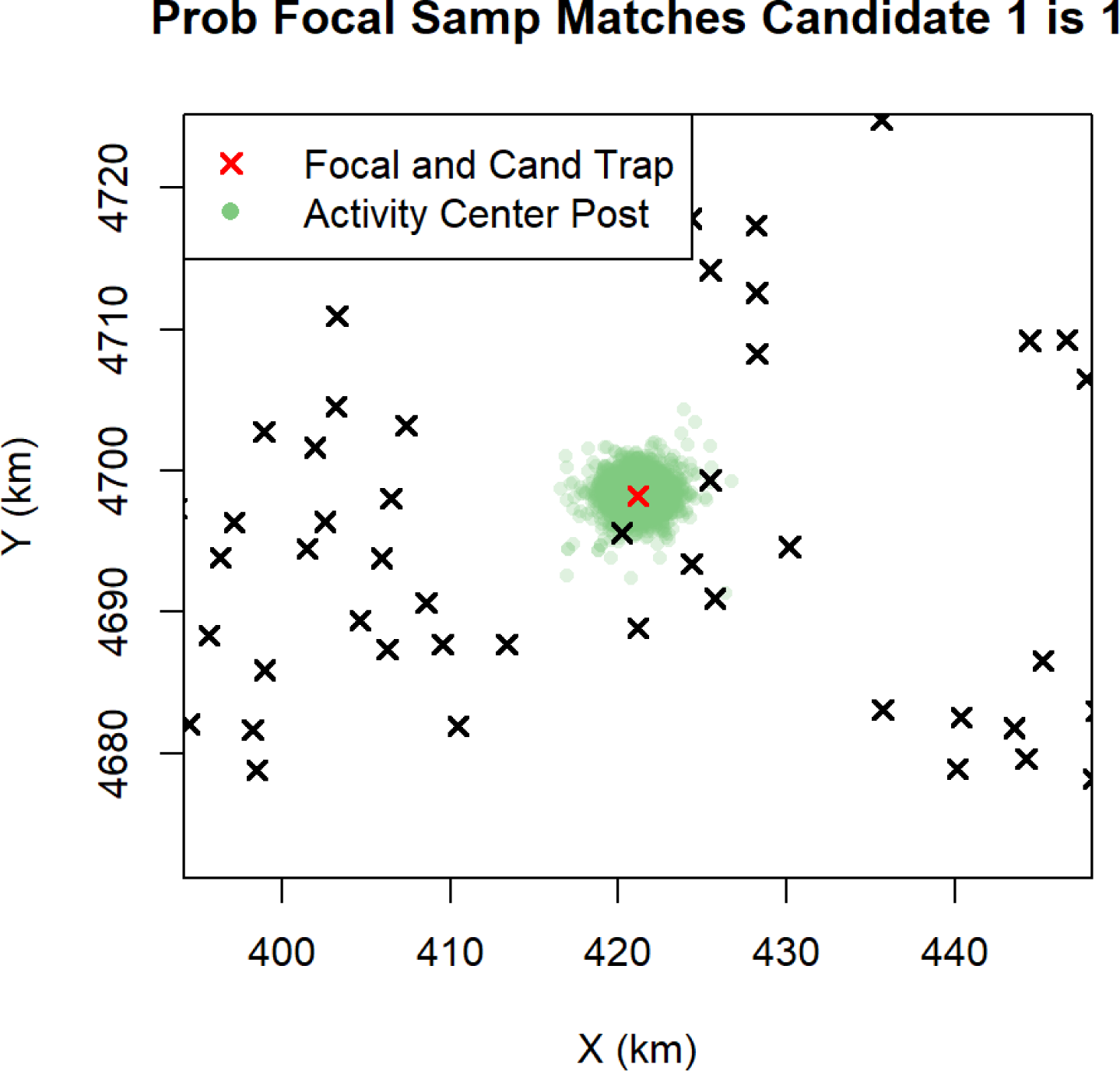

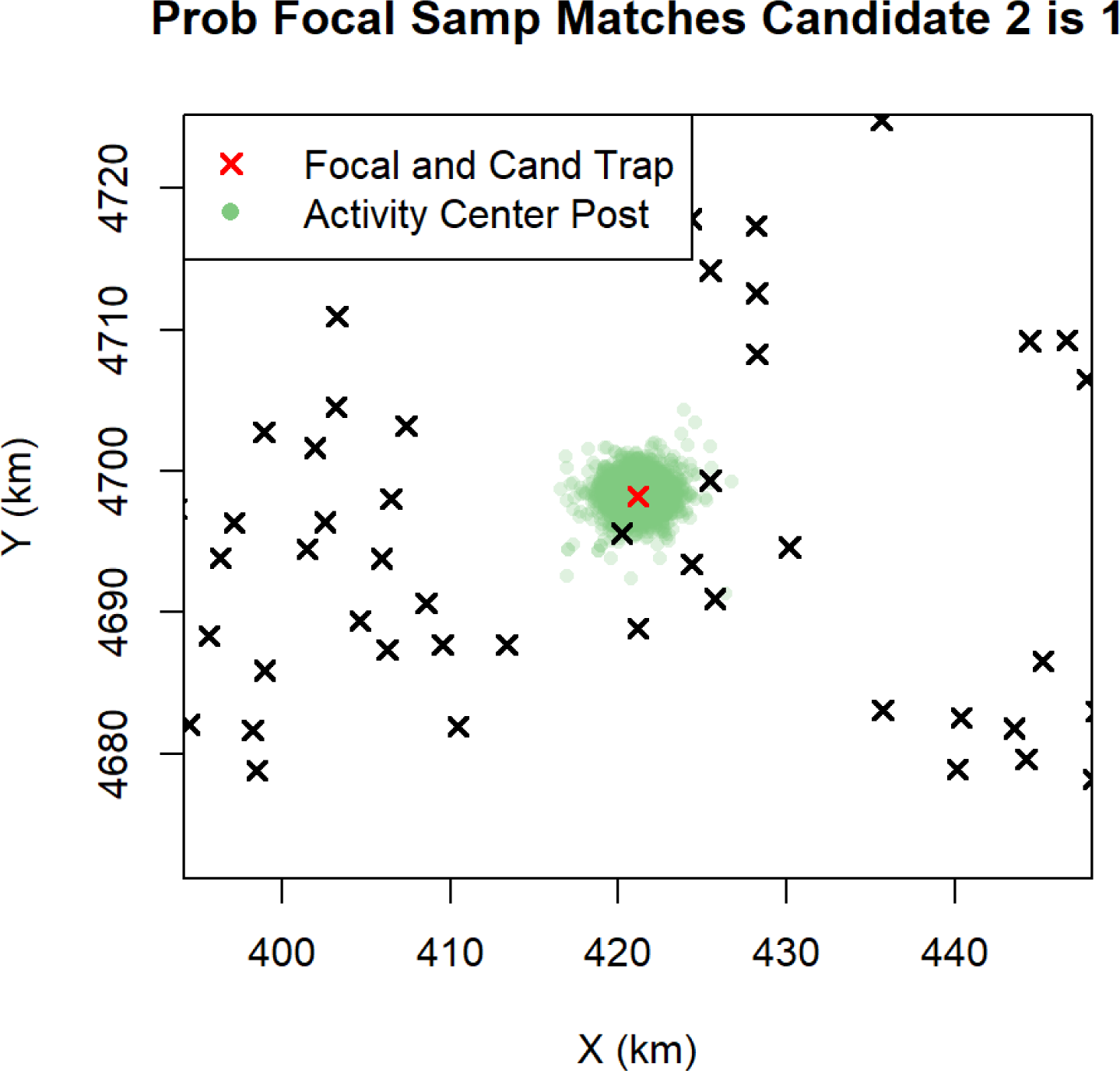

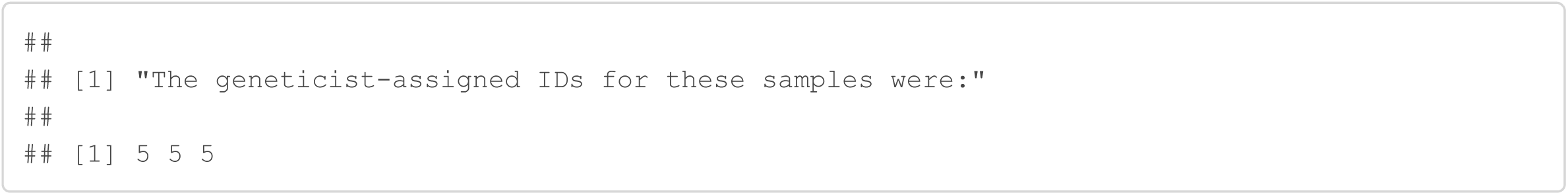

#### Samples 281 and 282

The geneticist grouped samples 281 and 282 together, which implies there were 4 allelic dropouts at locus 1 and 3 allelic dropouts at locus 3 for sample 282. The Genotype SPIM assigned a probability of 0.71 to this grouping, allowing a 0.29 probability these samples belonged to 2 different individuals and there were no or fewer allelic dropouts. This example illustrates the fact that the number of allelic dropout events across replicate assignments that lead to locus mismatches between samples provide critical information about the probability the two samples were produced by the same individual. Seven allelic dropout events under the state that the two samples came from one individual was enough to give some probability that the two samples came from two different individuals without the allelic dropout events. If there was more replication for sample 282 producing more homozygous scores, we would be more confident in the assignment of this sample to another individual than that which produced sample 281.

The assignment of a non-negligible probability that sample 282 was a distinct individual from 281 is also likely due to the mismatching scores for sample 281 being relatively common genotypes. 156.156 at locus 1 had an estimated frequency of 0.0523 and 207.207 at locus 3 had an estimated frequency of 0.0948. If these loci were scored as allelic dropouts leading to genotypes that were much more rare in the population, Genotype SPIM would place much less probability that the two samples came from different individuals. The estimated genotype frequencies can be found in the last section of this document.

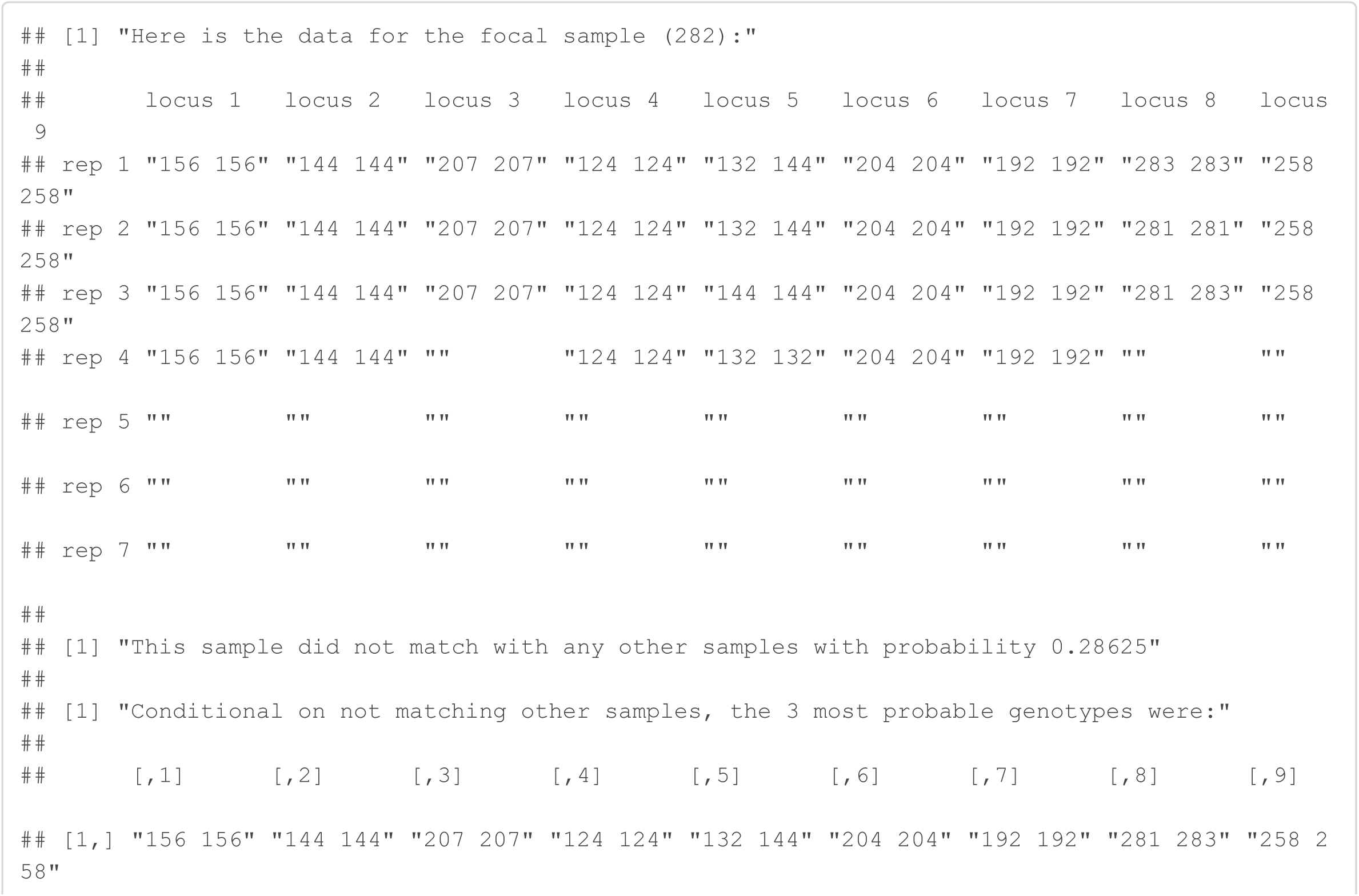

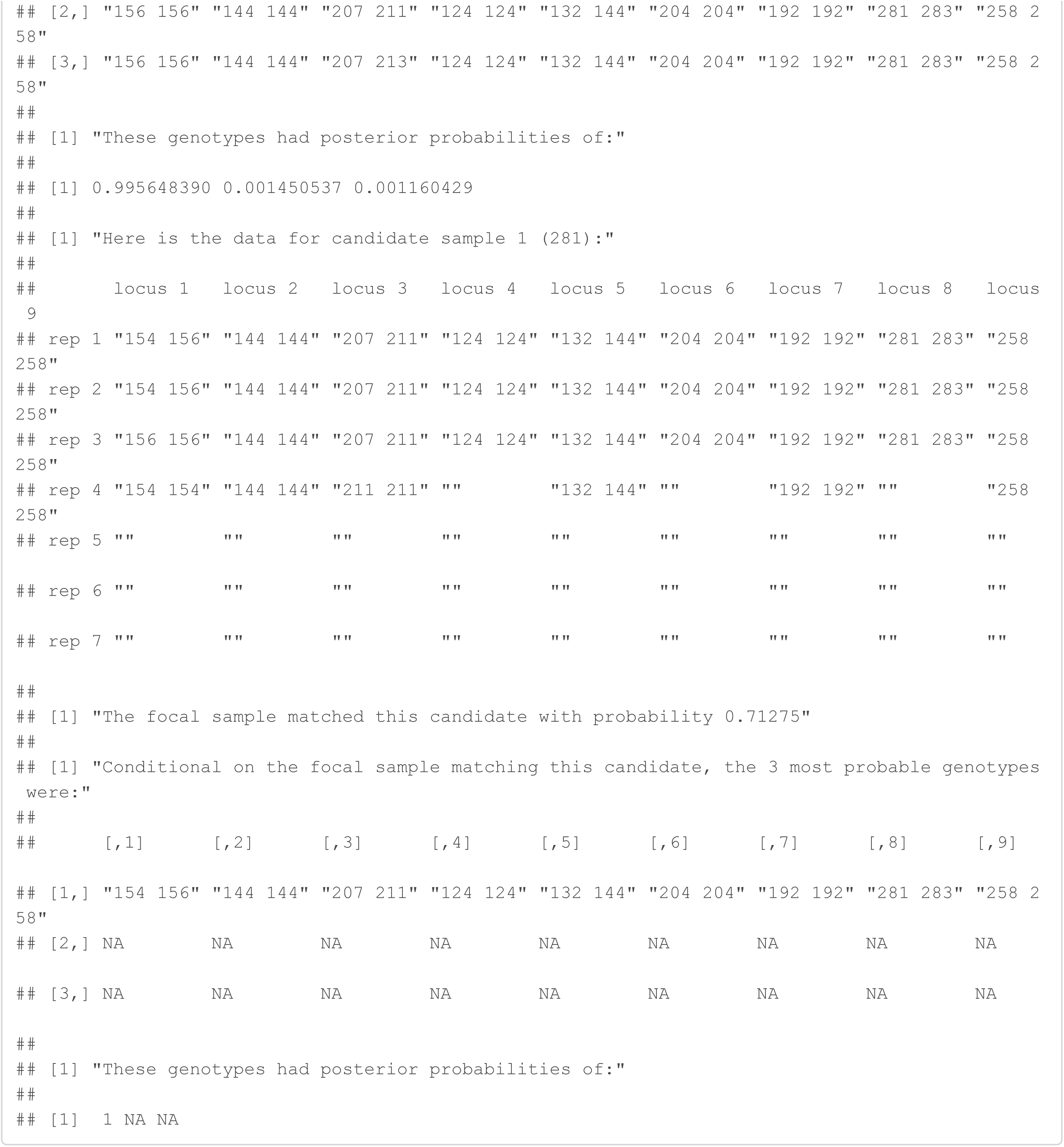

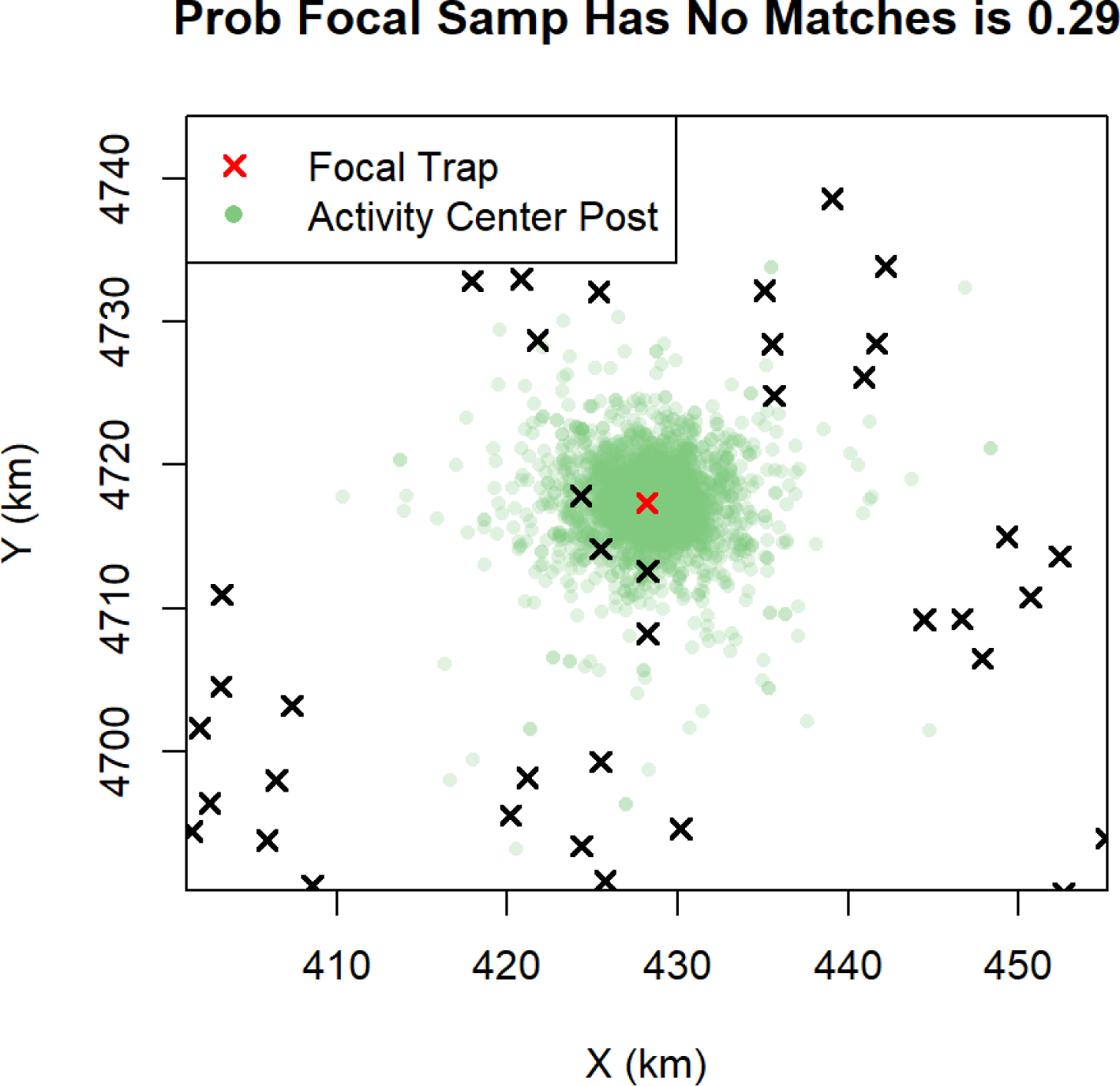

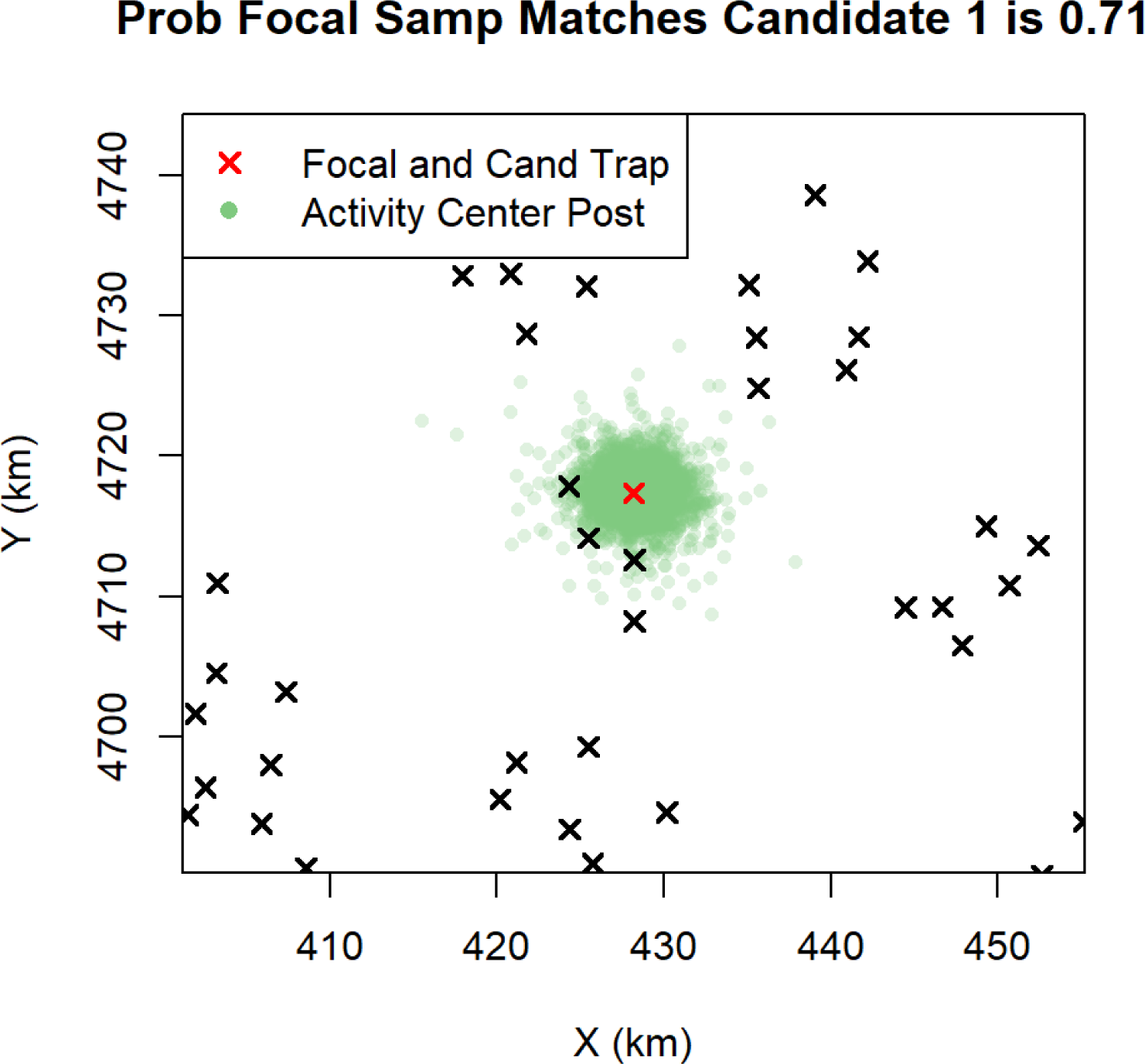

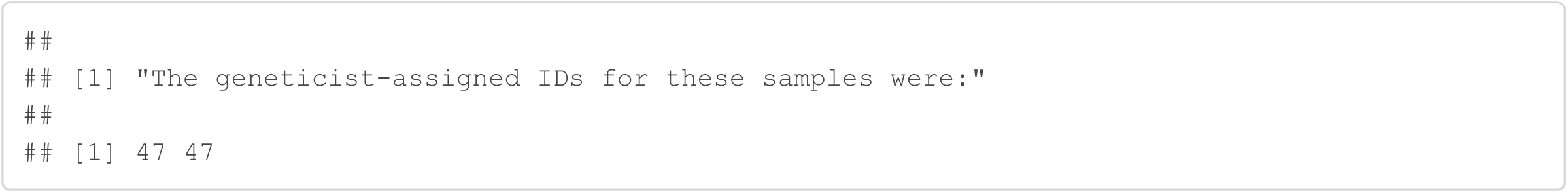

#### Samples 46 and 82

The geneticist assigned different individual identities to samples 46 and 82, but the Genotype SPIM assigned them the same identity with probability 1. The discrepancy between the samples that led the geneticist to assign different individual identities is found at locus 6 where sample 82 was scored as an allelic dropout 4 times if these samples were in fact the same individual. This is not an unlikely set of events because sample 82 is a “low quality” sample with a high allelic dropout probability. Both of these samples were scored as “male”, providing further support that these two samples came from the same individual.

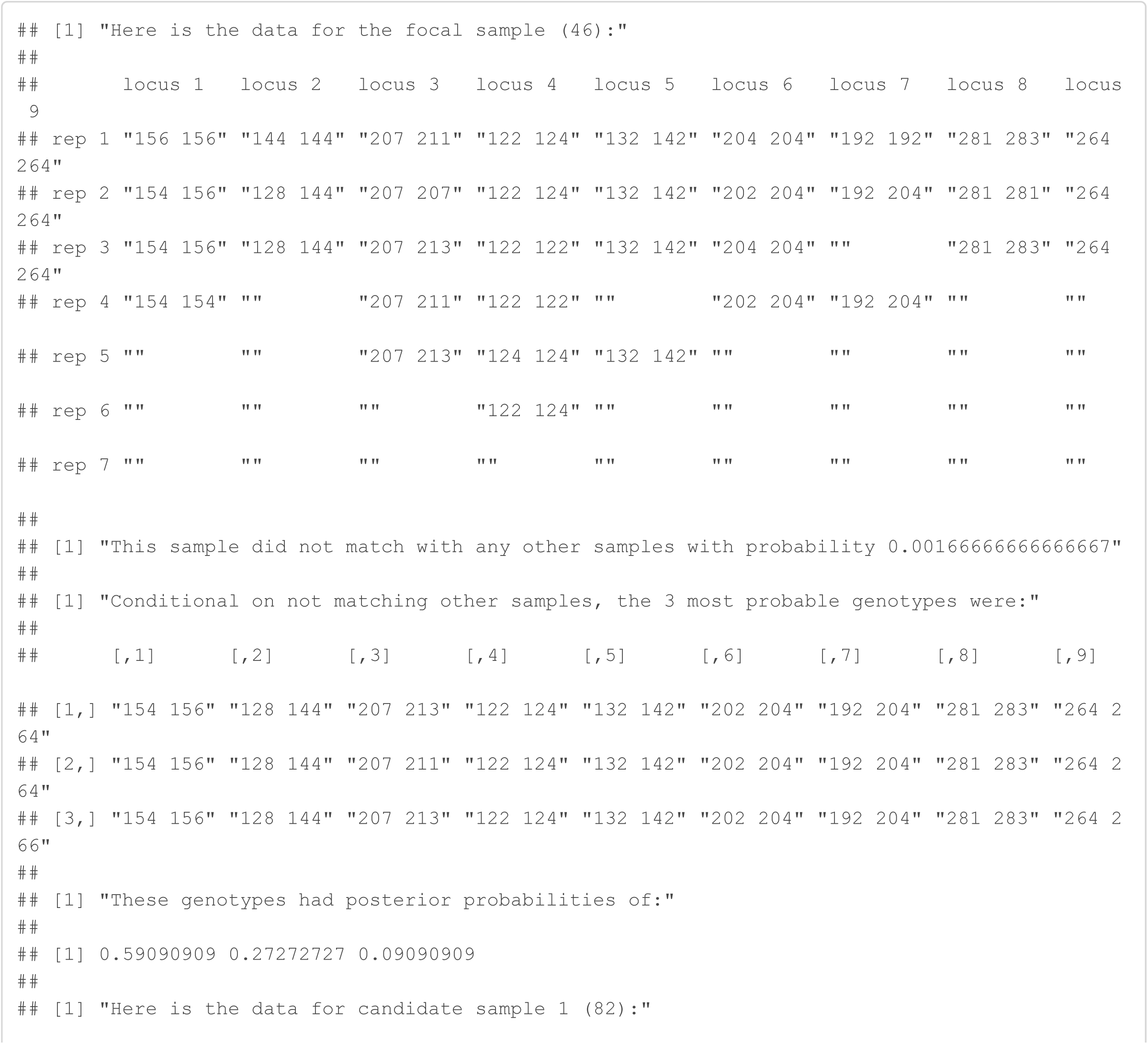

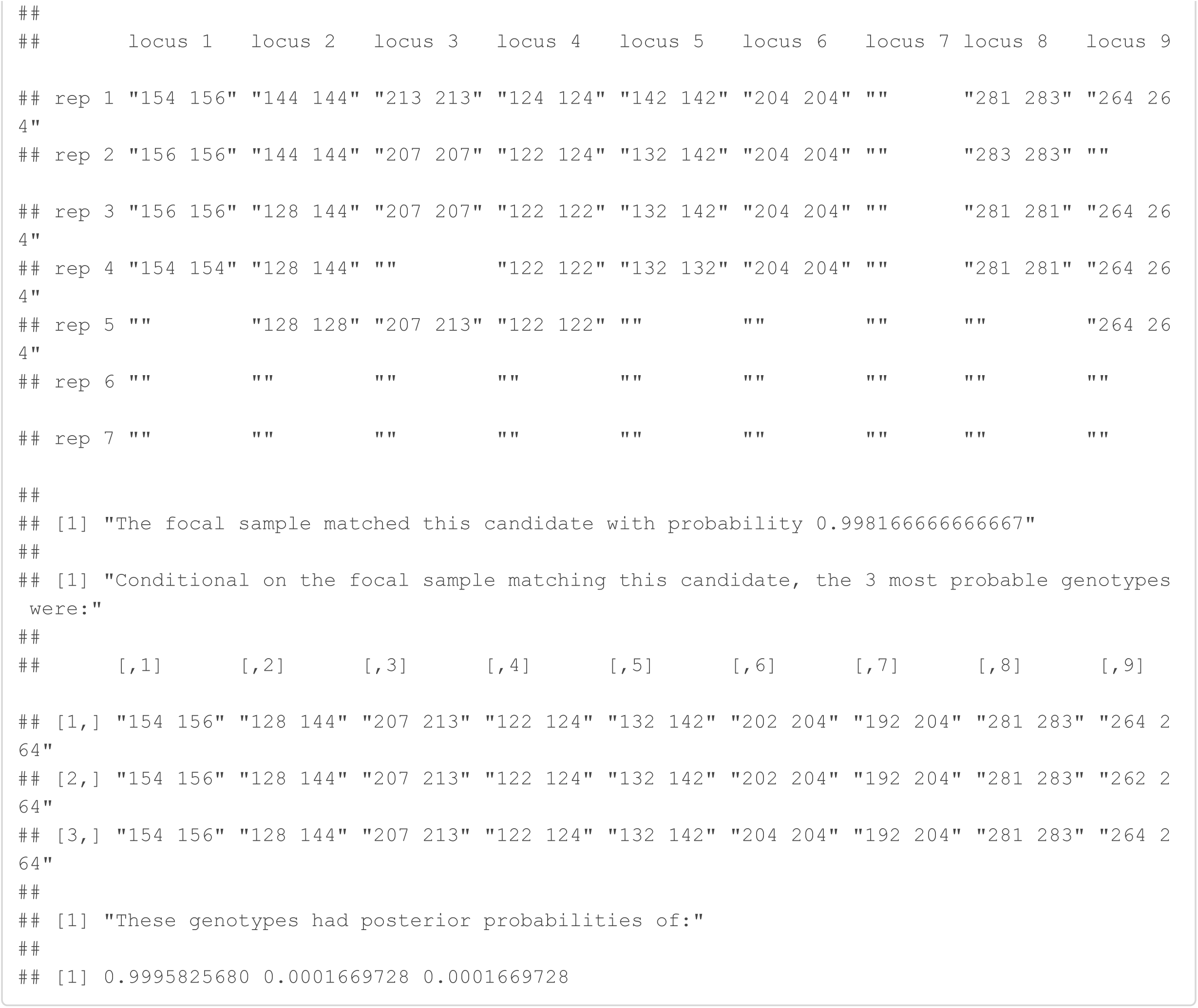

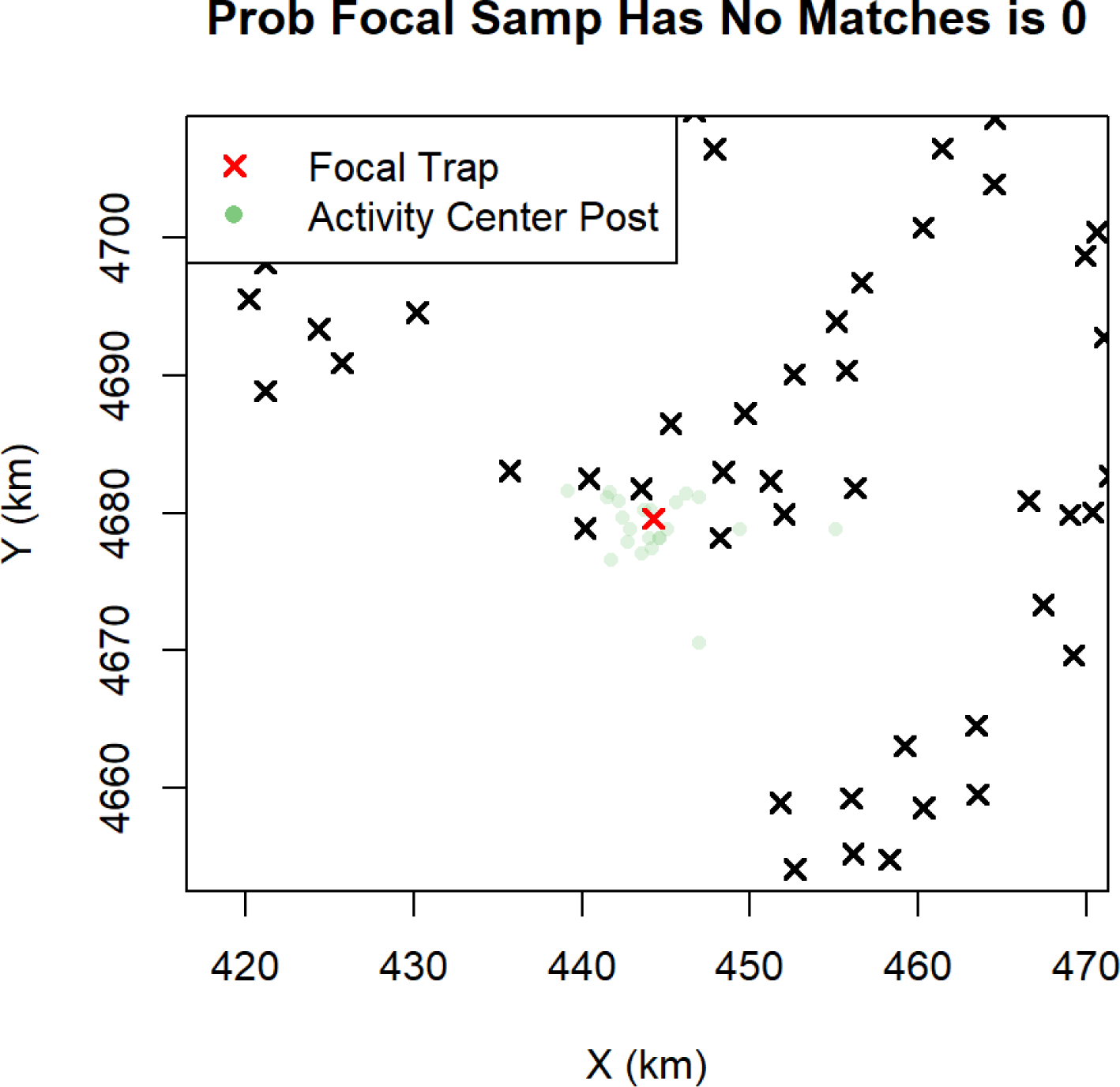

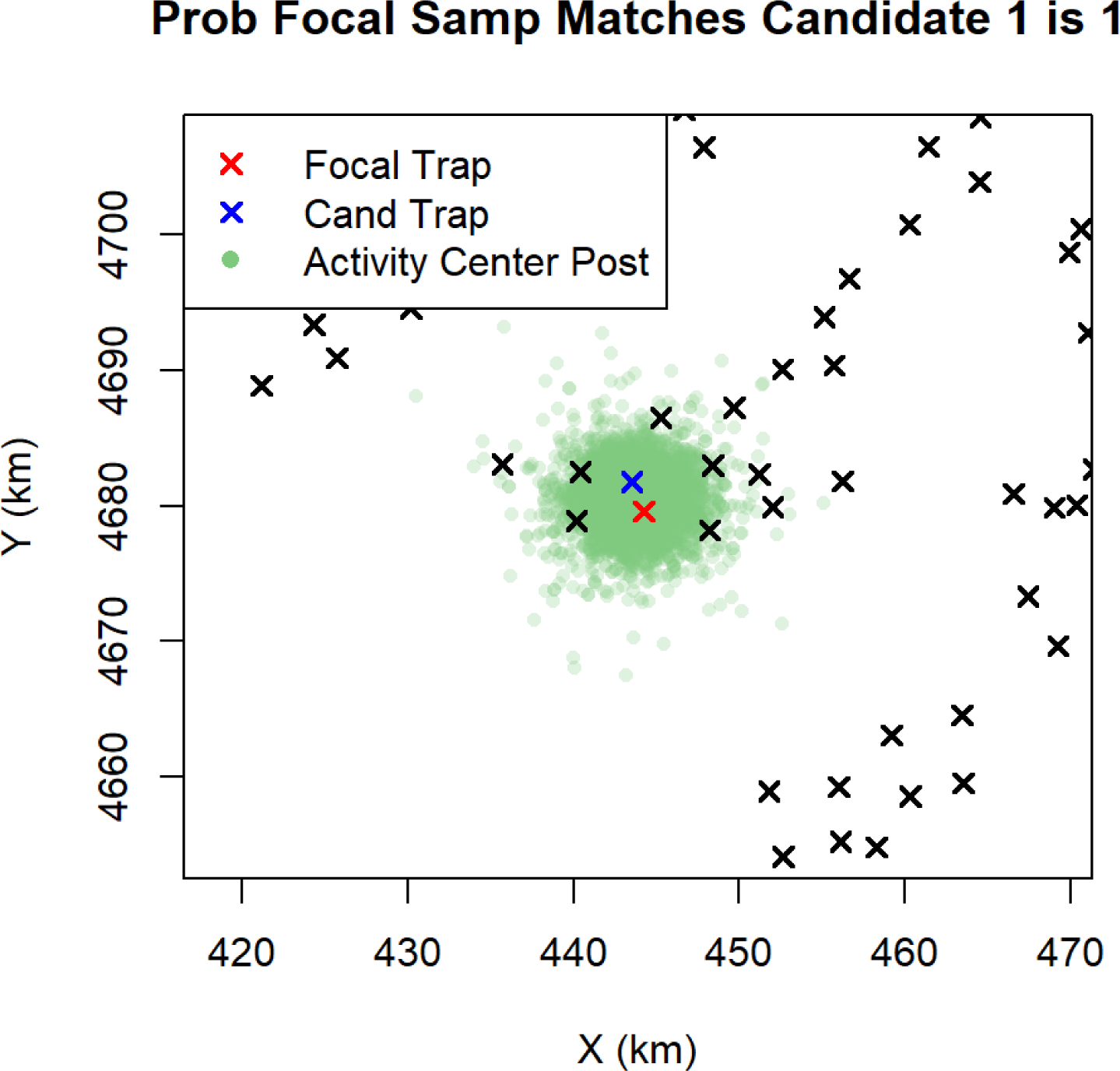

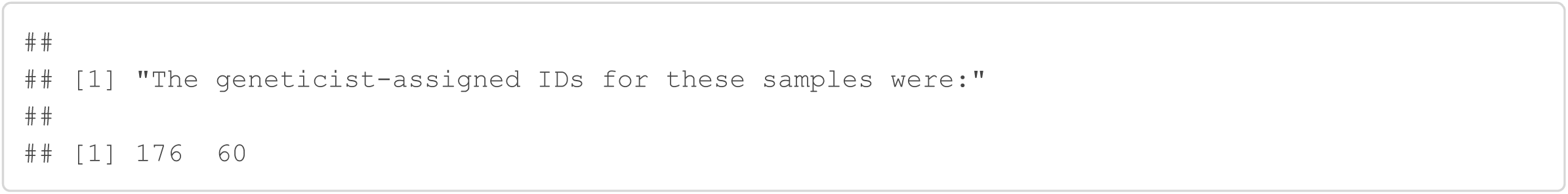

#### Samples 214 and 215

The geneticist assigned different individual identities to samples 214 and 215, an event that Genotype SPIM placed a probability of 0.08. The Genotype SPIM placed a probability of 0.92 that these two samples came from the same individual. The discrepancies between these two samples can be found at loci 3 and 9. If these samples belong to the same individual, there were 4 allelic dropout events at locus 3 for sample 214 and 4 allelic dropout events for sample 215 at locus 9. Because allelic dropout events were not rare, the Genotype SPIM places a much higher probability that these samples belong to the same individual than different individuals, but this later event is not conclusively ruled out. The sexes for both of these samples were female, further supporting the conclusion of the Genotype SPIM.

Interestingly, locus 9 for sample 214 was assigned the consensus 264.266 by the geneticist despite the scoring rules suggesting that it should have been called 264.264 because the heterozygote was never seen and 264.264 was seen 4 times. We are unaware of a scoring rule for calling a heterozygote when each corresponding homozygote is seen a certain number of times. No rule like this is mentioned in the original study. In this case, the presumed deviation from the typical scoring rules resulting in the most probable score being assigned, 264.266, and the typical scoring rules would have required assigning 264.264, which is very improbable.

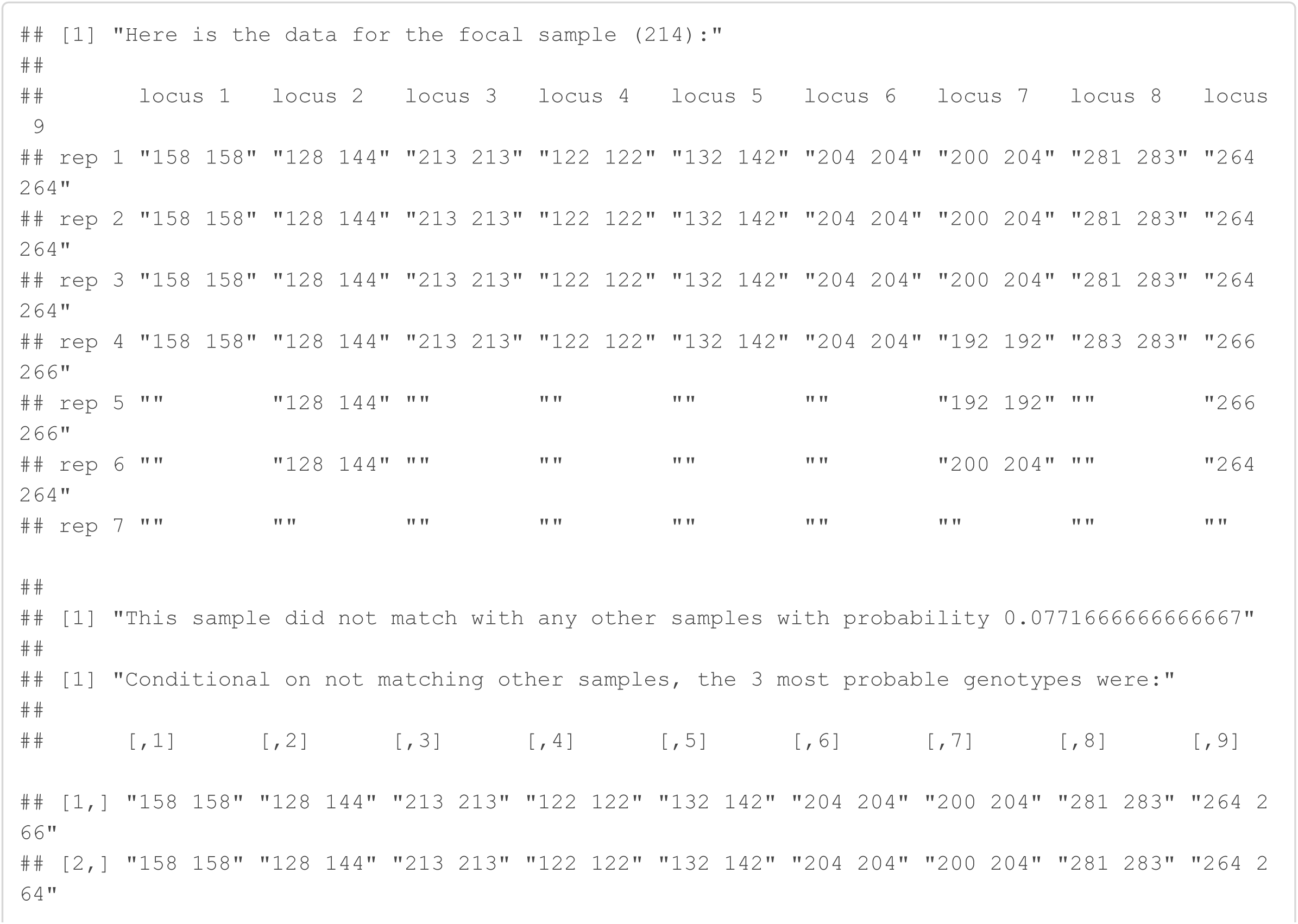

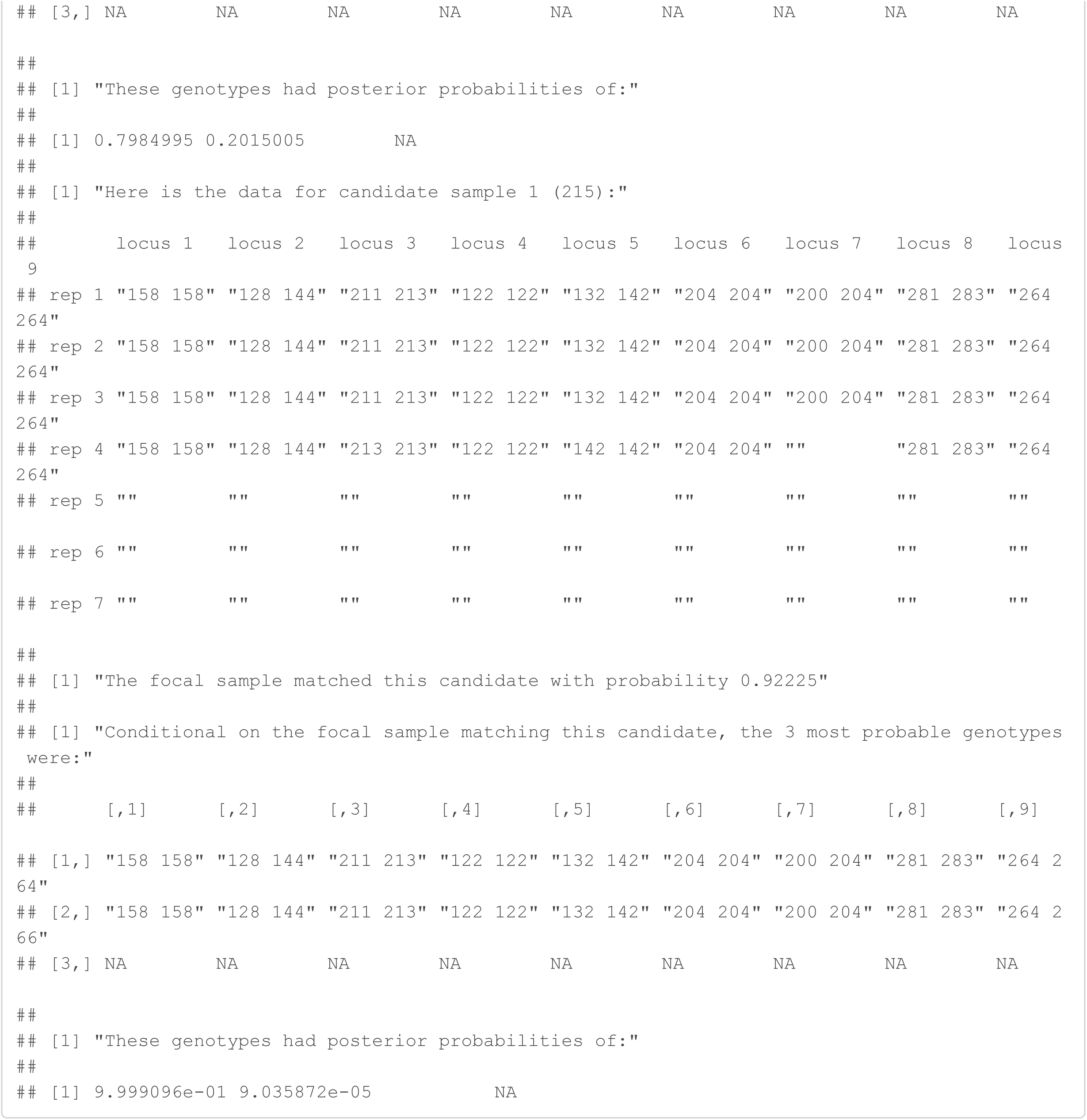

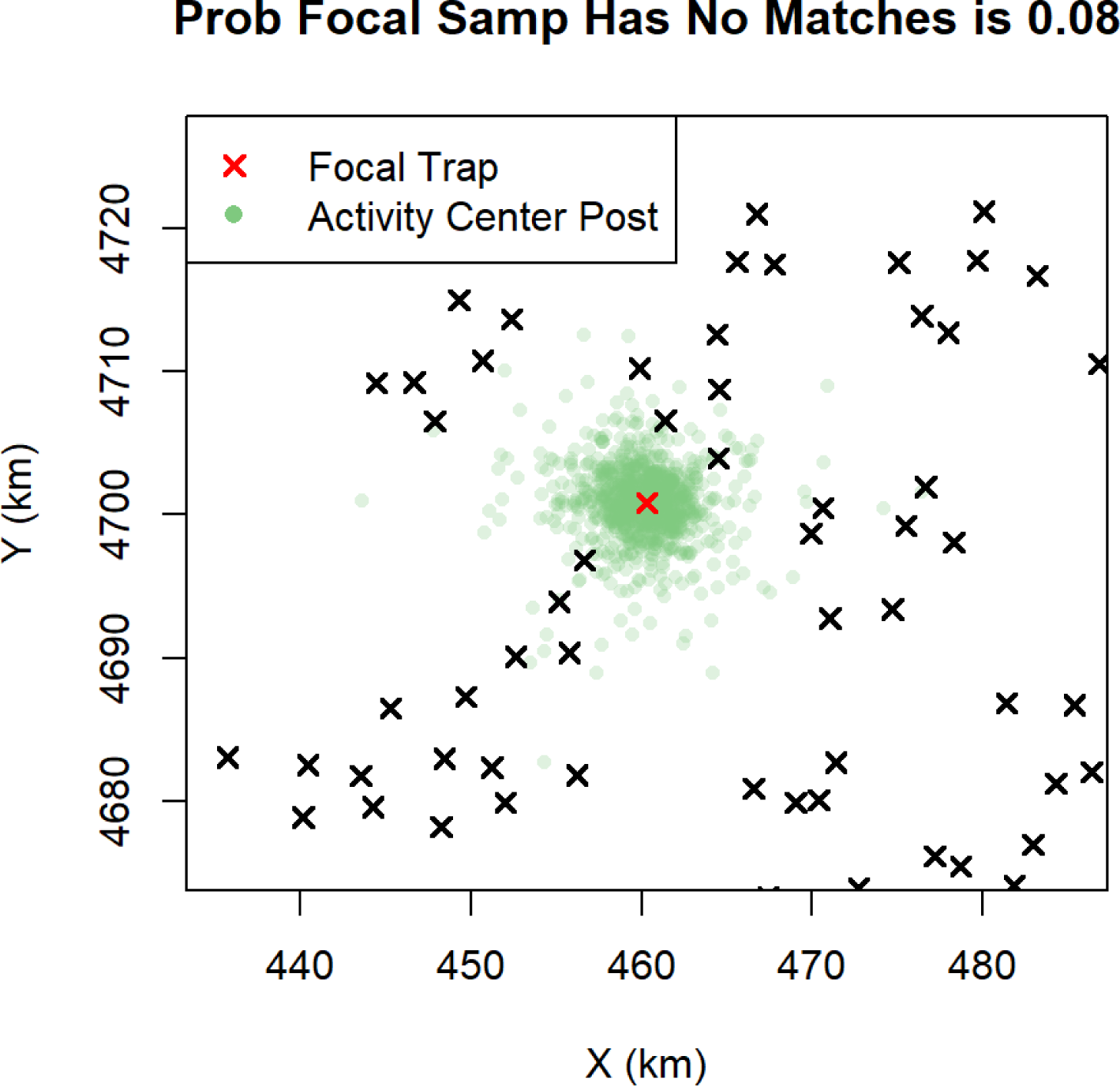

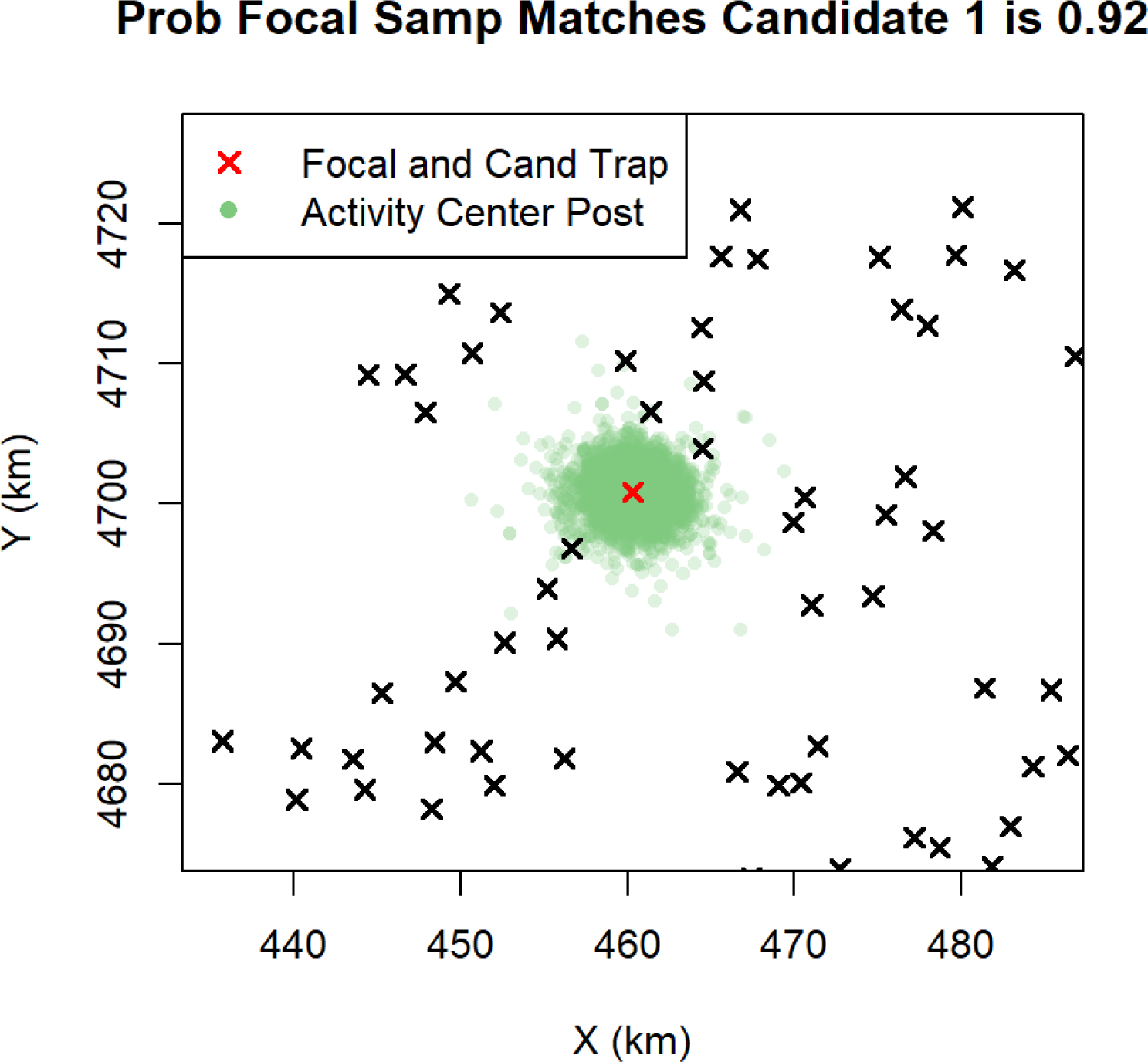

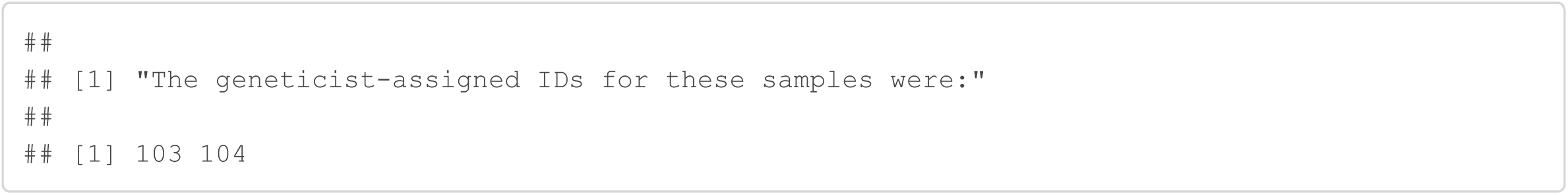

### High probability matches from samples discarded by geneticist to samples originally used

Here, we will look at some high probability assignments from the samples originally discarded by the geneticist because a certain identity assignment could not be made.

#### Sample 51

Sample 51 was originally discarded because it only amplified at 5/9 loci; however, the Genotype SPIM assigns it to the same individual as samples 42, 43, and 80 with probability 1 - a certain spatial recapture. For this to be the true state, there was most likely 2 allelic dropouts at locus 1 and 3 allelic dropouts at locus 6 for sample 51; however, allelic dropouts were estimated to be very likely for poor quality samples like sample 51. Thus, a partial genotype with only 5 amplified loci, 2 of which were scored incorrectly contains enough information to assign it an individual identity with certainty. Note samples 45 and 50, amplified at 0 and 1 locus, respectively, match the focal sample of 51 with probability 0.20 and 0.48, demonstrating that the spatial information alone carries substantial information about individual identity, at least in this low density population.

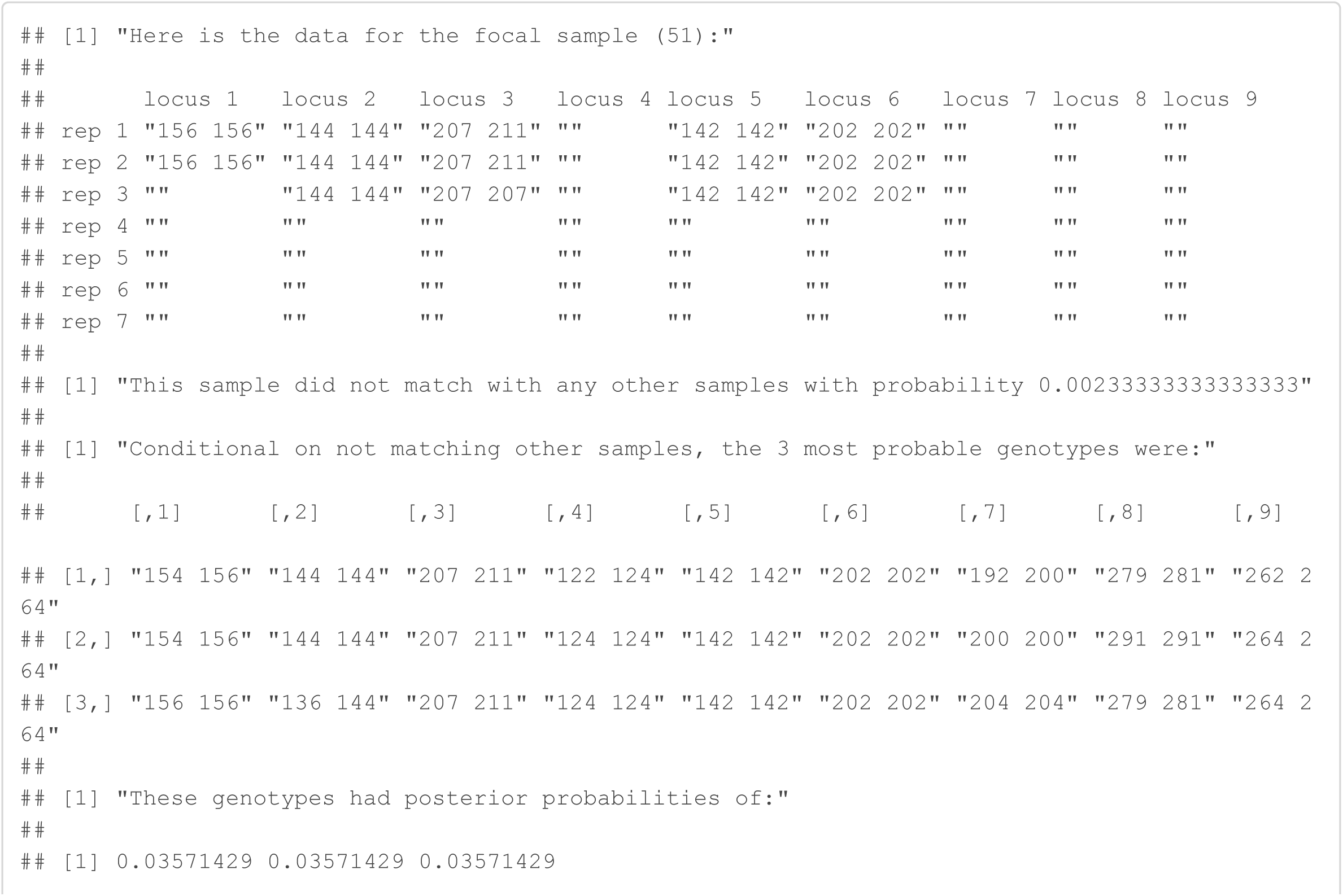

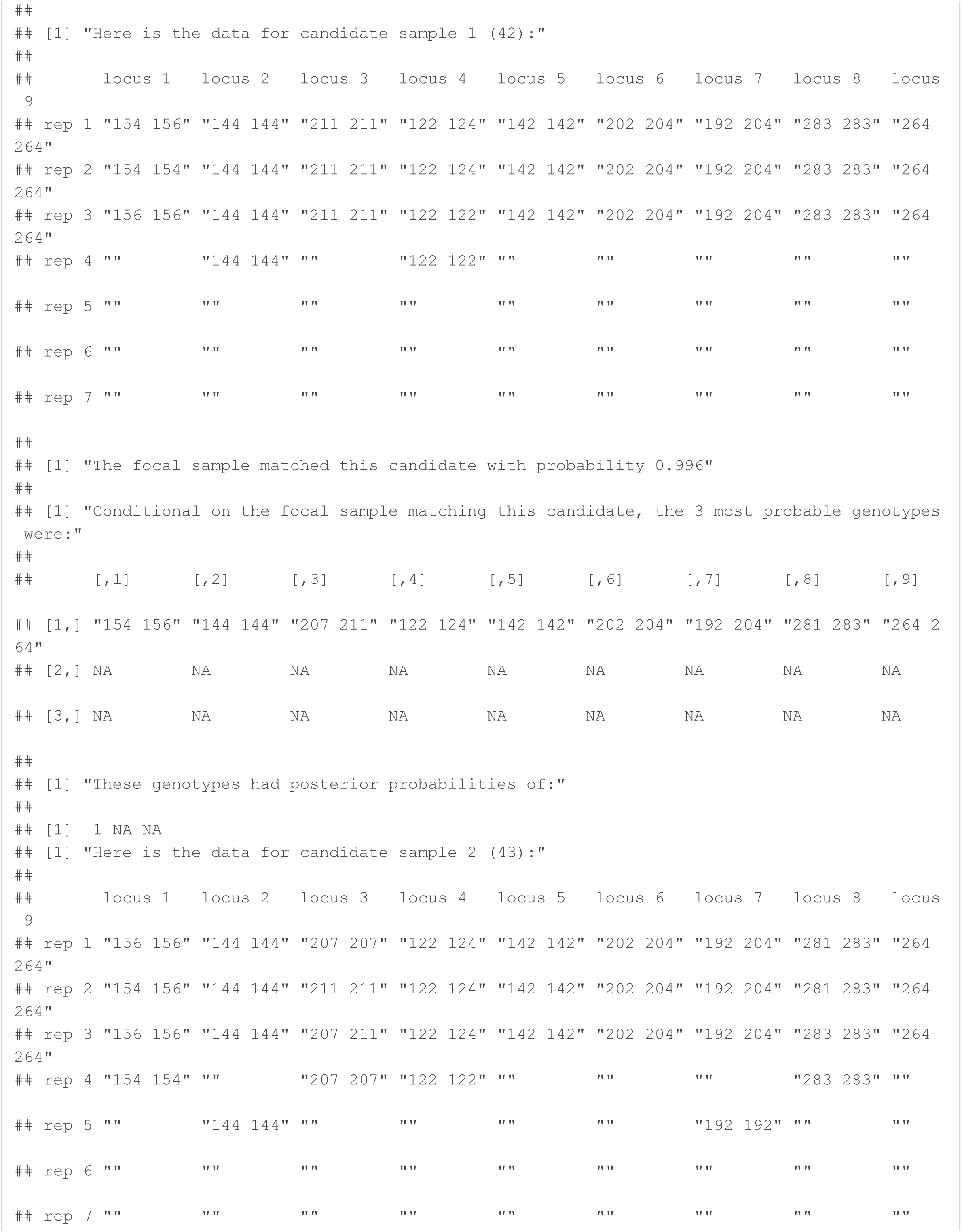

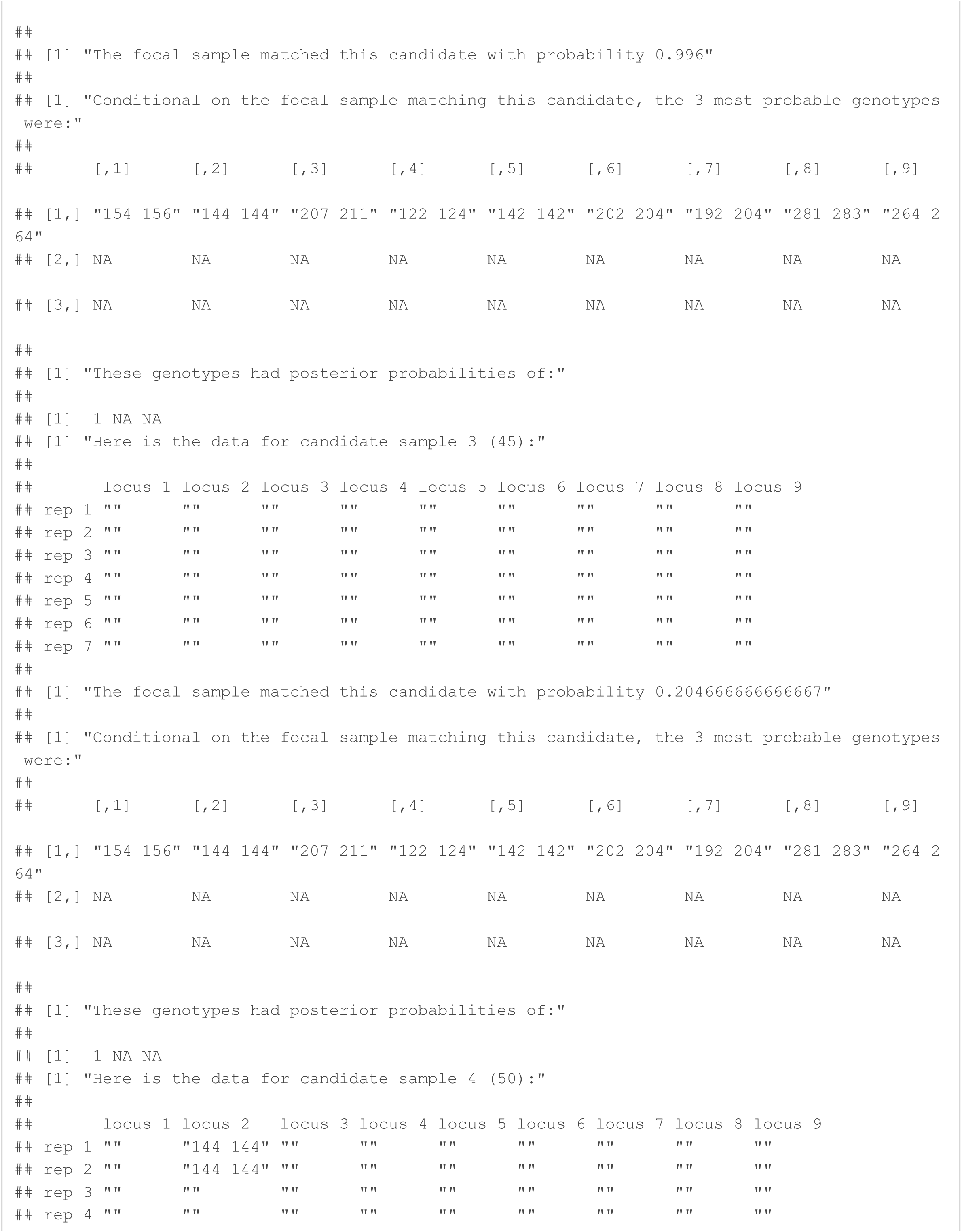

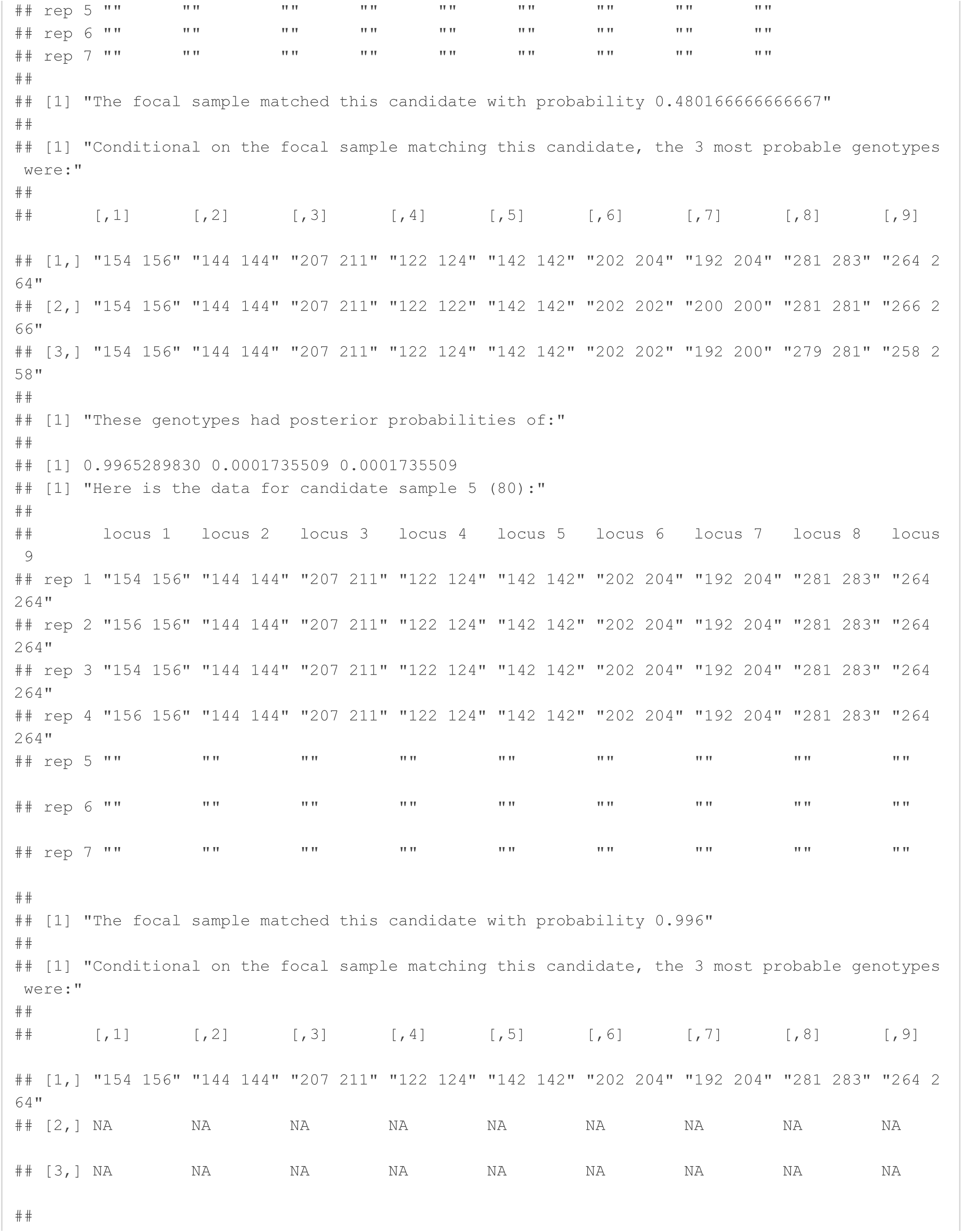

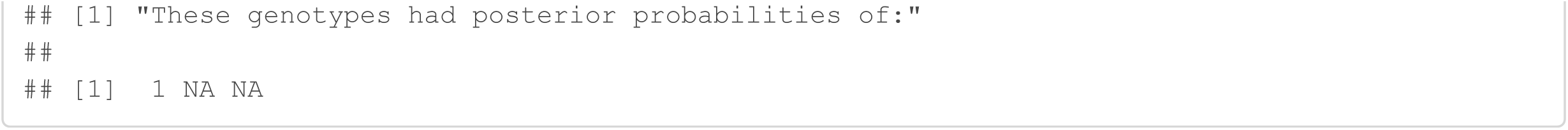

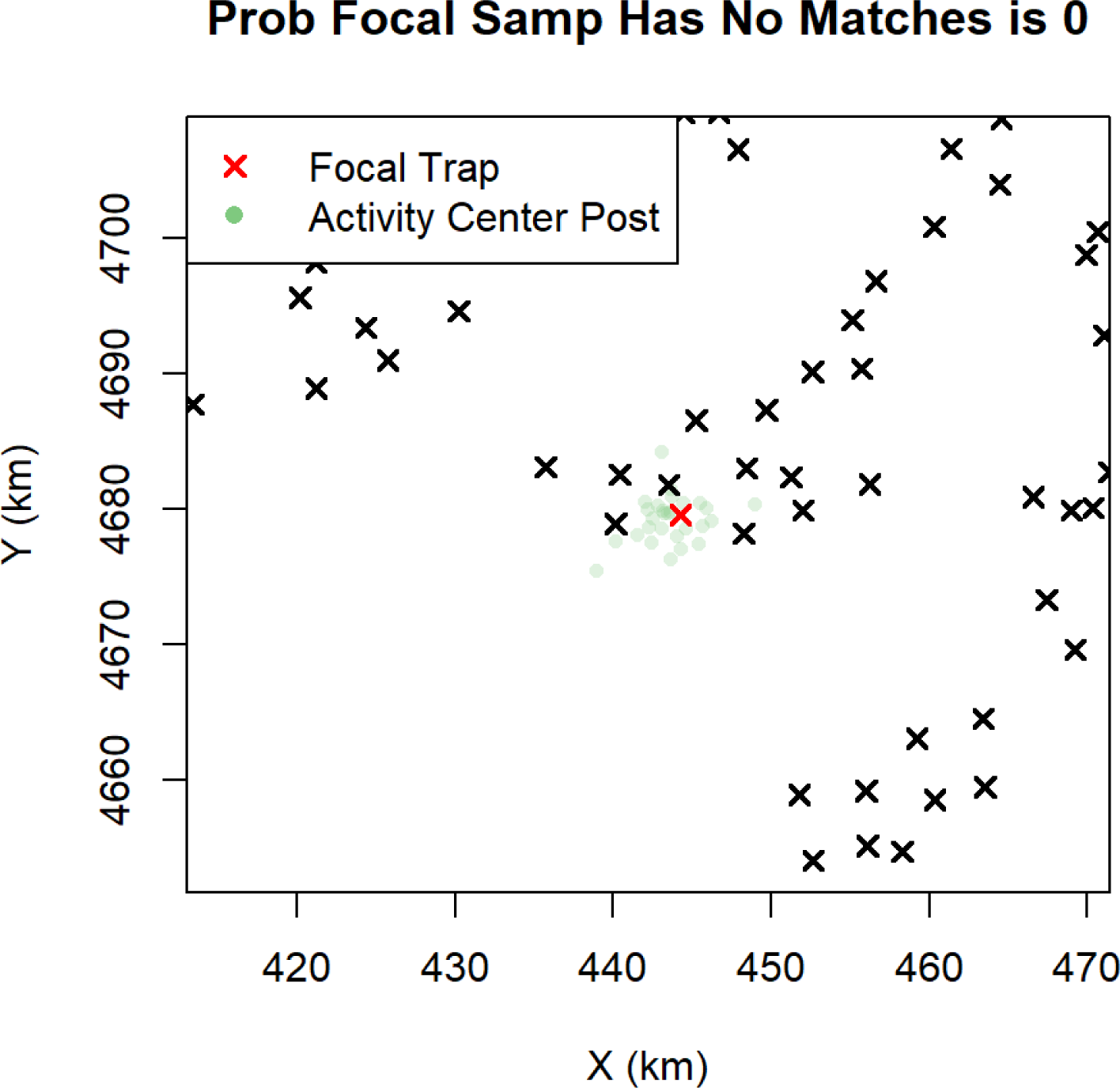

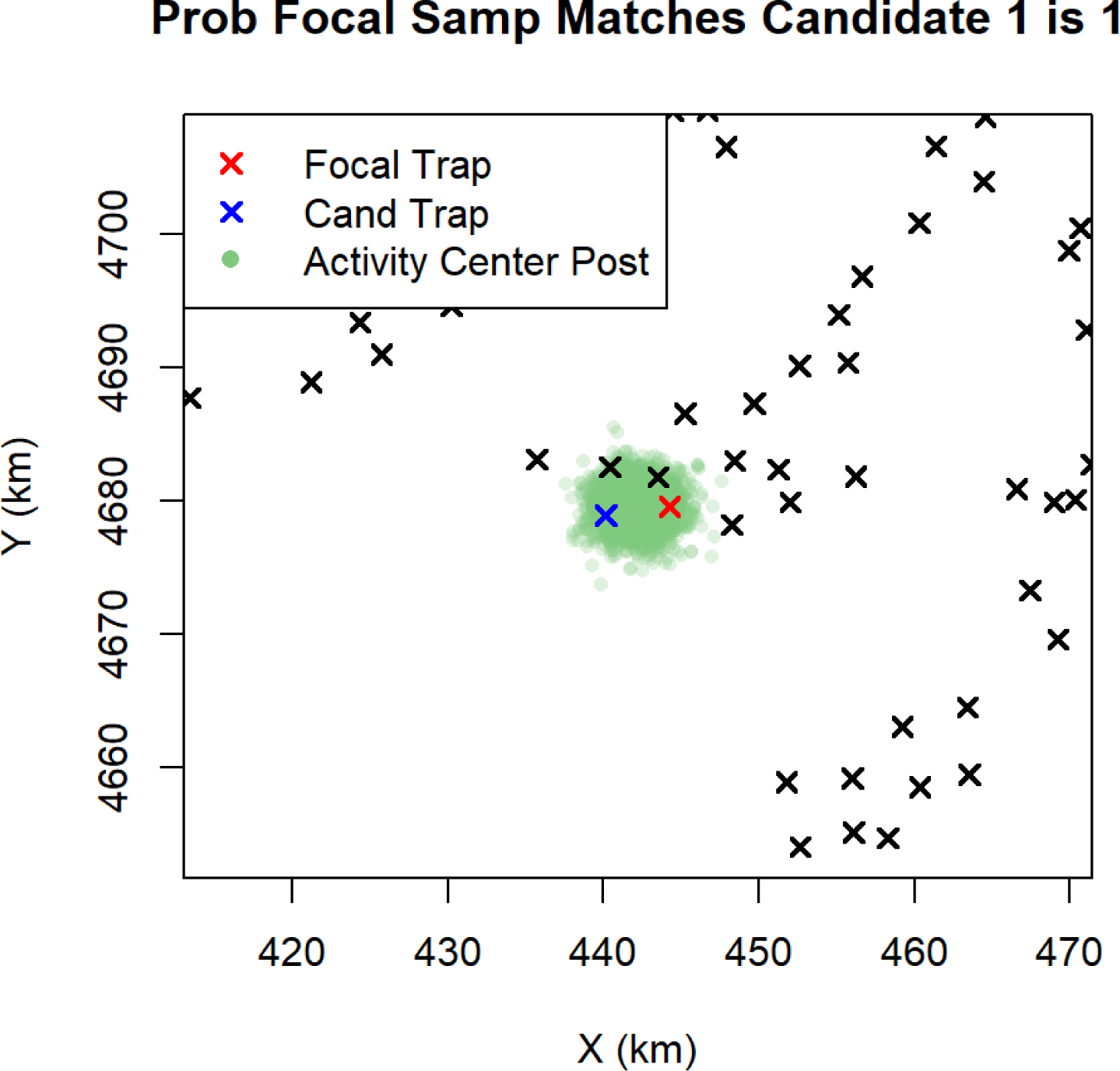

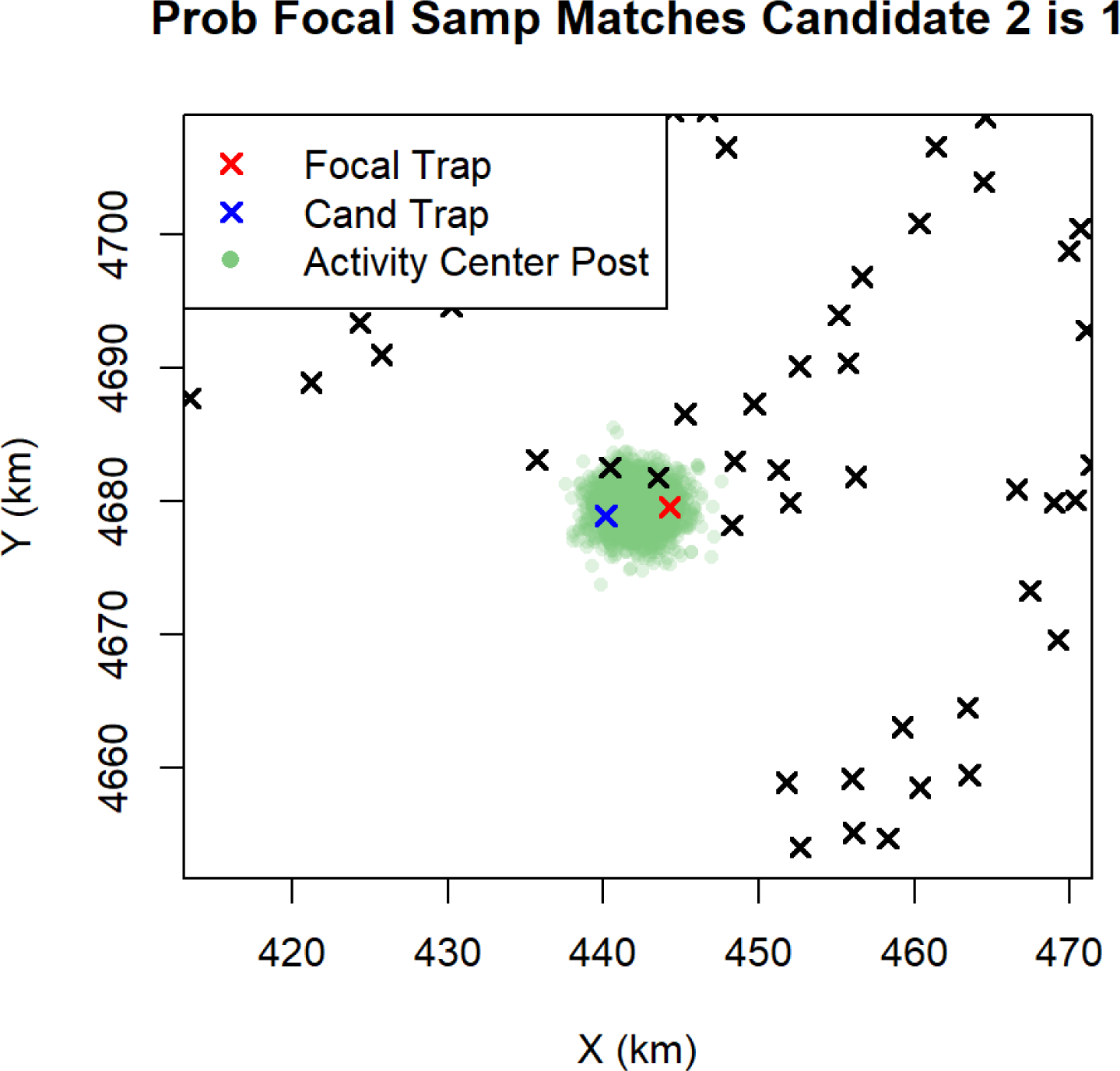

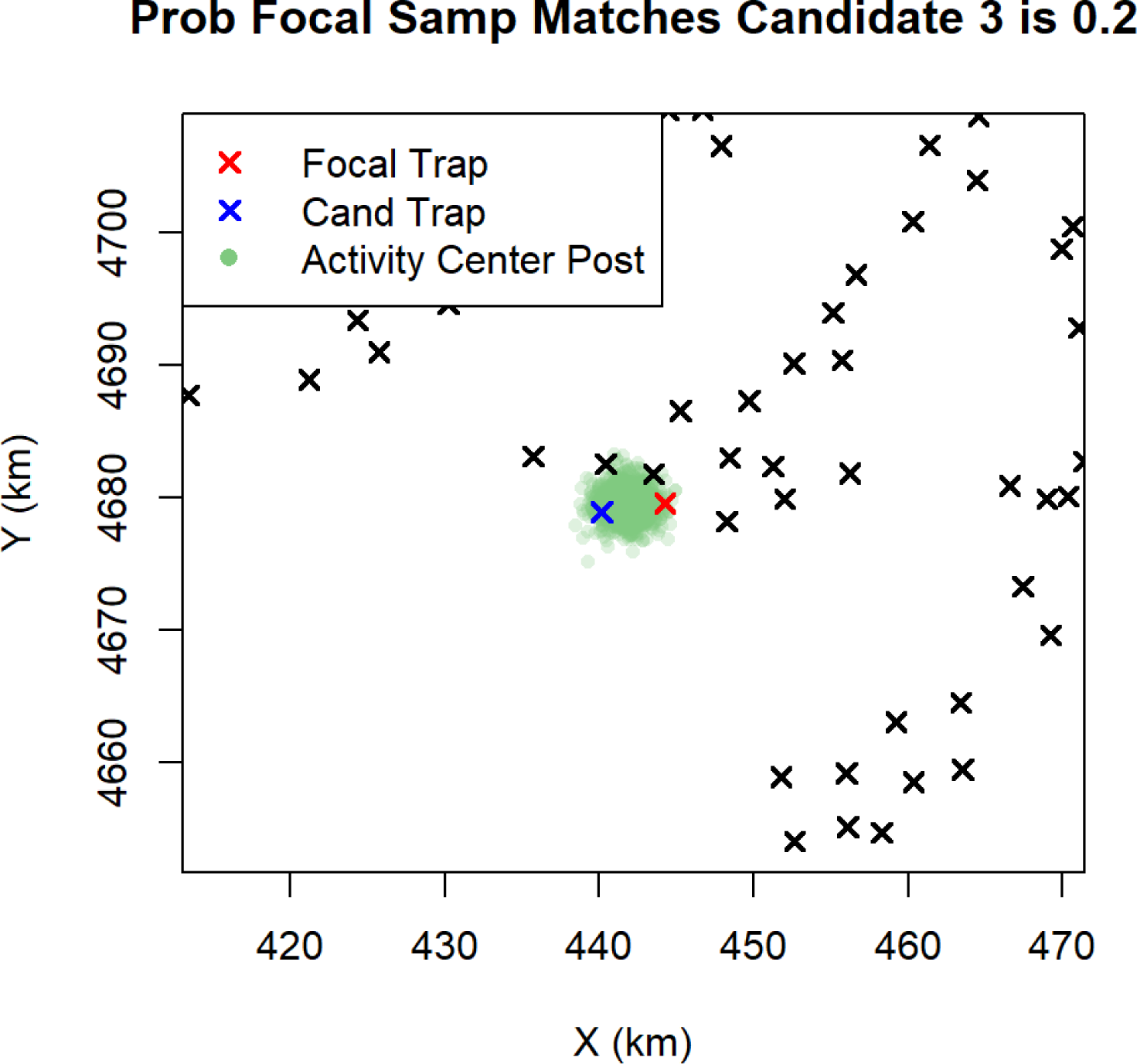

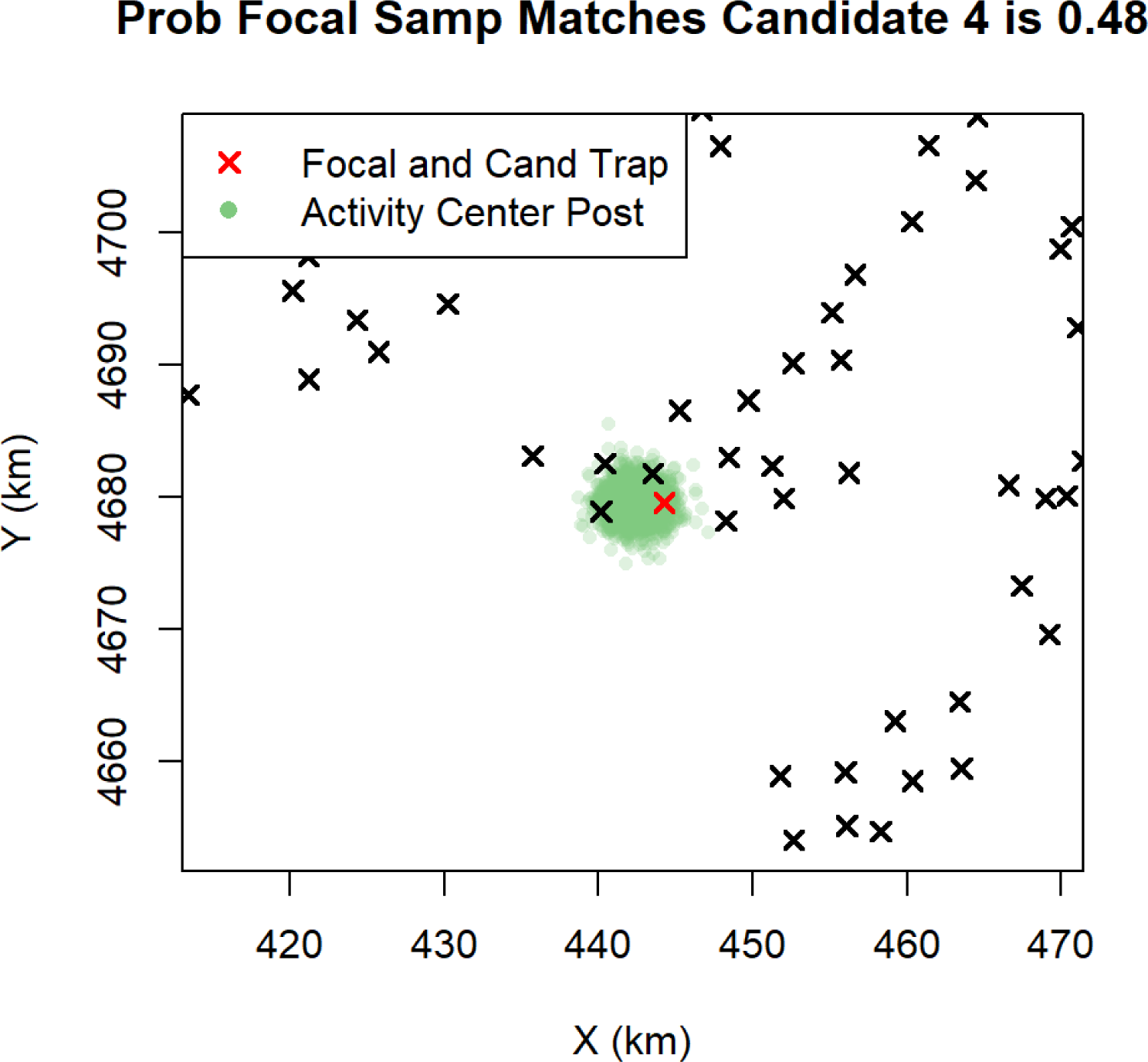

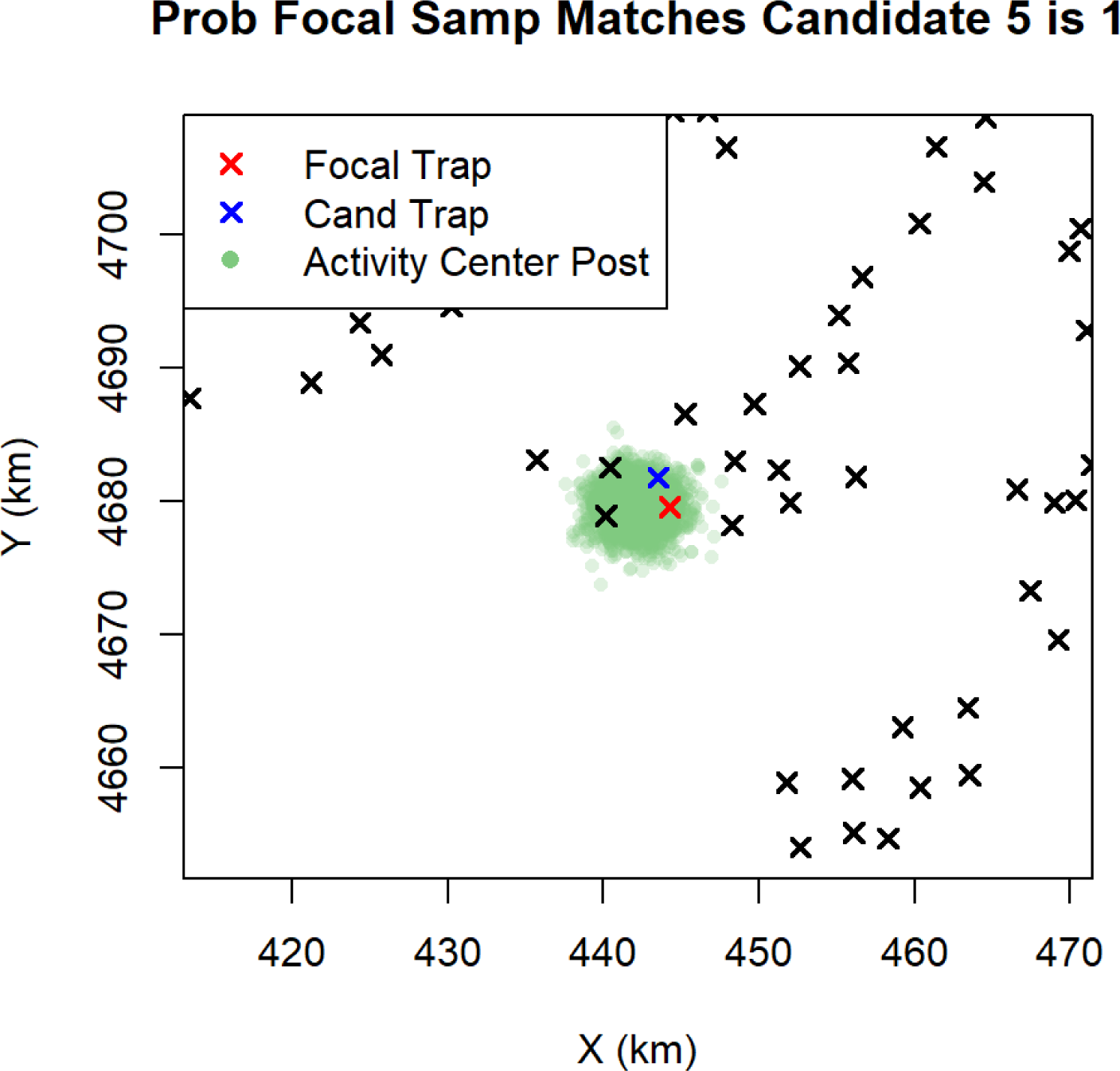

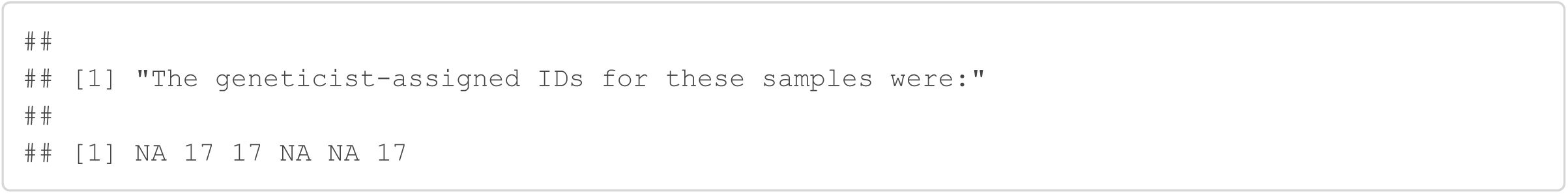

#### Sample 67

Sample 67 was originally discarded because it only amplified at 3/9 loci and it only amplified in 2 replications at each of these 3 loci. The Genotype SPIM assigns it to the same individual as sample 64, captured in the same trap, with probability 0.96. For this to be a correct match, it implies that both replicates for sample 67 at loci 2 and 6 were allelic dropouts and the only correct scores for sample 67 were the two at locus 7. The high probability of this match likely stems from the fact that the observed genotypes at locus 2 and 6 were estimated to be very rare, especially the genotype of 142.142 at locus 2, estimated to occur with probability 0.0002, while 142.144 was estimated to occur with probability 0.0315. The genotype frequency estimates can be found in the last section of this document.

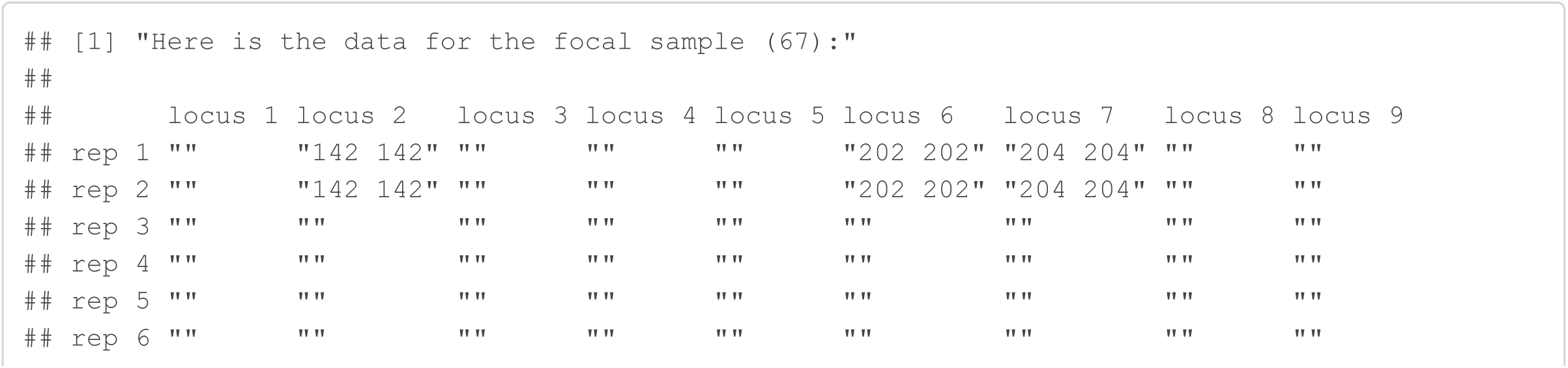

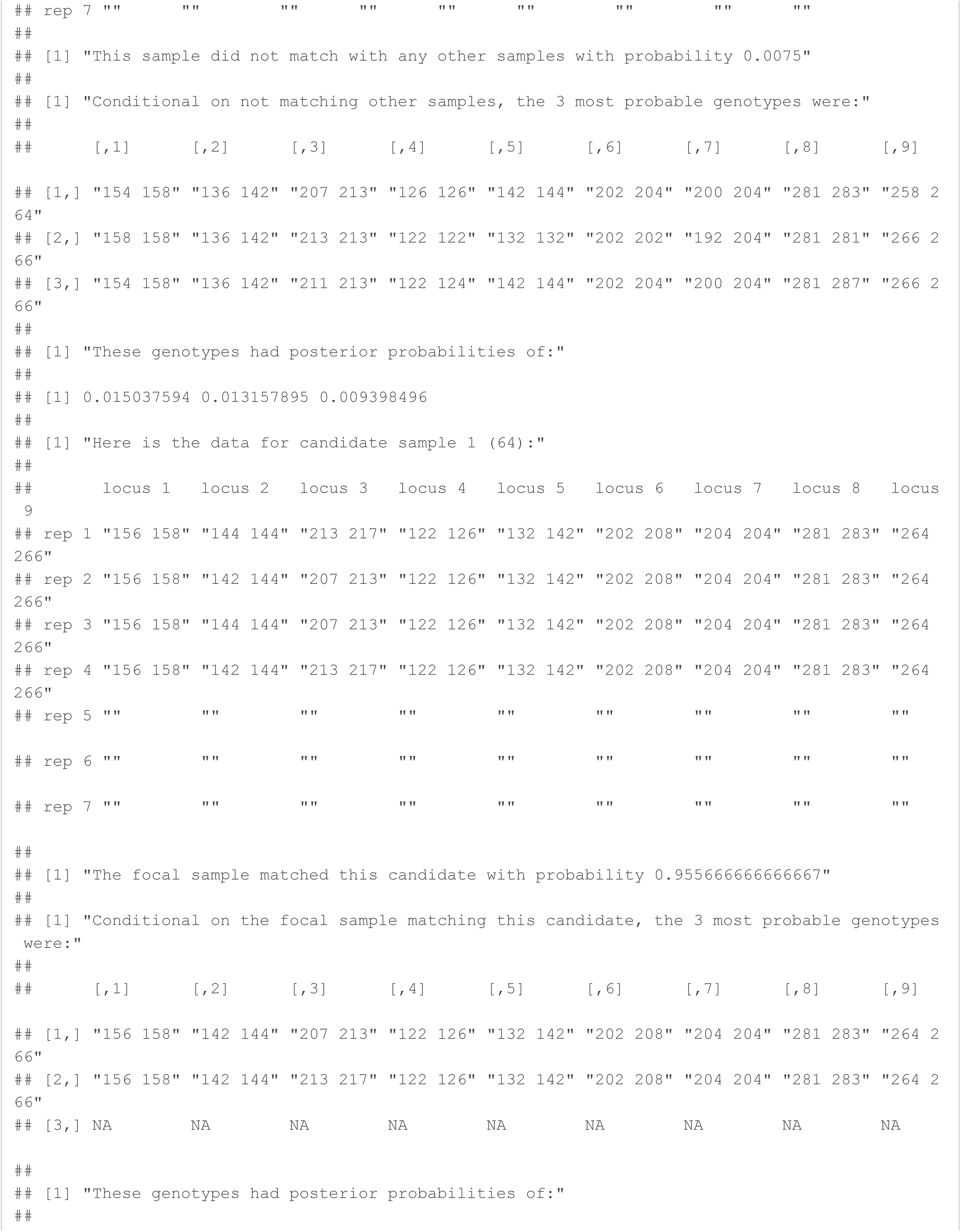

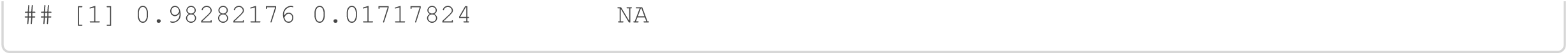

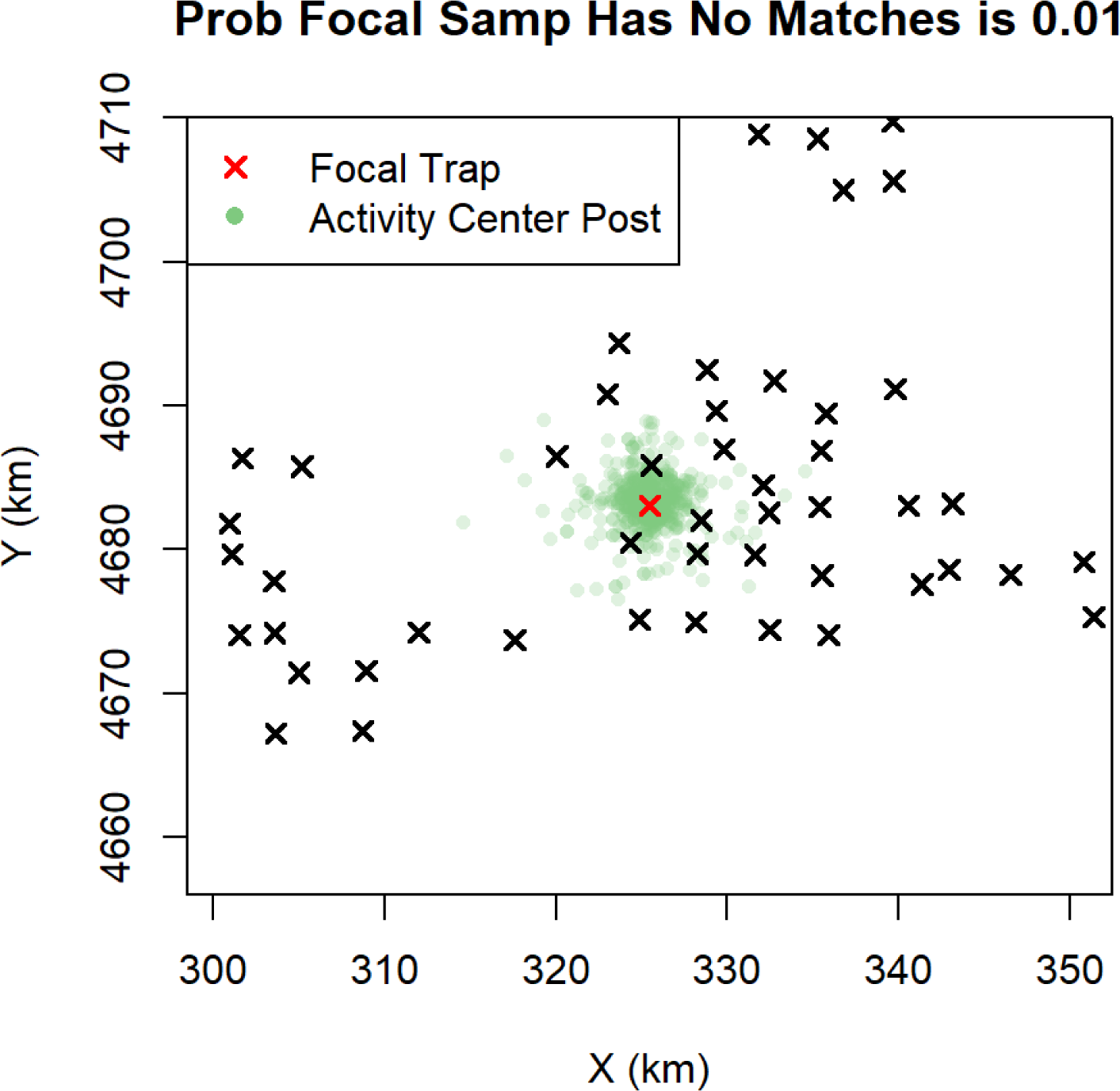

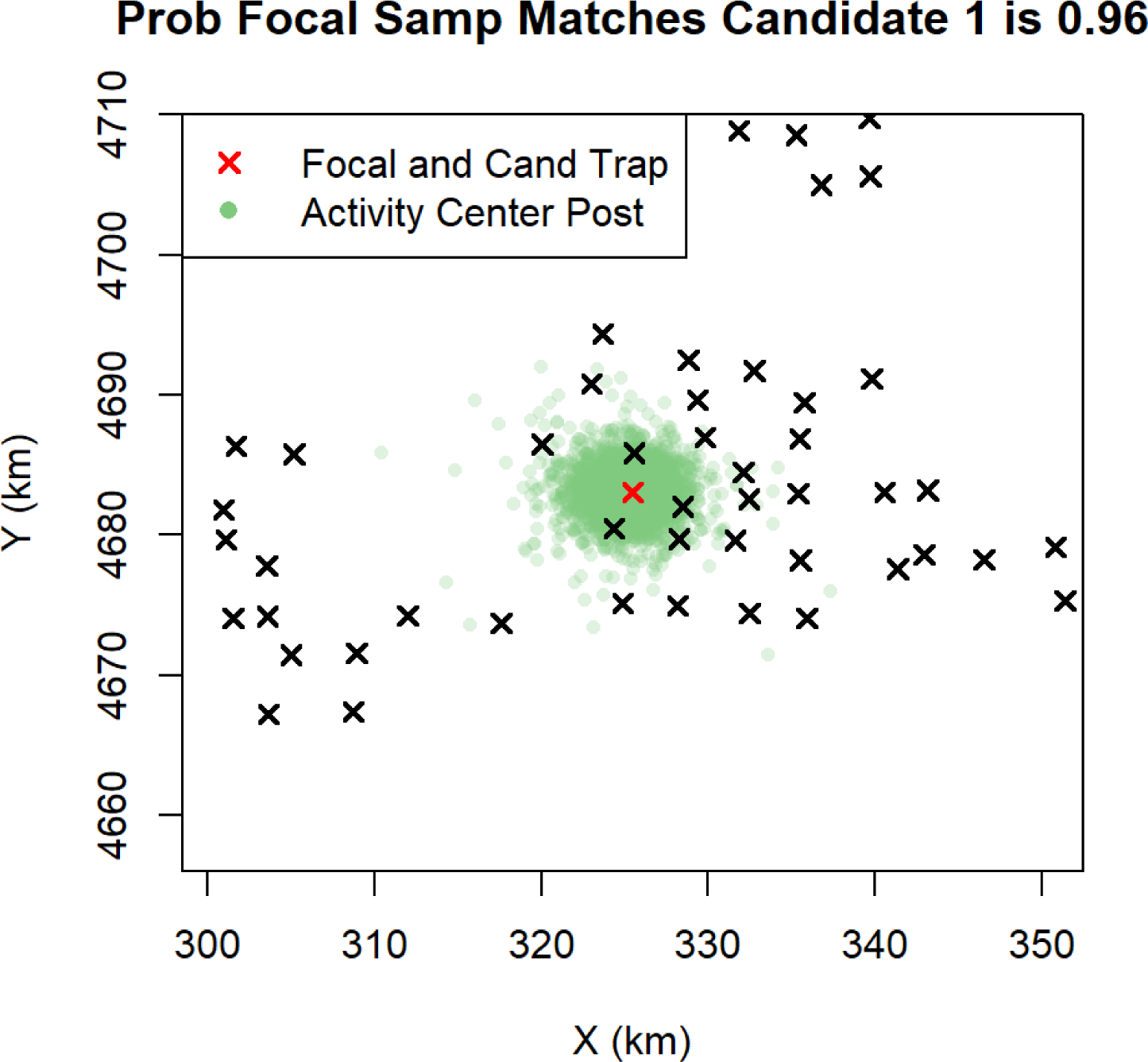

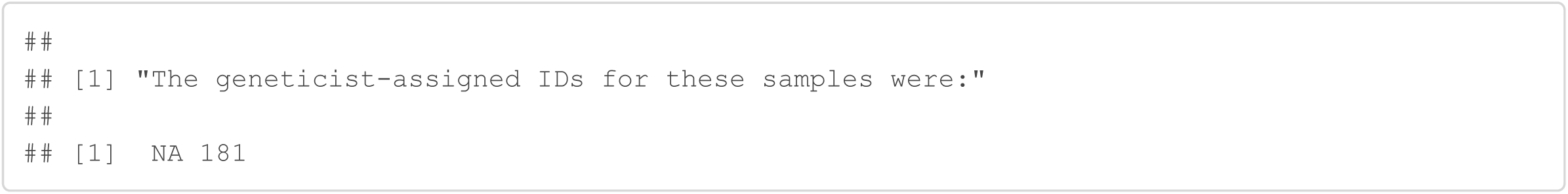

#### Sample 141

Sample 141 was originally discarded because it only amplified at 5/9 loci and it only amplified in 2 replications at 3 of these loci. The Genotype SPIM assigns it to the same individual as sample 138, captured in the same trap, with probability 0.99. For this to be a correct match, there were 2 allelic dropouts for sample 141 at locus 8. Given that sample 141 is “low quality”, this is not an unlikely event.

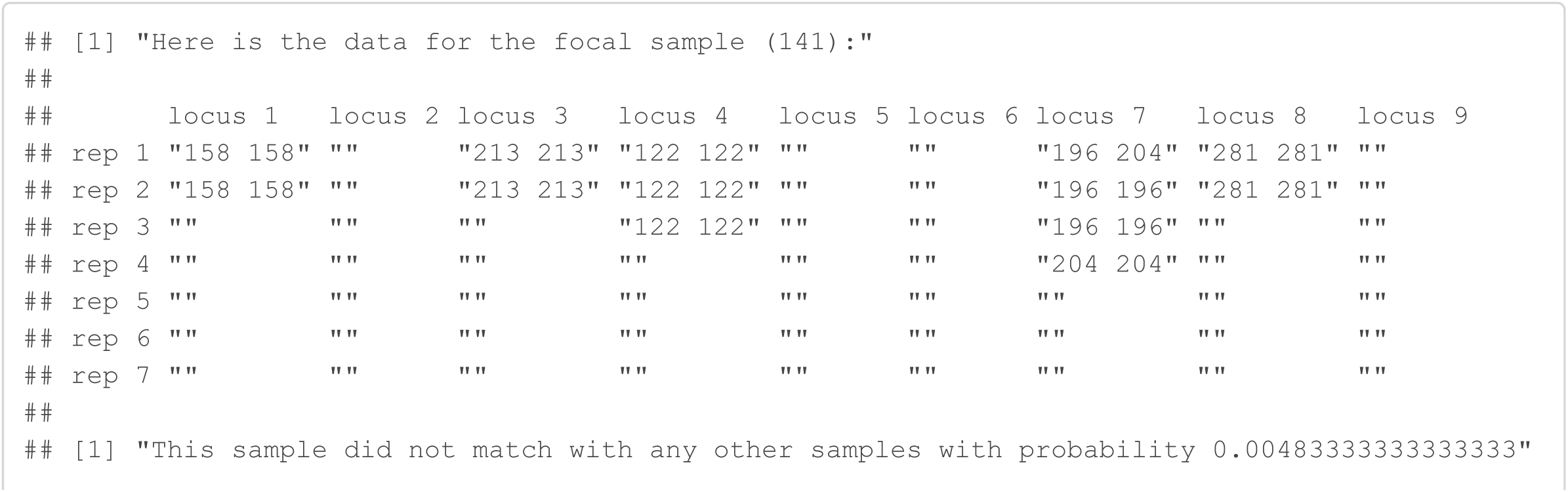

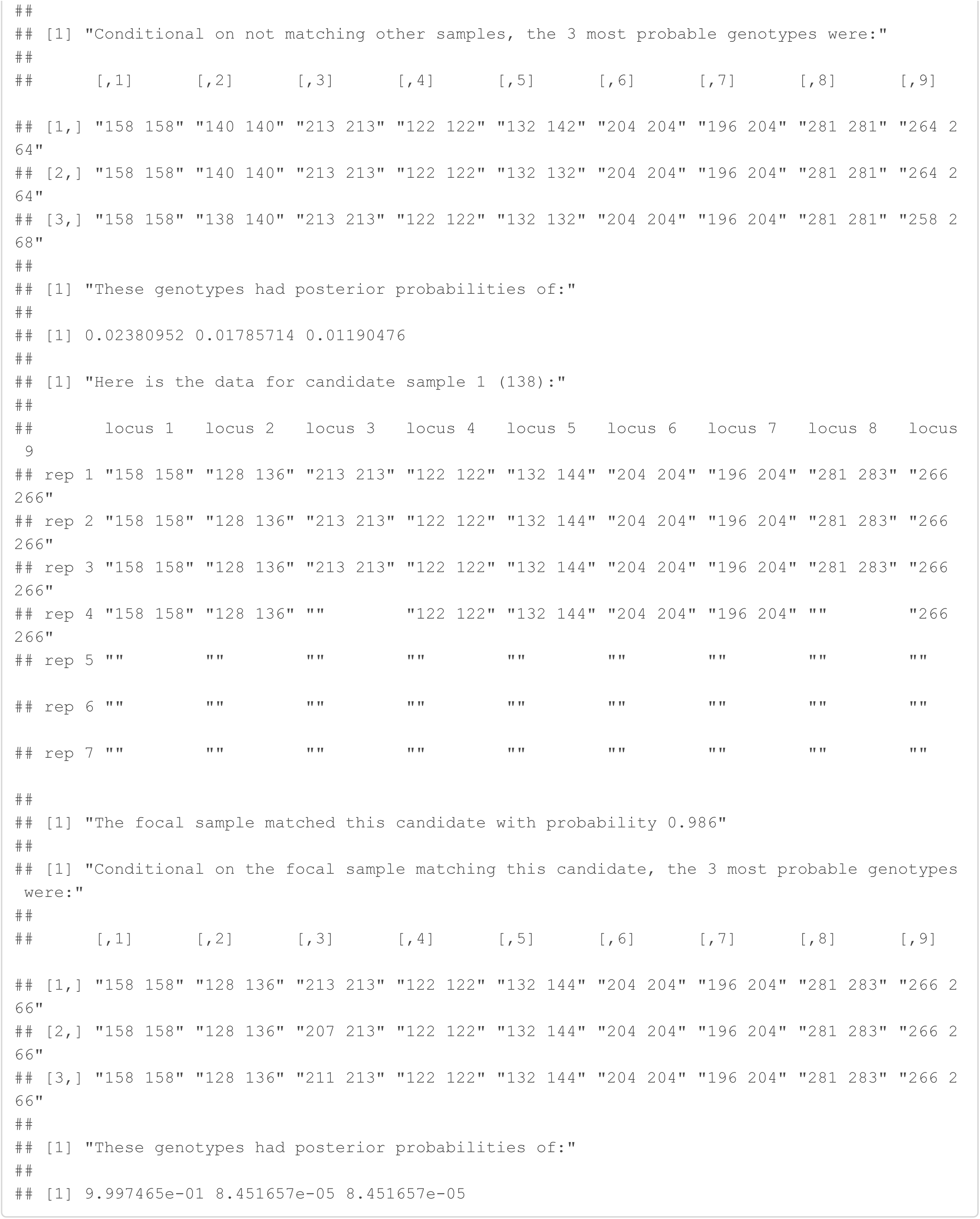

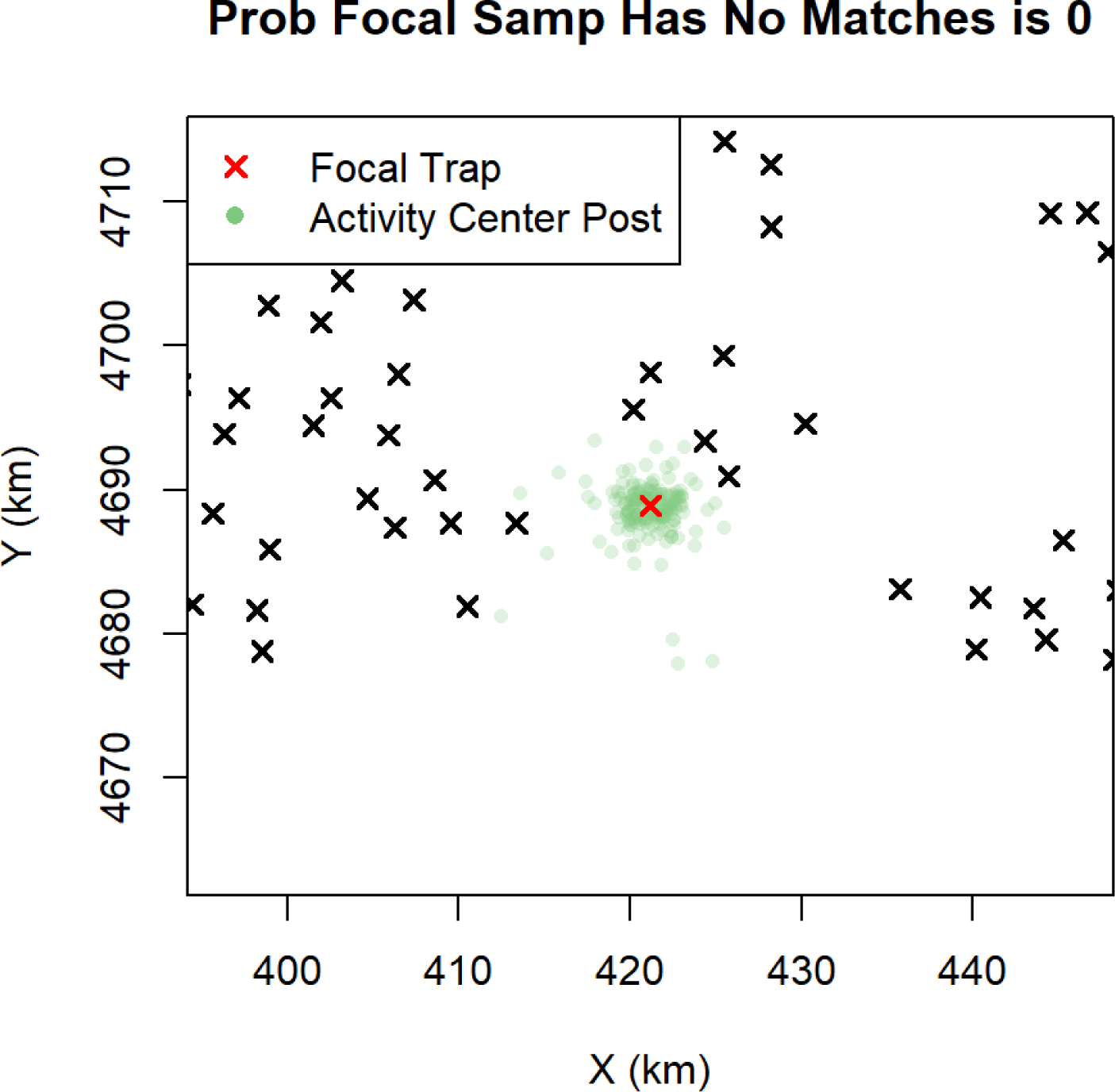

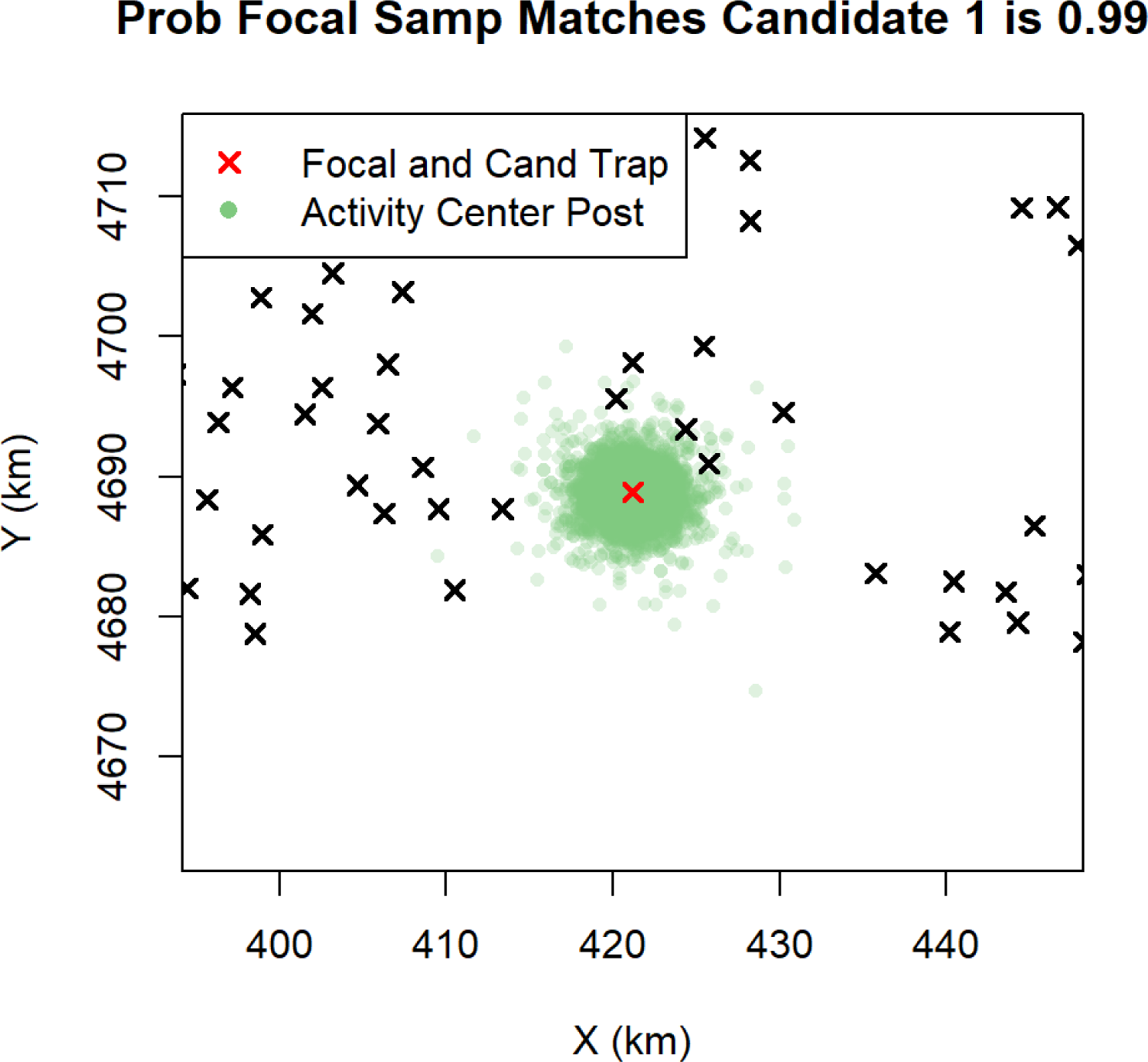

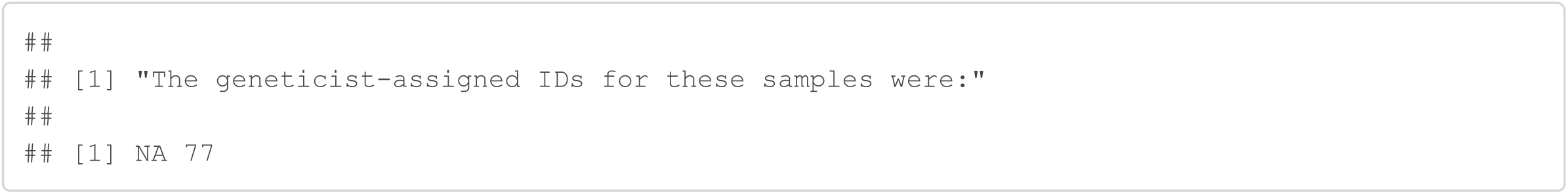

#### Sample 144

Sample 144 was originally discarded because it only amplified at 2/9 loci and it only amplified in 2 replications at each of these loci. The Genotype SPIM assigns it to the same individual as sample 143, captured in the same trap, with probability 0.98. No genotyping errors occurred for sample 144 if this is a correct match.

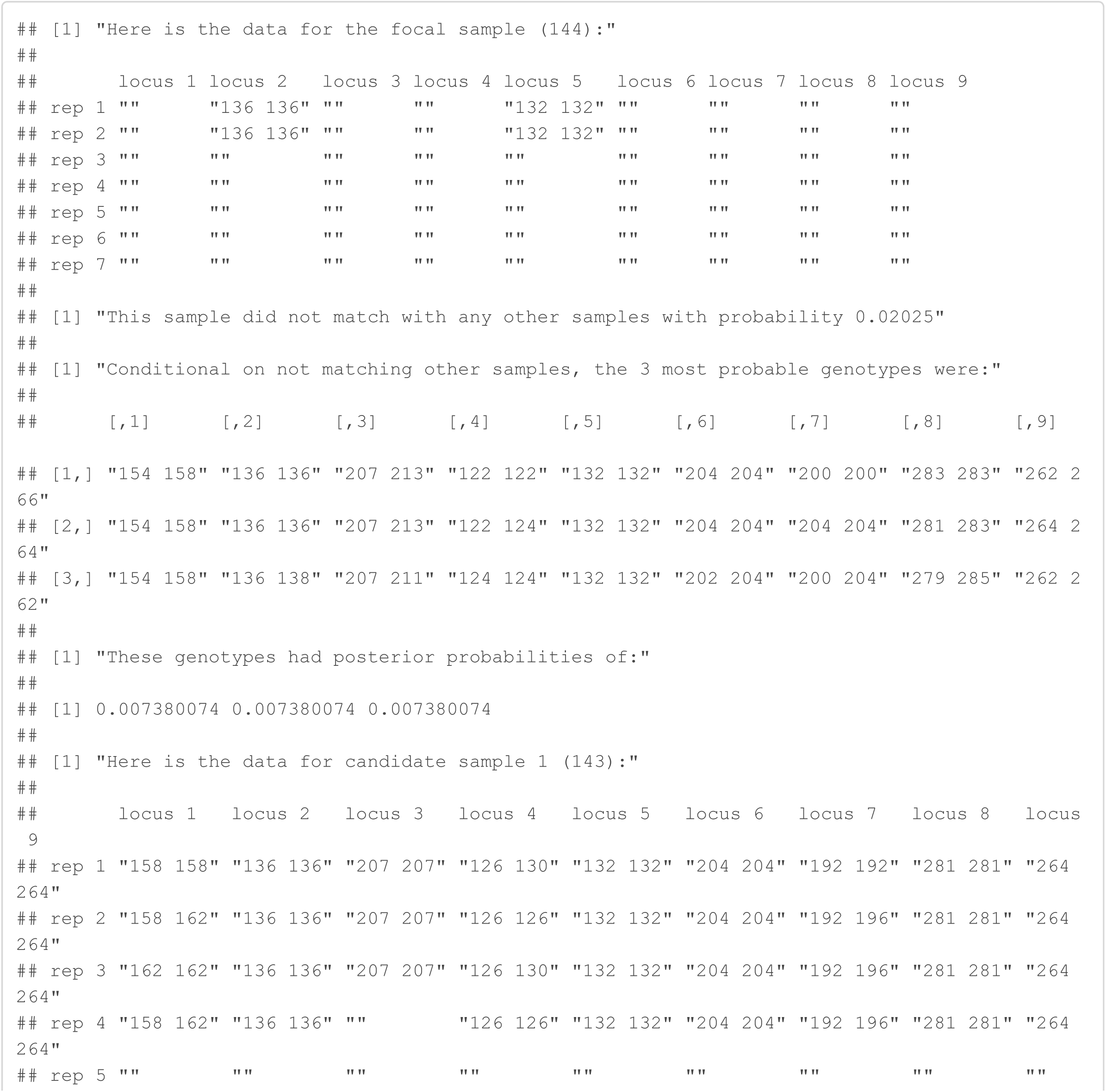

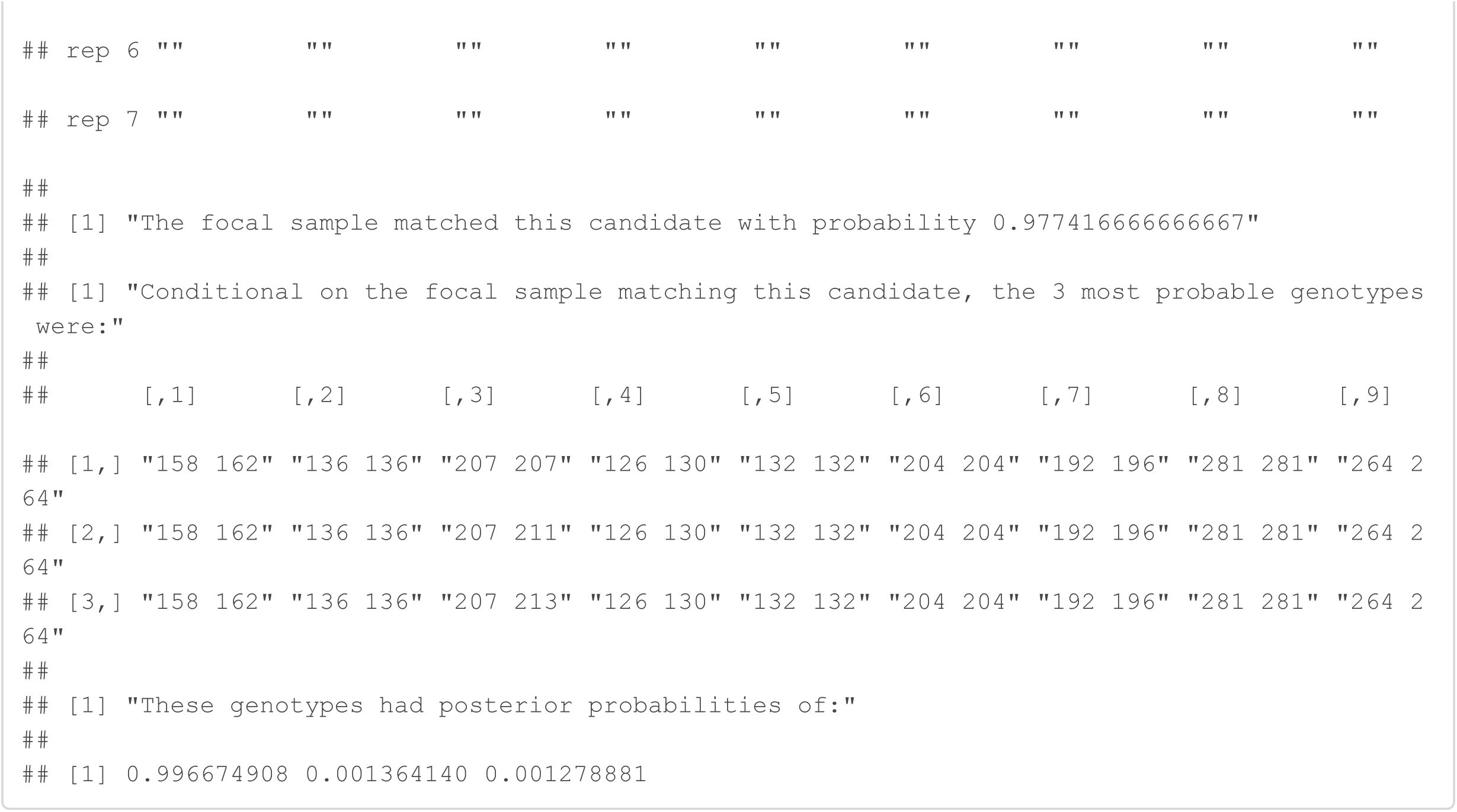

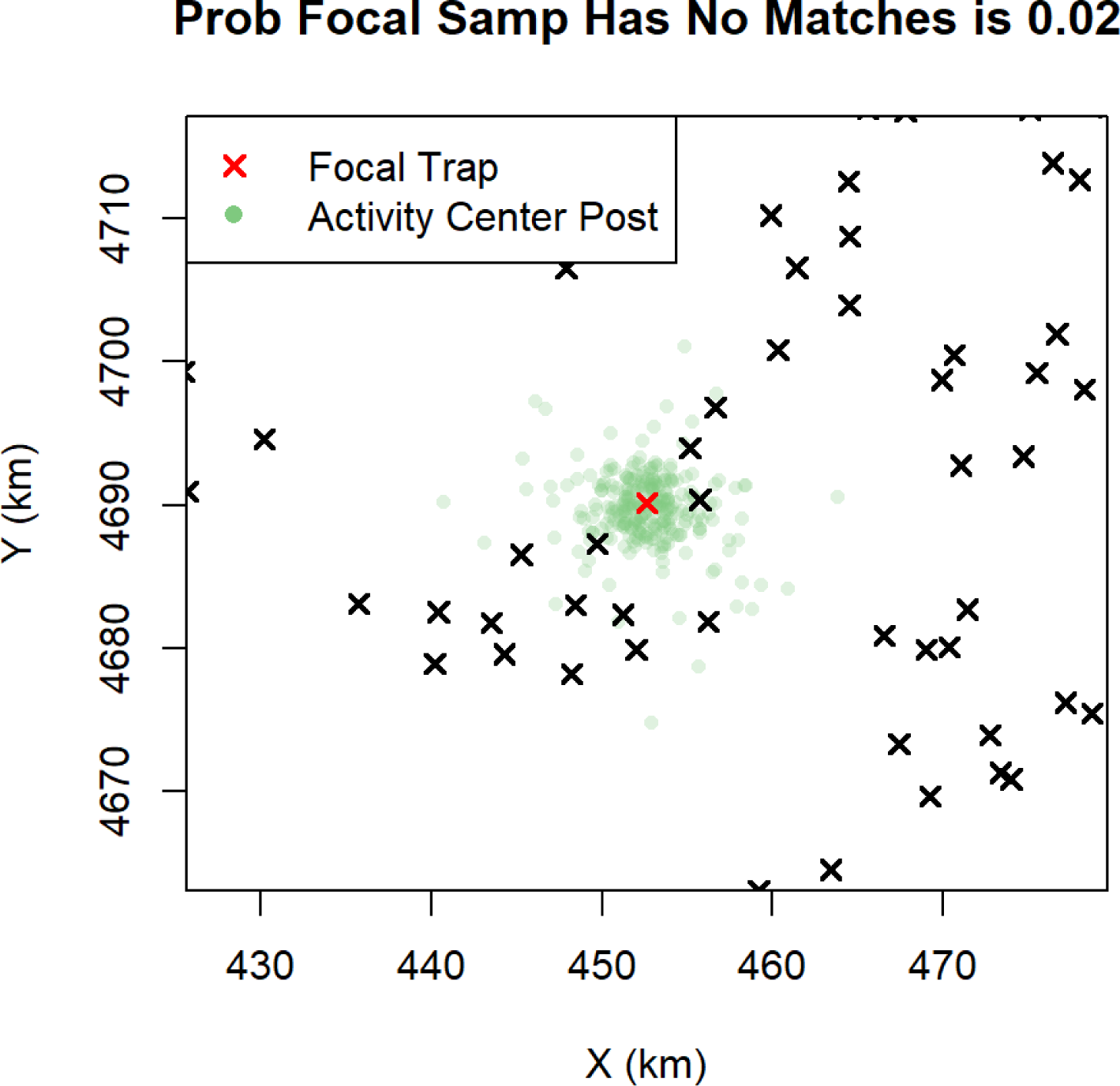

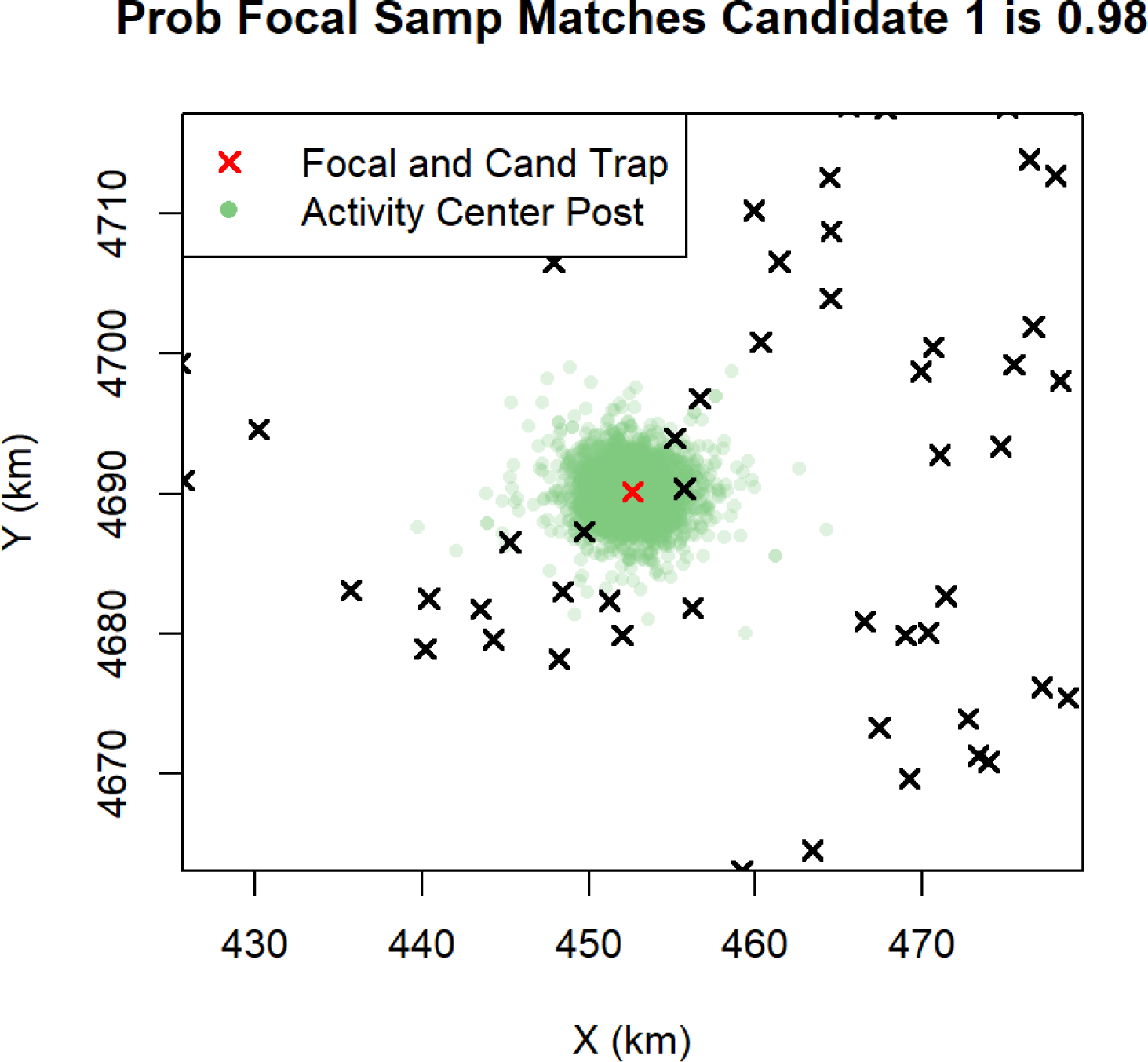

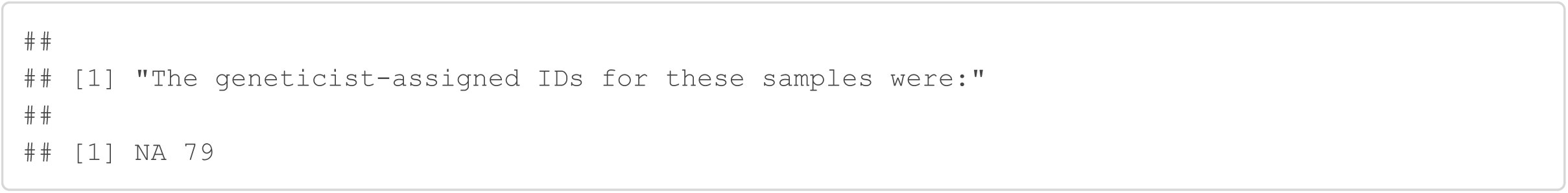

#### Sample 191

Sample 191 was originally discarded because it only amplified at 5/9 loci in 3 or more replications. Genotype SPIM matches this sample with 190 with probability 1. For this to be a correct match, there were 2 allelic dropouts for sample 191 at locus 1 and 2 allelic dropouts for sample 191 at locus 7.

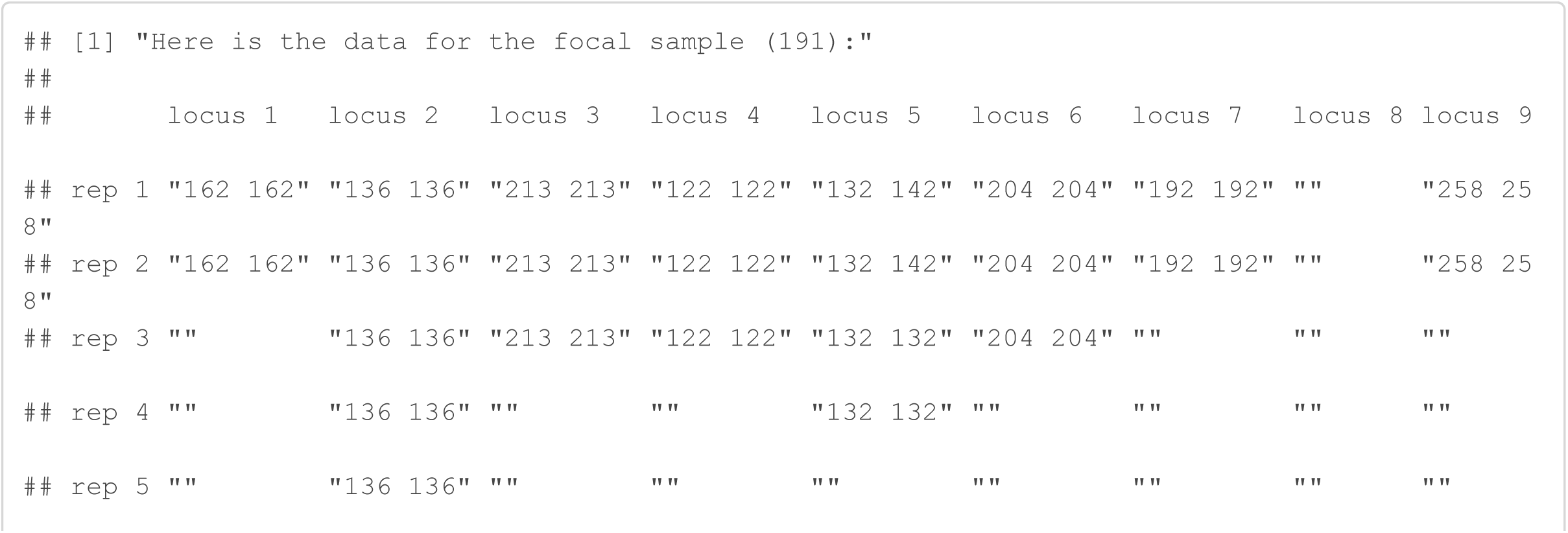

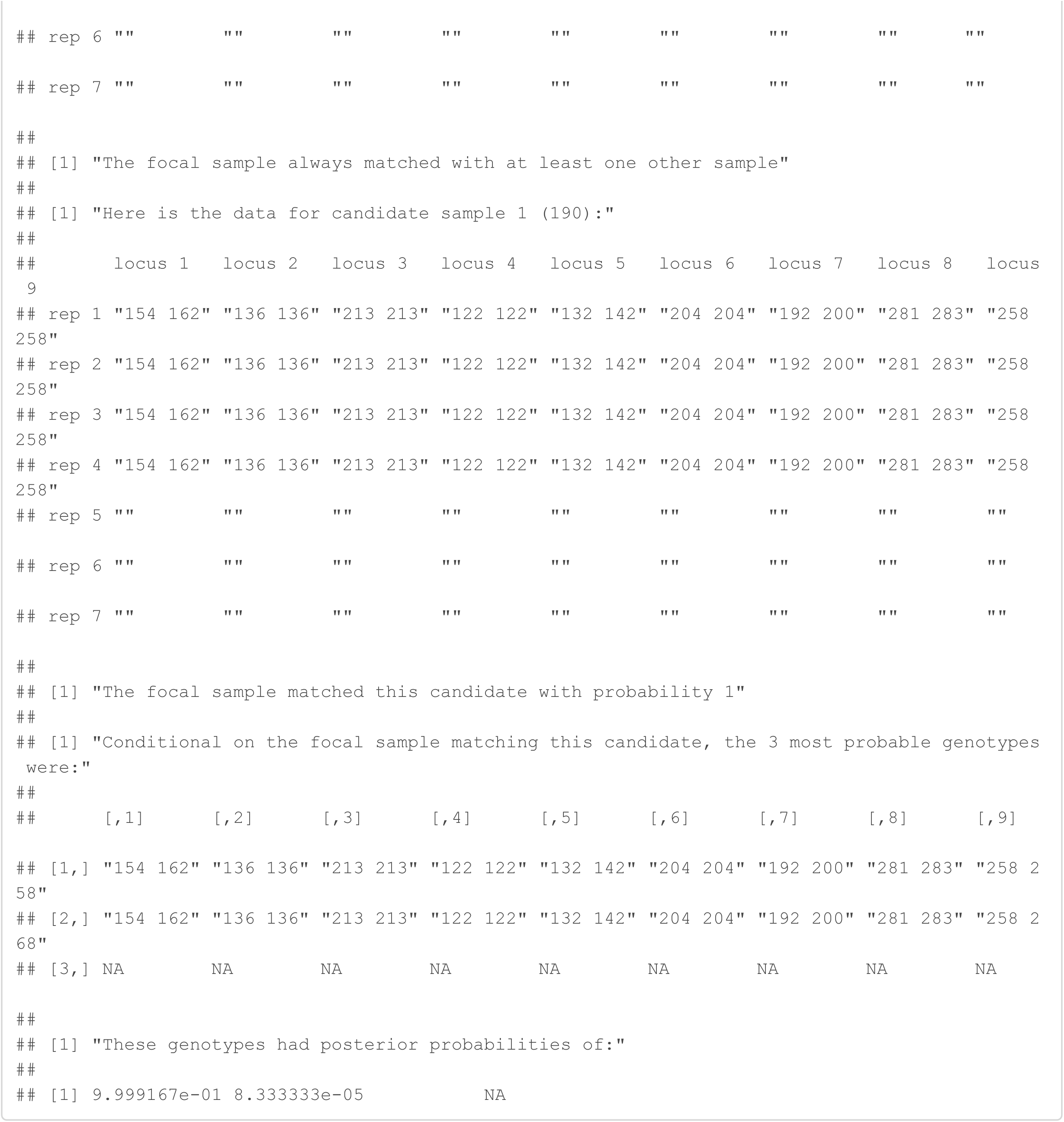

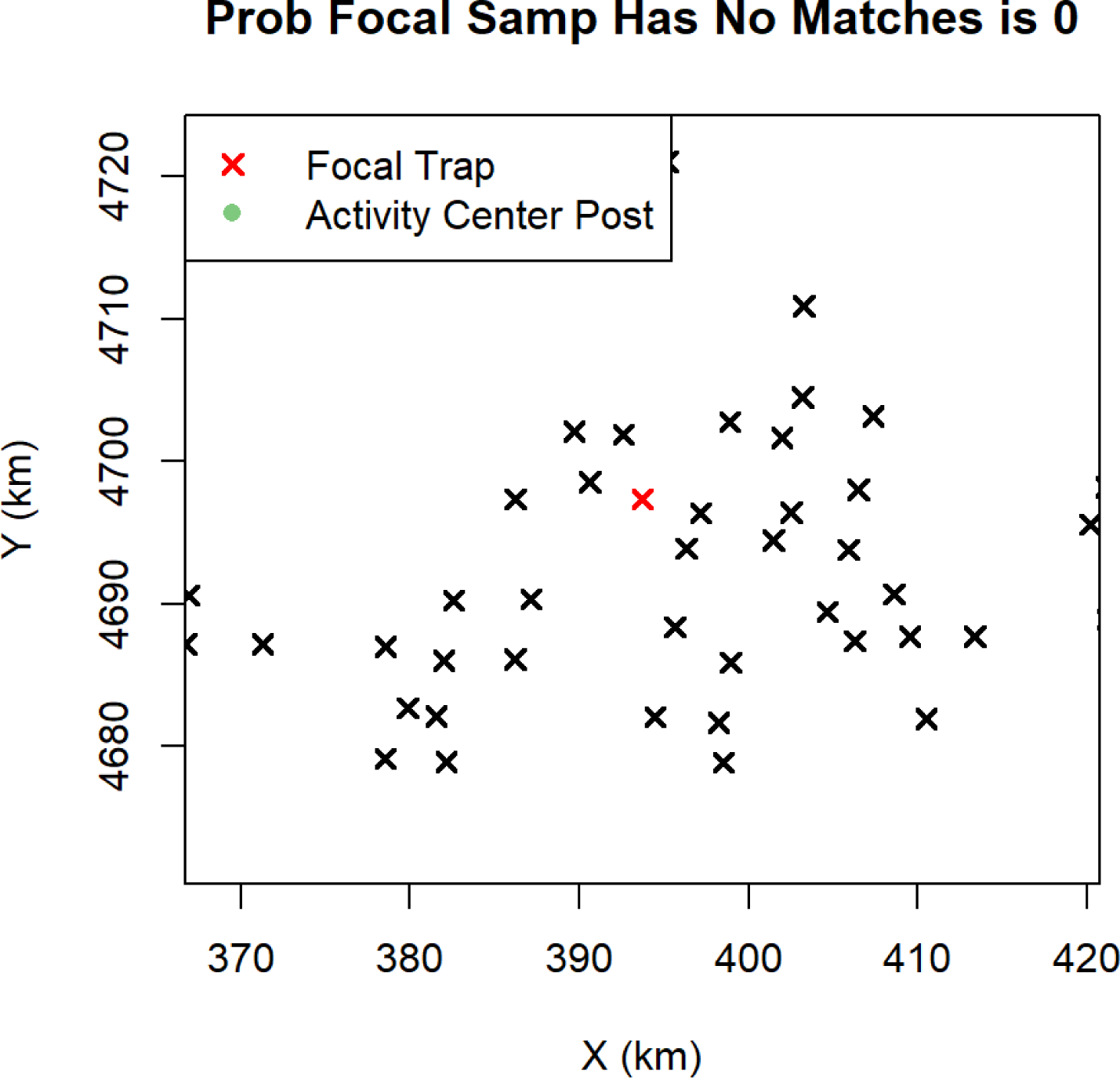

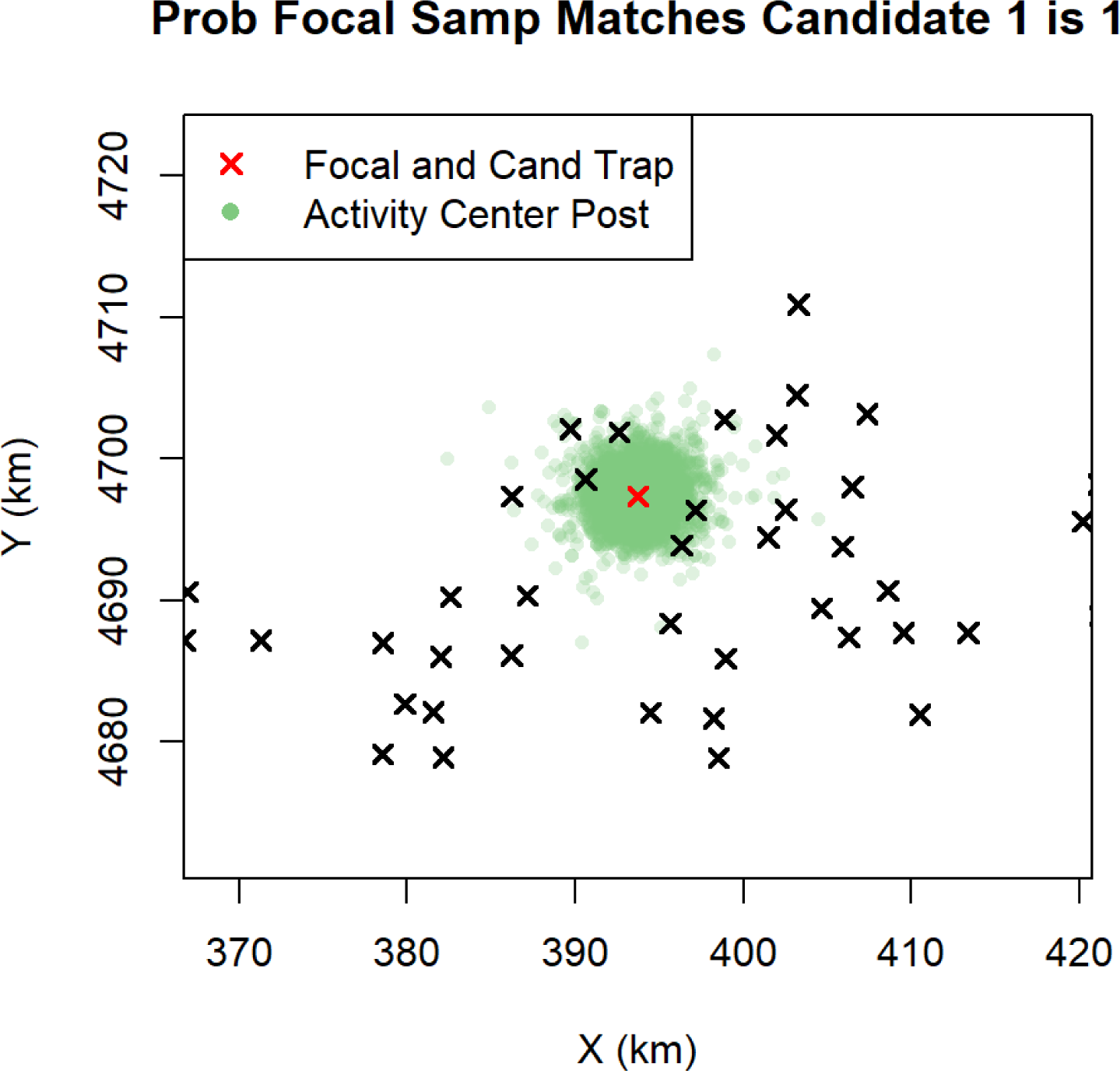

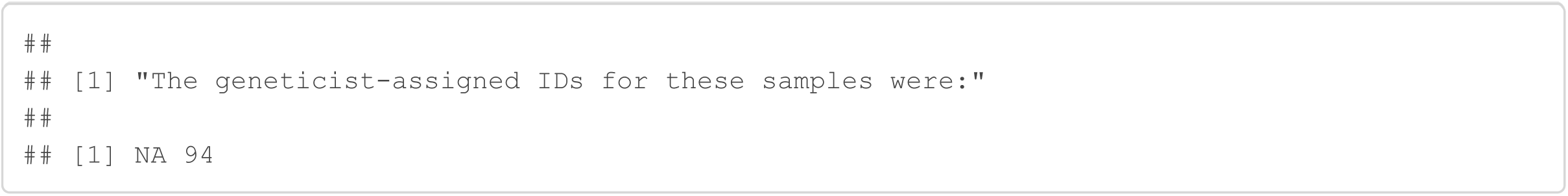

#### Sample 300

Sample 300 was originally discarded because it only amplified at 3/9 loci and only amplified 3 or more times at 1 locus. The Genotype SPIM matches this sample to samples 297, 298, and 299, all captured at the same trap, with probability 0.97. If this is the correct match, there were 2 allelic dropouts for sample 300 at locus 3.

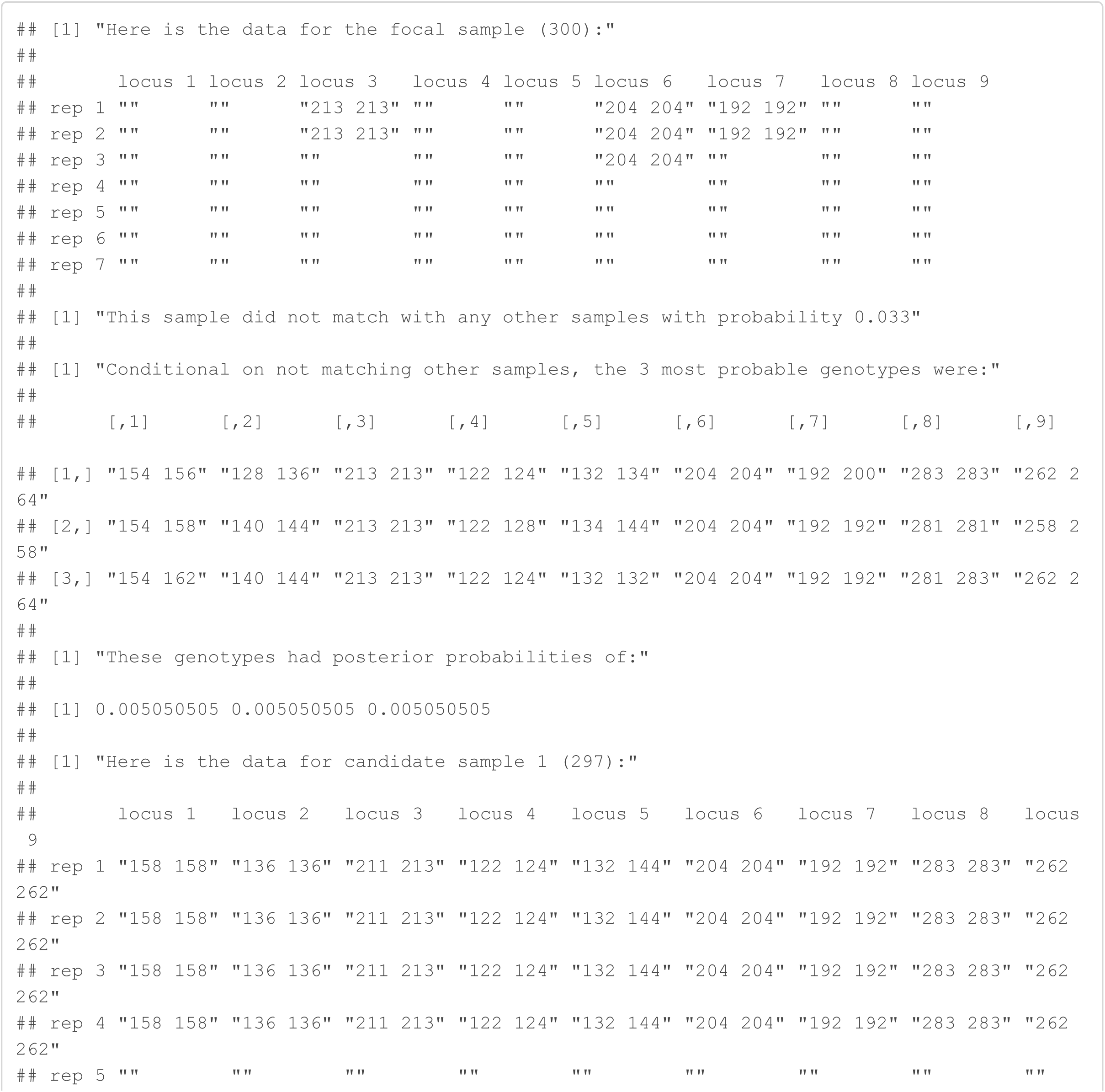

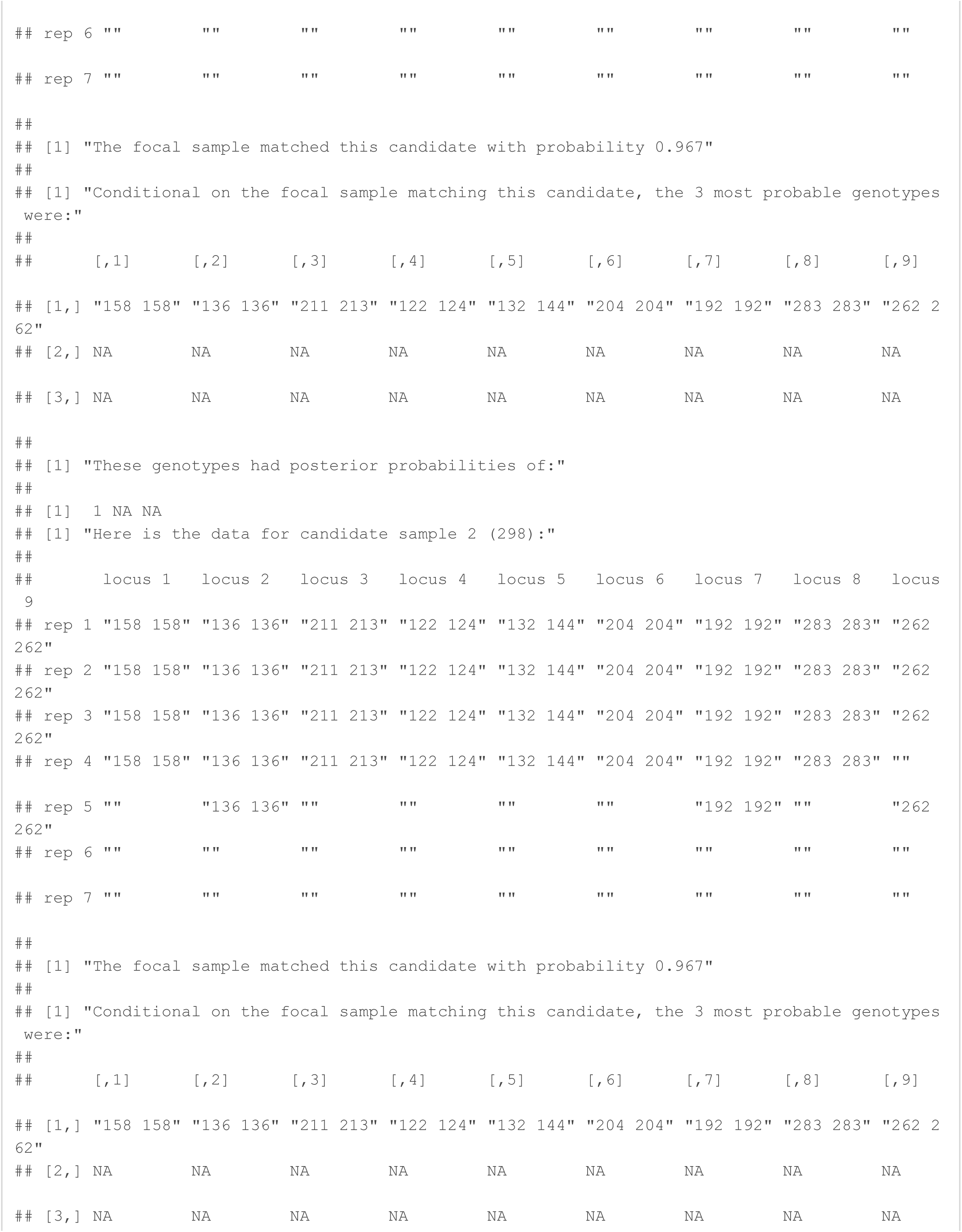

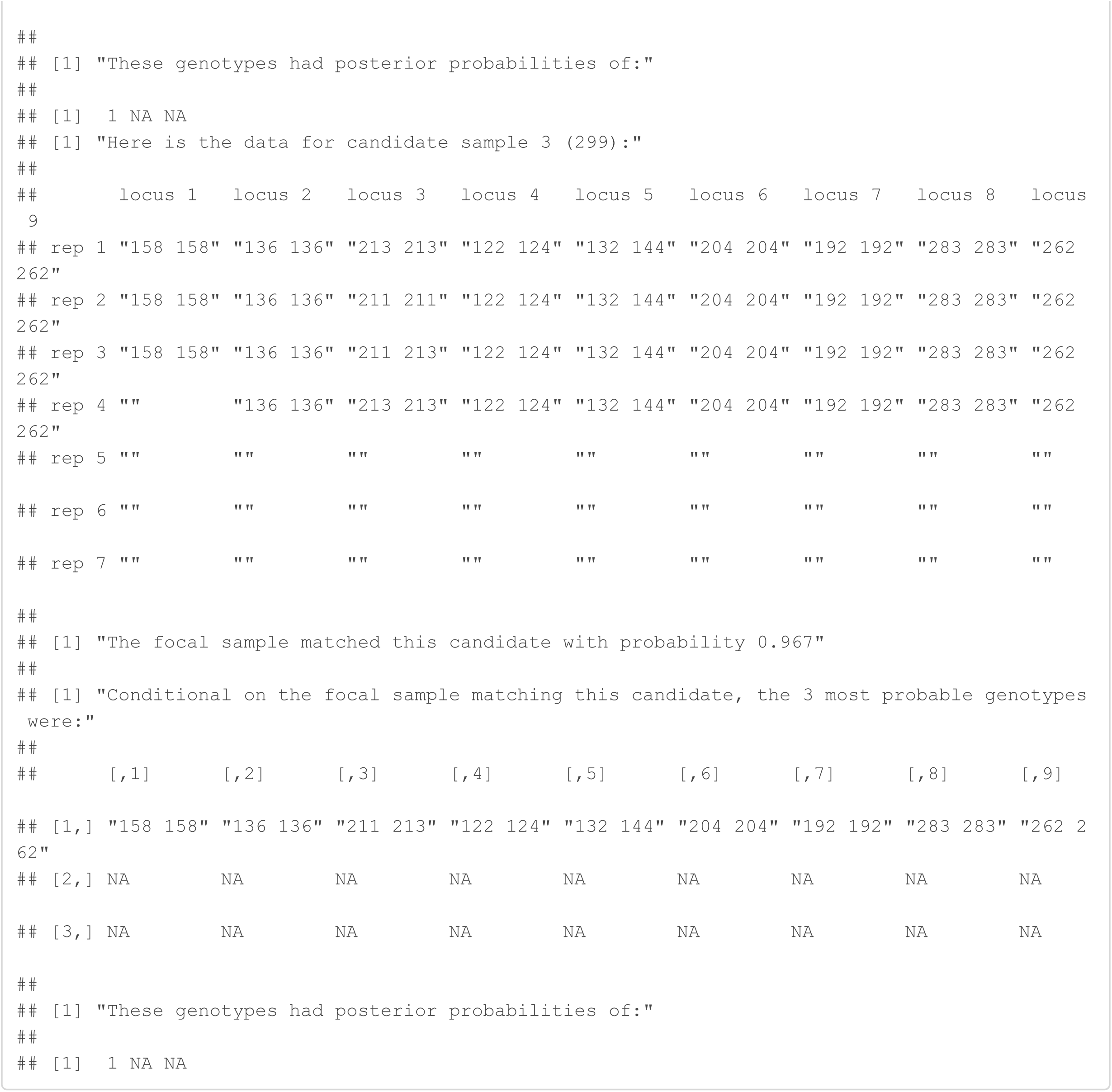

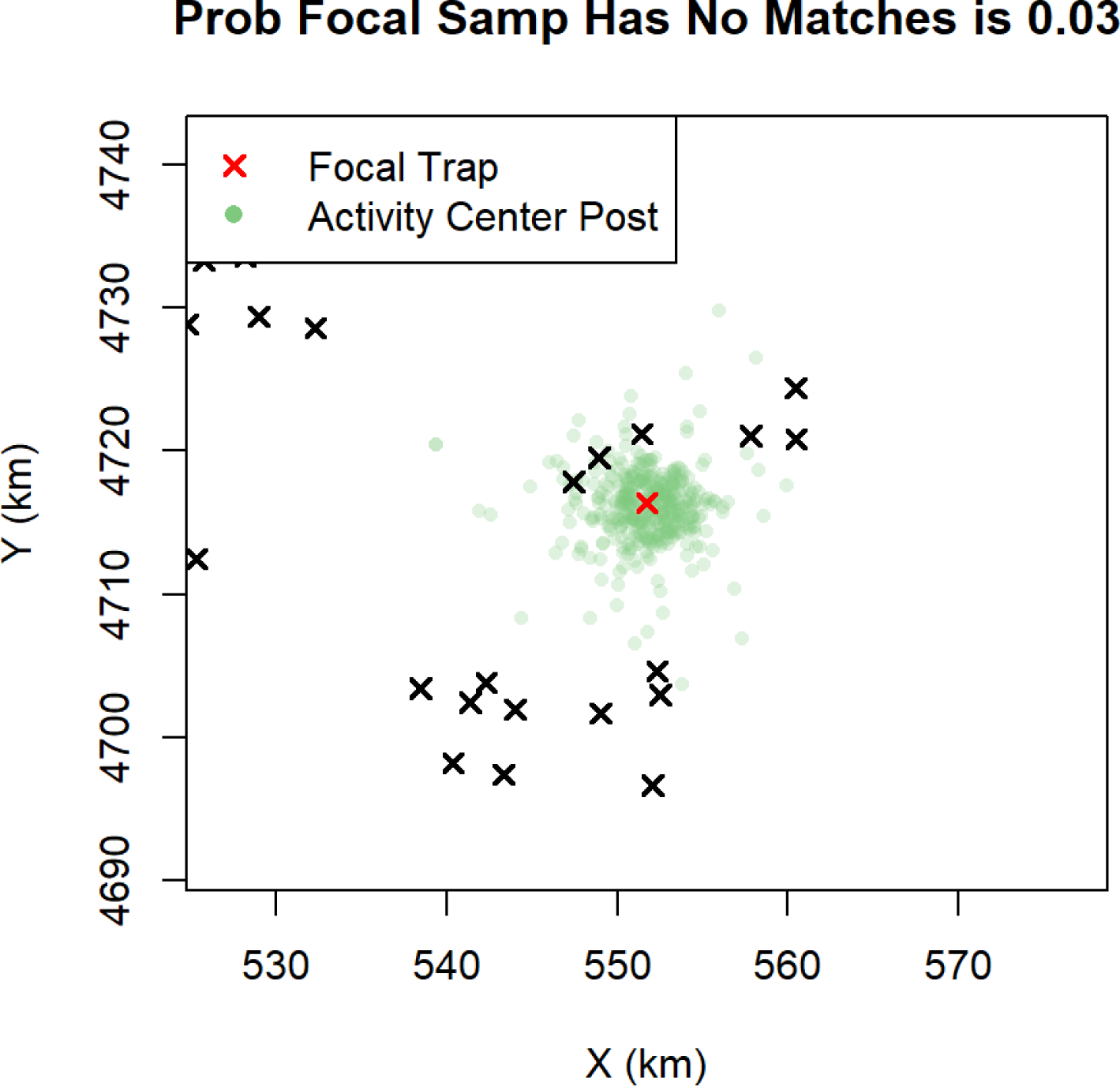

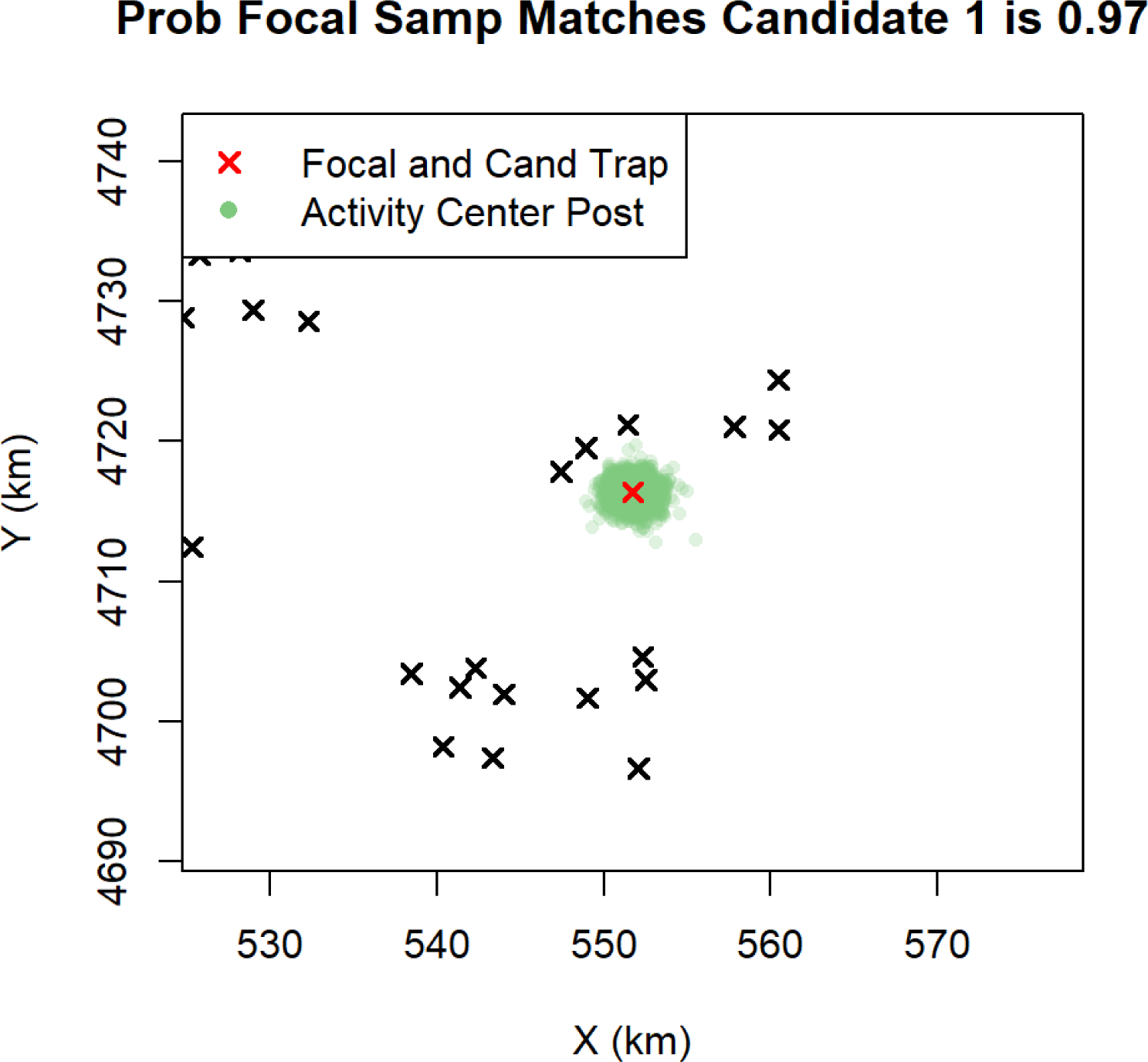

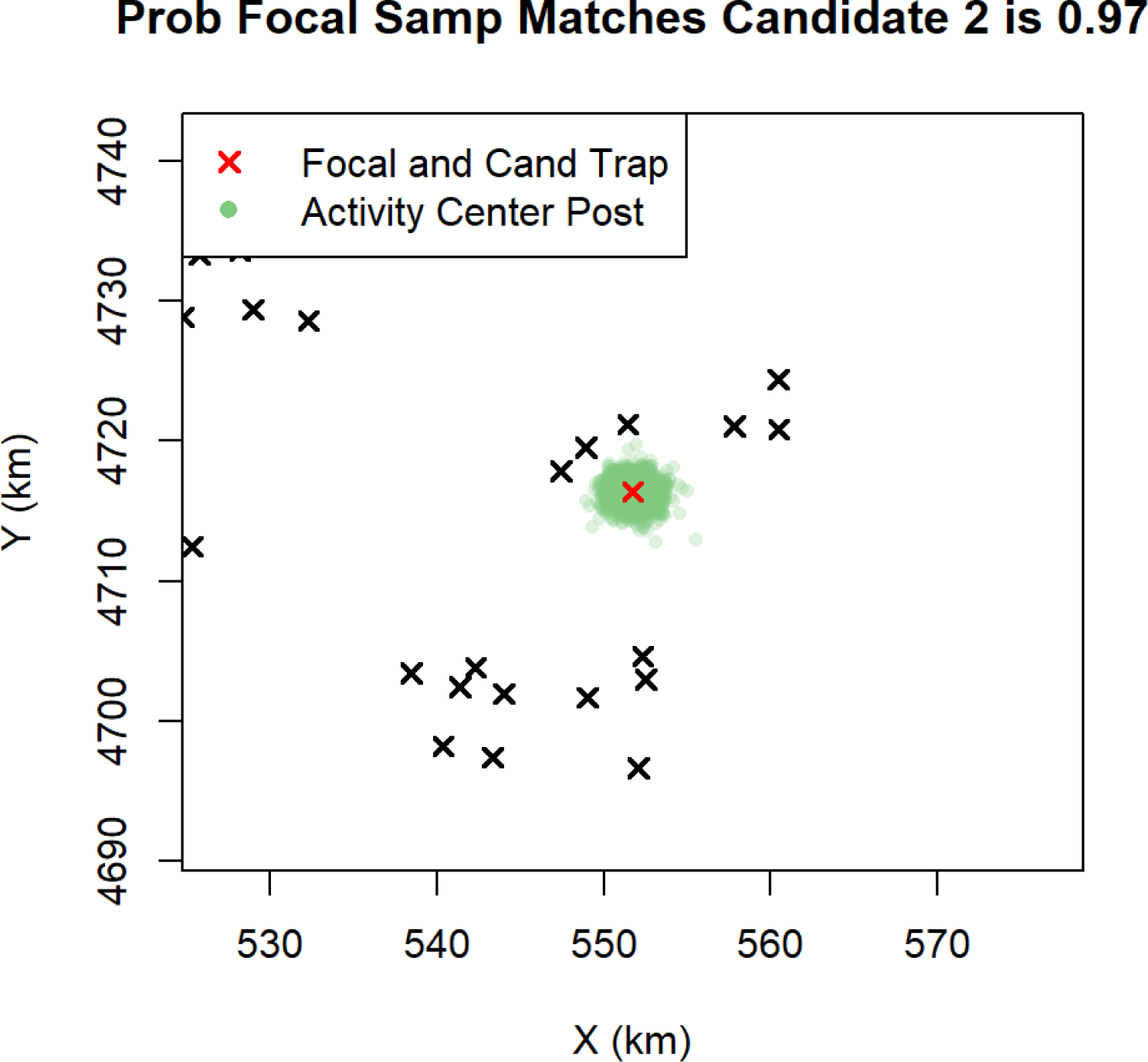

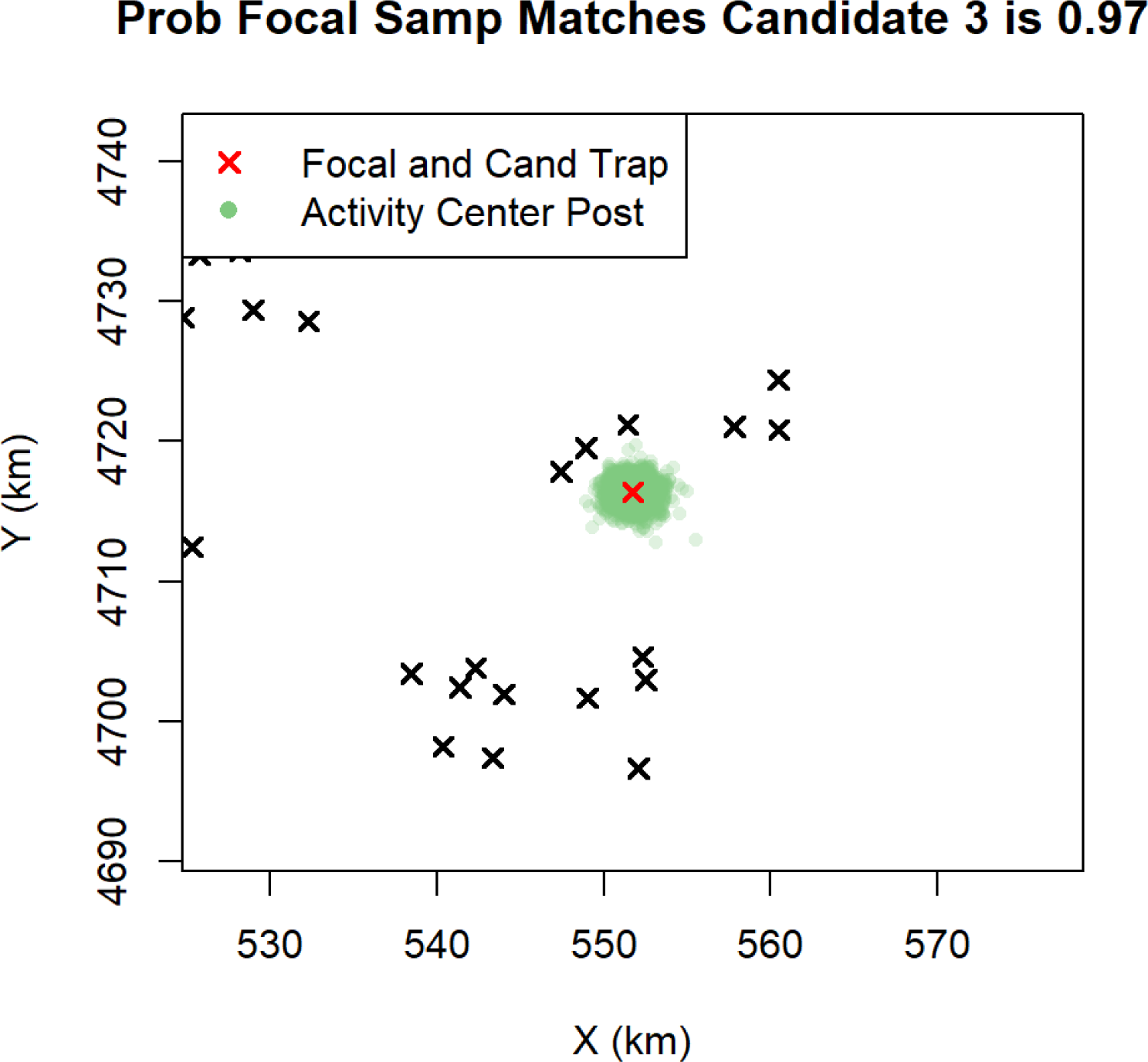

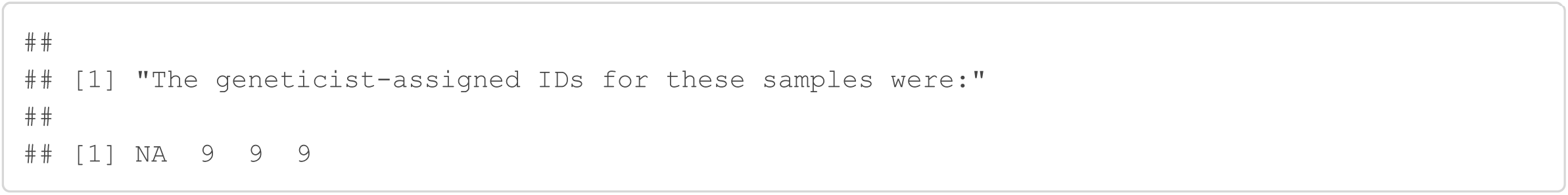

#### Sample 304

Sample 304 was originally discarded because it only amplified at 4/9 loci and only amplified 3 or more times at 2 loci. The Genotype SPIM matches this sample to sample 303, captured at the same trap, with probability 1. If this is the correct match, there were 3 allelic dropouts for sample 304 at locus 2. Further, there was 1 allelic dropout and 2 false allele events at locus 9 for sample 304.

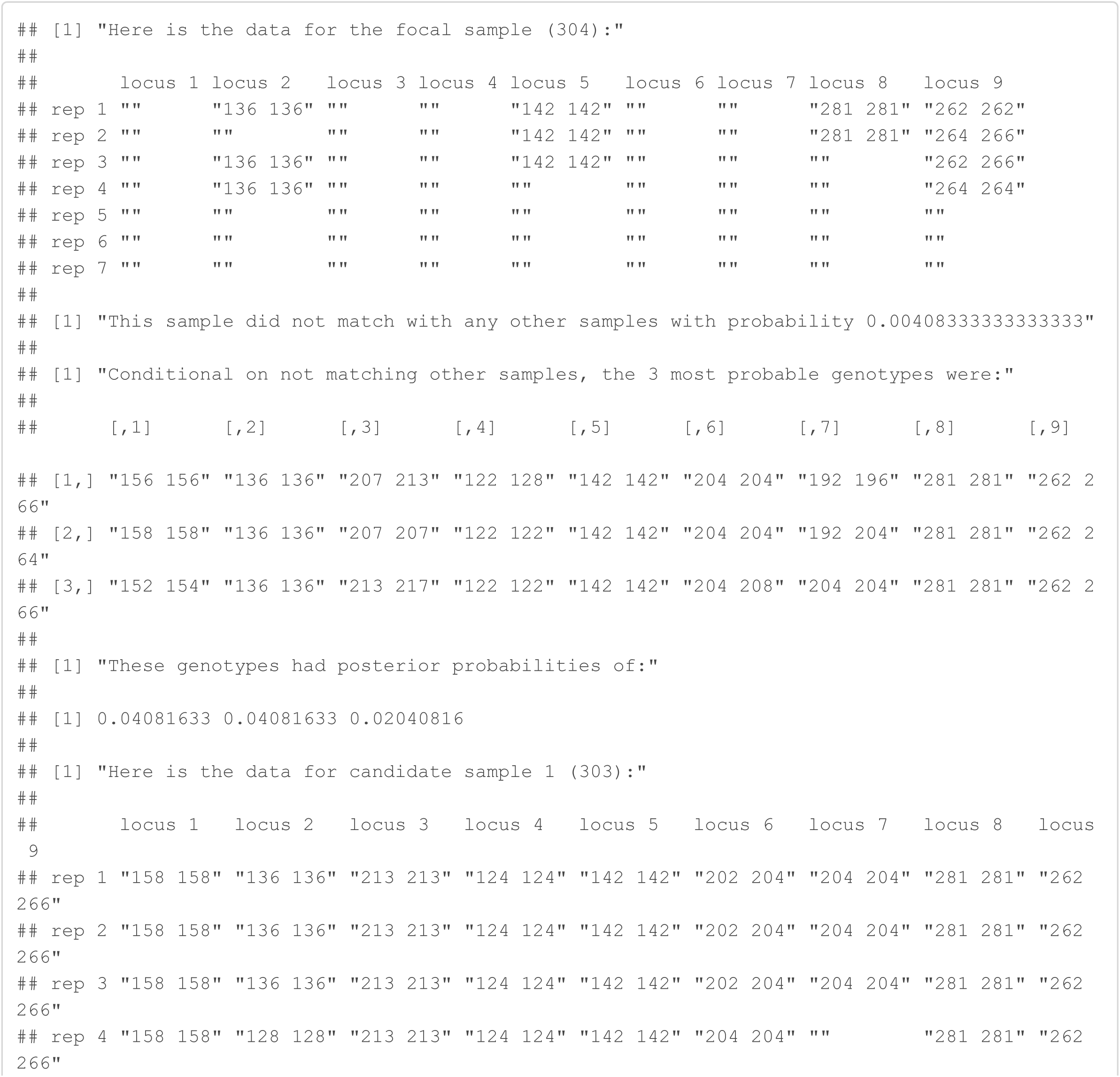

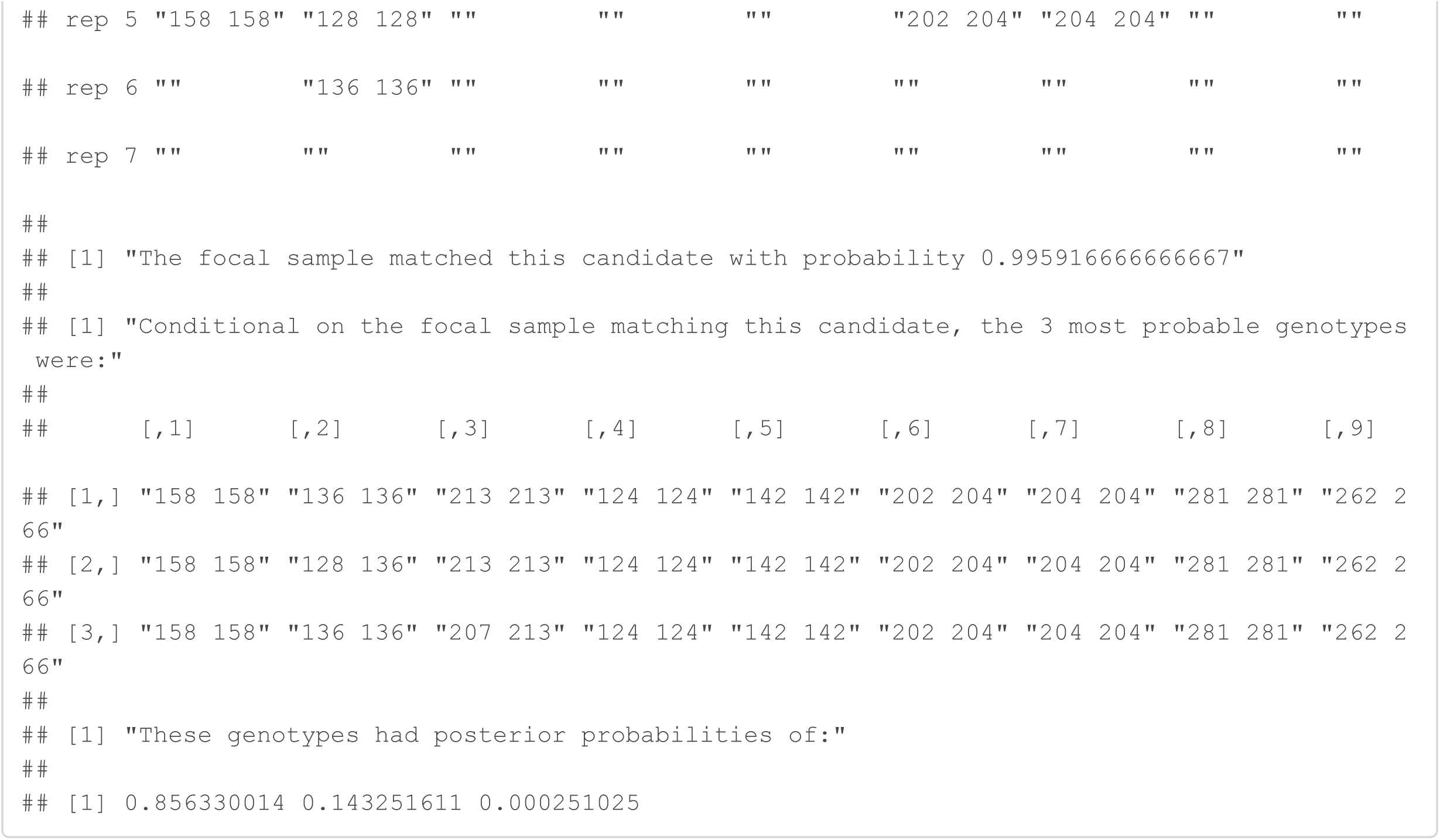

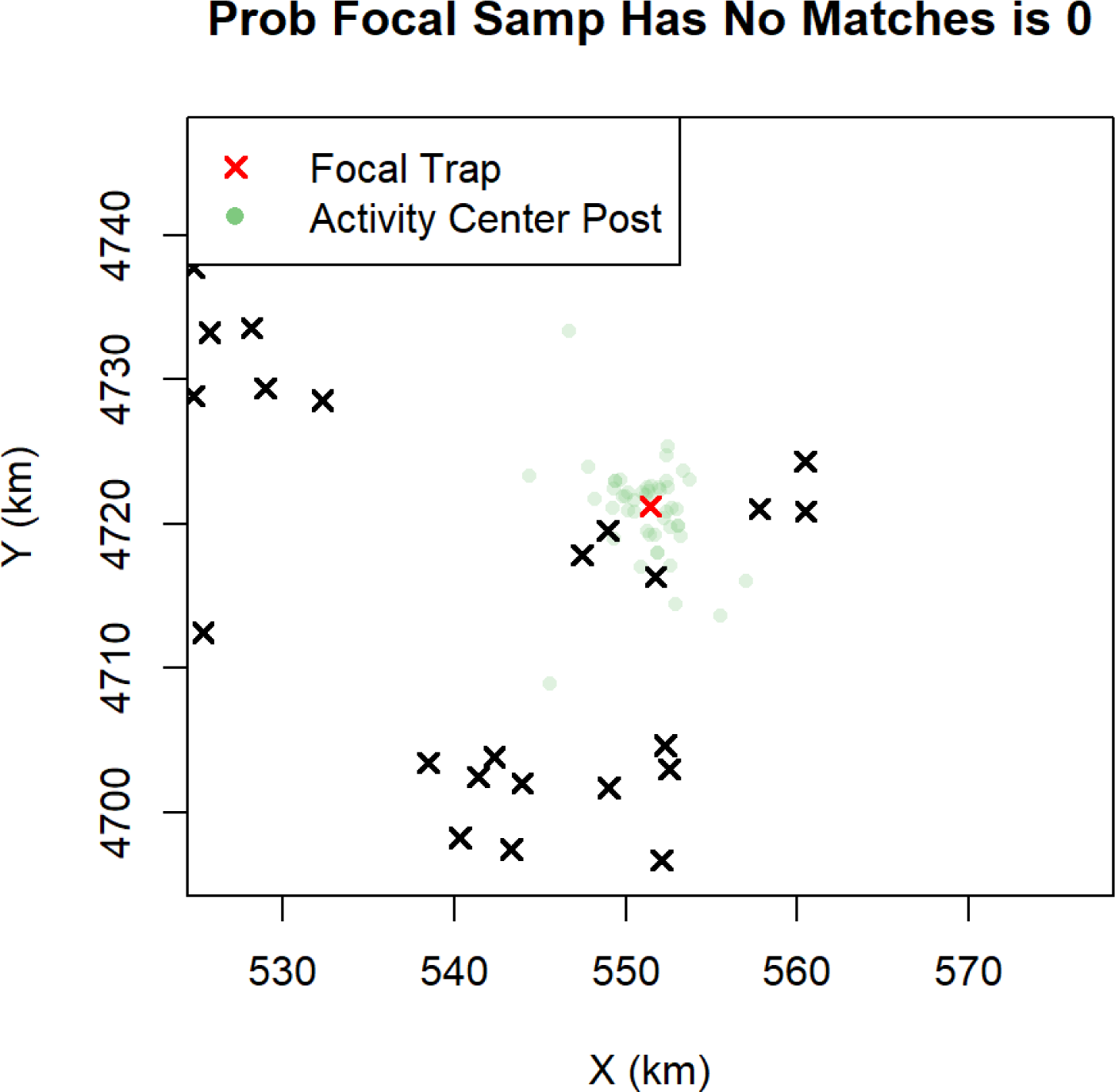

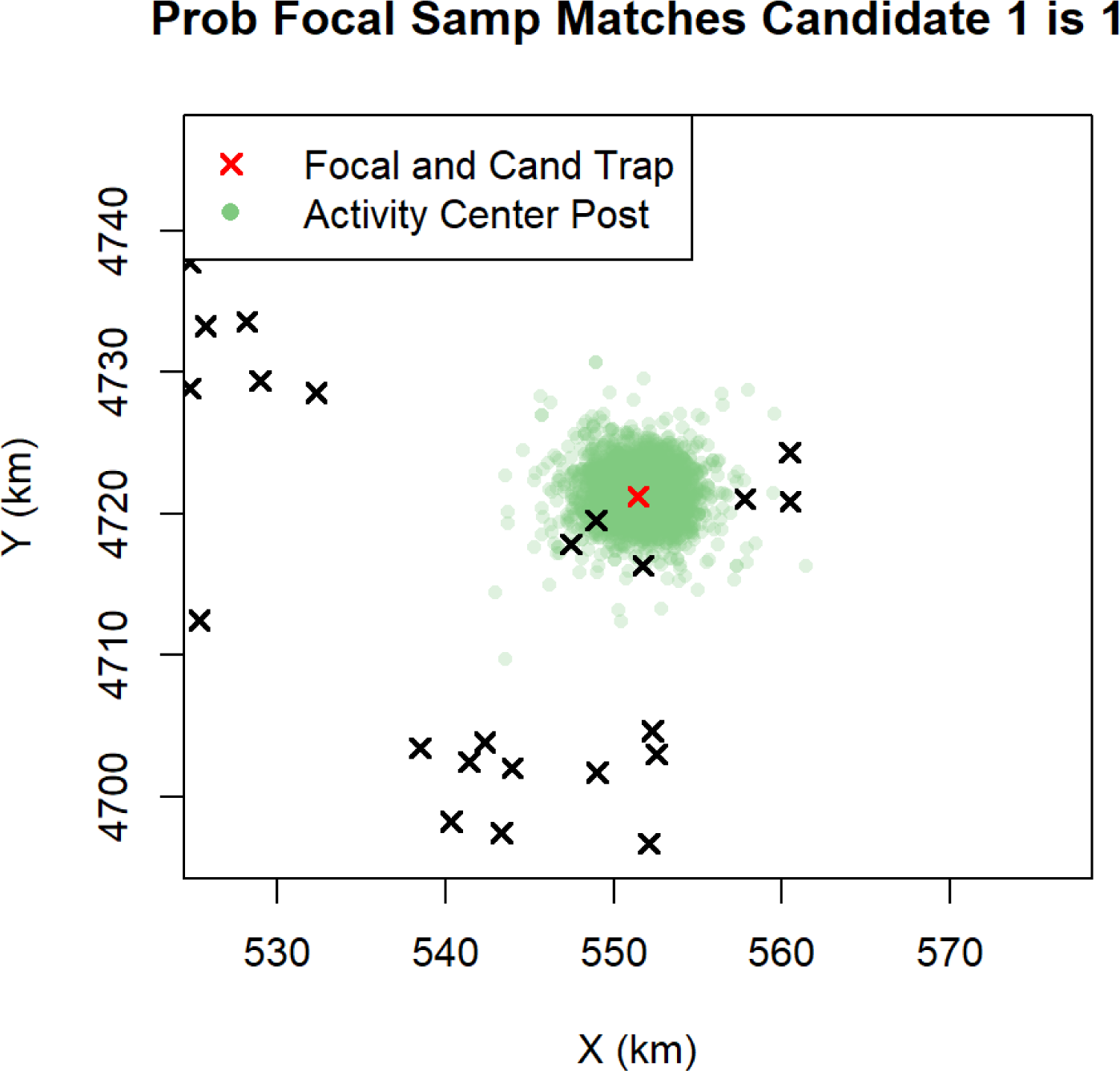

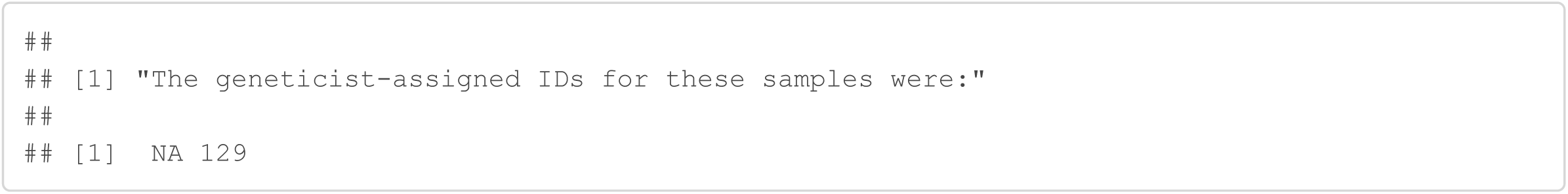

#### Sample 416

Sample 416 was originally discarded because it only amplified at 5/9 loci and only amplified 3 or more times at 1 loci. The Genotype SPIM matches this sample to samples 418 and 419, captured at a nearby trap, with probability 0.99. If this is the correct match, there were 2 allelic dropouts for sample 416 at locus 6, and 2 allelic dropouts for sample 416 at locus 9.

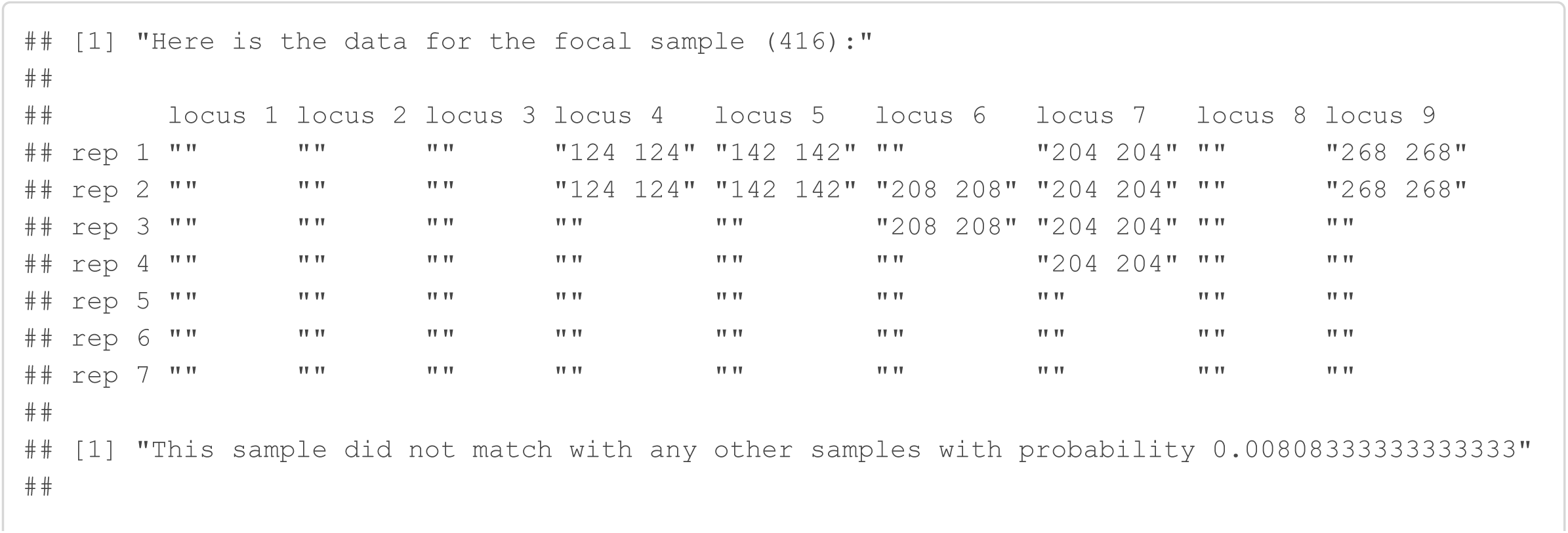

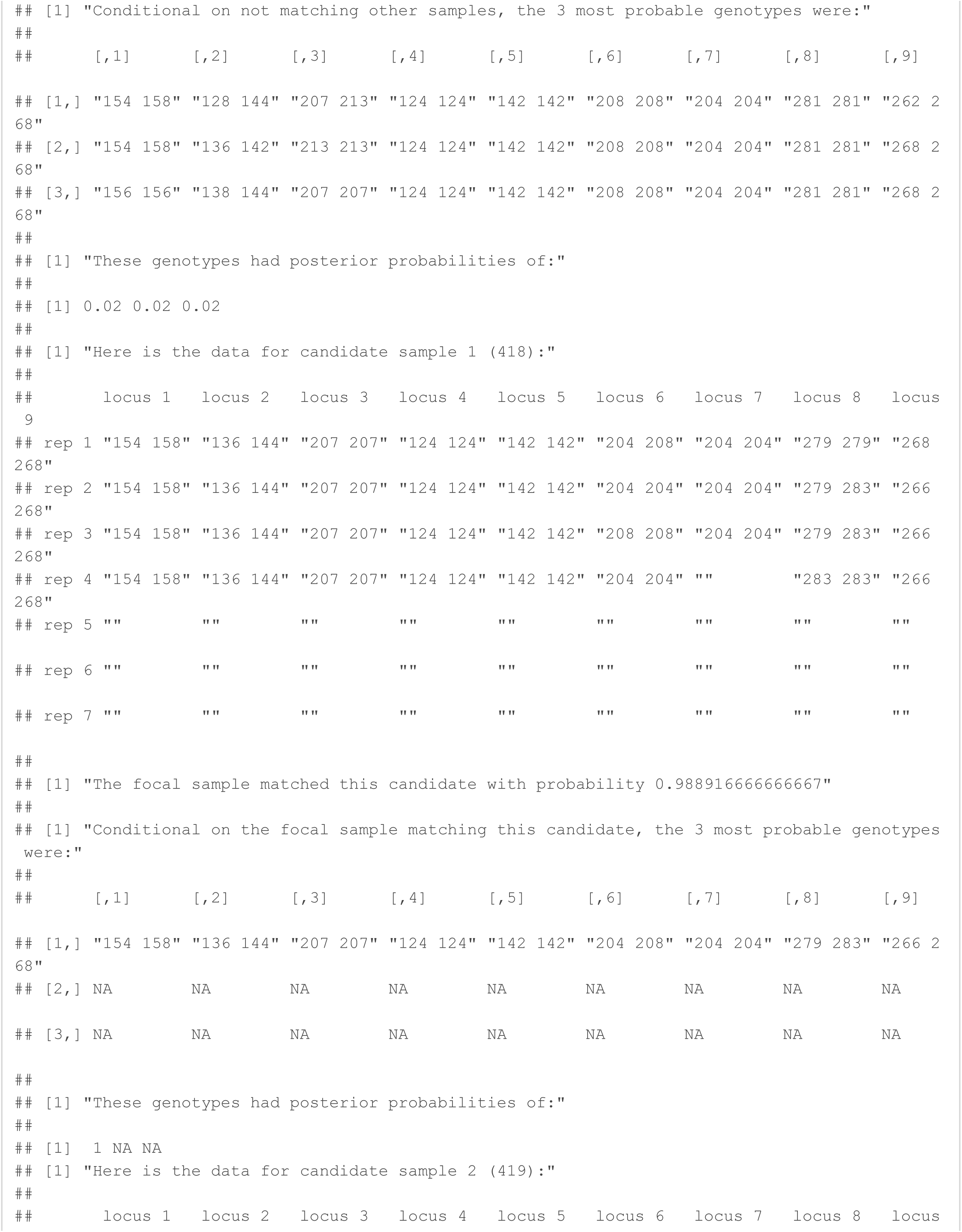

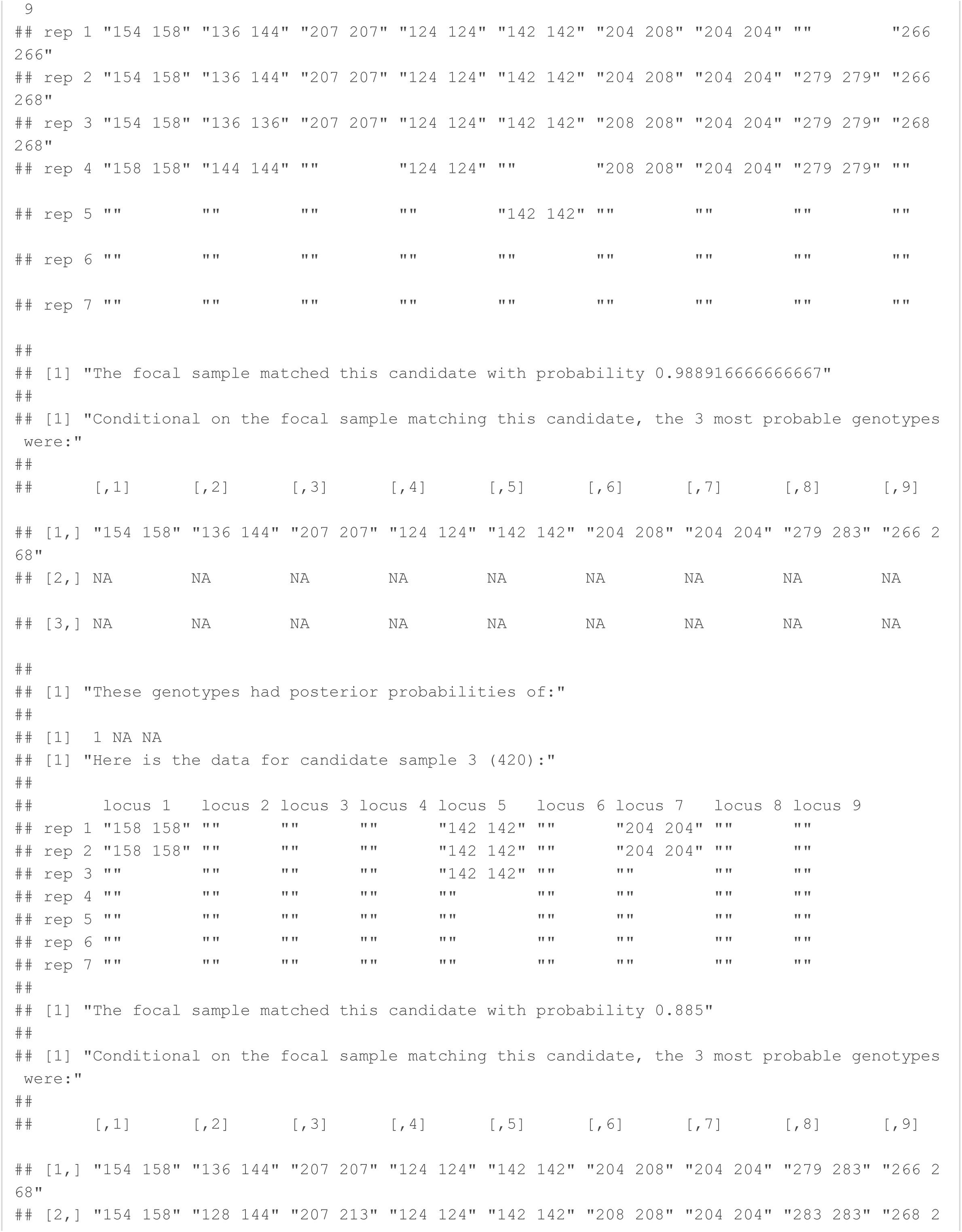

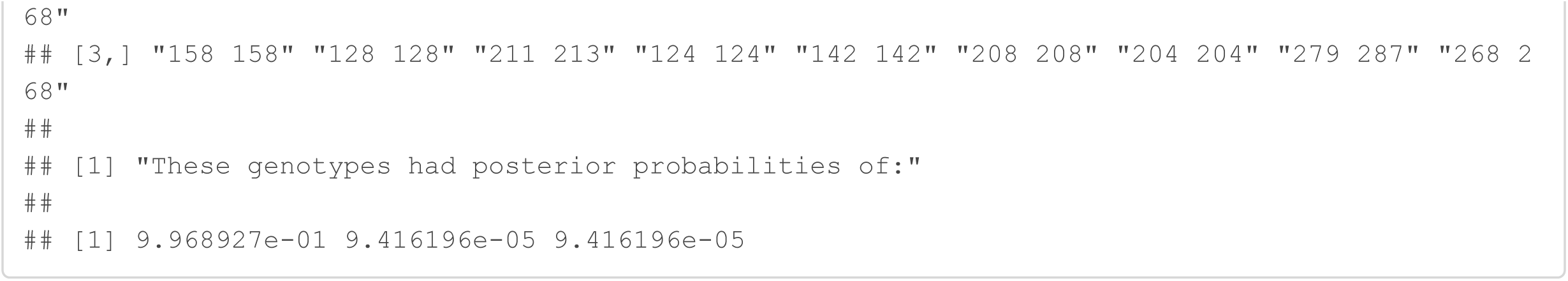

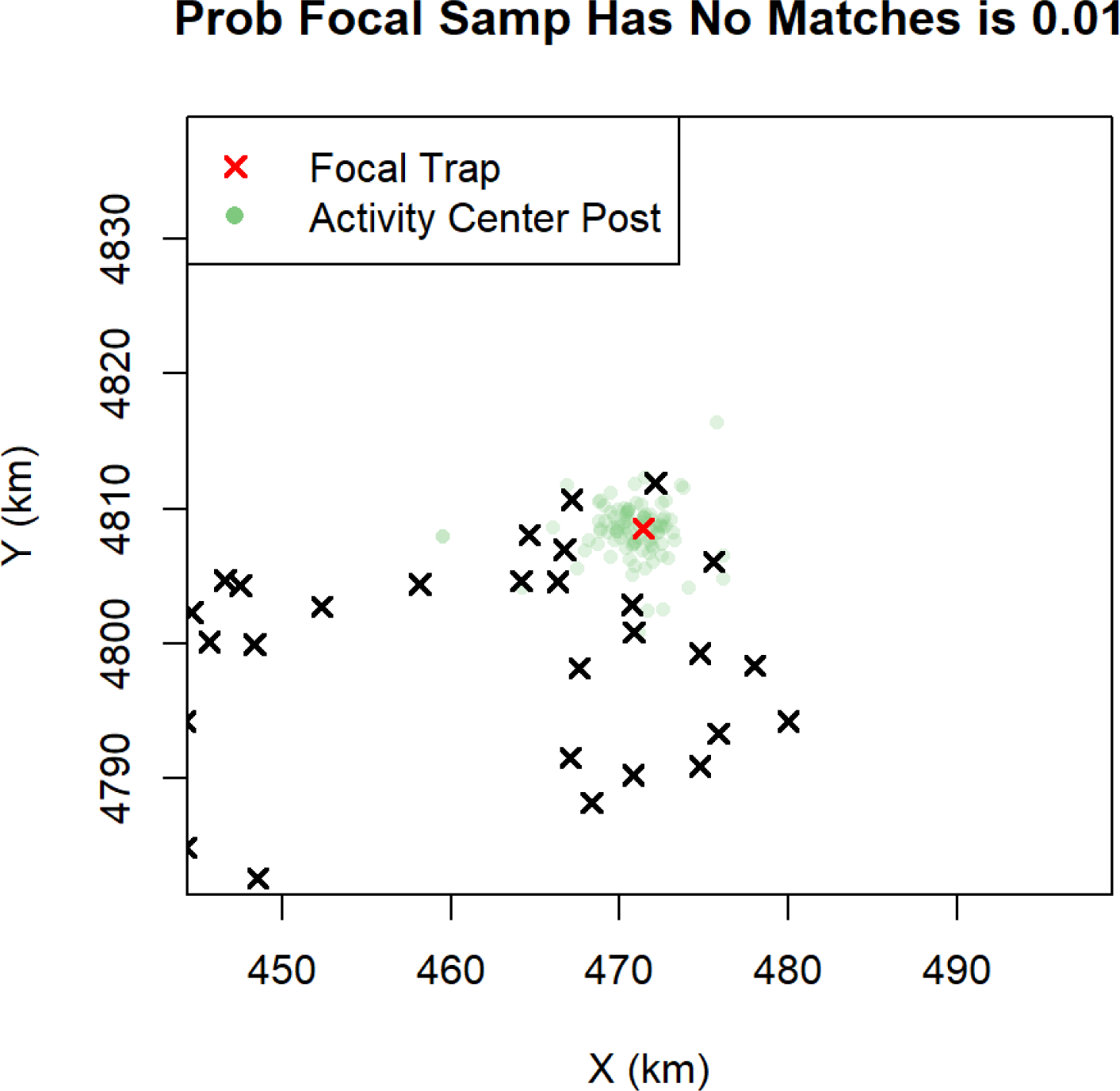

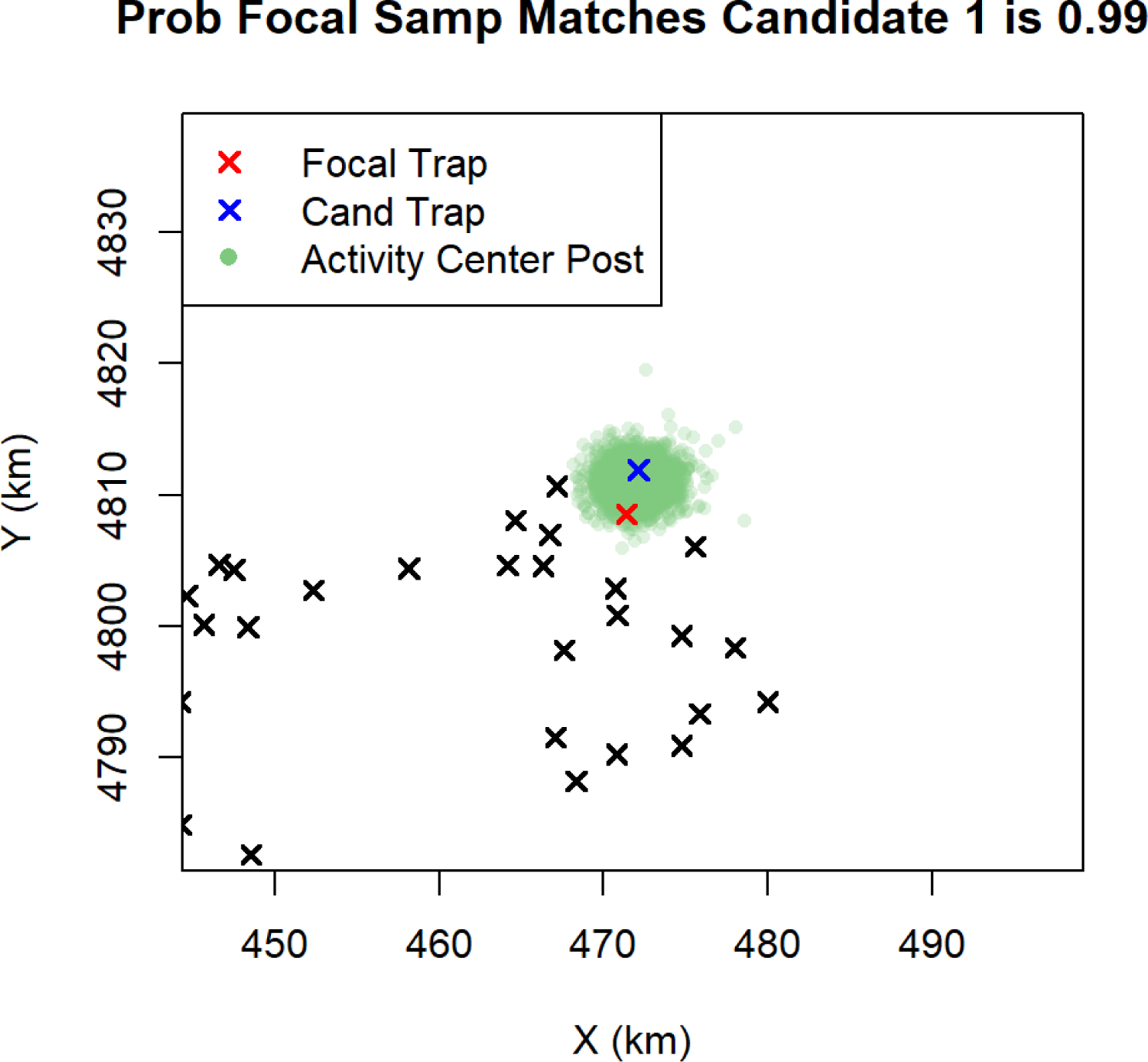

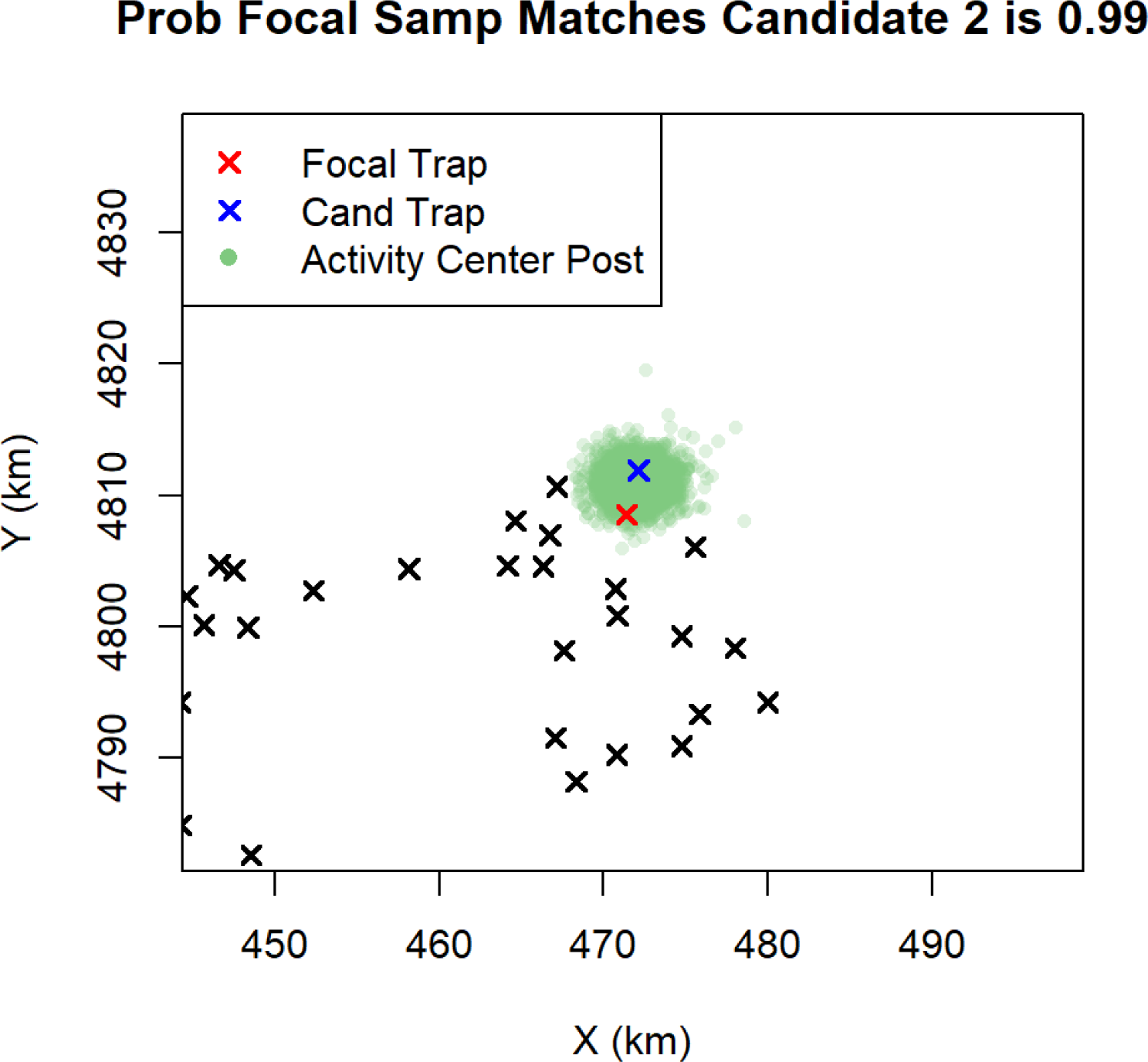

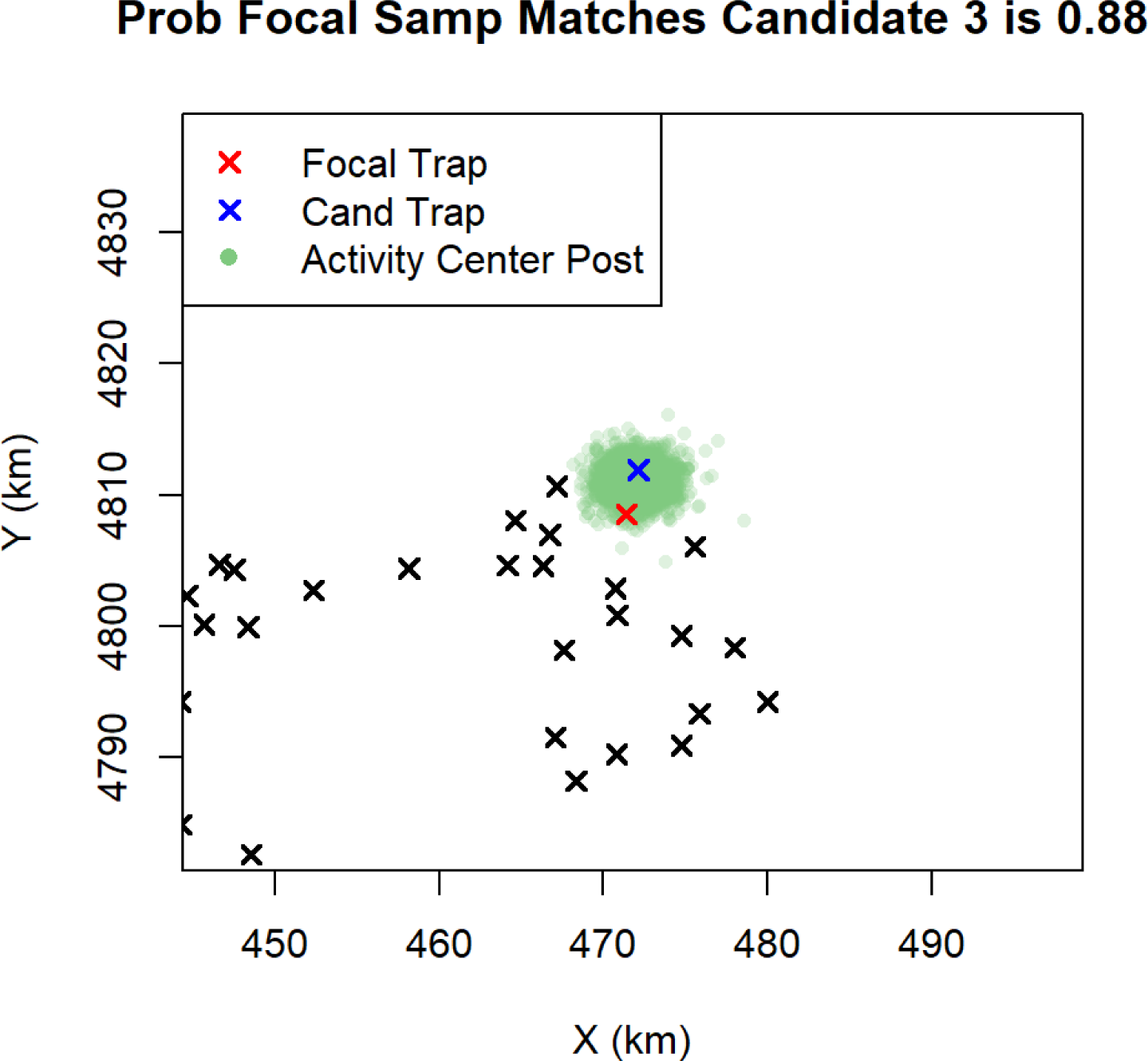

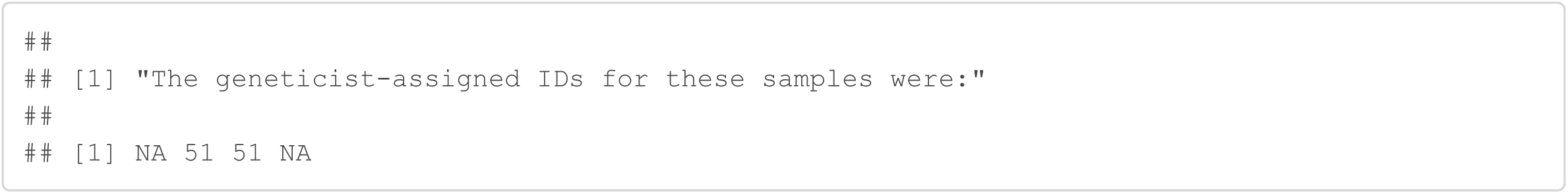

### High probability assignments that discarded samples are unique individuals with no matches to originally used samples

#### Sample 6

Sample 6 was originally discarded because it did not amplify at any loci. The Genotype SPIM gives a 0.95 probability that this sample does not match any other samples. This high probability assignment was only possible because this sample did not have any nearby neighboring samples to match with.

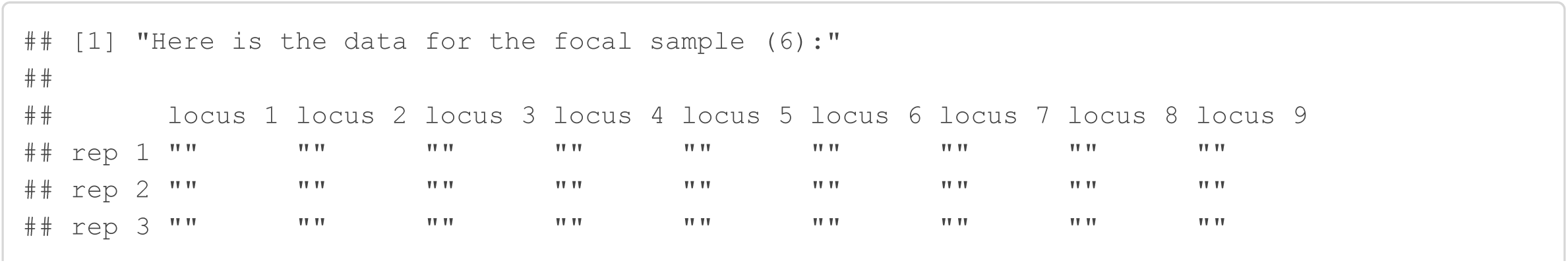

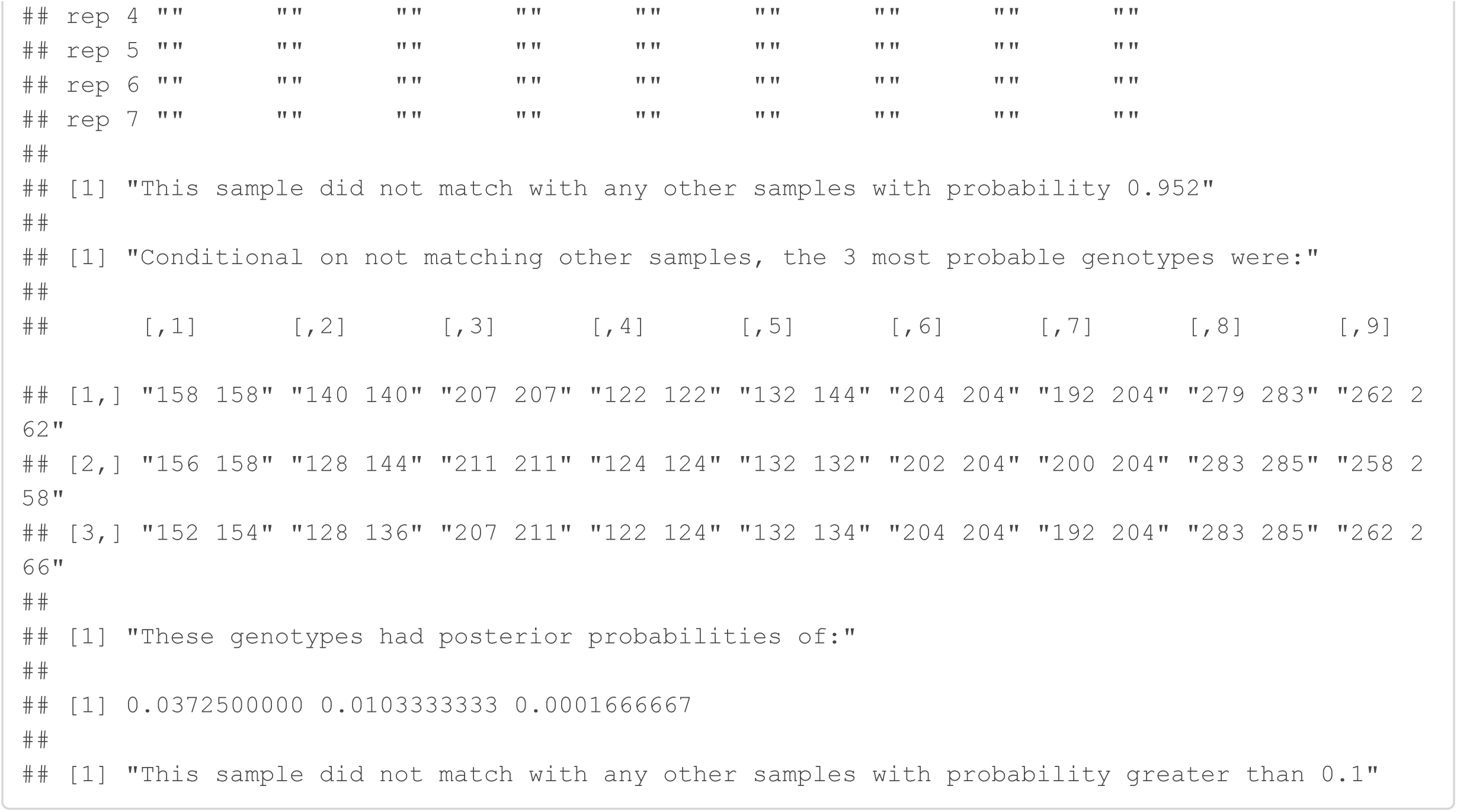

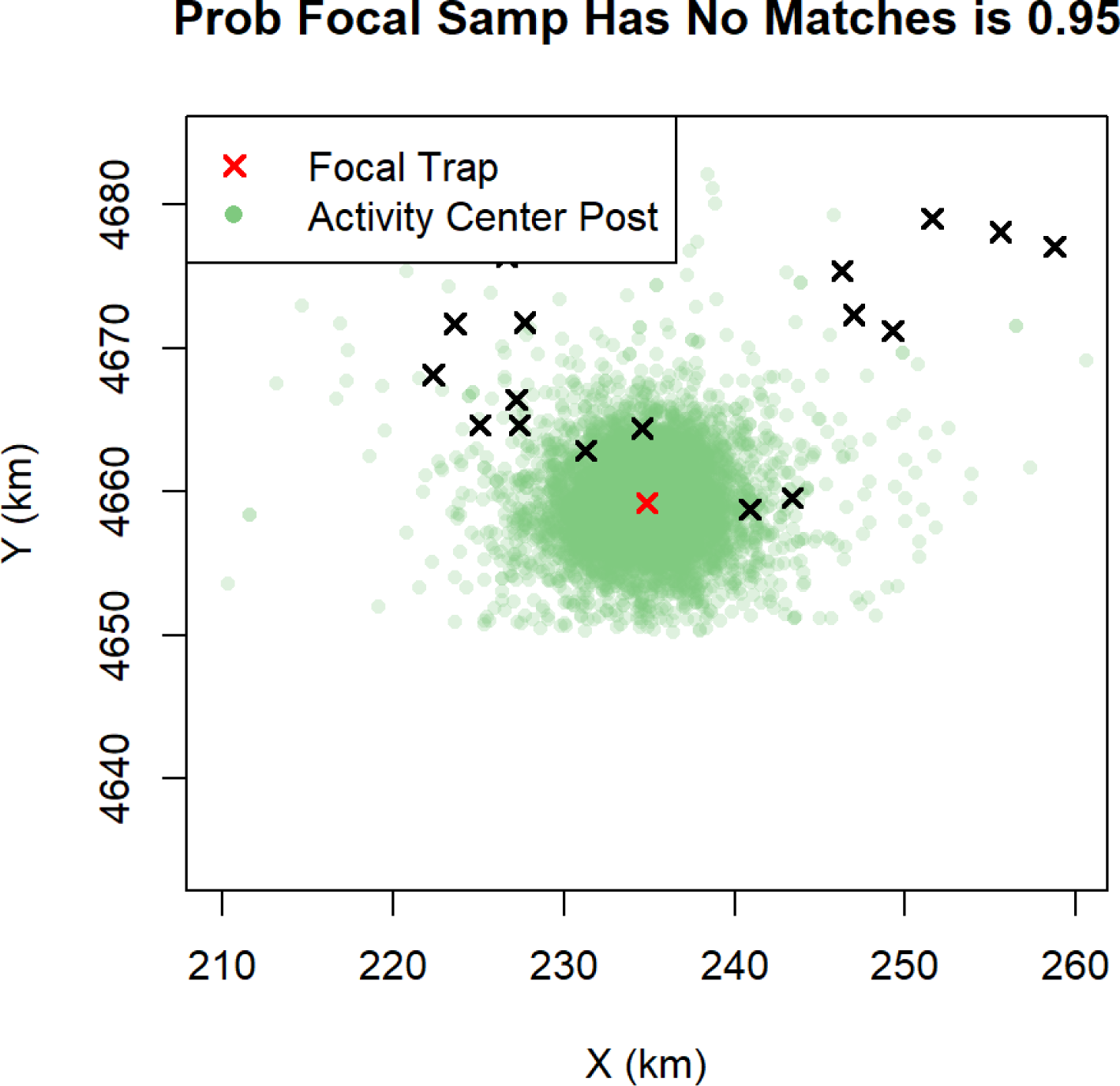

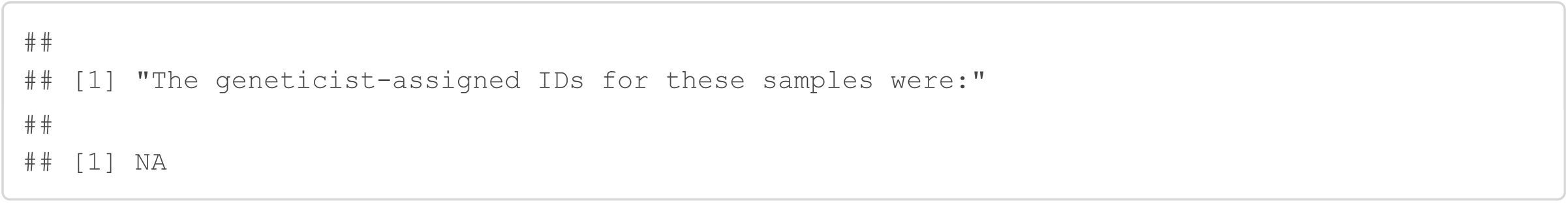

#### Sample 17

Sample 17 was originally discarded because it only amplified at 2/9 loci. The Genotype SPIM gives a 0.99 probability that this sample does not match any other samples. The rarity of genotype 213.221 at locus 3 (0.0034) and genotype 100.128 at locus 4 (0.0029) contributed to this high probability assignment of uniqueness. The genotype frequency estimates can be found in the last section of this document.

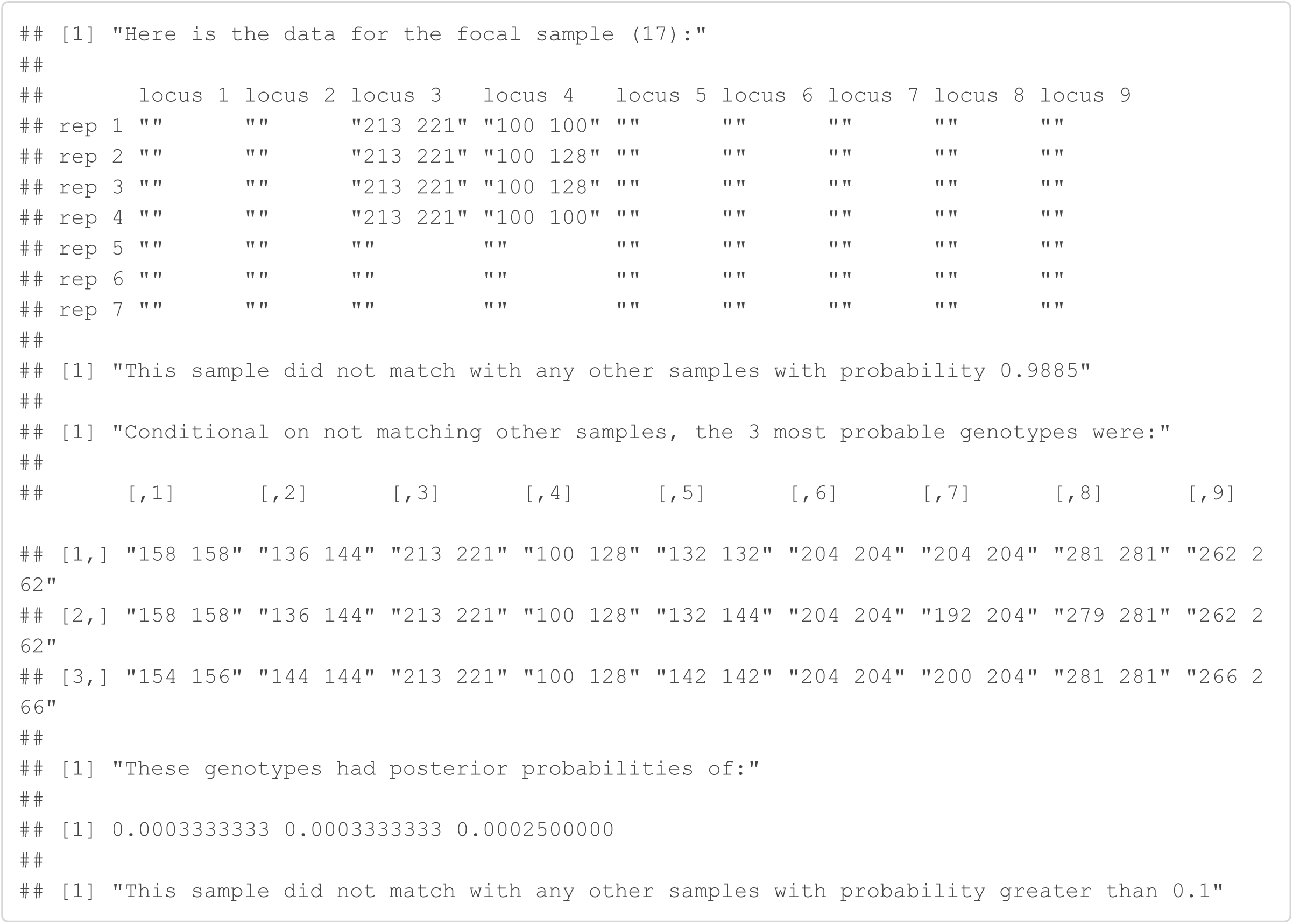

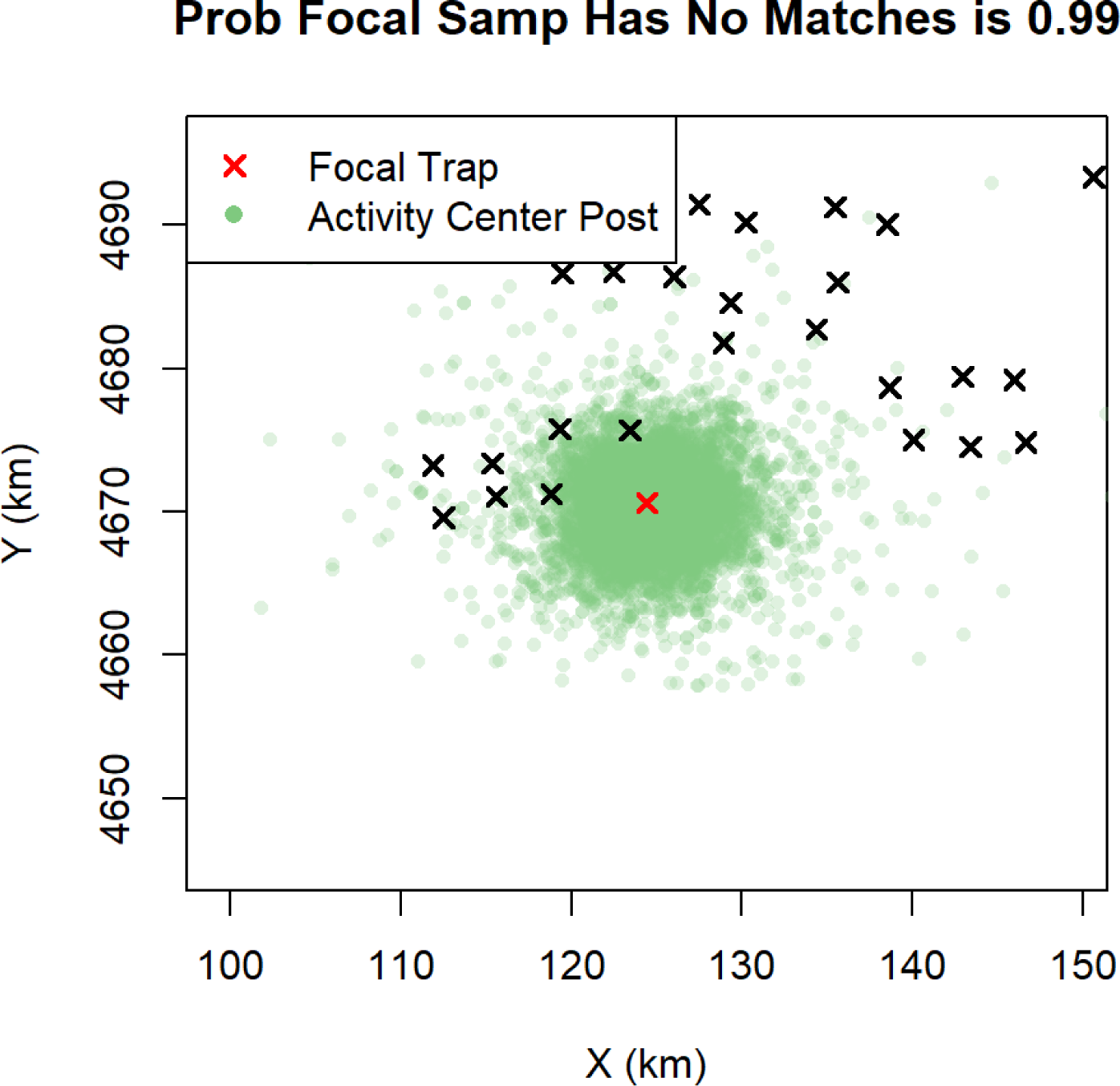

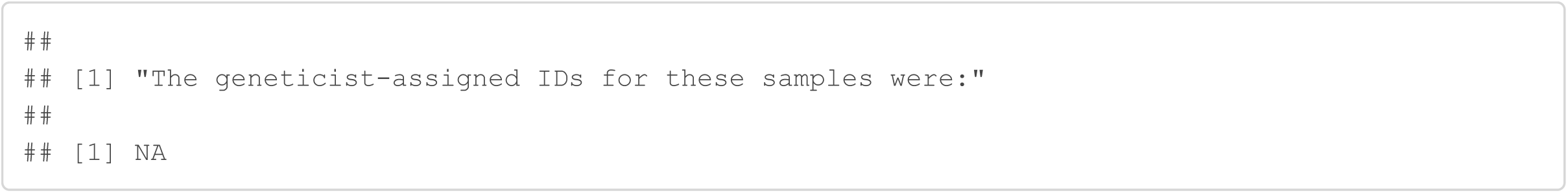

### Genotype frequency estimates

Above, the genotype frequency estimates were referenced. We display these here, with each list element storing the genotype frequency estimates for each locus.

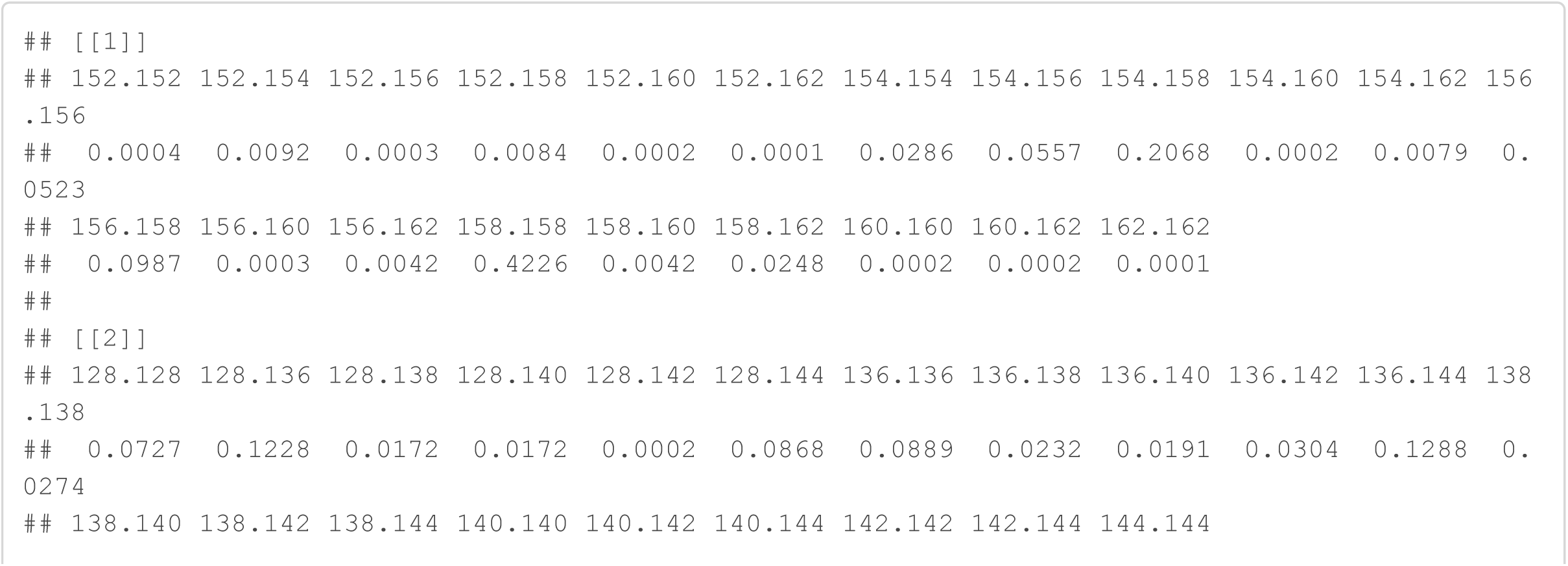

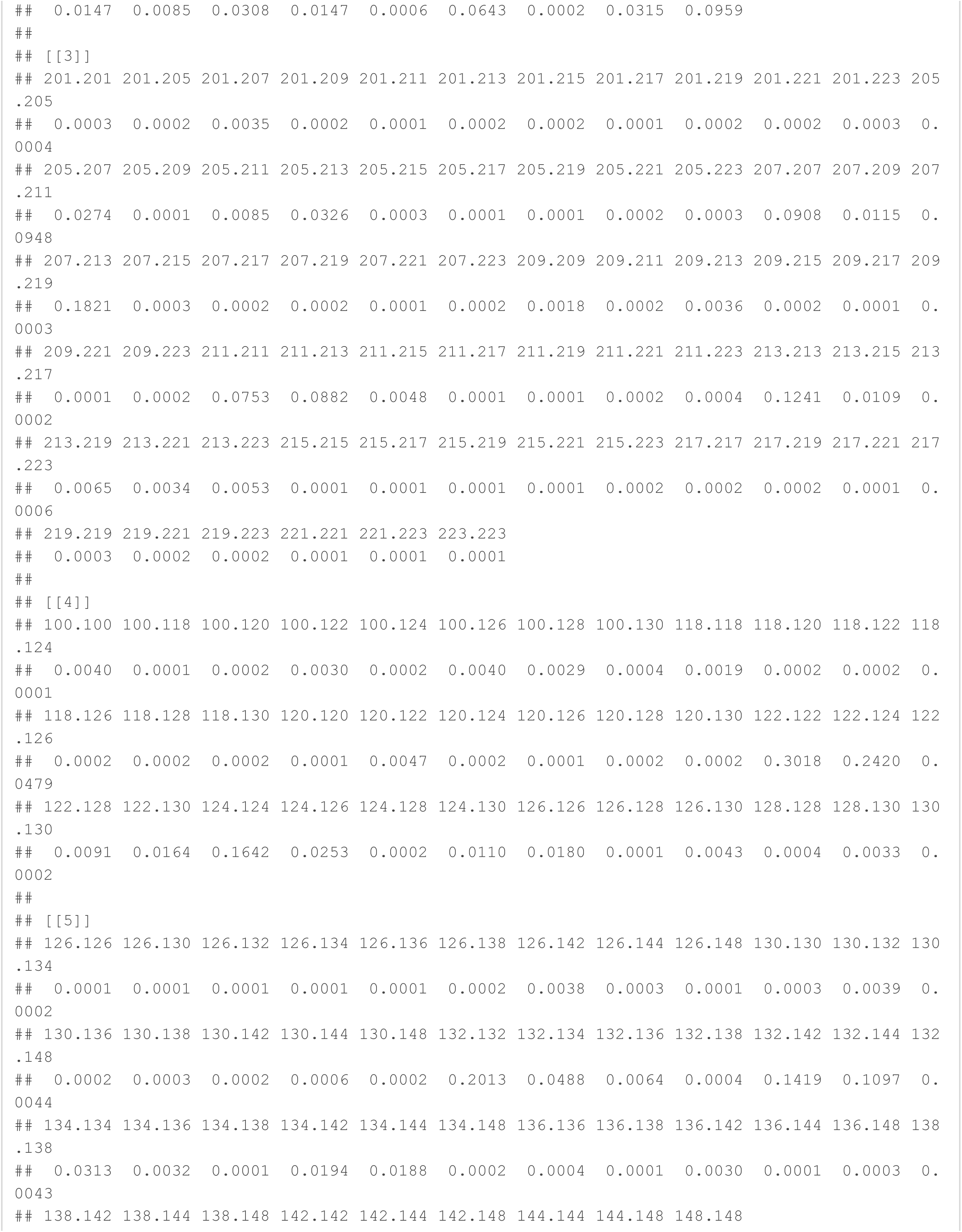

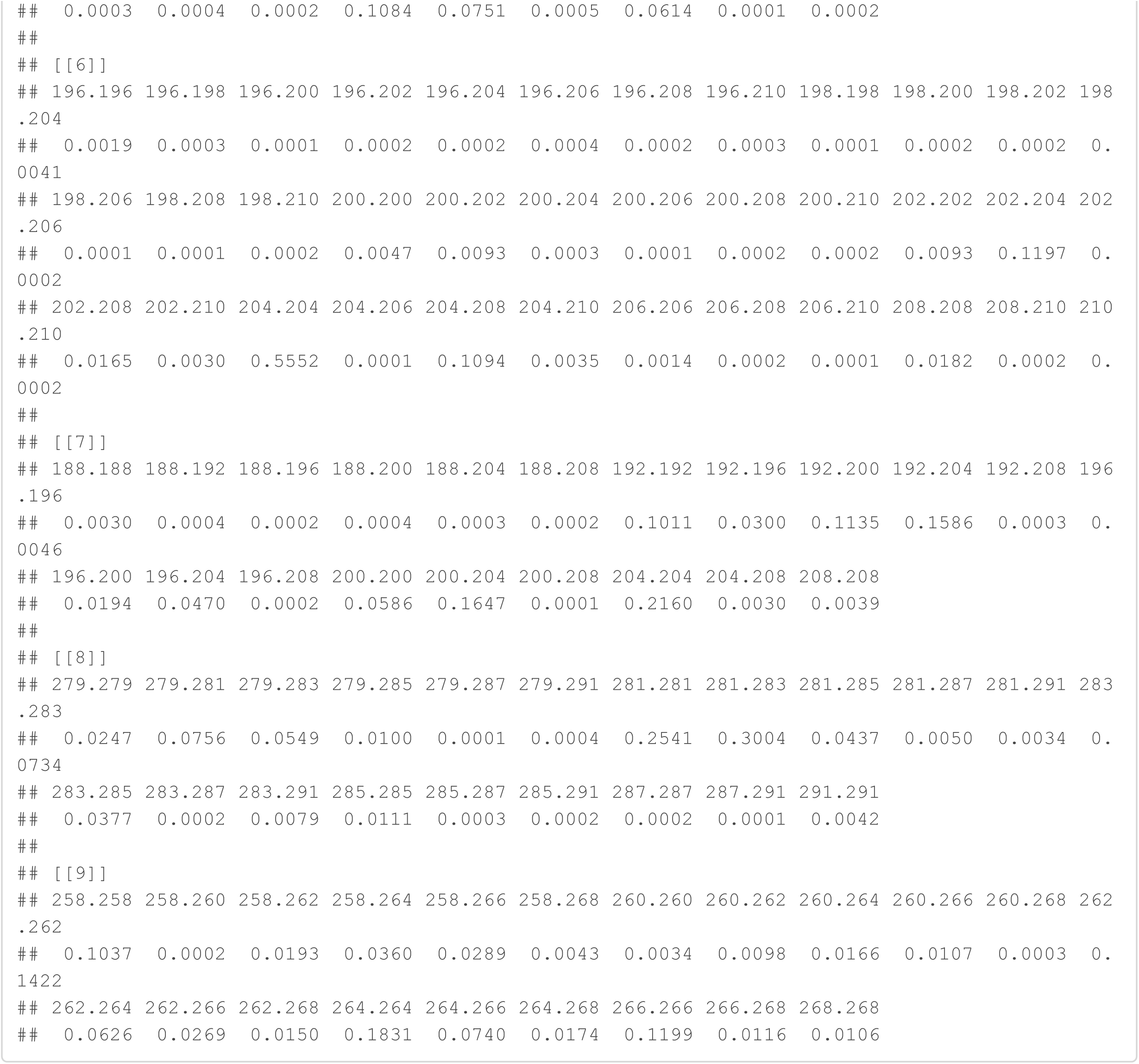

